# The choices we make and the impacts they have: Machine learning and species delimitation in North American box turtles (*Terrapene* spp.)

**DOI:** 10.1101/2020.05.19.103598

**Authors:** Bradley T. Martin, Tyler K. Chafin, Marlis R. Douglas, John S. Placyk, Roger D. Birkhead, Chris A. Phillips, Michael E. Douglas

**Affiliations:** Department of Biological Sciences, University of Arkansas, Fayetteville, Arkansas 72701, USA; Department of Biology, University of Texas, Tyler, Texas, 75799, USA; Alabama Science in Motion, Auburn University, Auburn, AL 36849, USA; Illinois Natural History Survey, Prairie Research Institute, University of Illinois, Champaign, IL 61820

**Keywords:** ddRAD, discordance, filtering, missing data, species tree, VAE

## Abstract

Model-based approaches that attempt to delimit species are hampered by computational limitations as well as the unfortunate tendency by users to disregard algorithmic assumptions. Alternatives are clearly needed, and machine-learning (M-L) is attractive in this regard as it functions without the need to explicitly define a species concept. Unfortunately, its performance will vary according to which (of several) bioinformatic parameters are invoked. Herein, we gauge the effectiveness of M-L-based species-delimitation algorithms by parsing 64 variably-filtered versions of a ddRAD-derived SNP dataset collected from North American box turtles (*Terrapene* spp.). Our filtering strategies included: (A) minor allele frequencies (MAF) of 5%, 3%, 1%, and 0% (=none), and (B) maximum missing data per-individual/per-population at 25%, 50%, 75%, and 100% (=no filtering). We found that species-delimitation via unsupervised M-L impacted the signal-to-noise ratio in our data, as well as the discordance among resolved clades. The latter may also reflect biogeographic history, gene flow, incomplete lineage sorting, or combinations thereof (as corroborated from previously observed patterns of differential introgression). Our results substantiate M-L as a viable species-delimitation method, but also demonstrate how commonly observed patterns of phylogenetic discordance can seriously impact M-L-classification.

## 1 INTRODUCTION

Species are recognized as the currency of biodiversity, yet defining what constitutes a species has been hampered by subjective interpretations. This in turn creates downstream issues for conservation (Mace 2004), where spurious ‘splitting’ or ‘lumping’ impede an equitable allocation of limited resources. Although genomic approaches based on the multispecies coalescent (MSC) are promising and have been commonly applied to the species problem (Allendorf *et al*. 2010), conflicting genome-wide signals are widely apparent due to incomplete lineage sorting (ILS) and gene flow (Funk & Omland 2003). Two MSC methods, BPP and BFD* (Yang & Rannala 2010; Leaché *et al*. 2014), seemingly over-split in the presence of strong population structure (Sukumaran & Knowles 2017) or with continuous geographic distributions (Chambers & Hillis 2019). Both are also computationally limited when applied to large datasets. As model complexity and data expand concomitantly, so also do: 1) efforts required to computationally explore appropriate parameter space; and 2) the probabilities that models fail to accommodate process. Herein, we explore alternative approaches for the parsing of high-dimensionality data by evaluating the performance of recently developed machine-learning (M-L) algorithms and classificatory approaches in successfully adjudicating variably-filtered versions of a ddRAD-derived SNP dataset.

‘Unsupervised’ machine learning methods (UML) are of particular interest for group delimitation, in that they do not require *a priori* designations to train the classification model. Several UML classifiers lend themselves to species delimitation, including: Random Forest (RF; Breiman 2001), t-distributed stochastic neighbor embedding (t-SNE; Maaten & Hinton 2008), and variational autoencoders (VAE; Kingma & Welling 2013). Each has distinct advantages: RF uses randomly replicated data subsets to develop ‘decision trees’ that are subsequently aggregated (=‘forest’), with classificatory decisions parsed as a majority vote. The random sub-setting approach is robust to correlations among features (=summary statistics or principal components used for prediction) as well as model overfitting (i.e., over-training the model such that it does not generalize to new data). One stipulation is that features must lack undue noise (Rodriguez-Galiano *et al*. 2012). By contrast, t-SNE creates clusters in reduced-dimension space, typically a 2D plane distilled from multi-dimensional data, and as such conceptually resembles principal components analysis (Maaten & Hinton 2008). On the other hand, VAE employs neural networks to ‘learn’ patterns within multidimensional data extracted from a compressed, low-dimensionality (=‘encoded’) representation. Again, an ordination technique is simulated but without imposing linear/orthogonal constraints, such that a statistically interpretable result emerges that is appropriate for highly-complex data (Derkarabetian *et al*. 2019).

Some algorithms are robust to gene flow (Derkarabetian *et al*. 2019; Newton *et al*. 2020; Smith & Carstens 2020), yet a greater number of tests must be performed across diverse systems so as to understand which parameters impinge upon performance. Potentials include: Data quantity (Newton *et al*. 2020), the proportion of missing data (Mussmann *et al*. 2020), and evolutionary complexity (Austerlitz *et al*. 2009). Here, we employ M-L algorithms alongside coalescent methods such as BFD* (Leaché *et al*. 2014) as vehicles to parse a taxonomically recalcitrant clade. Included algorithms are: Process-based RF (delimitR; Smith *et al*. 2017; Smith & Carstens 2020) and unsupervised RF, t-SNE, and VAE, as implemented in Derkarabetian *et al*. (2019).

### 1.1 Species concepts and their evolution in *Terrapene*

North American box turtles (Emydidae: *Terrapene*) are a primarily terrestrial group that includes five currently recognized species (Minx 1996; Iverson *et al*. 2017): Eastern (*Terrapene carolina*), Ornate (*T. ornata*), Florida (*T. bauri*), Coahuilan (*T. coahuila*), and Spotted (*T. nelsoni*), with a sixth (*T. mexicana*) proposed (Martin *et al*. 2013). *Terrapene carolina* is split into two subspecies east of the Mississippi River and south through the Gulf Coast [Woodland (*T. c. carolina*) and Gulf Coast (*T. c. major*); Figure 1]. *Terrapene mexicana* contains three subspecies: Three-toed (*T. m. triunguis*); Mexican (*T. m. mexicana*); and Yucatan (*T. m. yucatana*) that range across southeastern and midwestern United States, the Mexican state of Tamaulipas, and the Yucatan Peninsula. Ornate (*T. ornata ornata*) and Desert (*T. o. luteola*) inhabit the Midwest and Southwest U.S. and Northwest México, while Southern and Northern Spotted box turtles (*T. nelsoni nelsoni* and *T. n. klauberi*) occupy the Sonoran Desert in western México. *Terrapene coahuila* is semi-aquatic and restricted to Cuatro Ciénegas (Coahuila, México), while Florida box turtle occurs in Peninsular Florida.

**FIGURE 1.**
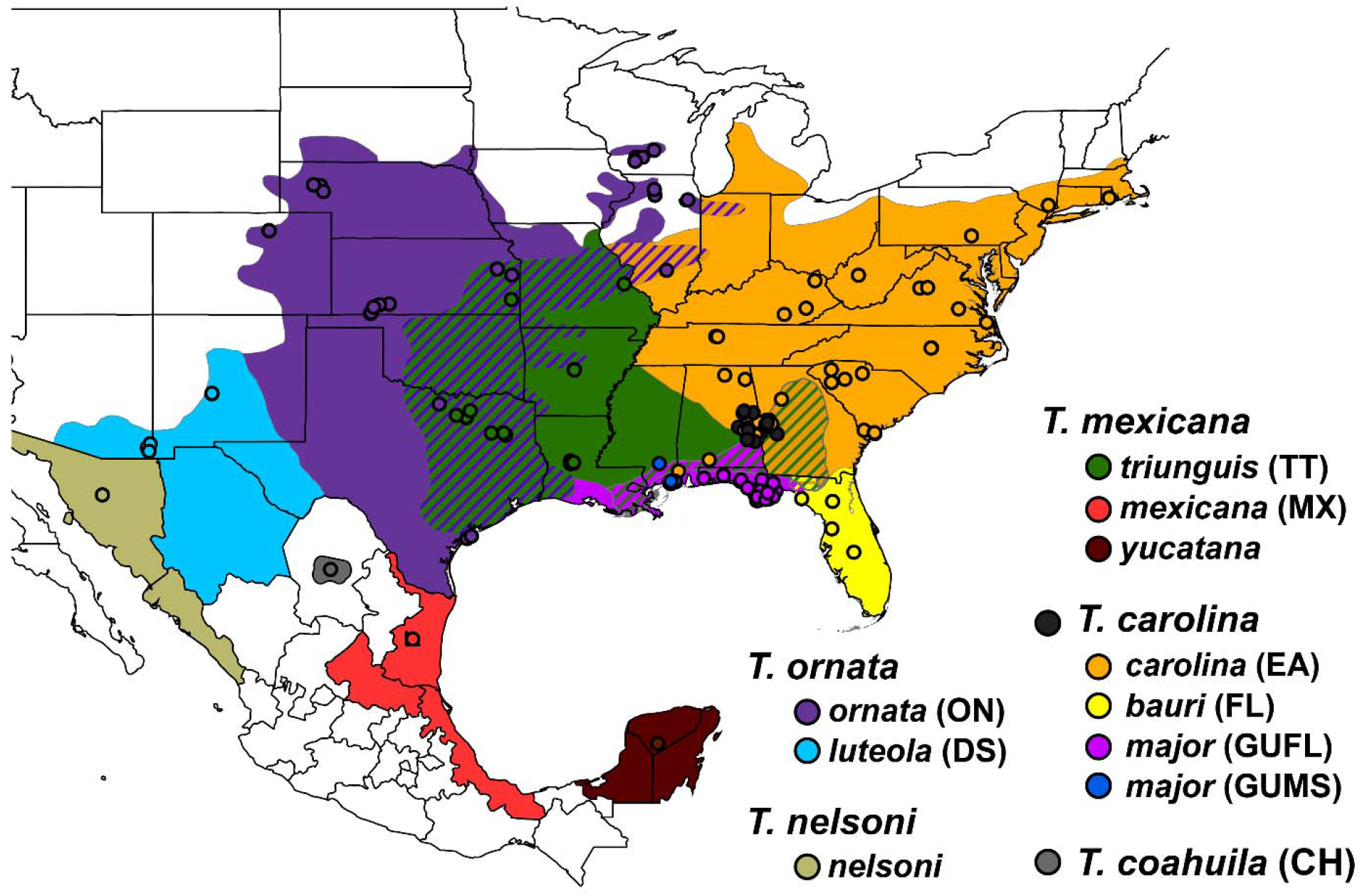
Range map and sample localities (=circles) for N*=*214 *Terrapene*. Closed circles=*T. carolina* samples without subspecific identification in the field. Cross-hatched areas=known hybrid zones. Headings and subheadings represent species and subspecies. *Terrapene carolina major*=*T. carolina major* and includes distinct subpopulations from Mississippi (GUMS) and Florida panhandle (GUFL). Parenthetical legend abbreviations correspond to Tables 2 and 3.

Morphological analyses delineate *T. carolina*/*mexicana* as a single species, sister to *T. coahuila* (Minx 1992, 1996), as supported by genetic studies (Feldman & Parham 2002; Stephens & Wiens 2003). Martin et al. (2013) elevated *T. mexicana*, and nested *T. coahuila* within *T. carolina. Terrapene carolina carolina* is sister to *T. c. major*/*T. coahuila*, although gene flow was suspected with *T. c. major. Terrapene carolina major* was recently demoted to an intergrade with subsequent loss of subspecific status (Butler *et al*. 2011; Iverson *et al*. 2017). However a recent genomic study supported pure *T. c. major* populations in Florida and Mississippi (Martin *et al*. 2020). Similarly, *T. bauri* (formerly *T. carolina bauri*) was recently elevated (Butler *et al*. 2011; Iverson *et al*. 2017), but more substantial evidence is needed (Martin *et al*. 2013). For clarity, we retain the nomenclature of Martin *et al*. (2013, 2014), with *T. c. major* and *bauri* representing *T. carolina* subspecies.

**TABLE 1.**
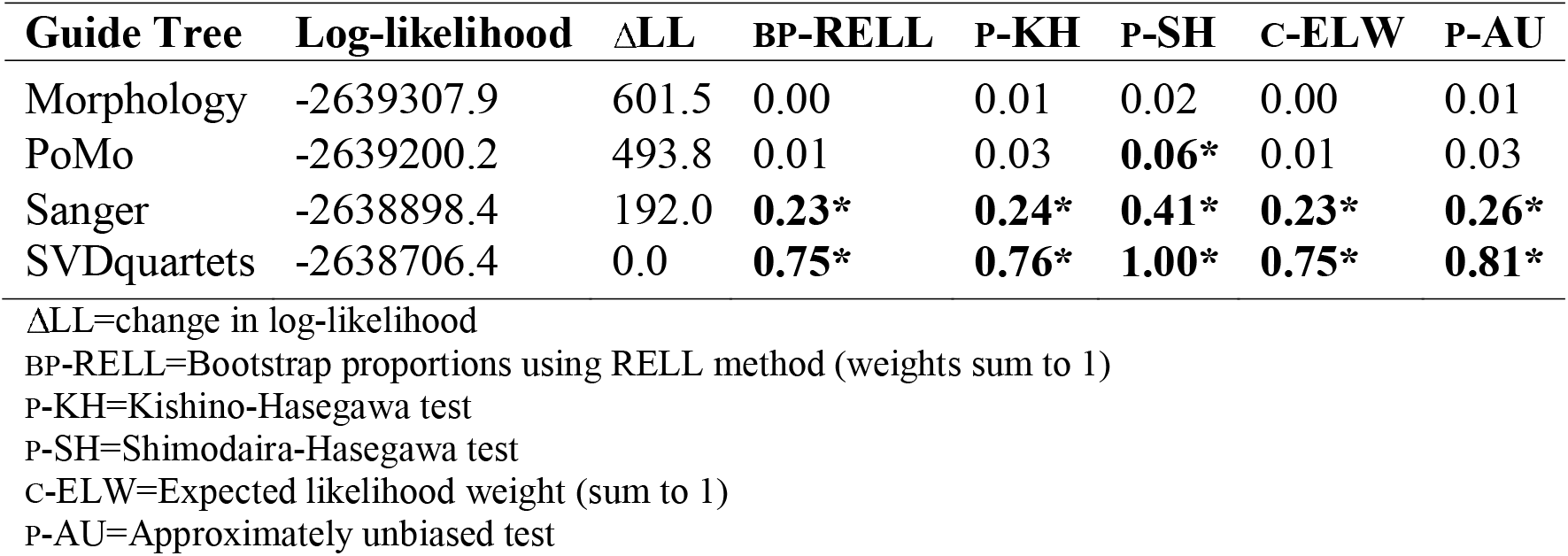
Topology tests for hypothesized *Terrapene* phylogenies. Sanger sequencing and morphology trees are based on previously published data whereas those representing SVDquartets and PoMo (Polymorphism-Aware Model) were generated in this study from ddRADseq data. *P*-values in bold with ‘*’ indicate significance (P>0.05/highly weighted).

**TABLE 2.** Species-delimitation results from Bayes Factor Delimitation (BFD) in *Terrapene*. Bayes factors (BF) depict support among models and were calculated as 2 × (MLE_1_-MLE_2_). ‘*’=best supported models; ‘+’=taxa grouped together; ‘/’=multiple groupings. DS=*T. o. luteola*, ON=*T. o. ornata*, EA=*T. c. carolina*, GUFL=*T. c. major* from Florida, GUMS=Mississippi *T. c. major*, CH=*T. coahuila*, FL=*T. c. bauri*, TT=*T. m. triunguis*, and MX=*T. m. mexicana*. East=all *T. carolina* and *T. mexicana*, West=all *T. ornata*. Outgroup (not shown) included *Clemmys guttata*.

| BFD* Model | MLE† | K‡ | Rank§ | BF¶ |
| --- | --- | --- | --- | --- |
| All Separate* | -2403.39 | 10 | 1 | - |
| DS+ON* | -2404.34 | 9 | 2 | 1.90 |
| EA+GUFL | -2417.84 | 9 | 3 | 28.91 |
| GUMS+GUFL | -2427.58 | 9 | 4 | 48.39 |
| GUMS+CH | -2448.61 | 9 | 5 | 90.44 |
| GUMS+CH/GUFL+EA | -2461.28 | 8 | 6 | 115.79 |
| GUMS+GUFL+CH | -2489.62 | 8 | 7 | 172.45 |
| EA+FL | -2511.83 | 9 | 8 | 216.89 |
| GUMS+GUFL+CH+EA | -2514.86 | 7 | 9 | 222.94 |
| EA+FL+GUFL | -2552.22 | 8 | 10 | 297.66 |
| EA+FL/CH+GUMS | -2555.16 | 8 | 11 | 303.53 |
| EA+FL+GUFL/CH+GUMS | -2594.91 | 7 | 12 | 383.04 |
| EA+CH+GUMS+GUFL+TT | -2607.72 | 6 | 13 | 408.66 |
| EA+CH+GUMS+GUFL+MX | -2657.48 | 6 | 14 | 508.19 |
| EA+FL+CH+GUMS+GUFL | -2693.37 | 6 | 15 | 579.96 |
| EA+CH+GUMS+GUFL+TT+MX | -2719.02 | 5 | 16 | 631.27 |
| ON+DS/EA+TT+MX+CH+GUMS+GUFL/FL | -2720.23 | 4 | 17 | 633.69 |
| EA+FL+CH+GUMS+GUFL+TT | -2800.56 | 5 | 18 | 794.35 |
| EA+FL+CH+GUMS+GUFL+TT+MX | -2926.20 | 4 | 19 | 1045.62 |
| East/West | -2926.56 | 3 | 20 | 1046.35 |

†MLE=Marginal likelihood estimates
‡K=# tips
§Rank=model ranking based on MLE (lower=better)
¶BF=Bayes factors

**TABLE 3.** The top five (of 51) delimitR models describing six *Terrapene* taxa. Model=rank determined by random forest (RF) vote counts (=# Votes). ‘*’=best supported model. Grouped taxa are separated by ‘+’, whereas ‘/’=distinct groups. ‘×’ separates migration events promoting secondary contact, with multiple migrations per model separated by commas. ON=*T. o. ornata*, TT=*T. m. triunguis*, FL=*T. c. bauri*, GUMS=*T. c. major* from Mississippi, GUFL=Florida *T. c. major*, EA=*T. c. carolina*. Error=proportion of incorrect model choices.

| Model | # Votes | Species (# delimited) | Secondary Contact | Error |
| --- | --- | --- | --- | --- |
| 17* | 464 | ON/TT/FL/GUMS+GUFL+EA (4) | ON × TT, TT × GU+EA, FL × GU+EA | 0.017 |
| 14 | 445 | ON/TT/FL/GUMS+GUFL+EA (4) | TT × GU+EA, FL × GU+EA | 0.036 |
| 3 | 441 | ON/TT+FL+GUMS+GUFL+EA (2) | ON × TT+FL+GU+EA | 0.009 |
| 8 | 359 | ON/TT/FL+GUMS+GUFL+EA (3) | ON × TT, TT × FL+GU+EA | 0.009 |
| 30 | 218 | ON/TT/FL/GUMS+GUFL/EA (5) | TT × GU, FL × EA, GU × EA | 0.007 |

One explanation for the enigmatic classification of *T. carolina* and *T. mexicana* involves hybridization (Auffenberg 1958, 1959; Milstead & Tinkle 1967; Milstead 1969). Some researchers (Fritz & Havaš 2013, 2014) interpreted reproductive semi-permeability as justification sufficient to collapse the southeastern taxa. However, their classificatory status must be re-examined, as indicated by results modulating the species boundaries of southeastern *Terrapene* (Martin *et al*. 2020).

Taxonomic disputes in *Terrapene* highlight the philosophical disparity among species definitions [e.g., biological (Mayr 1963) versus phylogenetic (Eldredge & Cracraft 1980)]. The approach advocated herein acknowledges that operational criteria among concepts are intimately related. Specifically, reproductive barriers (through time) beget genealogical concordance, while contemporary evaluations of gene flow are contextualized via phylogenetic/phylogeographic perspectives (Avise 2000a; b). We thus subscribe to a ‘unified species concept’ (De Queiroz 2007) wherein the primary criterion for formal taxonomic rank is the existence of evolutionary lineages (e.g., as distinct metapopulations), with evidence via reproductive isolation, phylogenetic-phylogeographic resolution, and phenotypic adaptation, with all acknowledged as being inherently linked. Here, our clustering and classificatory approaches define molecular diagnosability, and as such variably place *Terrapene* lineages along a speciation continuum (Via 2009; Nosil & Feder 2012; Edwards *et al*. 2016; Martin *et al*. 2020).

## 2 MATERIALS AND METHODS

### 2.1 DNA extraction and library preparation

Tissue samples were obtained from museums, agencies, and volunteers (Supplementary Information Table S1) and stored at-20°C. Genomic DNA was extracted via spin-column kits: DNeasy Blood and Tissue (QIAGEN), QIAamp Fast DNA (QIAGEN), and E.Z.N.A. Tissue DNA Kits (Omega Bio-tek). Extracted DNA was quantified using Qubit fluorometry (Thermo Fisher Scientific), and characterized using gel electrophoresis on 2% agarose.

Samples were processed via ddRADseq (Peterson et al. 2012), with ∼500-1,000ng of genomic DNA/sample digested with *PstI* and *MspI* at 37°C for 24 hours. Samples were bead-purified (Beckman-Coulter) at 1.5X concentration then standardized at 100ng. Barcoded adapters were ligated before pooling 48 samples per library. Taxa were spread across libraries to mitigate batch effects then size-selected (454-509 bp, including ligated adapters) on a Pippin Prep (Sage Science). Adapter-extension was performed via twelve-cycle PCR, followed by 1×100 sequencing on the Illumina Hi-Seq 4000 (University of Oregon/Eugene), with two indexed libraries pooled/lane.

### 2.2 Quality control and assembly

FastQCv.0.11.5 was used to assess sequence quality (Andrews 2010), with raw reads demultiplexed via ipyrad v.0.7.28 (Eaton & Overcast 2020), allowing for one barcode mismatch as a maximum. Low quality sequences (>5 bases with Q<33) and adapters were removed. Assembly was reference-guided using *Terrapene mexicana* (GCA_002925995.2), with unmapped reads discarded. To reduce error, only loci exhibiting ≥20X coverage were retained (Nielsen *et al*. 2011). We also excluded loci with excessive heterozygosity (≥75% of individual SNPs), <50% global occupancy, or >two alleles/sample.

### 2.3 Phylogenomic inferences

F_1_ and F_2_-generation hybrids previously identified in a population-level analysis (Martin *et al*. 2020) were excluded as a means of mitigating impacts of contemporary gene flow on species tree inference (Long & Kubatko 2018). We then employed SVDquartets (Chifman & Kubatko 2014) filtered to one SNP per locus to reduce linkage bias, with exhaustive quartet sampling and 100 bootstrap pseudo-replicates. Taxon partitions were grouped by subspecies and U.S./Mexican state locality, with *Emydoidea blandingii* and *Clemmys guttata* as outgroups.

We also employed a polymorphism-aware model (PoMo: Schrempf *et al*. 2016), as implemented in IQ-TREE v1.6.9 (Nguyen *et al*. 2015), with full-locus alignments and 1,000 ultrafast bootstrap (UFBoot) replicates (Hoang *et al*. 2017). The maximum virtual population size was 19, with discrete gamma-distributed rates=4.

Using ten-thousand re-samplings, we performed topology tests (IQ-TREE) with seven statistical criteria on the SVDquartets and PoMo trees, as well as a previously published morphological (Minx 1996) and a molecular hypothesis (Martin *et al*. 2013). Additional details are in Supplementary Information Appendix A.1.1.

A lineage tree was generated (IQ-TREE v2.0.6; Minh *et al*. 2020) and full-locus partitions merged (Chernomor *et al*. 2016), with the top 10% of combinations employed and a per-partition model search (ModelFinder: Kalyaanamoorthy *et al*. 2017). Node support was assessed using 1,000 UFBoot replicates and site-wise concordance factors (sCF; Minh *et al*. 2018). The sCF values were calculated from 10,000 randomly sampled quartets.

### 2.4 Divergence dating

A full concatenation tree was time-calibrated via least square dating (LSD2), as implemented in IQ-TREE (To *et al*. 2016). Four fossil calibration points were used (Holman & Fritz 2005; Spinks & Shaffer 2009), including the following most recent common ancestors (MRCAs): (1) *T. ornata* and *T. carolina/T. mexicana*, minimally constrained to 13 million years ago (Mya); (2) *T. o. ornata* and *T. o. luteola* (9.0-13.0 Mya); (3) *T. carolina* and *T. mexicana* (9.0-11.0 Mya); and (4) *Terrapene* and *Clemmys*/*Emydoidea* [(maximally constrained to 29.4 Mya) (per Martin *et al*. 2013)]. Branch lengths were simulated from a Poisson distribution with 1,000 replicates to assess 95% confidence intervals.

### 2.5 Species delimitation using BFD*

We employed Bayes Factor Delimitation (BFD*; Leaché *et al*. 2014) as a comparative baseline. Given its computationally-intense process, each taxon was subset to a maximum of five individuals containing the least missing data (N=37+outgroups). Sites with >50% missing data in any population were removed (see Supplementary Information Appendix A.2.1 for prior selection and data formatting steps for BFD*).

For each BFD* model, we used 48 path-sampling steps, 200,000 burn-in, plus 400,000 MCMC iterations, sampling every 1,000 generations. Path-sampling was conducted with 200,000 burn-in+300,000 MCMC generations, α=0.3, 10 cross-validation replicates, and 100 repeats. Trace plots were visualized in Tracer v1.7.1 to evaluate parameter convergence and compute effective sample sizes (ESS; Rambaut *et al*. 2018). Bayes factors (BF) were calculated from normalized likelihood estimates (MLE) as [2 × (MLE_1_-MLE_2_)]. We considered the following scheme for model support: 0<BF<2=no differentiation; 2<BF<6=positive; 6<BF<10=strong; and BF>10=decisive support (Kass & Raftery 1995).

### 2.6 Preparing and executing UML datasets

To assess the influence of bioinformatic choices on M-L species delimitation, we performed missing data filtering sweeps to produce 64 datasets across three filtering options. Missing data was filtered per-individual and per-population, with the maximum permitted occupancy set to 25%, 50%, 75%, and no filtering (=100%). Datasets were also filtered by minor allele frequency (MAF) at values of 5%, 3%, 1%, and 0% (=no MAF filter). Custom scripts were employed for all filtering steps (https://github.com/tkchafin/scripts).

RF and t-SNE (Breiman 2001; Maaten & Hinton 2008) were executed and visualized using an R script [Derkarabetian *et al*. (2019); https://github.com/shahanderkarabetian/uml_species_delim]. We ran 100 replicates for each of the 64 datasets, with data subsequently represented as scaled principal components (Adegenetv2.1.1; Jombart & Ahmed 2011) in Rv3.5.1 (R Development Core Team 2018). To generate RF predictions, we averaged 10,000 majority-vote decision trees. Clustered RF output was visualized using both classic and isotonic multidimensional scaling (cMDS and isoMDS; Shepard *et al*. 1972; Kruskal & Wish 1978). We ran t-SNE for 20,000 iterations, with equilibria of the clusters visually observed. Perplexity, which limits the effective number of t-SNE neighbors, was subjected to a grid search with values from 5-50, incremented by five.

VAE (Derkarabetian *et al*. 2019) employs neural networks to infer the marginal likelihood distribution of sample means (μ) and standard deviations [(s) (i.e. ‘latent variables’)]. As with RF and t-SNE analyses, VAE was also run with 100 replicates to assess cluster stochasticity. Each of the 64 datasets were split into 80% training/20% validation datasets using the *train_test_split* module (*scikit-learn*: Pedregosa *et al*. 2011), with model loss (∼error) visualized to determine the optimal number of ‘epochs’ (=cycles through the training dataset). VAE should ideally be terminated when loss converges on a minimal difference between training and validation datasets [the ‘Goldilocks zone’; Supplementary Information Figure S1 (Al’Aref *et al*. 2019)].

Overfitting is indicated when model loss in the validation dataset escalates, whereas underfitting is a failure to reach minimum points (=inability to generalize to unseen data). Thus, we added minor modifications to the original Python script (Derkarabetian *et al*. 2019) by implementing an early stopping callback (*keras*.*callbacks* Python module; Chollet 2015), which terminates training when model loss fails to improve for 50 epochs, then restores the best model prior to the tolerance period (see Supplementary Information Appendix A.2).

### 2.7 K-selection for RF, t-SNE, and VAE

Two clustering algorithms (R-scripts: Derkarabetian *et al*. 2019), were used to identify clusters and derive optimal *K* for RF and t-SNE analyses. The first [Partitioning Around Medoids (PAM); Kaufman and Rousseeuw 1987] minimizes the distance of intra-cluster points to a centroid. The program requires *K* to be defined *a priori*, and thus *K*=1-10 were tested. The second (hierarchical clustering, HC; Fraley & Raftery 1998) iteratively merges points with minimal dissimilarity. After clustering, optimal *K* was chosen using the gap statistic (GS) and highest mean silhouette width [HMSW; Rousseeuw (1987), Tibshirani *et al*. (2001)].

VAE used DBSCAN (Ester *et al*. 1996), as implemented in a custom Python script (*vae_dbscan*.*py*), to derive clusters using a distance threshold (ε) rather than *a priori* setting of *K*. Here we used 2 × the standard deviation, but averaged globally across all samples (following Derkarabetian *et al*. 2019).

For plotting, we implemented a permutation-based heuristic search to align *K* across all replicates and the 64 datasets [‘Cluster Markov Packager Across K;’ Kopelman *et al*. (2015) implemented in PopHelper (Francis 2017)]. Assignment probabilities were then visualized as stacked bar plots for each method (via a custom script: *plotUML_missData_maf*.*R*). For each dataset, we plotted as heatmaps the optimal *K* and standard deviation (SD) among replicates [(*plot_missData_comparison_maf*.*R*) (Scripts deposited at: https://github.com/btmartin721/mecr_boxturtle)].

### 2.8 Demography, migration history, and species-delimitation

We tested for reticulation in our phylogenomic dataset, as complementary to a range-wide evaluation of introgression in *Terrapene* (Martin *et al*. 2020). We first explored reticulation by identifying candidate edges (TreeMix; Pickrell & Pritchard 2012), with populations having but one sample (*T. nelsoni* and *T. m. yucatana*) being excluded from input, which was then thinned to bi-allelic SNPs. TreeMix was run 10X with subsets of SNPs randomly sampled per locus at 1,000 bootstrap replicates using the ‘global search’ option. The optimal number of admixture edges (*m*) was determined by running for *m*=1-10 and choosing the inflection point of log-likelihood scores.

TreeMix results and introgression (Martin *et al*. 2020) were used to generate gene flow hypotheses in a species-delimitation framework (delimitR: Smith *et al*. 2017; Smith & Carstens 2020). delimitR uses the joint site-frequency spectrum (jSFS) and fastsimcoalv2.6 (Excoffier *et al*. 2013) to simulate demographic models, including possible variations of lumping/splitting taxa and primary divergence, secondary contact, or no gene flow. The program then builds an RF-classifier trained with the simulated models (i.e., ‘supervised’ M-L) to predict the best model. Input was generated using easySFS (https://github.com/isaacovercast/easySFS), with taxa reduced to N=6 given computational resources required by larger datasets. Those excluded (*T. m. mexicana, T. m. yucatana, T. o. luteola, T. coahuila, T. nelsoni*) were either limited in sample size or had clear taxonomic identities in the other analyses.

To improve efficiency, we also used easySFS to down-project the jSFS to six alleles for *T. c. bauri*, and ten each for the remaining taxa. Samples were selected to maximize per-individual occupancy, followed by a maximum 50% per-population missing data filter. The SVDquartets result served as our topological prior for delimitR. Models considered were: No gene flow, primary divergence, secondary contact, and up to four migration edges. Migration was permitted between: *T. c. carolina* x *T. c. major, T. c. carolina* x *T. c. bauri, T. c. major* x *T. m. triunguis*, and *T. m. triunguis* x *T. o. ornata*. Population size priors were set broadly (1,000-100,000) and divergence times were obtained from LSD2 results. We defined a rule set that ranked overlapping coalescence times for *T. c. bauri*/*T. m. triunguis* and *T. c. major* from Mississippi/Florida. The migration rate prior range (1.96 × 10^−6^–9.78 × 10^−5^) was estimated from the number of migrants (Genepop v4.7.5; Rousset 2008). We applied three jSFS binning classes and 5,000 RF trees to build the classifier and predict the models.

## 3 RESULTS

### 3.1 Sampling and data processing

We sequenced 214 geographically-widespread *Terrapene* (Figure 1; Supplementary Information Table S1) including all recognized species and subspecies save the rare *T. nelsoni klauberi*. ipyrad recovered 134,607 variable sites (of 1,163,463 total) across 14,760 retained loci, with 90,777 as parsimony informative. The mean per-individual depth was 56.3X (Supplementary Information Figure S2).

### 3.2 Species tree inference

The lineage tree contained N=214 tips (Figure 2), whereas those from SVDquartets (Figure 3a) and PoMo (Figure 3b) grouped individuals into N=26 populations, again per locality and subspecies. SVDquartets examined 10,299 unlinked SNPs and the species tree was assembled from 87,395,061 quartets. Full loci were used for PoMo. All trees clearly delineated eastern *versus* western clades, with *T. mexicana, T. carolina*, and *T. coahuila* composing the eastern clade, with western represented by *T. ornata* and *T. nelsoni*.

**FIGURE 2.**
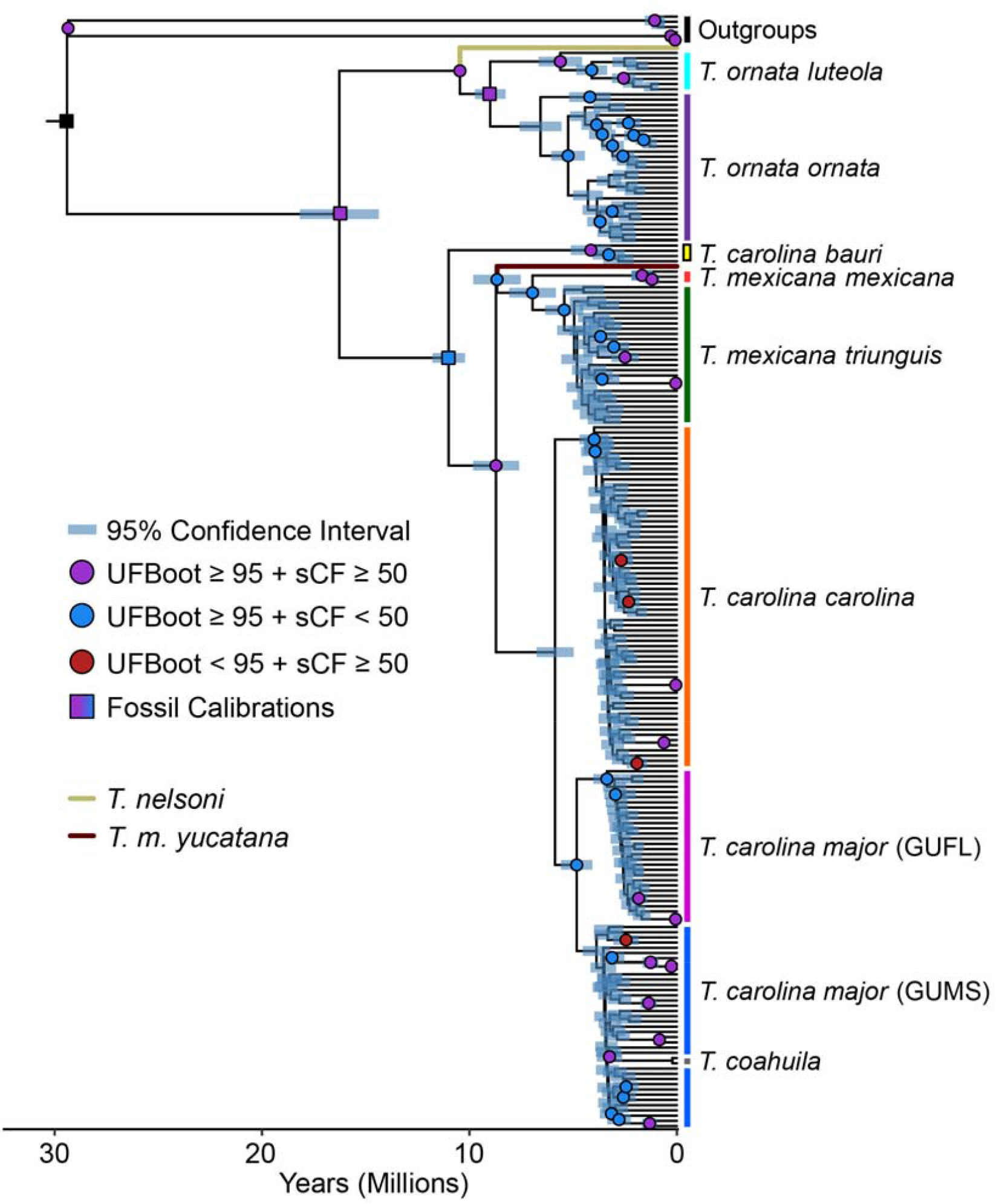
Chronogram reflecting relationships among 214 *Terrapene* ddRADseq samples as generated in IQ-TREE v2.1.2 and time-calibrated using LSD2. Node support was assessed with 1,000 ultrafast bootstrap (UFBoot) replicates, and site concordance-factors (sCF) calculated from 10,000 randomly-sampled quartets. Well-supported nodes (UFBoot≥95%, sCF≥50%) are represented by color-coded circles or squares, with squares showing fossil calibration points. Node bars reflect 95% confidence intervals based on 1,000 simulated trees. *Clemmys guttata* and *Emydoidea blandingii* represent outgroups.

**FIGURE 3.**
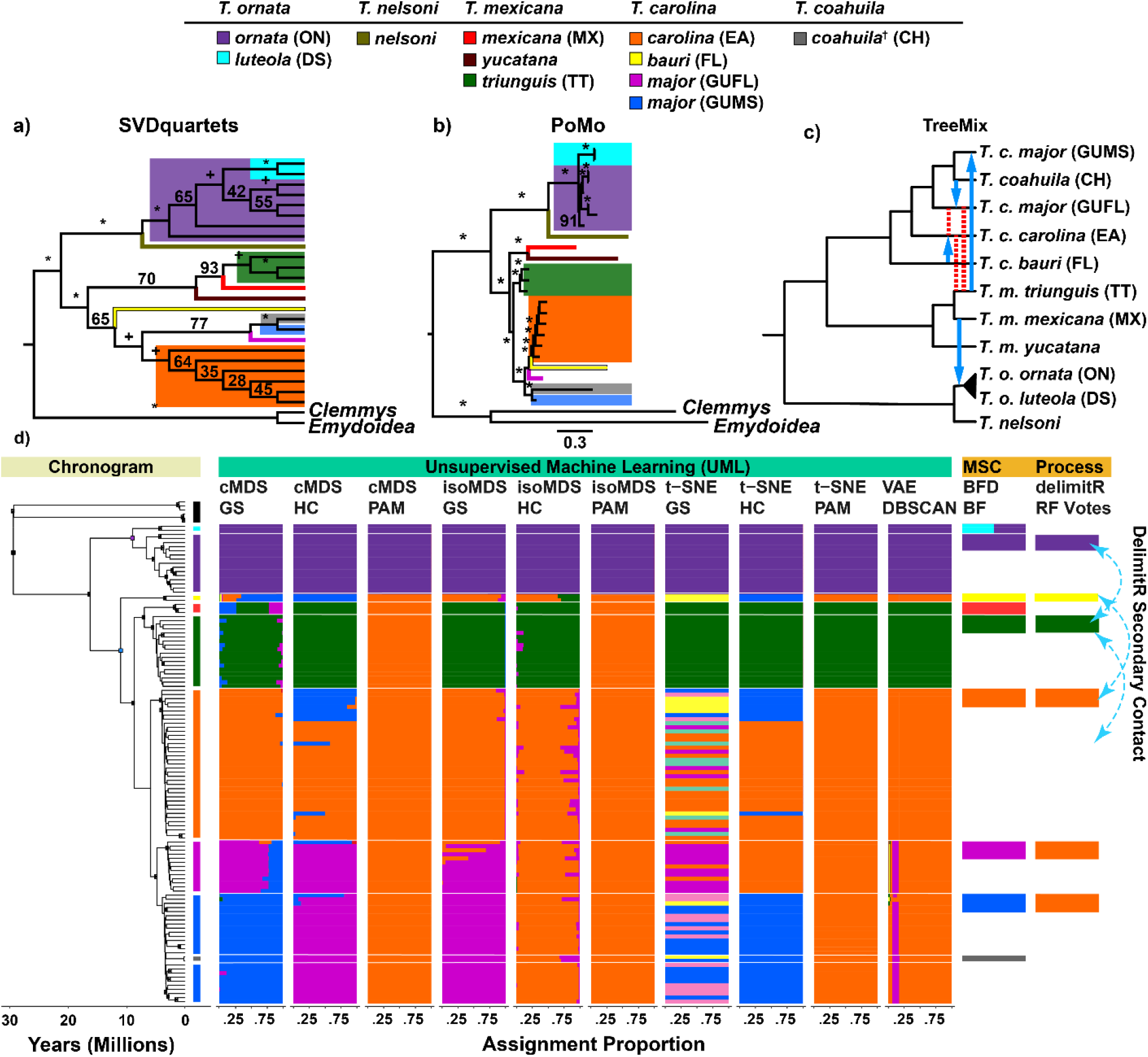
Species trees, TreeMix, and species delimitation results among *Terrapene* ddRADseq samples. Parenthetical legend abbreviations correspond to Tables 2 and 3. Phylogenies (N*=*214) were generated by (a) SVDquartets and (b) PoMo with 26 populations grouped by subspecies and state locality. ‘*’ and ‘+’ indicate 100% and ≥95% bootstrap support. (c) Migration supported by TreeMix (blue arrows) and previously published results (red/dashed lines; Martin *et al*. 2020). Outgroups were omitted for clarity. (d) Species delimitations for UML (N=117), multispecies coalescent (MSC; BFD=Bayes Factor Delimitation; N=37), and process-based (delimitR; N=28) methods. UML data filtering allowed ≤25% missing data per-individual and per-population, with minor allele frequency filters=5% (cMDS/t-SNE/VAE) and 1% (isoMDS), and t-SNE perplexity=15. UML includes RF=random forest, visualized with cMDS and isoMDS ordination, t-SNE, and VAE, with bar plots depicting assignment proportions among 100 replicates and aligning with chronogram tips. RF and t-SNE optimal *K* were assessed using partition around medoids (PAM)+gap statistic (GS), PAM+highest mean silhouette width (HMSW), and hierarchical clustering (HC)+HMSW, whereas VAE, BFD, and delimitR used DBSCAN, Bayes Factors (BF) and RF votes. Blue/dashed arrows show gene flow supported by delimitR. ‘†’ indicates a monotypic *T. coahuila*.

All phylogenies delineated *T. ornata* and *T. nelsoni*. However, SVDquartets nested *T. o. luteola* within a paraphyletic *T. o. ornata*, whereas IQ-TREE and PoMo represented them as reciprocally monophyletic. In the eastern clade, SVDquartets displayed two subdivisions: *Terrapene mexicana* (all subspecies) and *T. carolina*+*T. coahuila*. PoMo included *T. m. triunguis* as sister to *T. c. carolina*+*T. c. major* but paraphyletic with respect to *T. m. mexicana*+*T. m. yucatana*. Furthermore, SVDquartets, PoMo, and IQ-TREE each differed with respect to the placement of *T. c. bauri, T. coahuila*, and two previously recognized populations within *T. c. major* (Martin *et al*. 2013, 2020). SVDquartets depicted *T. c. bauri* as sister to the *major*/*coahuila*/*carolina* clade, whereas PoMo placed *T. c. major* from Mississippi/*coahuila* as sister to *T. c. major* (FL)/*bauri/carolina*. IQ-TREE placed *T. c. bauri* sister to *T. carolina*/*T. mexicana*, and *T. coahuila*/*T. c. major* (MS) sister to *T. c. carolina*/*T. c. major* (FL).

The topology tests failed to reject either Martin *et al*. (2013) or the SVDquartets trees, whereas morphology-based and PoMo trees were significantly rejected (Table 1). Although the SVDquartets tree was ranked highest, site-likelihood scores indicated a minority of sites drove those topologies (Supplementary Information Figure S3).

### 3.3 Species delimitation via BFD* and delimitR

TreeMix converged upon four migration edges (Figure 3c; Supplementary Information Figure S4), with gene flow identified between: *Terrapene m. mexicana* × *T. o. ornata+T. o. luteola*; T. *c. carolina* × *T. c. bauri*; *T. m. triunguis* × *T. c. major* (MS); and *T. coahuila* × *T. c. major* (FL). To target specific reticulation hypotheses, delimitR was run with a reduced set of sub-species, in compliance with computational constraints. The best-fitting delimitR model within selected taxa (*T. m. triunguis, T. o. ornata, T. c. major, T. c. bauri*, and *T. c. carolina*) was *K*=4 (posterior probability=0.98; Table 3; Figure 3d). Also, *T. c. major* and *T. c. carolina* were collapsed, and three secondary contact migration edges were apparent: *T. o. ornata* × *T. c. carolina*+*T. c. major*; *T. c. bauri* × *T. c. carolina*+*T. c. major*; and *T. o. ornata* × *T. m. triunguis*. The second-best model was identical save for excluding the latter migration, although it also had the highest error (Table 3).

BFD* supported two top models (Table 2), each delimited (*K*=9), and all distinct except *T. o. ornata*/*T. o. luteola* (*K*=8; Figure 3d). Although not statistically distinguishable (BF<2), both were decisively better than others (BF>10). Convergence was confirmed for the likelihood traces, with mean per-model ESS>300 (Supplementary Information Table S2).

### 3.4 UML species delimitation

UML results varied considerably (Figures 4, 5; Supplementary Information Figures S5-S10), with mean optimal *K* greatest for t-SNE, followed by cMDS, VAE, and isoMDS (Figures 4a, 5a). Across datasets, PAM clustering with the gap statistic (PAM+GS) exhibited the largest *K*, whereas PAM with the highest mean silhouette width (PAM+HMSW) was lowest (Figure 5b). Hierarchical clustering (HC)+HMSW and VAE were intermediate (Figures 4a, 5a; Supplementary Information Figure S5). Each algorithm delimited *T. ornata* from *T. carolina*+*T. mexicana* in most datasets, save PAM+HMSW in some of the larger datasets, and among some t-SNE replicates (e.g., Supplementary Information Appendix B, B1). In all cases, cMDS with PAM+GS and HC+HMSW further delimited *T. m. triunguis*+*T. m. mexicana* from *T. carolina*, whereas cMDS with PAM+HMSW did not. Whether the remaining algorithms did so depended upon filtering parameters. Finally, cMDS with PAM+GS and HC+HMSW further partitioned subgroups within *T. carolina* in most datasets, whereas isoMDS did so in a limited fashion, and t-SNE split *T. carolina* into multiple clusters without a phylogenetic pattern. Bar plots for 64 filtered datasets are in Supplementary Information Appendix B1-B60.

**FIGURE 4.**
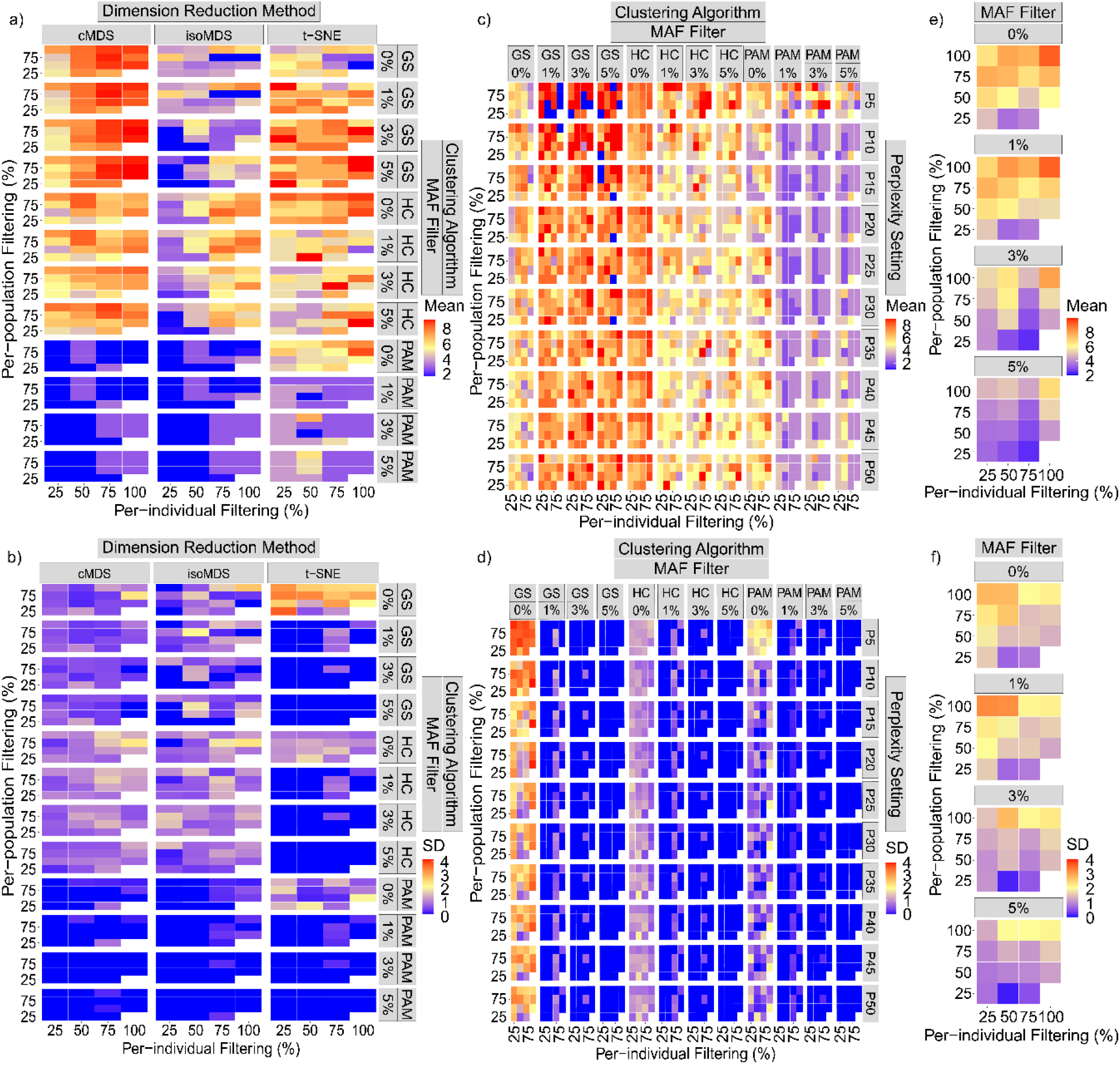
Heatmaps depicting mean and standard deviation (SD) of optimal *K* among 100 unsupervised machine learning species-delimitation replicates. Input ddRADseq alignments were filtered with a maximum of 25%, 50%, 75%, and 100% (=no filter) missing data allowed per-individual and per-population, and with minor allele frequency (MAF) filters as 5%, 3%, 1%, and 0% (=no filter). (a) and (b)=Pairwise missing data heatmaps for three dimensionality-reduction methods (cMDS and isoMDS=classical and isotonic multidimensional scaling), t-SNE=t-distributed stochastic neighbor embedding *versus* three clustering algorithms [(partition around medoids+gap statistic (GS)]; HC=hierarchical clustering+highest mean silhouette width (HMSW); PAM=partition around medoids+HMSW. (c) and (d)=t-SNE heatmap panels comparing clustering algorithms with ten perplexity (P) settings. (e) and (f)=VAE (variational autoencoder) heatmaps with optimal *K* chosen via DBSCAN.

**Figure 5.**
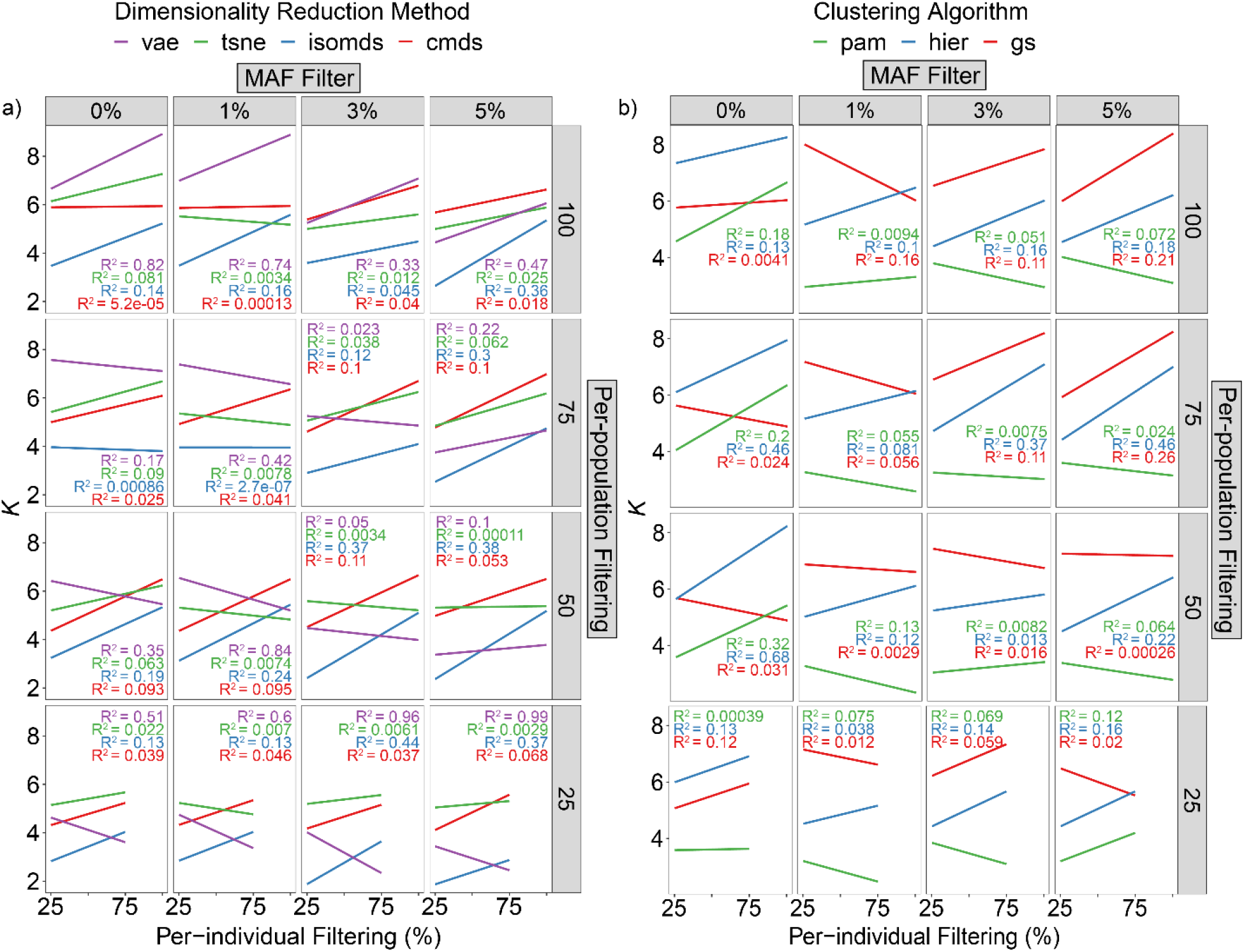
Regressions showing relationship between mean optimal *K (*y-axes), missing data, and minor allele frequency (MAF) filtering parameters. Missing data was filtered both per-individual (x-axes) and per-population (panel rows), with a maximum allowed of 25%, 50%, 75%, and 100% (=no filtering). Minor allele frequency (MAF) filters of 5%, 3%, 1%, and 0% (=no filtering) were also applied (panel columns). (a) Colors correspond to the dimensionality-reduction methods: cMDS and isoMDS=classical and isotonic multidimensional scaling, t-SNE=t-distributed stochastic neighbor embedding, VAE=variational autoencoder. (b) Colors indicate three clustering algorithms: GS=partition around medoids+gap statistic, HC=hierarchical clustering+highest mean silhouette width (HMSW), PAM=partition around medoids+HMSW.

We present representative results (Figure 3d) that displayed minimal inconsistencies among replicates and with respect to the phylogeny, with parameter choice also reflecting how each algorithm interacted with filtering values (below). This included 25% per-individual and per-population filters for all algorithms, a 5% MAF filter for cMDS, t-SNE, and VAE, and a 1% MAF filter for isoMDS. Five groups were delineated by cMDS with PAM+GS: *T. o. ornata* (ON)+*T. o. luteola* (DS), *T. c. major* from Mississippi (GUMS), *T. c. major* from Florida (GUFL), *T. c. carolina* (EA), and *T. m. mexicana* (MX)+*T. m. triunguis* (TT).

However, *T. c. bauri* displayed mixed assignment between *T. c. carolina* and GUMS. cMDS with HC+HMSW also delimited *K*=5 but lumped the two populations of *T. c. major*, splitting *T. c. bauri*, and grouped some *T. c. carolina* individuals with *T. c. bauri*. It also split *T. ornata* and *T. carolina*+*T. mexicana*. While isoMDS with PAM+GS resembled cMDS with HC+HMSW, it also clustered *T. c. bauri* with *T. c. carolina*. Similarly, isoMDS with HC+HMSW showed *T. o. ornata*+*T. o. luteola, T. c. carolina*+GUMS+GUFL, and *T. m. mexicana*+*T. m. triunguis*. However, isoMDS with PAM+HMSW only delimited *T. ornata* from *T. carolina*+*T. mexicana*. The model t-SNE (at perplexity=15) clearly partitioned *T. ornata, T. carolina*, and *T. mexicana*, though the PAM+GS algorithm exhibited spurious groupings within *T. carolina*. However, t-SNE with HC+HMSW clustered many *T. c. carolina* with GUFL and the remaining with GUMS. We found VAE and t-SNE with PAM+HMSW only delimited *T. ornata, T. carolina*, and *T. mexicana*.

### 3.5 Effects of data filtering

Among all dimensionality reduction and clustering algorithms, greater per-individual and per-population missing data generally increased mean optimal *K* and SD (Figures 4a-b and 5a-b; Supplementary Information Figure S5). PAM+HMSW deviated due to low *K*, regardless of filtering. This was manifested as two types of noise in the bar plots (Supplementary Information Appendix B1-B60): ‘vertical striping’ (inconsistency of assignment among replicates) and ‘horizontal striping’ (groupings inconsistent with phylogeny). We found the former largely driven by increased missing data per-individual, whereas the latter by increased missing data per-locus. However, performance varied among algorithms in how they interacted with both missing data parameters.

We found that t-SNE consistently resolved *T. ornata* and *T. carolina*+*T. mexicana*, but *T. mexicana* was only partitioned when per-population filtering was 25%. However, t-SNE did not further partition *T. carolina* in any dataset and displayed a tendency to form phylogenetically spurious groupings (=vertical striping). The perplexity grid search (Figures 4c-d and 5b; Supplementary Information Figures S6-S10) suggested that the highest *K* and SD among replicates was at perplexity=5-10, with a plateau at higher perplexities.

We also found cMDS with PAM+GS and HC+HMSW delineated most clades, save for inconsistency amongst the *T. c. major* populations and *T. coahuila*. In contrast, cMDS and isoMDS with PAM+HMSW typically displayed *K*=2 or 3 and contained no phylogenetically meaningful clusters with ≥75% missing data per-individual (e.g., Supplementary Information Appendix B62). Finally, VAE partitioned *T. ornata* from *T. carolina*+*T. mexicana* in all datasets, but *T. mexicana* was only delineated from *T. carolina* when per-individual missing data was ≤50% and with MAF filter.

Filtering by MAF ubiquitously reduced noise, although results varied by algorithm (Supplementary Information Appendix B1-B60). For t-SNE, optimal *K* and SD were reduced. In contrast, the clusters yielded by cMDS with PAM+GS and HC+HMSW were only marginally affected. We found cMDS and isoMDS with PAM+HMSW and MAF filters ≥3% were less noisy, but for isoMDS with PAM+GS and HC+HMSW the MAF filter effect was dependent on the number of individuals present in the dataset. With a maximum of 25% per-individual missing data (N=117), a 1% MAF filter shows minimal striping and higher *K* than did a >1% MAF filter. However, larger MAF filters have a greater effect above 25% per-individual filtering. Lastly, optimal *K*, SD, and striping in VAE were strongly influenced by MAF filters (Figures 4e-f, 5a, Supplementary Information Figure S5). With lower per-individual filters (≤50%) and a 5% MAF filter, VAE consistently delineated *T. mexicana* from *T. carolina*, even with high per-population filters. However, lower MAF and higher per-individual (>50%) filters introduced progressively more noise and grouped *T. carolina* and *T. mexicana*.

### 3.6 Relative performance among approaches

The cMDS model with PAM+GS and HC+HMSW consistently displayed the highest *K* and was less susceptible to data filtering. However, isoMDS with PAM+GS and HC+HMSW was more influenced by filtering parameters, but still consistently resolved the highest level of hierarchical structure (*T. ornata/T. carolina+T. mexicana*). Both cMDS and isoMDS with PAM+HMSW consistently displayed the lowest *K* at the top hierarchy and were usually in complete agreement. We note that t-SNE was highly susceptible to horizontal and vertical striping, and only partitioned *T. mexicana* from *T. carolina* ssp. at 25% per-individual filtering. Similarly, VAE performed far more consistently with a 5% MAF filter and ≤50% per-individual filtering. VAE also consistently hovered between K=2 and K=3, making it the second most conservative algorithm next to PAM+HMSW. In contrast, BFD* delimited the most taxa among all the approaches, splitting all save *T. o. luteola* and *T. o. ornata*, and delimitR partitioned *T. ornata, T. carolina, T. mexicana*, and *T. c. bauri*.

In terms of computational resources, the UML algorithms were far less intensive than BFD* and delimitR, enabling stochasticity to be assessed in many replicates. Each UML algorithm needed ∼1-3GB RAM per replicate and ∼2-3 days runtime for 100 replicates. Comparatively, BFD* required the greatest memory and time, often using >200GB RAM (with 16 CPU threads) and a ∼10-day runtime per model. We note delimitR used much less memory and was faster than BFD*, but output ∼3.2 TB with six tips and 51 models.

## 4 DISCUSSION

We observed substantial heterogeneity in resolving *Terrapene* via M-L approaches, which echoed previous morphological and single-gene results (Milstead 1967, 1969; Milstead & Tinkle 1967; Butler *et al*. 2011; Martin *et al*. 2013). We interpret this variability as reflecting inherent differences in dimensionality-reduction, clustering, and *K*-selection, as well how methodologies interact with biological aspects of the data and user-defined filtering.

### 4.1 Delimitation hypotheses and biological interpretations reconciled

Two factors likely contribute to the observed heterogeneity: 1) An hierarchical arrangement of phylogenetic signal (Martin *et al*. 2013); and 2) Phylogenetic discord (Martin *et al*. 2020). Both reverberate noticeably within prior literature and phylogenetic evaluations.

The most consistent grouping was eastern (*T. carolina+T. mexicana*) versus western (*T. ornata*) clades, representing the deepest *Terrapene* divergence (Figures 3a-b). This is unsurprising given it is the most prominent axis of molecular variation (morphologically corroborated; Milstead & Tinkle 1967; Dodd 2001) Nominal species have been identifiable since late Miocene (Holman & Fritz 2005), as corroborated by molecular dating (Figure 2).

#### 4.1.1 Terrapene ornata

Although introgression between *T. o. ornata* and *T. m. triunguis* occurred during secondary contact (Table 3; Figure 3d), no contemporary evidence for introgression among these clades emerged from previous evaluations, except rare *F*_1_ hybrids between *T. o. ornata* and *T. carolina* (Martin *et al*. 2020). TreeMix also suggested introgression between *T. ornata* and *T. m. mexicana* (Figure 3c). Although contact with *T. mexicana* was certainty possible during glacial expansion-contraction (Martin *et al*. 2020), we echo earlier conclusions that hybridization lacks justifiable taxonomic implications, per hybridization between *T. ornata* and *T. carolina* (Martin *et al*. 2020).

Regarding *T. ornata*, algorithms failed to further partition *T. o. ornata*/*T. o. luteola*, suggesting a lack of diagnosability at our most recent scale. Notably, both also lack reciprocal monophyly in some phylogenomic (Figure 3a) and single-gene analyses (Martin et al. 2013). They also lack clear morphological synapomorphies (Minx 1996). Although *T. o. luteola* exhibits habitat and movement patterns markedly different from mesic conspecifics (Nieuwolt 1996), few investigations have similarly compared *T. ornata* subspecies, such that inferences regarding reproductive isolation (or potential thereof) are difficult. Populations of *T. o. luteola* also do not exhibit thermal adaptations that are mutually exclusive from *T. o. ornata*, as might be surmised given other desert-dwelling tortoises (Plummer 2003).

Previous authors hypothesized *T. o. luteola* as a relict population (Milstead & Tinkle 1967). Weak differentiation [molecular: Martin *et al*. (2013); morphological: Dodd (2001)], as well as possible paraphyly of *T. o. ornata* (Figure 3a) suggest isolation was recent. Although phylogenetic structuring was present in some analyses (e.g., Figure 2), it is insufficient to mandate recognition beyond the subspecific level. However, special guidelines that delineate relictual lineages may be warranted (Mussmann *et al*. 2020), particularly given the isolation and reduced *N*_e_ in *T. o. luteola* (Nieuwolt 1996).

#### 4.1.2 Terrapene mexicana

The second most frequent split (Figures 2, 3a) divided *T. mexicana* and *T. carolina*, corresponding to the second-deepest phylogenetic node (Figures 2, 3a). This lends further support to a prior elevation of *T. mexicana* (Martin *et al*. 2013). Conspecifics of *T. mexicana* also share multiple morphological characteristics, such as carapace coloration and a degree of concavity to the posterior plastron, that separate the group from *T. carolina* (Minx 1996). *Terrapene mexicana mexicana* (as well as *T. m. yucatana*, excluded due to sample size) have isolated, allopatric ranges (Smith & Smith 1980; Ernst & Lovich 2009), with reproductive isolation difficult to assume.

Evidence for interbreeding of *T. m. triunguis* with *T. carolina* subspecies in the southeastern United States (Butler *et al*. 2011) has led some to conclude that species-level recognition of *T. mexicana sensu lato* is unwarranted (Fritz & Havaš 2014). Indeed, our own results suggest introgression between *T. m. triunguis* and *T. carolina* in secondary contact (Figure 3d). Martin *et al*. (2020) confirmed hybridization of *T. m. triunguis* with both *T. c. major* and *T. c. carolina* in the southeast, yet found genetic exchange was restricted, given that: (1) Genetically ‘pure’ individuals are predominant throughout the contact zone; and (2) patterns of gene-level exchange exhibit strong sigmoidal patterns, suggesting selection against interspecific heterozygotes. Additionally, the sigmoidal pattern was strongest within a subset of genes involved in thermal adaptation (Martin *et al*. 2020), suggesting species boundaries are modulated by an adaptive barrier between co-occurring *T. mexicana* and *T. carolina* sub-species. This functional perspective corroborates the proposed taxonomy herein, and by Martin et al. (2013).

#### 4.1.3 Terrapene carolina

Partitioning within *T. carolina* echoed inconsistencies in our phylogenies (Figures 2, 3a-b), and seemingly depended upon algorithm and filtering regime (Figure 3d; Supplementary Information B). *Terrapene carolina major*, for example, occasionally split from the remaining *T. carolina* (usually including *T. coahuila*; cMDS+HC, Figure 3d), whereas in other cases, *T. c. major* (FL and MS) were separated (with the former grouped into *T. c. carolina*) (t-SNE+HC, Fig. 3d).

In contrast to steep clines in interspecific comparisons (Martin *et al*. 2020; see above), a transect of the *T. c. carolina* and *T. c. major* contact zone revealed shallow genetic transition, with multiple loci showing potential signatures of selection-driven introgression. Previous authors have hypothesized either direct ancestry (Bentley & Knight 1998) or historic admixture with a now extinct taxon, [*T. c. putnami*; Butler *et al*. (2011)]. While such ‘ghost’ admixture can mislead population structure (Lawson *et al*. 2018), such a signal is unlikely manufactured in entirety. In contrast to Butler *et al*. (2011), Martin *et al*. (2020) found a pervasive signal of population structure and strong molecular diagnosability in *T. c. major*, with a cryptic east-west division roughly defined by the Apalachicola River [a recurring phylogeographic discontinuity reflecting recolonization from disparate Gulf Coast refugia; Soltis *et al*. (2006)]. Our interpretations refuted the ‘genetic melting pot’ assertion (Fritz & Havaš 2014) and favored instead recognition of the two as distinct evolutionarily significant units (ESUs). Additionally, differences in habitat use and movement patterns distinguish *T. c. major* (Meck *et al*. 2020), which spends greater time in mesic habitats (e.g., floodplain swamps). In support, early studies observed a distinct webbing of the hind foot in *T. c. major* (Taylor 1895). Given the genetic data herein, we reject the taxonomic coalescence of *T. c. major*.

*Terrapene carolina bauri* was similarly resistant to straightforward classification, although generally grouping with *T. c. major* (when the latter was separated from *T. c. carolina*; Figure 3b). We found *T. c. bauri* as sister to either the remaining *T. carolina* group, *T. c. carolina*+*T. c. major*, or only *T. c. carolina* (Figures 2-3; Martin *et al*. 2013). This argues against it being sister to *T. m. triunguis* (per Spinks *et al*. 2009). Osteologically, it alone shares a complete zygomatic arch with *T. c. major* (Taylor 1895; Ditmars 1934), although other morphological investigations have allied it more closely with *T. c. carolina* (Minx 1996). Thus, phylogenetic inconsistency for *T. c. bauri* clearly extends beyond our results.

Although hybridization likely contributes to this issue (as with *T. c. major*), the biogeography of the region may provide insight, with peninsular Florida recognized as a distinct biogeographic province (Ennen *et al*. 2017). Intraspecific division are recognized in multiple species [e.g., *Chelydra serpentina, Deirochelys reticularia* (Walker & Avise 1998)], a phylogenetic legacy likely reflecting periodic isolation from the mainland that may have inflated genetic divergences (Douglas *et al*. 2006), and facilitated secondary contact. This scenario is supported by delimitR and TreeMix (Figures 3c-d). Here, we again stress that evidence is sufficient to support continued recognition, yet not for taxonomic elevation.

#### 4.1.4 Terrapene coahuila

*Terrapene coahuila* represents a persistent phylogenetic uncertainty (Spinks *et al*. 2009; Wiens *et al*. 2010; Martin *et al*. 2013). It is unique in that it occupies streams, ponds, and marshes, with terrestrial movements restricted to the rainy seasons (Webb *et al*. 1963). Milstead (1967) postulated that *T. coahuila* evolved as a relictual population of a *Terrapene* ancestor (potentially the extinct *T. c. putnami*) during pluvial periods associated with Pleistocene glacial-interglacial cycles across the broad eastern coastal plain of Mexico. In this scenario, relictual populations are what remains from those north-south migrations, as hypothesized for *T. m. mexicana* and *T. m. yucatana*. The scenario is plausible, given semi-aquatic adaptations in the presumed ancestor (*T. c. putnami*) and closely related *T. c. major*, as well as shared morphologies between extinct *T. c. putnami* and modern *T. coahuila* (Milstead 1967). The phylogenetic placement of *T. coahuila*, as nested within *T. c. major*, offers further evidence (Figure 2-3), as does the almost unanimous UML grouping in our results (Figure 3d; Supplementary Information Appendix B1-B60). As with *T. o. luteola*, small, isolated populations that differ in evolutionary rates could contribute to a lack of molecular similarity with extant *T. c. major*, despite a unique functional morphology (Brown 1971).

### 4.2 Relative performance of species-delimitation methods

As with prior studies (Derkarabetian *et al*. 2019; Mussmann *et al*. 2020), we also found considerable variation among methods, some of which can be attributed either to idiosyncrasies in the data or to algorithms and their implementation. First, among RF methods cMDS with PAM+GS and HC+HMSW displayed higher *K* and isoMDS generally yielded smaller *K* (Figure 3d), with the latter being attributed by Derkarabetian *et al*. (2019) to the retention of only two dimensions. PAM+HMSW (Figure 3d) also trended towards a small *K*=2, corresponding to the deepest *Terrapene* bifurcation, and suggesting a potential failure in identifying hierarchical clusters. Here, a solution might include partitioning divergent subtrees for separate analyses.

In contrast to Derkarabetian *et al*. (2019), we found t-SNE the most inclined to produce inconsistent groupings, a pattern most prevalent with the gap statistic (Supplementary Information Appendix B1-B60). Mussmann *et al*. (2020) concurred, although in their case it was PAM+HMSW. We see this as an inherent problem relating to data structure. Previous comparisons of t-SNE found low fidelity with global data patterns, and latent space distances were poor proxies for ‘true’ among-group distances, particularly when compared to VAE (Becht *et al*. 2019; Battey *et al*. 2020). This potentially explains our observed ‘plateau’ of mean optimal *K* and SD in the t-SNE perplexity grid-search, in that perplexity defines relative weighting of local versus global components (Wattenberg *et al*. 2016). It may also explain the formation of spurious clusters even at higher perplexities, in that clusters are formed *post hoc* (PAM or HC). Thus, t-SNE may perform poorly when inter-cluster distances/dispersion in global data structure are skewed, although it is not clear to what degree hyperparameter choice and initializations contribute (Belkina *et al*. 2019; Kobak & Berens 2019).

In our case, VAE with DBSCAN yielded higher fidelity to the underlying phylogeny (Figure 3a) and was also more robust to missing data (Figures 4e-f). A particular benefit of the VAE approach is the output of a standard deviation around samples in latent space (Derkarabetian *et al*. 2019). Our DBSCAN hyperparameters were informed directly from latent variable uncertainties, and in so doing, we circumvented the issue of *K-*selection that drove heterogeneity in the RF and t-SNE methods [also recognized with other clustering approaches (Janes *et al*. 2017)].

By comparison, BFD* partitioned all groups, which may reflect a vulnerability to local structure at the population level, as reported by others for MSC methods (Sukumaran & Knowles 2017). BFD* and VAE partitioned equally in Mussmann *et al*. (2020), although their populations were relictual and without contemporary connectivity, whereas *Terrapene* reflects both historical (Figure 3d) and contemporary gene flow (Martin *et al*. 2020). In corroboration, other studies have also demonstrated reticulation to condense VAE clusters (Derkarabetian *et al*. 2019; Newton *et al*. 2020). Although not run on a full dataset, delimitR formed clusters consistent with (or similar to) several of the UML methods (e.g., isoMDS+GS; Figure 3d, Table 3). The latter displayed a particular utility regarding testing targeted hypotheses relating to demographic processes such as migration, whereas these must be applied to UML results *post hoc*.

### 4.3 Data treatment and assignment consistency

We generally found a tendency for UML methods to ‘over-split’ given large amounts of missing data, and phylogenetically inconsistent groupings (‘horizontal striping’) were most pronounced when missing data was elevated per-individual (Supplementary Information Appendix B1-B60). However, low-level, undetected introgression could also drive such a pattern. Mussmann *et al*. (2020) noted a similar pattern with the RF methods, possibly reflecting an artificial similarity among samples generated by a non-random distribution of missing data. A similar ‘vertical striping’ effect was seen when missing data was elevated per-locus (e.g., Supplementary Information Appendix B13), often manifested as inconsistency among replicates. However, effects varied across methods, as per previous analyses [phylogeographic: Graham *et al*. (2020); phylogenetic: Molloy & Warnow (2018)].

Missing-data bias is a particular concern when patterns are non-random (i.e., presence or absence of observations are data-dependent; Rubin 1976). Here, the temptation is to filter stringently, yet we found highly filtered datasets were biased towards smaller *K*, generally retaining only nodes deepest within the phylogeny. The same pattern was identified using the VAE method (Newton *et al*. 2020), and is intuitive given expectations that a major subset of missing ddRAD data are systematically distributed [defined by mutation-disruption of restriction sites: Gautier *et al*. (2013); Eaton *et al*. (2017)]. Thus, indiscriminate exclusion may unintendedly bias information content leading to the underestimation of diversity (Arnold *et al*. 2013; Leaché *et al*. 2015; Huang & Knowles 2016). Again, care must be taken to filter the data such that sufficient discriminatory signal remains, while also being mindful of the signal-to-noise ratio, and the underlying biases driving interactions of sparse data versus information content (Nakagawa & Freckleton 2008).

A potential solution involves the input of genotypes to fill in missing values (per Howie *et al*. 2009; Durbin 2014; Das *et al*. 2016). However, a cautious *a priori* designation of population references is needed, particularly when group-delimitation is the goal. It may be appropriate to employ phylogenetically-informed methods previously applied in comparative studies (e.g., Goolsby *et al*. 2017).

We found MAF filters dampened the effect of missing data, likely by removing sequencing errors and uninformative variants at low-frequency (Mathieson & McVean 2012; Jakobsson *et al*. 2013). In a similar context, Linck & Battey (2019) found MAF filters to significantly increase in the discriminatory capacity of assignment-test methods (Structure; Pritchard *et al*. 2000). In our case, MAF filtering reduced noise and improved group differentiation (e.g., resulting in lower variability among replicates; Figures 4-5, Supplementary Information Figures S5-S6), although this might prompt the M-L algorithms to miss low levels of introgression. Thus, we view it as a parameter in need of further empirical exploration.

### 4.4 Conclusions

UML approaches identify groups based on the structure of the data, and as such, represent a natural extension to species-delimitation approaches. However, we found idiosyncrasies regarding: Phylogenetic context of the study system (e.g., hierarchical structure, reticulation); the manner by which clustering and *K*-selection approaches were applied *post hoc*; and the bioinformatic treatment of the data. We particularly note that lax filtering, performed to maximize size and information content, actually promote spurious groupings and inflate variability among replicates. An alternate method, i.e., filtering via MAF to promote informative characters, favorably altered the signal-to-noise ratio and increased the consistency of our delimitations. Thus, we recommend that UML practitioners test multiple algorithms, veer away from high levels of missing data, and utilize MAF filters. We conclude that UML approaches, when applied to formulate taxonomic hypotheses and reduce dimensionality of complex data, are valuable and computationally efficient tools for integrative species-delimitation, as demonstrated within our study system.

## ACKNOWLEDGEMENTS

Many thanks to those volunteers, organizations, and agencies that contributed tissue samples (Supplementary Information Table S1). We also thank colleagues for guidance at various stages of this project: A. Alverson, W. Anthonysamy, M. Bangs, J. Beaulieu, J. Koukl, P. Martin, S. Mussmann, J. Pummill, and Z. Zbinden. Sample collections were approved under Animal Care and Use Committee (IACUC) protocols: #113 (University of Texas/Tyler), #16160 and #18000 (University of Illinois/Champaign-Urbana). Funding sources included the Lucille F. Stickle Fund of the North American Box Turtle Committee, the American Turtle Observatory, and University of Arkansas endowments to MRD and MED. The Arkansas High Performance Computing Cluster (AHPCC) and Jetstream cloud (XSEDE #TG-BIO160065) provided computational resources.

## AUTHOR CONTRIBUTIONS

BTM and TKC designed the research, implemented laboratory protocols, authored scripts, and wrote the manuscript. BTM conducted lab work and data analyses. MRD and MED guided the study design, provided funding, and contributed to manuscript development. JSP facilitated collection of *Terrapene* tissues and provided methodological expertise. RDB collected *Terrapene* tissues from southeastern North America and facilitated access to additional samples. CAP provided taxon expertise and provided many *T. ornata* tissues. All authors contributed to and revised the manuscript.

## DATA AVAILABILITY STATEMENT

Raw ddRADseq data are available on the GenBank Nucleotide Database at https://www.ncbi.nlm.nih.gov/bioproject/563121 (BioProject ID: 563121). Scripts for parsing and plotting UML output are available on GitHub at https://github.com/btmartin721/mecr_boxturtle. Input and output files for all analyses can be found in a Dryad Digital Repository (DOI: https://doi.org/10.5061/dryad.xgxd254fc).

## REFERENCES

Al’Aref SJ, Anchouche K, Singh G, Slomka PJ, Kolli KK, Kumar A, Pandey M, Maliakal G, Van Rosendael AR, and Beecy AN (2019) Clinical applications of machine learning in cardiovascular disease and its relevance to cardiac imaging. European Heart Journal, 40, 1975–1986.

Allendorf FW, Hohenlohe PA, and Luikart G (2010) Genomics and the future of conservation genetics. Nature Reviews Genetics, 11, 697–709.

Andrews S (2010) FastQC: a quality control tool for high throughput sequence data. https://www.bibsonomy.org/bibtex/2b6052877491828ab53d3449be9b293b3/ozborn.

Arnold B, CorbettLJDetig RB, Hartl D, and Bomblies K (2013) RAD seq underestimates diversity and introduces genealogical biases due to nonrandom haplotype sampling. Molecular Ecology, 22, 3179–3190.

Auffenberg W (1958) Fossil turtles of the genus Terrapene in Florida. Bulletin of the Florida State Museum, 3, 53–92.

Auffenberg W (1959) A Pleistocene Terrapene hibernaculum, with remarks on a second complete box turtle skull from Florida. Quarterly Journal of the Florida Academy of Science, 22, 49–53.

Austerlitz F, David O, Schaeffer B, Bleakley K, Olteanu M, Leblois R, Veuille M, and Laredo C (2009) DNA barcode analysis: a comparison of phylogenetic and statistical classification methods. BMC Bioinformatics, 10, S10.

Avise JC (2000a) Cladists in Wonderland. Evolution, 54, 1828–1832.

Avise JC (2000b) Phylogeography: the history and formation of species. Harvard University Press, Cambridge, MA.

Battey CJ, Coffing GC, and Kern AD (2020) Visualizing population structure with variational autoencoders. bioRxiv, 248278.

Becht E, McInnes L, Healy J, Dutertre C-A, Kwok IWH, Ng LG, Ginhoux F, and Newell EW (2019) Dimensionality reduction for visualizing single-cell data using UMAP. Nature Biotechnology, 37, 38–44.

Belkina AC, Ciccolella CO, Anno R, Halpert R, Spidlen J, and Snyder-Cappione JE (2019) Automated optimized parameters for t-distributed stochastic neighbor embedding improve visualization and analysis of large datasets. Nature Communications, 10, 1–12.

Bentley CC and Knight JL (1998) Turtles (Reptilia: Testudines) of the Ardis local fauna late Pleistocene (Rancholabrean) of South Carolina. Brimleyana, 25, 1–33.

Breiman L (2001) Random Forests. Machine Learning, 45, 5–32.

Brown WS (1971) Morphometrics of Terrapene coahuila (Chelonia, Emydidae), with comments on its evolutionary status. The Southwestern Naturalist, 16, 171–184.

Butler JM, Dodd Jr. CK, Aresco M, and Austin JD (2011) Morphological and molecular evidence indicates that the Gulf Coast box turtle (Terrapene carolina major) is not a distinct evolutionary lineage in the Florida Panhandle. Biological Journal of the Linnean Society, 102, 889–901.

Chambers EA and Hillis DM (2019) The multispecies coalescent over-splits species in the case of geographically widespread taxa. Systematic Biology, 69, 184–193.

Chernomor O, Von Haeseler A, and Minh BQ (2016) Terrace aware data structure for phylogenomic inference from supermatrices. Systematic Biology, 65, 997–1008.

Chifman J and Kubatko L (2014) Quartet inference from SNP data under the coalescent model. Bioinformatics, 30, 3317–3324.

Chollet F (2015) Keras. https://keras.io.

Das S, Forer L, Schönherr S, Sidore C, Locke AE, Kwong A, Vrieze SI, Chew EY, Levy S, and McGue M (2016) Next-generation genotype imputation service and methods. Nature Genetics, 48, 1284–1287.

Derkarabetian S, Castillo S, Koo PK, Ovchinnikov S, and Hedin M (2019) A demonstration of unsupervised machine learning in species delimitation. Molecular Phylogenetics and Evolution, 139, 106562.

Ditmars RL (1934) A review of the box turtles. Zoologica, 17, 1–44.

Dodd KC (2001) North American Box Turtles, A Natural History. University of Oklahoma Press, Norman, OK, USA.

Douglas MRE, Douglas MRE, Schuett GW, and Porras LW (2006) Evolution of rattlesnakes (Viperidae; Crotalus) in the warm deserts of western North America shaped by Neogene vicariance and Quaternary climate change. Molecular Ecology, 15, 3353–3374.

Durbin R (2014) Efficient haplotype matching and storage using the positional Burrows–Wheeler transform (PBWT). Bioinformatics, 30, 1266–1272.

Eaton DAR and Overcast I (2020) ipyrad: Interactive assembly and analysis of RADseq datasets. Bioinformatics, 36, 2592–2594.

Eaton DAR, Spriggs EL, Park B, and Donoghue MJ (2017) Misconceptions on missing data in RAD-seq phylogenetics with a deep-scale example from flowering plants. Systematic Biology, 66, 399–412.

Edwards S V, Potter S, Schmitt CJ, Bragg JG, and Moritz C (2016) Reticulation, divergence, and the phylogeography–phylogenetics continuum. Proceedings of the National Academy of Sciences, 113, 8025–8032.

Eldredge N and Cracraft J (1980) Phytigenetic Patterns and the Evolutinary Process: Methods and Theory in Comparative Biology. Columbia University Press, New York, NY, USA.

Ennen JR, Matamoros WA, Agha M, Lovich JE, Sweat SC, and Hoagstrom CW (2017) Hierarchical, quantitative biogeographic provinces for all North American turtles and their contribution to the biogeography of turtles and the continent. Herpetological Monographs, 31, 114–140.

Ernst CH and Lovich JE (2009) Turtles of the united states and Canada, 2nd Edition. The John Hopkins University Press, Baltimore, MD, USA.

Ester M, Kriegel H-P, Sander J, and Xu X (1996) A density-based algorithm for discovering clusters in large spatial databases with noise. In: Proceedings of the Second International Conference on Knowledge Discovery and Data Mining, pp. 226–231.

Excoffier L, Dupanloup I, Huerta-Sánchez E, Sousa VC, and Foll M (2013) Robust demographic inference from genomic and SNP data. PLoS Genetics, 9, e1003905.

Feldman CR and Parham JF (2002) Molecular phylogenetics of emydine turtles: Taxonomic revision and the evolution of shell kinesis. Molecular Phylogenetics and Evolution, 22, 388– 398.

Fraley C and Raftery AE (1998) How many clusters? Which clustering method? Answers via model-based cluster analysis. The Computer Journal, 41, 578–588.

Francis RM (2017) pophelper: an R package and web app to analyse and visualize population structure. Molecular Ecology Resources, 17, 27–32.

Fritz U and Havaš P (2013) Order Testudines: 2013 update. In: Zhang, Z.-Q. (Ed.) Animal Biodiversity: An Outline of Higher-level Classification and Survey of Taxonomic Richness (Addenda 2013). Zootaxa, 3703, 12–14.

Fritz U and Havaš P (2014) On the reclassification of Box Turtles (Terrapene): A response to Martin et al. (2014). Zootaxa, 3835, 295–298.

Funk DJ and Omland KE (2003) Species-level paraphyly and polyphyly: frequency, causes, and consequences, with insights from animal mitochondrial DNA. Annual Review of Ecology, Evolution, and Systematics, 34, 397–423.

Gautier M, Gharbi K, Cezard T, Foucaud J, Kerdelhué C, Pudlo P, Cornuet J-M, and Estoup A (2013) The effect of RAD allele dropout on the estimation of genetic variation within and between populations. Molecular Ecology, 22, 3165–3178.

Goolsby EW, Bruggeman J, and Ané C (2017) Rphylopars: fast multivariate phylogenetic comparative methods for missing data and withinLJspecies variation. Methods in Ecology and Evolution, 8, 22–27.

Graham MR, SantibáñezLJLópez CE, Derkarabetian S, and Hendrixson BE (2020) Pleistocene persistence and expansion in tarantulas on the Colorado Plateau and the effects of missing data on phylogeographical inferences from RADseq. Molecular Ecology, 29, 3684–3701.

Hoang DT, Chernomor O, von Haeseler A, Minh BQ, and Vinh LS (2017) UFBoot2: improving the ultrafast bootstrap approximation. Molecular Biology and Evolution, 35, 518–522.

Holman JA and Fritz U (2005) The box turtle genus Terrapene (TestudinesLJ: Emydidae) in the Miocene of the USA. Journal of Herpetology, 15, 81–90.

Howie BN, Donnelly P, and Marchini J (2009) A flexible and accurate genotype imputation method for the next generation of genome-wide association studies. PLoS Genetics, 5, e1000529.

Huang H and Knowles LL (2016) Unforeseen Consequences of Excluding Missing Data from Next-Generation Sequences: Simulation Study of RAD Sequences. Systematic Biology, 65, 357–365.

Iverson JB, Meylan PA, and Seidel ME (2017) Testudines—Turtles. In: Scientific and Standard English Names of Amphibians and Reptiles of North America North of Mexico, with Comments Regarding Confidence in Our Understanding (ed Crother BI), pp. 82–91. SSAR Herpetological Circular 43.

Jakobsson M, Edge MD, and Rosenberg NA (2013) The relationship between FST and the frequency of the most frequent allele. Genetics, 193, 515–528.

Janes JK, Miller JM, Dupuis JR, Malenfant RM, Gorrell JC, Cullingham CI, and Andrew RL (2017) The K = 2 conundrum. Molecular Ecology, 26, 3594–3602.

Jombart T and Ahmed I (2011) adegenet 1.3-1: new tools for the analysis of genome-wide SNP data. Bioinformatics, 27, 3070–3071.

Kalyaanamoorthy S, Minh BQ, Wong TKF, von Haeseler A, and Jermiin LS (2017) ModelFinder: fast model selection for accurate phylogenetic estimates. Nature Methods, 14, 587–589.

Kass RE and Raftery AE (1995) Bayes Factors. Journal of the American Statistical Association, 90, 773–795.

Kaufman L and Rousseeuw P (1987) Clustering by means of medoids. Statistical Data Analysis Based on the L1-Norm and Related Methods, 405–416.

Kingma DP and Welling M (2013) Auto-encoding variational bayes. In: Proceedings of the International Conference on Learning Representations (ICLR). arXiv:1312.6114 [stat.ML].

Kobak D and Berens P (2019) The art of using t-SNE for single-cell transcriptomics. Nature Communications, 10, 1–14.

Kopelman NM, Mayzel J, Jakobsson M, Rosenberg NA, and Mayrose I (2015) CLUMPAK: a program for identifying clustering modes and packaging population structure inferences across K. Molecular Ecology Resources, 15, 1179–1191.

Kruskal JB and Wish M (1978) Multidimensional Scaling. Sage Publishing, Thousand Oaks, CA, USA.

Lawson DJ, van Dorp L, and Falush D (2018) A tutorial on how not to over-interpret STRUCTURE and ADMIXTURE bar plots. Nature Communications, 9, 3258.

Leaché AD, Banbury BL, Felsenstein J, De Oca AN-M, and Stamatakis A (2015) Short tree, long tree, right tree, wrong tree: new acquisition bias corrections for inferring SNP phylogenies. Systematic Biology, 64, 1032–1047.

Leaché AD, Fujita MK, Minin VN, and Bouckaert RR (2014) Species delimitation using genome-wide SNP data. Systematic Biology, 63, 534–542.

Linck EB and Battey CJ (2019) Minor allele frequency thresholds strongly affect population structure inference with genomic datasets. Molecular Ecology Resources, 19, 639–647.

Long C and Kubatko L (2018) The effect of gene flow on coalescent-based species-tree inference. Systematic Biology, 67, 770–785.

Maaten L van der and Hinton G (2008) Visualizing data using t-SNE. Journal of Machine Learning Research, 9, 2579–2605.

Mace GM (2004) The role of taxonomy in species conservation. Philosophical Transactions of the Royal Society B: Biological Sciences, 359, 711–719.

Martin BT, Bernstein NP, Birkhead RD, Koukl JF, Mussmann SM, and Placyk JS (2013) Sequence-based molecular phylogenetics and phylogeography of the American box turtles (Terrapene spp.) with support from DNA barcoding. Molecular Phylogenetics and Evolution, 68, 119–134.

Martin BT, Bernstein NP, Birkhead RD, Koukl JF, Mussmann SM, and Placyk Jr JS (2014) On the reclassification of the Terrapene (Testudines: Emydidae): a response to Fritz & Havaš. Zootaxa, 3835, 292–294.

Martin BT, Douglas MR, Chafin TK, Placyk JS, Birkhead RD, Phillips CA, and Douglas ME (2020) Contrasting signatures of introgression in North American box turtle (Terrapene spp.) contact zones. Molecular Ecology, 29, 4186–4202.

Mathieson I and McVean G (2012) Differential confounding of rare and common variants in spatially structured populations. Nature Genetics, 44, 243–246.

Mayr E (1963) Animal Species and Evolution. Belknap Press at Harvard University Press, Cambridge, MA.

Meck JR, Jones MT, Willey LL, and Mays JD (2020) Autecological study of Gulf Coast box turtles (Terrapene carolina major) in the Florida Panhandle, USA, reveals unique spatial and behavioral characteristics. Herpetological Conservation and Biology, 15, 293–305.

Milstead WW (1967) Fossil box turtles (Terrapene) from central North America, and box turtles of eastern Mexico. Copeia, 1967, 168–179.

Milstead WW (1969) Studies on the evolution of the box turtles (genus Terrapene). Bulletin of the Florida State Museum, Biological Science Series, 14, 1–113.

Milstead WW and Tinkle DW (1967) Terrapene of Western Mexico, with comments on species groups in the genus. Copeia, 1967, 180–187.

Minh BQ, Hahn MW, and Lanfear R (2018) New methods to calculate concordance factors for phylogenomic datasets. bioRxiv, 487801.

Minh BQ, Schmidt HA, Chernomor O, Schrempf D, Woodhams MD, Von Haeseler A, and Lanfear R (2020) IQ-TREE 2: New models and efficient methods for phylogenetic inference in the genomic era. Molecular Biology and Evolution, 37, 1530–1534.

Minx P (1992) Variation in phalangeal formulas in the turtle genus Terrapene. Journal of Herpetology, 26, 234–238.

Minx P (1996) Phylogenetic relationships among the box turtles, Genus Terrapene. Herpetologica, 52, 584–597.

Molloy EK and Warnow T (2018) To include or not to include: the impact of gene filtering on species tree estimation methods. Systematic Biology, 67, 285–303.

Mussmann SM, Douglas MR, Oakey DD, and Douglas ME (2020) Defining relictual biodiversity: Conservation units in speckled dace (Leuciscidae: Rhinichthys osculus) of the Greater Death Valley ecosystem. Ecology and Evolution, 10, 10798–10817.

Nakagawa S and Freckleton RP (2008) Missing inaction: the dangers of ignoring missing data. Trends in Ecology & Evolution, 23, 592–596.

Newton LG, Starrett J, Hendrixson BE, Derkarabetian S, and Bond JE (2020) Integrative species delimitation reveals cryptic diversity in the southern Appalachian Antrodiaetus unicolor (Araneae: Antrodiaetidae) species complex. Molecular Ecology, 29, 2269–2287.

Nguyen L-T, Schmidt HA, von Haeseler A, and Minh BQ (2015) IQ-TREE: A fast and effective stochastic algorithm for estimating maximum-likelihood phylogenies. Molecular Biology and Evolution, 32, 268–274.

Nielsen R, Paul JS, Albrechtsen A, and Song YS (2011) Genotype and SNP calling from next-generation sequencing data. Nature Reviews Genetics, 12, 443.

Nieuwolt PM (1996) Movement, activity, and microhabitat selection in the western box turtle, Terrapene ornata luteola, in New Mexico. Herpetologica, 487–495.

Nosil P and Feder JL (2012) Genomic divergence during speciation: causes and consequences. Philosophical Transactions of the Royal Society B: Biological Sciences, 367, 332–342.

Pedregosa F, Varoquaux G, Gramfort A, Michel V, Thirion B, Grisel O, Blondel M, Prettenhofer P, Weiss R, and Dubourg V (2011) Scikit-learn: Machine learning in Python. Journal of Machine Learning Research, 12, 2825–2830.

Peterson BK, Weber JN, Kay EH, Fisher HS, and Hoekstra HE (2012) Double digest RADseq: an inexpensive method for de novo SNP discovery and genotyping in model and non-model species. PLoS One, 7, e37135.

Pickrell JK and Pritchard JK (2012) Inference of population splits and mixtures from genome-wide allele frequency data. PLoS Genetics, 8, e1002967.

Plummer M V (2003) Activity and thermal ecology of the box turtle, Terrapene ornata, at its southwestern range limit in Arizona. Chelonian Conservation and Biology, 4, 569–577.

Pritchard JK, Stephens M, and Donnelly P (2000) Inference of population structure using multilocus genotype data. Genetics, 155, 945–959.

De Queiroz K (2007) Species concepts and species delimitation. Systematic Biology, 56, 879– 886.

R Development Core Team (2018) R: A language and environment for statistical computing. https://cran.r-project.org/.

Rambaut A, Drummond AJ, Xie D, Baele G, and Suchard MA (2018) Posterior summarization in bayesian phylogenetics using Tracer 1.7 (E Susko, Ed,). Systematic Biology, 67, 901–904.

Rodriguez-Galiano VF, Ghimire B, Rogan J, Chica-Olmo M, and Rigol-Sanchez JP (2012) An assessment of the effectiveness of a random forest classifier for land-cover classification. ISPRS Journal of Photogrammetry and Remote Sensing, 67, 93–104.

Rousseeuw PJ (1987) Silhouettes: A graphical aid to the interpretation and validation of cluster analysis. Journal of Computational and Applied Mathematics, 20, 53–65.

Rousset F (2008) genepop ‘007: a complete re-implementation of the genepop software for Windows and Linux. Molecular Ecology Resources, 8, 103–106.

Rubin DB (1976) Inference and missing data. Biometrika, 63, 581–592.

Schrempf D, Minh BQ, De Maio N, von Haeseler A, and Kosiol C (2016) Reversible polymorphism-aware phylogenetic models and their application to tree inference. Journal of Theoretical Biology, 407, 362–370.

Shepard RN, Romney AK, and Nerlove SB (1972) Multidimensional Scaling: Theory and Applications in the Behavioral Sciences: I. Theory. Seminar Press, New York City, NY, USA.

Smith ML and Carstens BC (2020) ProcessLJbased species delimitation leads to identification of more biologically relevant species. Evolution, 74, 216–229.

Smith ML, Ruffley M, Espíndola A, Tank DC, Sullivan J, and Carstens BC (2017) Demographic model selection using random forests and the site frequency spectrum. Molecular Ecology, 26, 4562–4573.

Smith HM and Smith RB (1980) Synopsis of the herpetofauna of Mexico: Volume VI, guide to Mexican turtles, bibliographic addendum III. John Johnson, North Bennington, Vermont (“1979”), xviii + 1044 pp.

Soltis DE, Morris AB, McLachlan JS, Manos PS, and Soltis PS (2006) Comparative phylogeography of unglaciated eastern North America. Molecular Ecology, 15, 4261–4293.

Spinks PQ and Shaffer HB (2009) Conflicting mitochondrial and nuclear phylogenies for the widely disjunct Emys (Testudines: Emydidae) species complex, and what they tell us about biogeography and hybridization. Systematic Biology, 58, 1–20.

Spinks PQ, Thomson RC, Lovely GA, and Shaffer HB (2009) Assessing what is needed to resolve a molecular phylogeny: Simulations and empirical data from emydid turtles. BMC Evolutionary Biology, 9, 56.

Stephens PR and Wiens JJ (2003) Ecological diversification and phylogeny of emydid turtles. Biological Journal of the Linnaean Society, 79, 577–610.

Sukumaran J and Knowles LL (2017) Multispecies coalescent delimits structure, not species. Proceedings of the National Academy of Sciences of the United States of America, 114, 1607–1611.

Taylor WE (1895) The box tortoises of North America. Proceedings of the United States National Museum, 17, 573–588.

Tibshirani R, Walther G, and Hastie T (2001) Estimating the number of clusters in a data set via the gap statistic. Journal of the Royal Statistical Society: Series B (Statistical Methodology), 63, 411–423.

To T-H, Jung M, Lycett S, and Gascuel O (2016) Fast dating using least-squares criteria and algorithms. Systematic Biology, 65, 82–97.

Via S (2009) Natural selection in action during speciation. Proceedings of the National Academy of Sciences, 106, 9939–9946.

Walker DE and Avise JC (1998) Principles of phylogeography as illustrated by freshwater and terrestrial turtles in the southeastern United States. Annual Review of Ecology and Systematics, 29, 23–58.

Wattenberg M, Viégas F, and Johnson I (2016) How to use t-SNE effectively. Distill, 1, e2.

Webb RG, Minckley WL, and Craddock JE (1963) Remarks on the Coahuilan box turtle, Terrapene coahuila (Testudines, Emydidae). The Southwestern Naturalist, 8, 89–99.

Wiens JJ, Kuczynski CA, and Stephens PR (2010) Discordant mitochondrial and nuclear gene phylogenies in emydid turtles: implications for speciation and conservation. Biological Journal of the Linnaean Society, 99, 445–461.

Yang Z and Rannala B (2010) Bayesian species delimitation using multilocus sequence data. Proceedings of the National Academy of Sciences, 107, 9264–9269.

