## Supplementary Information Appendix A for "The choices we make and the impacts they have: Machine learning and species delimitation in North American box turtles (*Terrapene* spp.)"

### 1. Additional phylogenomic analyses

#### 1.1. Topology tests

We performed IQ-TREE topology tests using for four *Terrapene* phylogenetic hypotheses: (a) The SVDquartets and (b) PoMo topologies, as generated herein; (c) Sanger sequencing with mtDNA and nuclear introns (Martin *et al.* 2013); and (d) Morphological data (Minx 1996). ModelFinder was again employed to optimize per-partition substitution models (Kalyaanamoorthy *et al.* 2017), and nodal confidence of each tree was assessed using 1,000 ultrafast bootstrap (UFB) replicates (Hoang *et al.* 2017). We then compared support among the constraint trees using seven topological tests, each with 10,000 re-samplings: (a) Raw log-likelihoods; (b) bootstrap proportion test using the RELL approximation (bpRELL; Kishino *et al.* 1990); (c) Kishino-Hasegawa test (KH; Kishino & Hasegawa 1989); (d) Shimodaira-Hasegawa test (SH; Shimodaira & Hasegawa 1999); (e) Approximately Unbiased test (AU; Shimodaira 2002); and (f) Expected Likelihood Weights (ELW; Strimmer & Rambaut 2002). To visualize support for each topology across the genome, site-likelihood probabilities and pairwise site-likelihood score differences (Δ*SLS*) were calculated between the best-supported *versus* remaining trees.

### 2. Species delimitation analyses

#### 2.1. BFD* Prior Selection and Data Filtering

Here, we derived appropriate priors following (Bangs *et al.* 2020). We first calculated a pairwise distance matrix using the DIVEIN web server (Deng *et al.* 2010). We did so with a random subset of the full concatenated alignment (N_sites_=36,800, the maximum allowed by DIVEIN) derived using a custom Perl script, *nremover.pl* (<https://github.com/tkchafin/scripts>). Average within-species divergence was calculated from the DIVEIN pairwise distance matrix across all taxa to represent our prior for ancestral population size (*θ*=0.000730885) which served as the mean (α/β) for a gamma-distributed prior. The coalescent rate was set to 2/*θ*=2736.4086. The lineage birth-rate for the Yule process (λ=196.5038) was determined with *pyule* (<https://github.com/joaks1/pyule>), which invokes tree height and number of species to determine λ. Tree height was calculated as ½ the maximum among-group pairwise distance (=0.002549775), and the number of species was conservatively set to three to limit potential biases from over-splitting (a tendency for multi-species coalescent species delimitation approaches). The mutation rate priors were fixed to 1.0 per recommendations from the BFD* tutorial (Leaché & Bouckaert 2018).

Before running BFD*, we first removed loci containing >50% missing data, both globally and per-population, using a custom Perl script *phylipFilterPops.pl* (<https://github.com/tkchafin/scripts>). Thus, all retained sites contained SNP data in at least 50% of individuals from each population. We further filtered the alignment by removing non-binary SNPs and invariant sites via the *Phrynomics* R package (<https://github.com/bbanbury/phrynomics>). We then generated XML files for 20 models using Beauti v2.5.2 and ran BFD* via the SNAPP v1.4.2 plug-in for BEAST v2.5.2 (Bryant *et al.* 2012; Bouckaert *et al.* 2019).

#### 2.2. Machine learning data preparation

Using R v3.5.1 (R Core R Development Core Team 3.0.1. 2013), we ran a slightly modified version of the R script developed by Derkarabetian *et al.* [(2019) ([*PCA-DAPC-RF-tSNE_str.r*](https://github.com/shahanderkarabetian/uml_species_delim/blob/master/PCA-DAPC-RF-tSNE_str.r)*;* <https://github.com/shahanderkarabetian/uml_species_delim>)] to load and prepare the input alignments, perform the random forest (RF) and t-distributed stochastic neighbor embedding (t-SNE) machine learning algorithms, and identify taxon clusters. The modified script, *PCA-DAPC-RF-tSNE_gridSearch_maf.r*, adds the capability of performing multiple independent runs among multiple datasets to assess RF and t-SNE variability and evaluate model performance among differently filtered datasets. It also performs a t-SNE grid search for the perplexity setting. Generally, the script used the R package *adegenet* v2.1.1 (Jombart & Ahmed 2011) to load the input alignments from Structure-formatted files. The data were scaled using the *scaleGen* function and subjected to dimensionality reduction via principle component analysis (PCA; *dudi.pca* function in *adegenet*). The full suite of PCA axes were assessed using DAPC (discriminant analysis of principle components; Jombart *et al.* 2010) cross-validation with 1,000 replicates (*xvalDapc* function in *adegenet*) to determine the optimal number of principle components and discriminant functions to retain. The scaled PCA data with the optimal number of axes were ultimately used as input for the RF and t-SNE analyses.

Input alignments for VAE (variational autoencoder; Kingma & Welling 2013) were generated from PHYLIP-formatted files using a custom python script, *phylip2onehotsnps.py*. VAE was then run via the Python3 script developed by Derkarabetian *et al.* (2019), with some minor modifications ([*sp_deli_clust_commandline_noClust.py*](https://github.com/btmartin721/mecr_boxturtle/blob/main/Derkarabetian_etal_2019/sp_deli_clust_commandline_noClust.py); [*vae_dbscan.py*](https://github.com/btmartin721/mecr_boxturtle/blob/main/vae_dbscan.py)). Changes included the implementation of a training/test data split to assess model performance, an early stopping callback to reduce overfitting, support for multiple independent runs to evaluate variability and cluster stability, and the DBSCAN clustering algorithm to determine the optimal number of clusters (*K*) in an unsupervised manner. Modified scripts can be found in a GitHub repository: <https://github.com/btmartin721/mecr_boxturtle>.

#### 2.3. Random forest

A user-specified number of classification/ decision trees (i.e., a “forest”) are created by the Random Forest (RF) algorithm (Breiman 2001), and classification trees (*N*=10,000) are then trained with random data subsets from which majority-vote class predictions are made. Nodes containing overlapping among-sample distances elevate a “proximity score” that is bootstrapped and aggregated (i.e., “bagged”) over all classification trees, with higher proximity scores indicating similar individuals. The output proximity matrix was visualized using two dimensionality reduction algorithms, classic and isotonic multidimensional scaling (cMDS and isoMDS; Shepard *et al.* 1972; Kruskal & Wish 1978). cMDS utilizes the full dissimilarity matrix from the RF classifier, whereas isoMDS forces a monotonic transformation that only uses the ranks from the proximity scores. Thus, cMDS preserves among-sample distances, whereas isoMDS does not.

#### 2.4. t-SNE

Similar to PCA, t-SNE is a dimensionality reduction algorithm (Maaten & Hinton 2008). Rather than using proximity scores to generate probability distributions representing similarities between samples in multidimensional space, it instead employs non-parametric, non-linear algorithms to estimate pairwise distances. It then attempts to minimize differences between high-dimensional space versus low-dimensional embedding. Samples with low similarity continue to repel each other as each iteration occurs, such that they become diffuse across parameter space. t-SNE was run for 20,000 iterations, within which the equilibria of the clusters were visually confirmed. Perplexity, which limits the effective number of neighbors, was tested at values ranging from 5-50 (incrementing by five), with the initial number of dimensions parameter set to five.

#### 2.5. Variational autoencoders

SNPs were first converted from a PHYLIP file to a binary ‘one-hot’ format, from which two latent variables representing the sample mean (µ) and standard deviation (σ) can be inferred by VAE (as implemented by Derkarabetian *et al.* 2019). VAE reconstructs the SNP dataset using the latent variables and self-trains by minimizing the difference (i.e., model loss) between the input and reconstructed datasets. Following training, latent variables are predicted from the full dataset and represented in two-dimensional space.

VAE was run with three encoder and decoder layers, each containing 100 neurons subjected to a dropout rate of 0.5 to reduce overfitting. Encoded SNP data was normalized and scaled to reduce the impact of stochasticity, with input split into datasets representing 80% training/ 20% validation (as is standard with machine learning). Model loss was assessed using an early stopping callback function from the scikit-learn Python package (Pedregosa *et al.* 2011) to determine an optimal number of epochs (i.e. cycles through the training dataset). Ideally, this should terminate when loss (~error) has converged and is minimized among both the training and validation datasets [(i.e. the ‘Goldilocks zone’; Al’Aref *et al.* 2019) (Fig. S3)]. An escalating loss in the validation dataset indicates overfitting. On the other hand, losses that have not yet reached their minimum value suggest model underfitting (i.e. a lack of generalization for both training and unseen data). Other parameters were chosen following Derkarabetian *et. al.* (2019).

### 3. REFERENCES

Al’Aref SJ, Anchouche K, Singh G, Slomka PJ, Kolli KK, Kumar A, Pandey M, Maliakal G, Van Rosendael AR, and Beecy AN (2019) Clinical applications of machine learning in cardiovascular disease and its relevance to cardiac imaging. *European Heart Journal*, **40**, 1975–1986.

Bangs MR, Douglas MR, Chafin TK, and Douglas ME (2020) Gene flow and species delimitation in fishes of Western North America: Flannelmouth (*Catostomus latipinnis*) and Bluehead sucker (*C. Pantosteus discobolus*). *Ecology and Evolution*, **10**, 6477–6493.

Bouckaert R, Vaughan TG, Barido-Sottani J, Duchêne S, Fourment M, Gavryushkina A, Heled J, Jones G, Kühnert D, De Maio N, Matschiner M, Mendes FK, Müller NF, Ogilvie HA, du Plessis L, Popinga A, Rambaut A, Rasmussen D, Siveroni I *et al.* (2019) BEAST 2.5: An advanced software platform for Bayesian evolutionary analysis (M Pertea, Ed,). *PLOS Computational Biology*, **15**, e1006650.

Breiman L (2001) Random Forests. *Machine Learning*, **45**, 5–32.

Bryant D, Bouckaert R, Felsenstein J, Rosenberg NA, and RoyChoudhury A (2012) Inferring species trees directly from biallelic genetic markers: bypassing gene trees in a full coalescent analysis. *Molecular Biology and Evolution*, **29**, 1917–1932.

Deng W, Maust B, Nickle D, Learn G, Liu Y, Heath L, Kosakovsky Pond S, and Mullins J (2010) DIVEIN: a web server to analyze phylogenies, sequence divergence, diversity, and informative sites. *BioTechniques*, **48**, 405–408.

Derkarabetian S, Castillo S, Koo PK, Ovchinnikov S, and Hedin M (2019) A demonstration of unsupervised machine learning in species delimitation. *Molecular Phylogenetics and Evolution*, **139**, 106562.

Hoang DT, Chernomor O, von Haeseler A, Minh BQ, and Vinh LS (2017) UFBoot2: improving the ultrafast bootstrap approximation. *Molecular Biology and Evolution*, **35**, 518–522.

Jombart T and Ahmed I (2011) adegenet 1.3-1: new tools for the analysis of genome-wide SNP data. *Bioinformatics*, **27**, 3070–3071.

Jombart T, Devillard S, and Balloux F (2010) Discriminant analysis of principal components: a new method for the analysis of genetically structured populations. *BMC Genetics*, **11**, 94.

Kalyaanamoorthy S, Minh BQ, Wong TKF, von Haeseler A, and Jermiin LS (2017) ModelFinder: fast model selection for accurate phylogenetic estimates. *Nature Methods*, **14**, 587–589.

Kingma DP and Welling M (2013) Auto-encoding variational bayes. In: Proceedings of the International Conference on Learning Representations (ICLR). arXiv:1312.6114 [stat.ML].

Kishino H and Hasegawa M (1989) Evaluation of the maximum likelihood estimate of the evolutionary tree topologies from DNA sequence data, and the branching order in hominoidea. *Journal of Molecular Evolution*, **29**, 170–179.

Kishino H, Miyata T, and Hasegawa M (1990) Maximum likelihood inference of protein phylogeny and the origin of chloroplasts. *Journal of Molecular Evolution*, **31**, 151–160.

Kruskal JB and Wish M (1978) *Multidimensional Scaling*. Sage Publisiinig, Thousand Oaks, CA, USA.

Leaché A and Bouckaert R (2018) Species trees and species delimitation with SNAPP: a tutorial and worked example. http://evomicsorg.wpengine.netdna-cdn.com/wp-content/uploads/2018/01/BFD-tutorial-1.pdf.

Maaten L van der and Hinton G (2008) Visualizing data using t-SNE. *Journal of Machine Learning Research*, **9**, 2579–2605.

Minx P (1996) Phylogenetic relationships among the box turtles, Genus *Terrapene*. *Herpetologica*, **52**, 584–597.

Pedregosa F, Varoquaux G, Gramfort A, Michel V, Thirion B, Grisel O, Blondel M, Prettenhofer P, Weiss R, and Dubourg V (2011) Scikit-learn: Machine learning in Python. *Journal of Machine Learning Research*, **12**, 2825–2830.

R Development Core Team 3.0.1. (2013) A language and environment for statistical computing. *R Foundation for Statistical Computing*, **2**, https://www.R-project.org.

Shepard RN, Romney AK, and Nerlove SB (1972) *Multidimensional Scaling: Theory and Applications in the Behavioral Sciences: I. Theory.* Seminar Press.

Shimodaira H (2002) An approximately unbiased test of phylogenetic tree selection. *Systematic Biology*, **51**, 492–508.

Shimodaira H and Hasegawa M (1999) Multiple comparisons of log-likelihoods with applications to phylogenetic inference. *Molecular Biology and Evolution*, **16**, 1114–1116.

Strimmer K and Rambaut A (2002) Inferring confidence sets of possibly misspecified gene trees. *Proceedings of the Royal Society of London. Series B: Biological Sciences*, **269**, 137–142.
