## Supplementary Information Appendix B for "The choices we make and the impacts they have: Machine learning and species delimitation in North American box turtles (*Terrapene* spp.)"

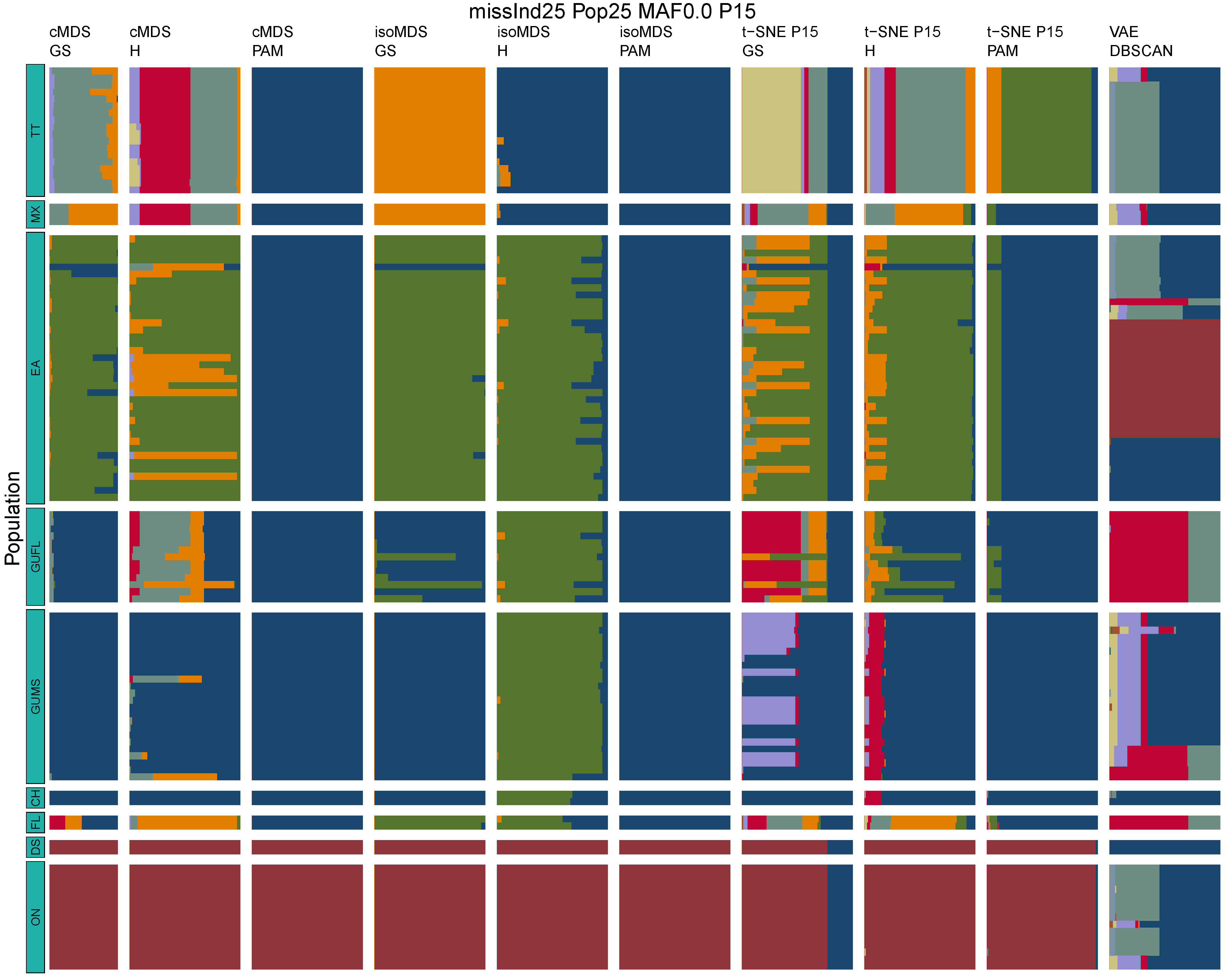

**Figure B1:** *Terrapene* unsupervised machine learning (UML) barplots depicting assignment proportions (x axis) among 100 replicates. Filtering parameters allowed maximum per-individual and per-population missing data and minimum minor allele frequency (MAF) filters. The included maximum missing data and minimum MAF filter proportions are shown in the plot title (missInd=per-individual, Pop=per-population filtering, MAF=MAF filter, P=perplexity). Each barplot on the page represents one UML algorithm: random forest, visualized with classical and isotonic multidimensional scaling (cMDS and isoMDS) ordination. t-SNE=t-distributed stochastic neighbor embedding with perplexity=15 (P15), and VAE=variational autoencoder. cMDS, isoMDS, and t-SNE were clustered with the partition around medoids (PAM) and hierarchical clustering (H) algorithms, and optimal *K* was chosen using PAM with the gap statistic (GS), H with the highest mean silhouette width (HMSW), and PAM with the HMSW. For VAE, clustering was performed, and optimal *K* chosen, with DBSCAN. *Terrapene* populations are separated by the horizontal white strips. Population IDs (blue left-aligned strip) include the ON=Ornate (*T. ornata ornata*), DS=Desert (*T. o. luteola*), FL=Florida (*T. c. bauri*), CH=Coahuilan (*T. coahuila*), GUMS=Gulf Coast (*T. carolina major*) from Mississippi, GUFL=Gulf Coast from the Florida Panhandle, EA=Woodland (*T. c. carolina*), MX=Mexican (*T. mexicana mexicana*), and TT=Three-toed (*T. m. triunguis*).

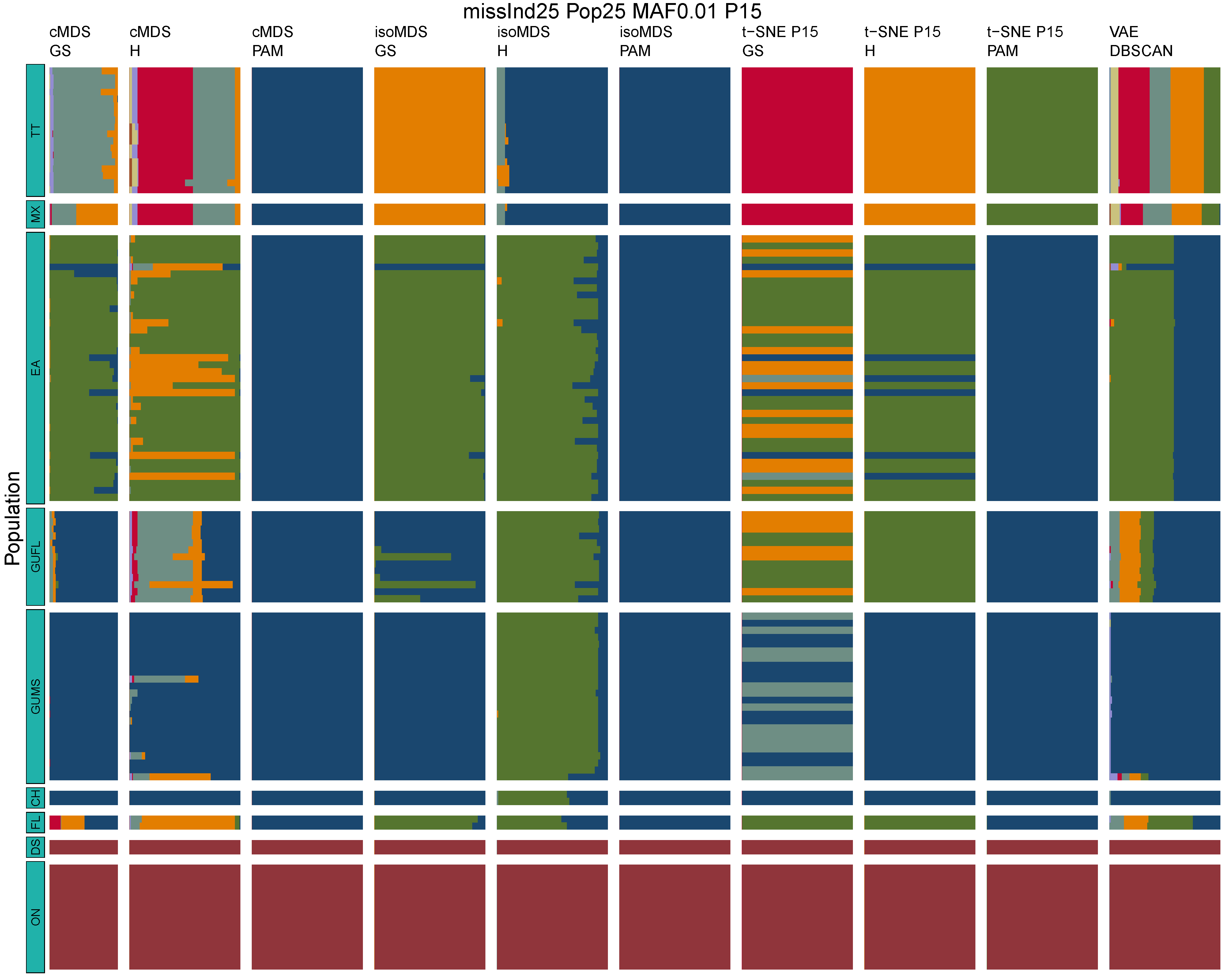

**Figure B2:** *Terrapene* unsupervised machine learning (UML) barplots depicting assignment proportions (x axis) among 100 replicates. Filtering parameters allowed maximum per-individual and per-population missing data and minimum minor allele frequency (MAF) filters. The included maximum missing data and minimum MAF filter proportions are shown in the plot title (missInd=per-individual, Pop=per-population filtering, MAF=MAF filter, P=perplexity). Each barplot on the page represents one UML algorithm: random forest, visualized with classical and isotonic multidimensional scaling (cMDS and isoMDS) ordination. t-SNE=t-distributed stochastic neighbor embedding with perplexity=15 (P15), and VAE=variational autoencoder. cMDS, isoMDS, and t-SNE were clustered with the partition around medoids (PAM) and hierarchical clustering (H) algorithms, and optimal *K* was chosen using PAM with the gap statistic (GS), H with the highest mean silhouette width (HMSW), and PAM with the HMSW. For VAE, clustering was performed, and optimal *K* chosen, with DBSCAN. *Terrapene* populations are separated by the horizontal white strips. Population IDs (blue left-aligned strip) include the ON=Ornate (*T. ornata ornata*), DS=Desert (*T. o. luteola*), FL=Florida (*T. c. bauri*), CH=Coahuilan (*T. coahuila*), GUMS=Gulf Coast (*T. carolina major*) from Mississippi, GUFL=Gulf Coast from the Florida Panhandle, EA=Woodland (*T. c. carolina*), MX=Mexican (*T. mexicana mexicana*), and TT=Three-toed (*T. m. triunguis*).

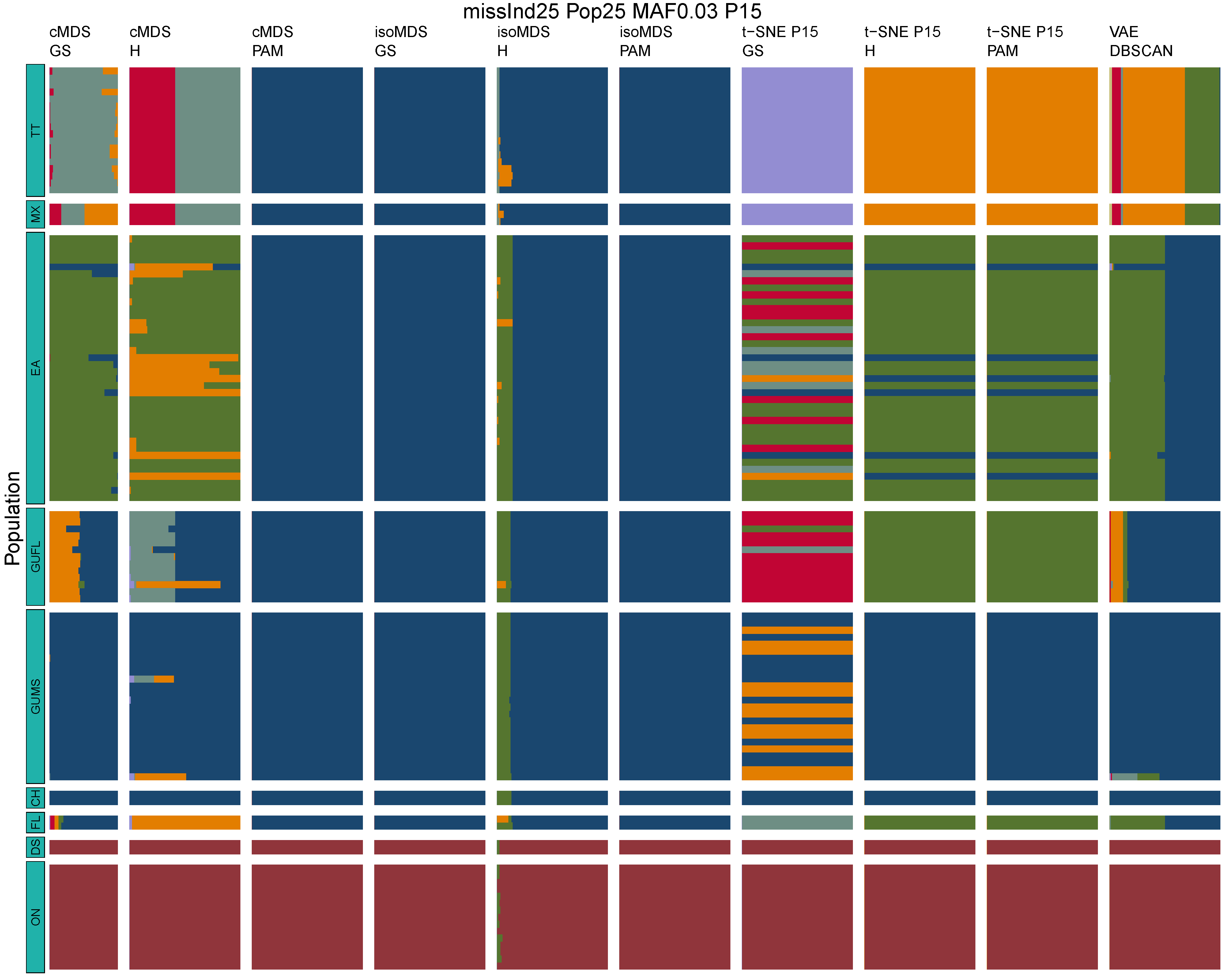

**Figure B3:** *Terrapene* unsupervised machine learning (UML) barplots depicting assignment proportions (x axis) among 100 replicates. Filtering parameters allowed maximum per-individual and per-population missing data and minimum minor allele frequency (MAF) filters. The included maximum missing data and minimum MAF filter proportions are shown in the plot title (missInd=per-individual, Pop=per-population filtering, MAF=MAF filter, P=perplexity). Each barplot on the page represents one UML algorithm: random forest, visualized with classical and isotonic multidimensional scaling (cMDS and isoMDS) ordination. t-SNE=t-distributed stochastic neighbor embedding with perplexity=15 (P15), and VAE=variational autoencoder. cMDS, isoMDS, and t-SNE were clustered with the partition around medoids (PAM) and hierarchical clustering (H) algorithms, and optimal *K* was chosen using PAM with the gap statistic (GS), H with the highest mean silhouette width (HMSW), and PAM with the HMSW. For VAE, clustering was performed, and optimal *K* chosen, with DBSCAN. *Terrapene* populations are separated by the horizontal white strips. Population IDs (blue left-aligned strip) include the ON=Ornate (*T. ornata ornata*), DS=Desert (*T. o. luteola*), FL=Florida (*T. c. bauri*), CH=Coahuilan (*T. coahuila*), GUMS=Gulf Coast (*T. carolina major*) from Mississippi, GUFL=Gulf Coast from the Florida Panhandle, EA=Woodland (*T. c. carolina*), MX=Mexican (*T. mexicana mexicana*), and TT=Three-toed (*T. m. triunguis*).
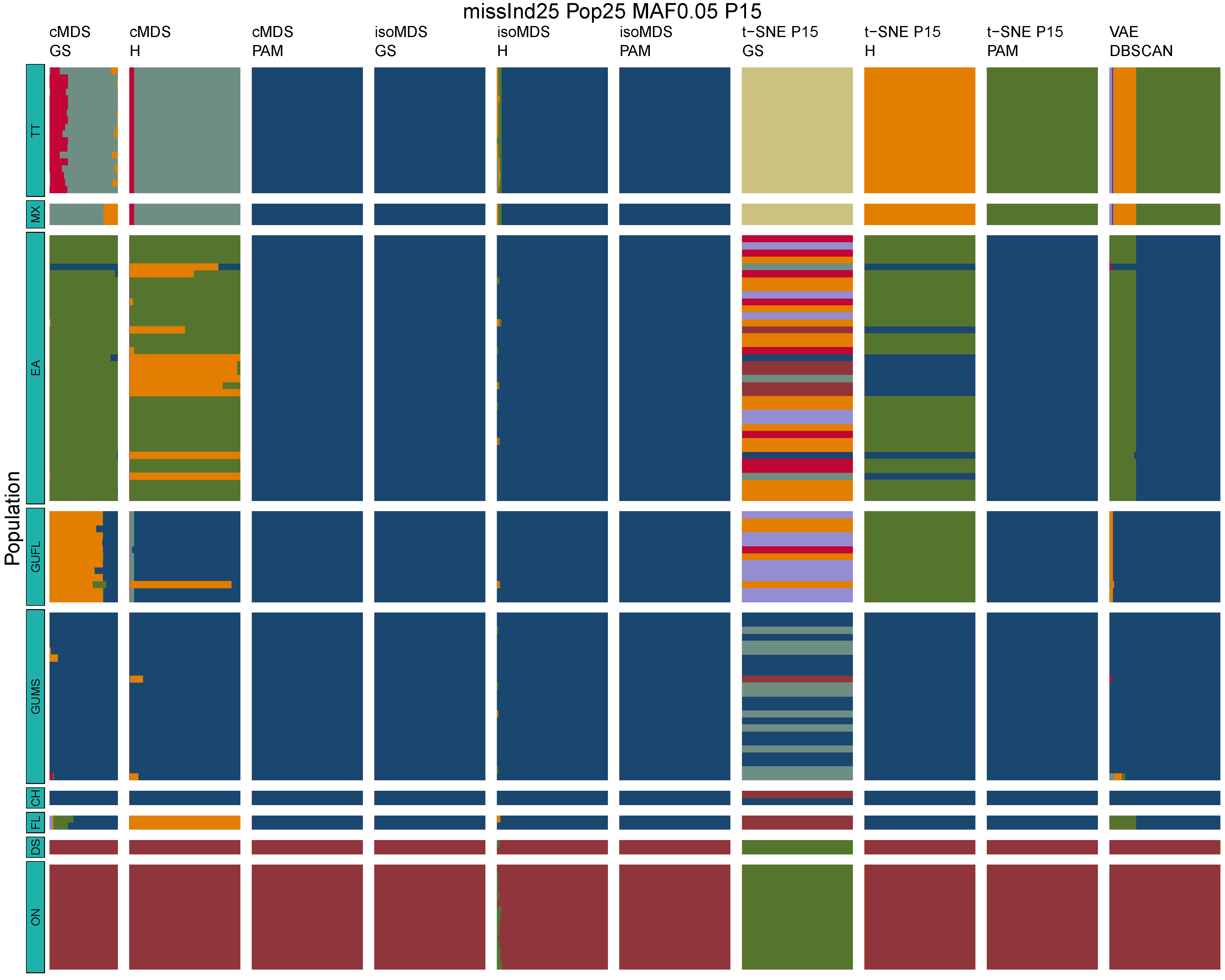

**Figure B4:** *Terrapene* unsupervised machine learning (UML) barplots depicting assignment proportions (x axis) among 100 replicates. Filtering parameters allowed maximum per-individual and per-population missing data and minimum minor allele frequency (MAF) filters. The included maximum missing data and minimum MAF filter proportions are shown in the plot title (missInd=per-individual, Pop=per-population filtering, MAF=MAF filter, P=perplexity). Each barplot on the page represents one UML algorithm: random forest, visualized with classical and isotonic multidimensional scaling (cMDS and isoMDS) ordination. t-SNE=t-distributed stochastic neighbor embedding with perplexity=15 (P15), and VAE=variational autoencoder. cMDS, isoMDS, and t-SNE were clustered with the partition around medoids (PAM) and hierarchical clustering (H) algorithms, and optimal *K* was chosen using PAM with the gap statistic (GS), H with the highest mean silhouette width (HMSW), and PAM with the HMSW. For VAE, clustering was performed, and optimal *K* chosen, with DBSCAN. *Terrapene* populations are separated by the horizontal white strips. Population IDs (blue left-aligned strip) include the ON=Ornate (*T. ornata ornata*), DS=Desert (*T. o. luteola*), FL=Florida (*T. c. bauri*), CH=Coahuilan (*T. coahuila*), GUMS=Gulf Coast (*T. carolina major*) from Mississippi, GUFL=Gulf Coast from the Florida Panhandle, EA=Woodland (*T. c. carolina*), MX=Mexican (*T. mexicana mexicana*), and TT=Three-toed (*T. m. triunguis*).
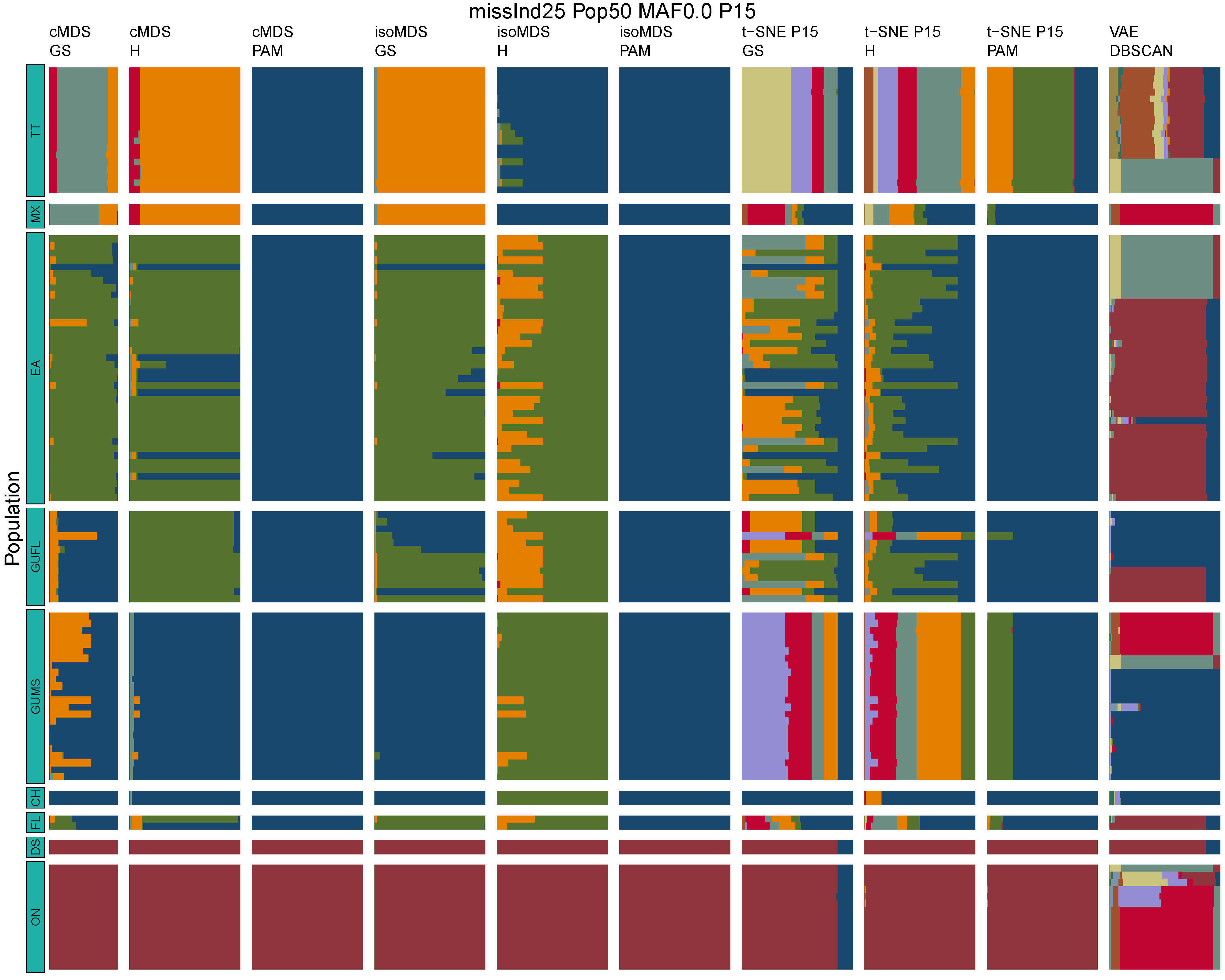

**Figure B5:** *Terrapene* unsupervised machine learning (UML) barplots depicting assignment proportions (x axis) among 100 replicates. Filtering parameters allowed maximum per-individual and per-population missing data and minimum minor allele frequency (MAF) filters. The included maximum missing data and minimum MAF filter proportions are shown in the plot title (missInd=per-individual, Pop=per-population filtering, MAF=MAF filter, P=perplexity). Each barplot on the page represents one UML algorithm: random forest, visualized with classical and isotonic multidimensional scaling (cMDS and isoMDS) ordination. t-SNE=t-distributed stochastic neighbor embedding with perplexity=15 (P15), and VAE=variational autoencoder. cMDS, isoMDS, and t-SNE were clustered with the partition around medoids (PAM) and hierarchical clustering (H) algorithms, and optimal *K* was chosen using PAM with the gap statistic (GS), H with the highest mean silhouette width (HMSW), and PAM with the HMSW. For VAE, clustering was performed, and optimal *K* chosen, with DBSCAN. *Terrapene* populations are separated by the horizontal white strips. Population IDs (blue left-aligned strip) include the ON=Ornate (*T. ornata ornata*), DS=Desert (*T. o. luteola*), FL=Florida (*T. c. bauri*), CH=Coahuilan (*T. coahuila*), GUMS=Gulf Coast (*T. carolina major*) from Mississippi, GUFL=Gulf Coast from the Florida Panhandle, EA=Woodland (*T. c. carolina*), MX=Mexican (*T. mexicana mexicana*), and TT=Three-toed (*T. m. triunguis*).
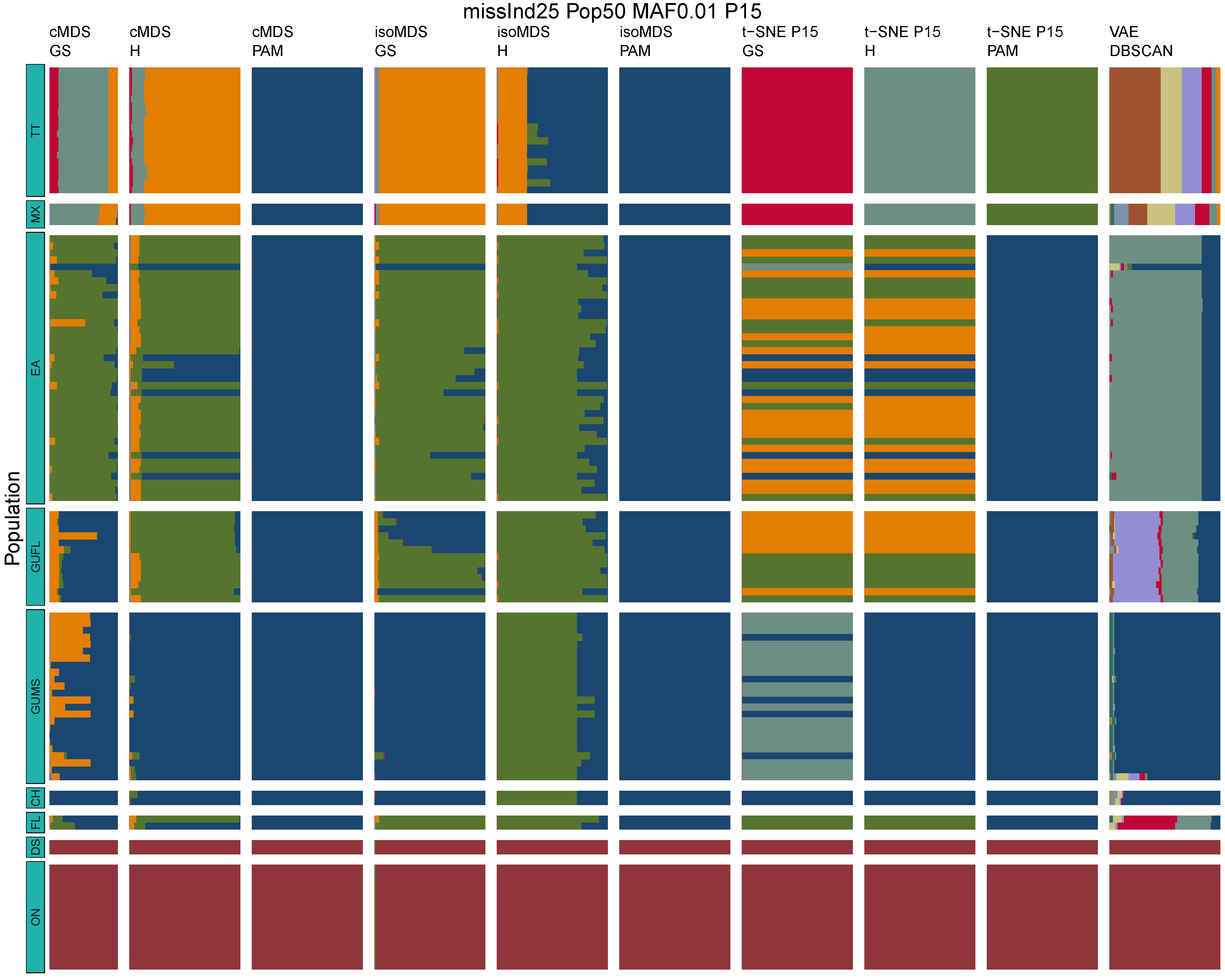

**Figure B6:** *Terrapene* unsupervised machine learning (UML) barplots depicting assignment proportions (x axis) among 100 replicates. Filtering parameters allowed maximum per-individual and per-population missing data and minimum minor allele frequency (MAF) filters. The included maximum missing data and minimum MAF filter proportions are shown in the plot title (missInd=per-individual, Pop=per-population filtering, MAF=MAF filter, P=perplexity). Each barplot on the page represents one UML algorithm: random forest, visualized with classical and isotonic multidimensional scaling (cMDS and isoMDS) ordination. t-SNE=t-distributed stochastic neighbor embedding with perplexity=15 (P15), and VAE=variational autoencoder. cMDS, isoMDS, and t-SNE were clustered with the partition around medoids (PAM) and hierarchical clustering (H) algorithms, and optimal *K* was chosen using PAM with the gap statistic (GS), H with the highest mean silhouette width (HMSW), and PAM with the HMSW. For VAE, clustering was performed, and optimal *K* chosen, with DBSCAN. *Terrapene* populations are separated by the horizontal white strips. Population IDs (blue left-aligned strip) include the ON=Ornate (*T. ornata ornata*), DS=Desert (*T. o. luteola*), FL=Florida (*T. c. bauri*), CH=Coahuilan (*T. coahuila*), GUMS=Gulf Coast (*T. carolina major*) from Mississippi, GUFL=Gulf Coast from the Florida Panhandle, EA=Woodland (*T. c. carolina*), MX=Mexican (*T. mexicana mexicana*), and TT=Three-toed (*T. m. triunguis*).

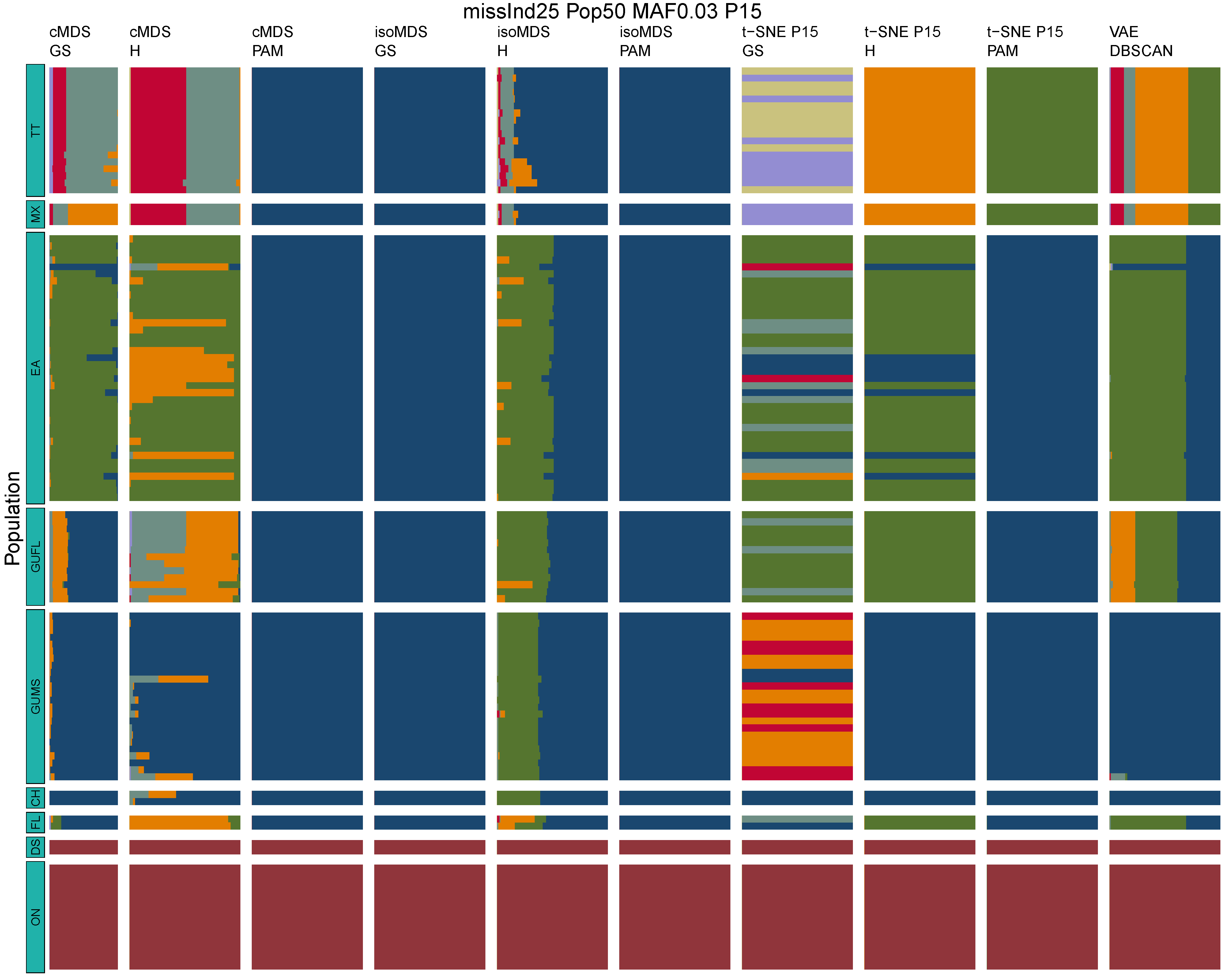

**Figure B7:** *Terrapene* unsupervised machine learning (UML) barplots depicting assignment proportions (x axis) among 100 replicates. Filtering parameters allowed maximum per-individual and per-population missing data and minimum minor allele frequency (MAF) filters. The included maximum missing data and minimum MAF filter proportions are shown in the plot title (missInd=per-individual, Pop=per-population filtering, MAF=MAF filter, P=perplexity). Each barplot on the page represents one UML algorithm: random forest, visualized with classical and isotonic multidimensional scaling (cMDS and isoMDS) ordination. t-SNE=t-distributed stochastic neighbor embedding with perplexity=15 (P15), and VAE=variational autoencoder. cMDS, isoMDS, and t-SNE were clustered with the partition around medoids (PAM) and hierarchical clustering (H) algorithms, and optimal *K* was chosen using PAM with the gap statistic (GS), H with the highest mean silhouette width (HMSW), and PAM with the HMSW. For VAE, clustering was performed, and optimal *K* chosen, with DBSCAN. *Terrapene* populations are separated by the horizontal white strips. Population IDs (blue left-aligned strip) include the ON=Ornate (*T. ornata ornata*), DS=Desert (*T. o. luteola*), FL=Florida (*T. c. bauri*), CH=Coahuilan (*T. coahuila*), GUMS=Gulf Coast (*T. carolina major*) from Mississippi, GUFL=Gulf Coast from the Florida Panhandle, EA=Woodland (*T. c. carolina*), MX=Mexican (*T. mexicana mexicana*), and TT=Three-toed (*T. m. triunguis*).

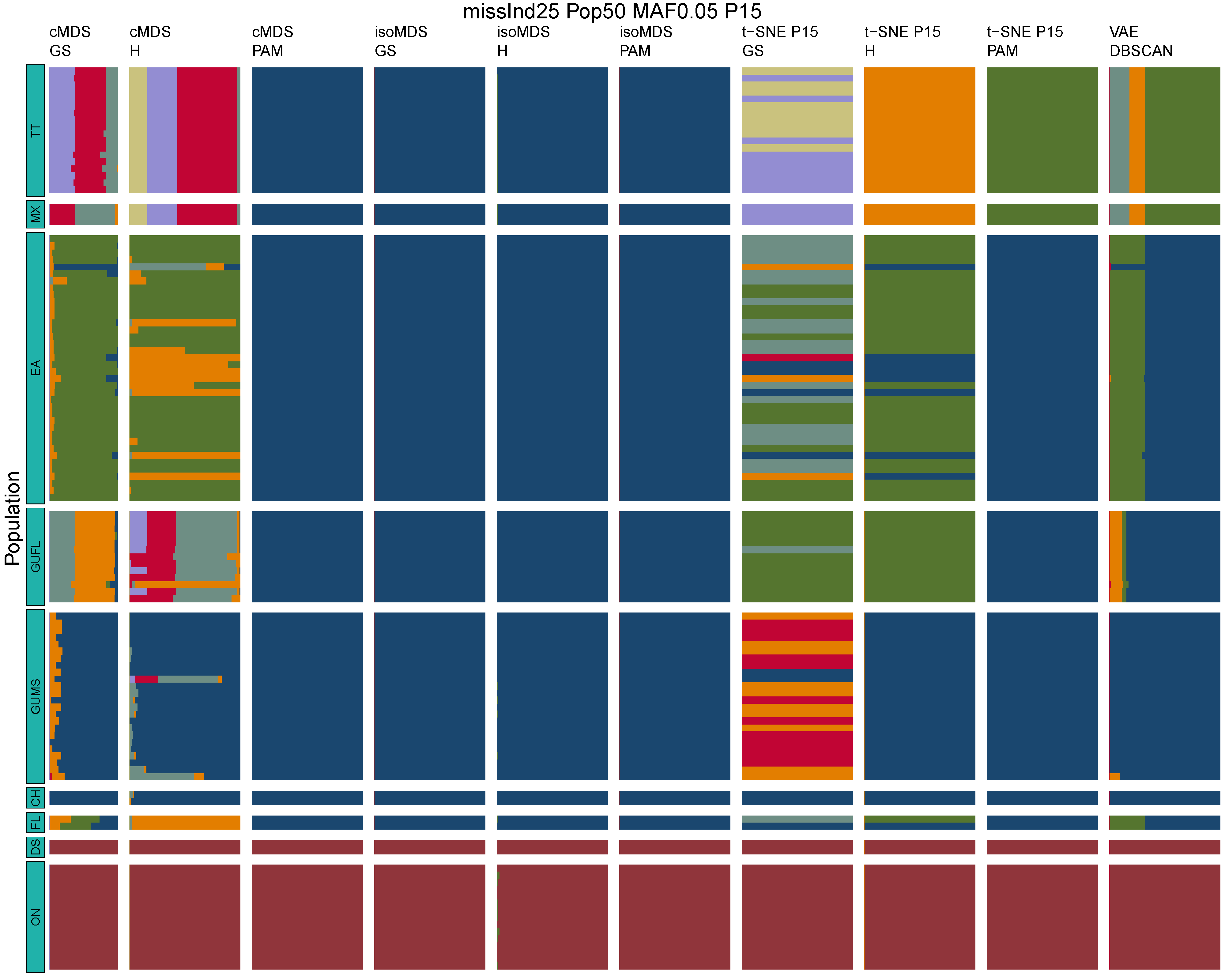

**Figure B8:** *Terrapene* unsupervised machine learning (UML) barplots depicting assignment proportions (x axis) among 100 replicates. Filtering parameters allowed maximum per-individual and per-population missing data and minimum minor allele frequency (MAF) filters. The included maximum missing data and minimum MAF filter proportions are shown in the plot title (missInd=per-individual, Pop=per-population filtering, MAF=MAF filter, P=perplexity). Each barplot on the page represents one UML algorithm: random forest, visualized with classical and isotonic multidimensional scaling (cMDS and isoMDS) ordination. t-SNE=t-distributed stochastic neighbor embedding with perplexity=15 (P15), and VAE=variational autoencoder. cMDS, isoMDS, and t-SNE were clustered with the partition around medoids (PAM) and hierarchical clustering (H) algorithms, and optimal *K* was chosen using PAM with the gap statistic (GS), H with the highest mean silhouette width (HMSW), and PAM with the HMSW. For VAE, clustering was performed, and optimal *K* chosen, with DBSCAN. *Terrapene* populations are separated by the horizontal white strips. Population IDs (blue left-aligned strip) include the ON=Ornate (*T. ornata ornata*), DS=Desert (*T. o. luteola*), FL=Florida (*T. c. bauri*), CH=Coahuilan (*T. coahuila*), GUMS=Gulf Coast (*T. carolina major*) from Mississippi, GUFL=Gulf Coast from the Florida Panhandle, EA=Woodland (*T. c. carolina*), MX=Mexican (*T. mexicana mexicana*), and TT=Three-toed (*T. m. triunguis*).
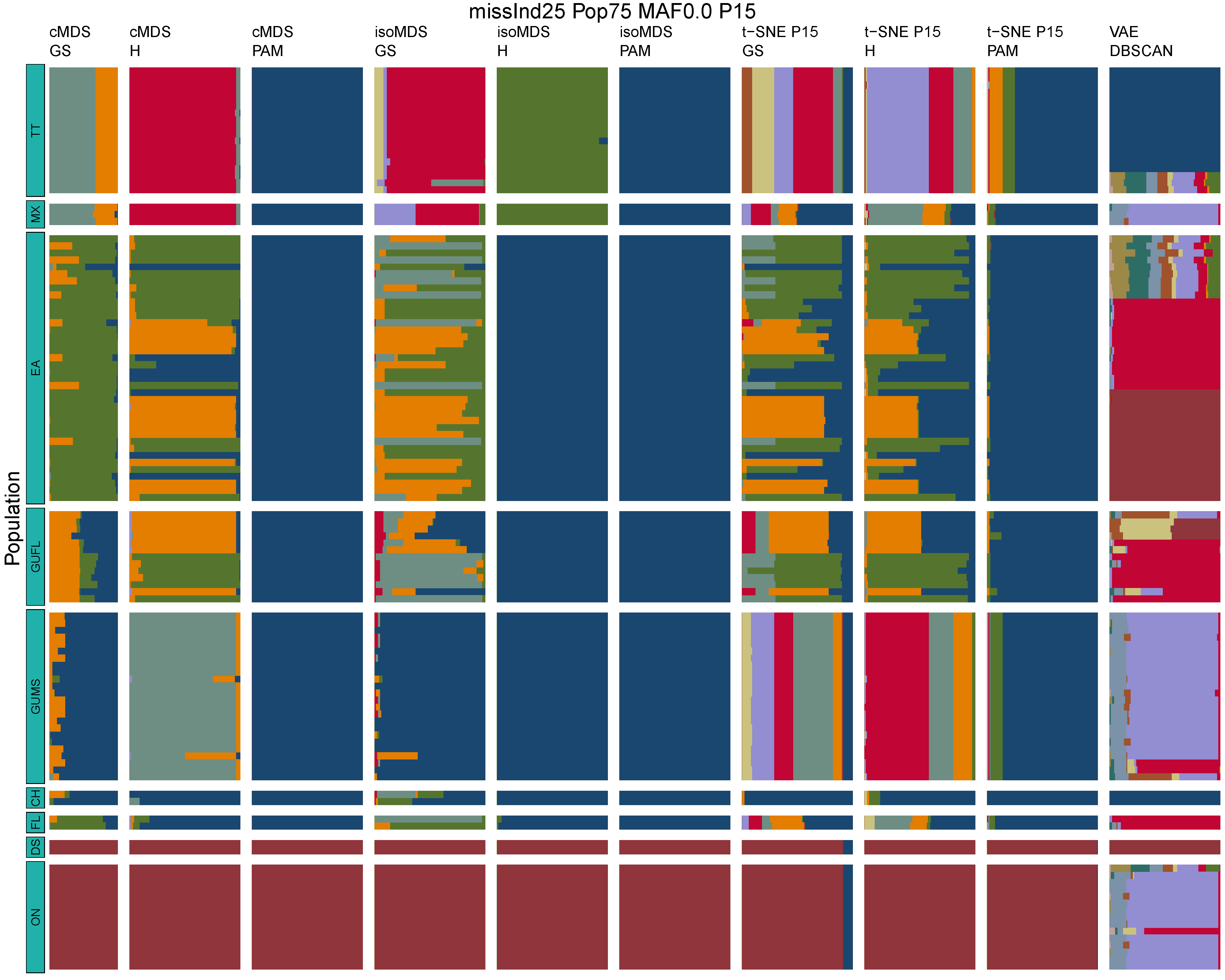

**Figure B9:** *Terrapene* unsupervised machine learning (UML) barplots depicting assignment proportions (x axis) among 100 replicates. Filtering parameters allowed maximum per-individual and per-population missing data and minimum minor allele frequency (MAF) filters. The included maximum missing data and minimum MAF filter proportions are shown in the plot title (missInd=per-individual, Pop=per-population filtering, MAF=MAF filter, P=perplexity). Each barplot on the page represents one UML algorithm: random forest, visualized with classical and isotonic multidimensional scaling (cMDS and isoMDS) ordination. t-SNE=t-distributed stochastic neighbor embedding with perplexity=15 (P15), and VAE=variational autoencoder. cMDS, isoMDS, and t-SNE were clustered with the partition around medoids (PAM) and hierarchical clustering (H) algorithms, and optimal *K* was chosen using PAM with the gap statistic (GS), H with the highest mean silhouette width (HMSW), and PAM with the HMSW. For VAE, clustering was performed, and optimal *K* chosen, with DBSCAN. *Terrapene* populations are separated by the horizontal white strips. Population IDs (blue left-aligned strip) include the ON=Ornate (*T. ornata ornata*), DS=Desert (*T. o. luteola*), FL=Florida (*T. c. bauri*), CH=Coahuilan (*T. coahuila*), GUMS=Gulf Coast (*T. carolina major*) from Mississippi, GUFL=Gulf Coast from the Florida Panhandle, EA=Woodland (*T. c. carolina*), MX=Mexican (*T. mexicana mexicana*), and TT=Three-toed (*T. m. triunguis*).
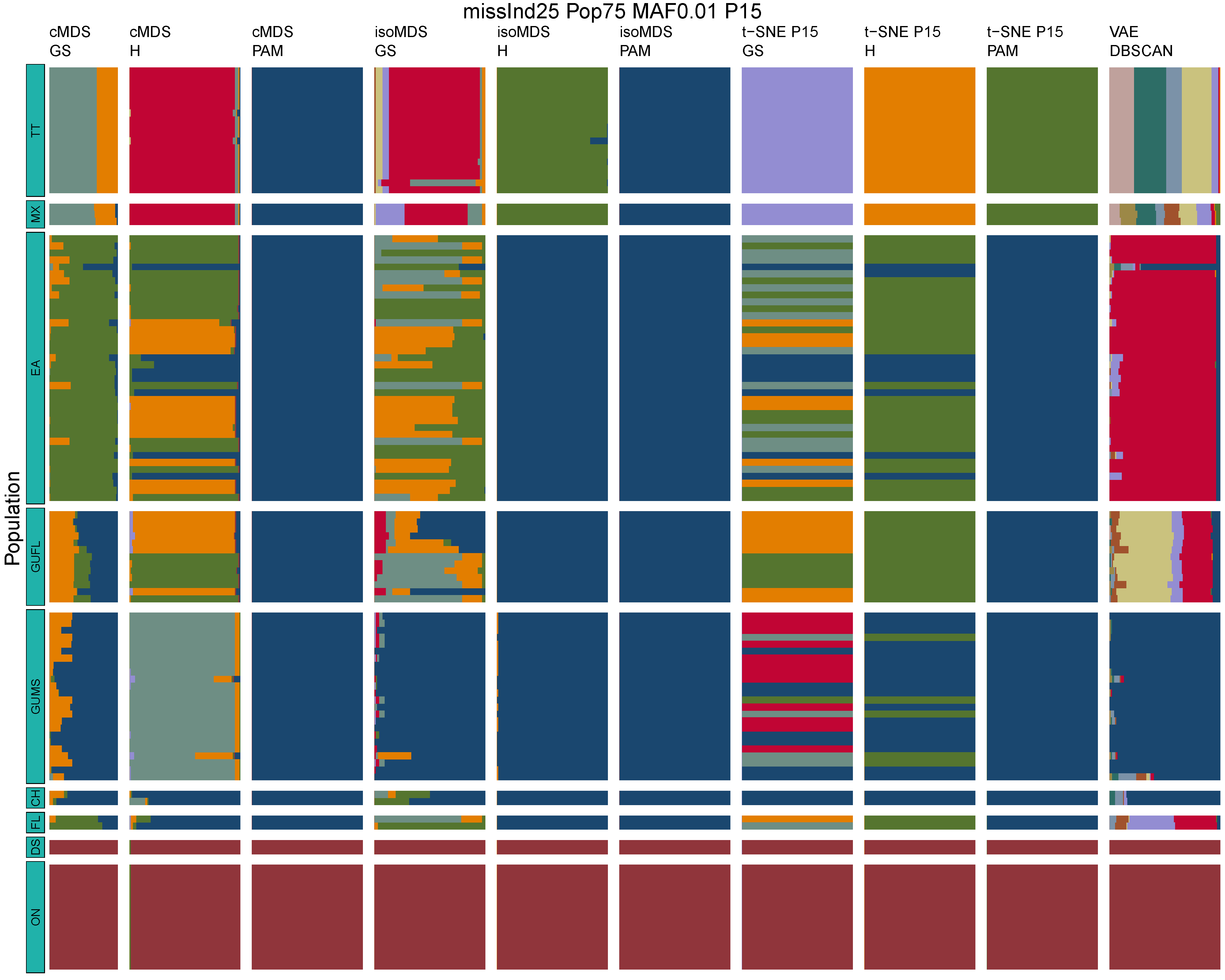

**Figure B10:** *Terrapene* unsupervised machine learning (UML) barplots depicting assignment proportions (x axis) among 100 replicates. Filtering parameters allowed maximum per-individual and per-population missing data and minimum minor allele frequency (MAF) filters. The included maximum missing data and minimum MAF filter proportions are shown in the plot title (missInd=per-individual, Pop=per-population filtering, MAF=MAF filter, P=perplexity). Each barplot on the page represents one UML algorithm: random forest, visualized with classical and isotonic multidimensional scaling (cMDS and isoMDS) ordination. t-SNE=t-distributed stochastic neighbor embedding with perplexity=15 (P15), and VAE=variational autoencoder. cMDS, isoMDS, and t-SNE were clustered with the partition around medoids (PAM) and hierarchical clustering (H) algorithms, and optimal *K* was chosen using PAM with the gap statistic (GS), H with the highest mean silhouette width (HMSW), and PAM with the HMSW. For VAE, clustering was performed, and optimal *K* chosen, with DBSCAN. *Terrapene* populations are separated by the horizontal white strips. Population IDs (blue left-aligned strip) include the ON=Ornate (*T. ornata ornata*), DS=Desert (*T. o. luteola*), FL=Florida (*T. c. bauri*), CH=Coahuilan (*T. coahuila*), GUMS=Gulf Coast (*T. carolina major*) from Mississippi, GUFL=Gulf Coast from the Florida Panhandle, EA=Woodland (*T. c. carolina*), MX=Mexican (*T. mexicana mexicana*), and TT=Three-toed (*T. m. triunguis*).
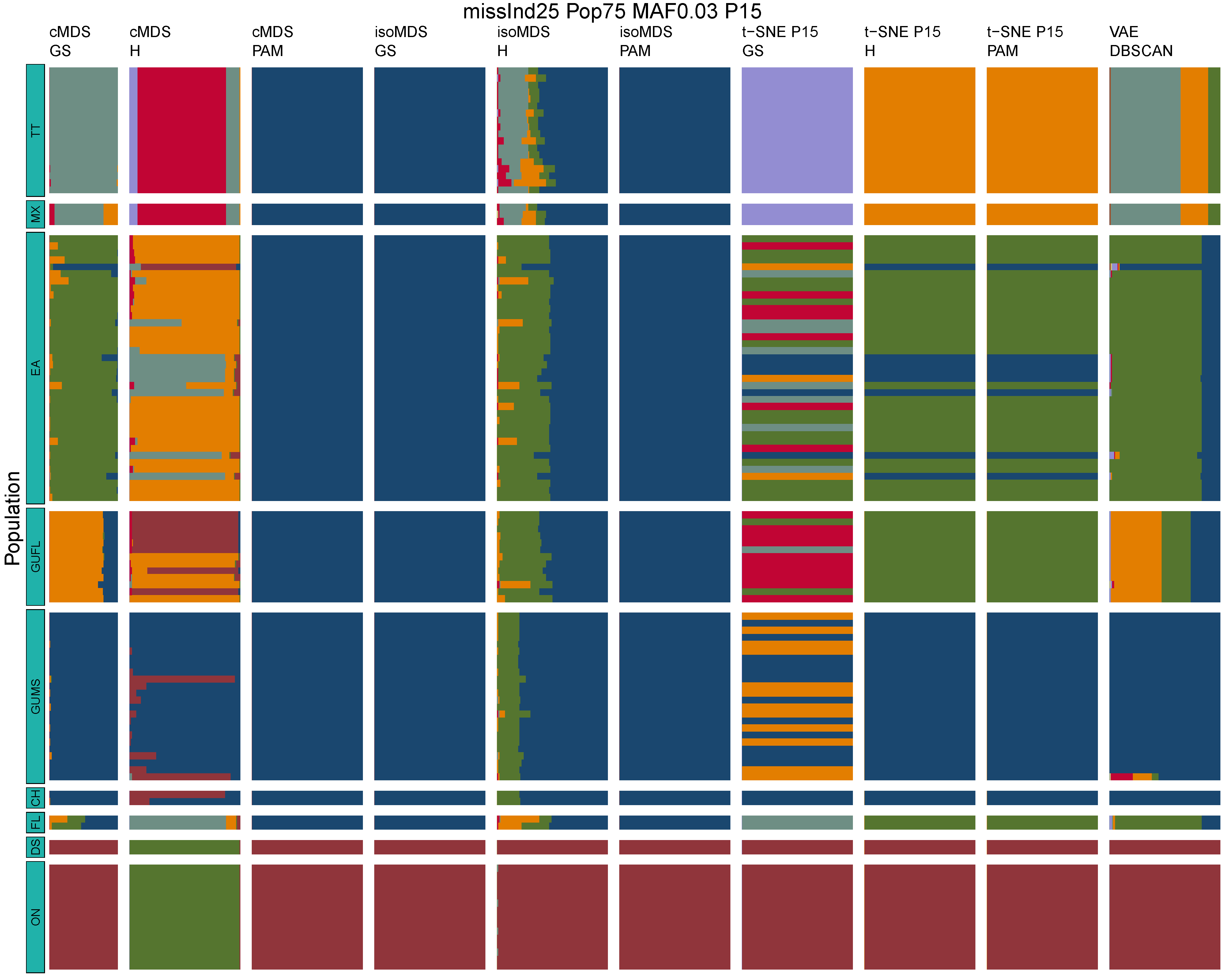

**Figure B11:** *Terrapene* unsupervised machine learning (UML) barplots depicting assignment proportions (x axis) among 100 replicates. Filtering parameters allowed maximum per-individual and per-population missing data and minimum minor allele frequency (MAF) filters. The included maximum missing data and minimum MAF filter proportions are shown in the plot title (missInd=per-individual, Pop=per-population filtering, MAF=MAF filter, P=perplexity). Each barplot on the page represents one UML algorithm: random forest, visualized with classical and isotonic multidimensional scaling (cMDS and isoMDS) ordination. t-SNE=t-distributed stochastic neighbor embedding with perplexity=15 (P15), and VAE=variational autoencoder. cMDS, isoMDS, and t-SNE were clustered with the partition around medoids (PAM) and hierarchical clustering (H) algorithms, and optimal *K* was chosen using PAM with the gap statistic (GS), H with the highest mean silhouette width (HMSW), and PAM with the HMSW. For VAE, clustering was performed, and optimal *K* chosen, with DBSCAN. *Terrapene* populations are separated by the horizontal white strips. Population IDs (blue left-aligned strip) include the ON=Ornate (*T. ornata ornata*), DS=Desert (*T. o. luteola*), FL=Florida (*T. c. bauri*), CH=Coahuilan (*T. coahuila*), GUMS=Gulf Coast (*T. carolina major*) from Mississippi, GUFL=Gulf Coast from the Florida Panhandle, EA=Woodland (*T. c. carolina*), MX=Mexican (*T. mexicana mexicana*), and TT=Three-toed (*T. m. triunguis*).
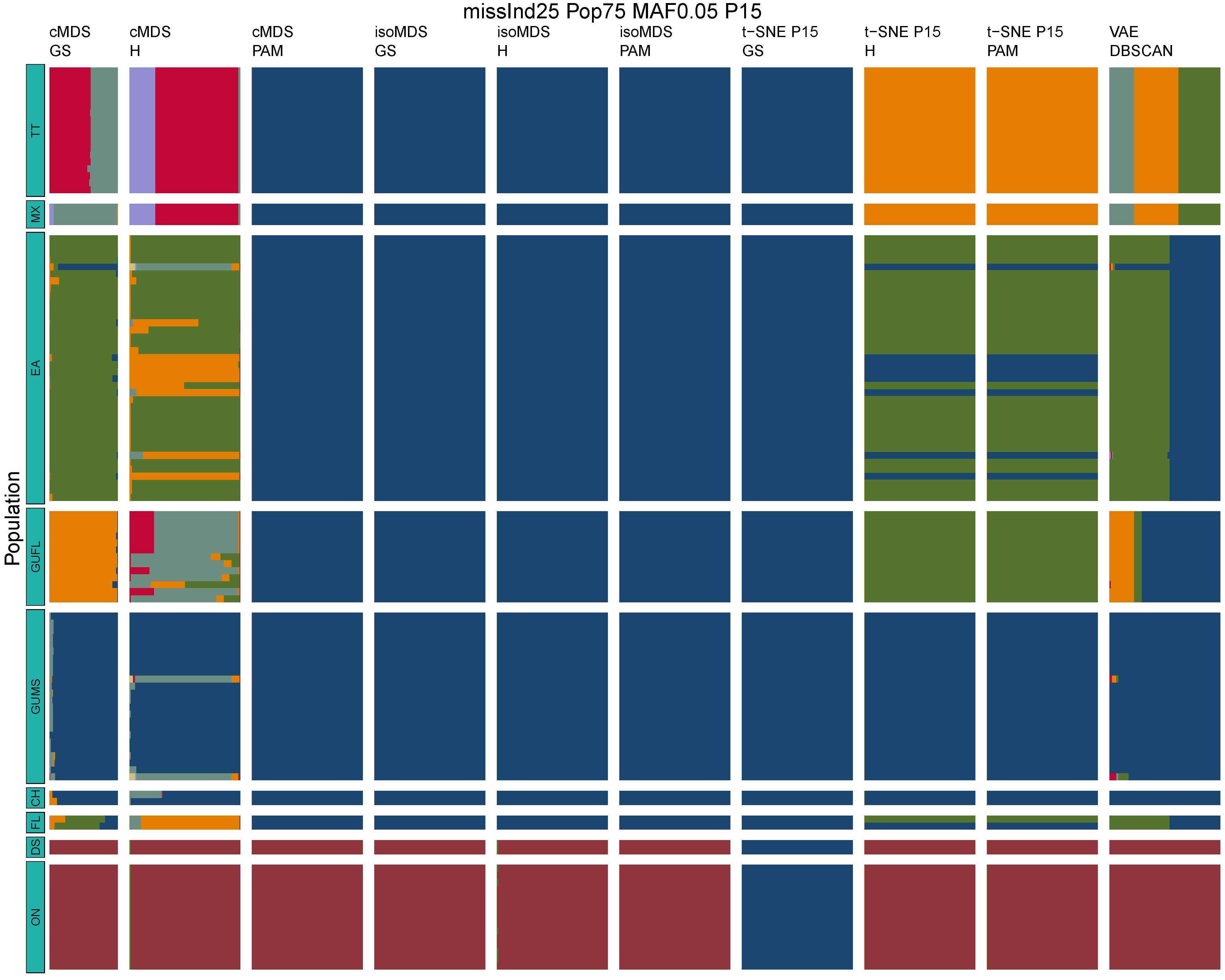

**Figure B12:** *Terrapene* unsupervised machine learning (UML) barplots depicting assignment proportions (x axis) among 100 replicates. Filtering parameters allowed maximum per-individual and per-population missing data and minimum minor allele frequency (MAF) filters. The included maximum missing data and minimum MAF filter proportions are shown in the plot title (missInd=per-individual, Pop=per-population filtering, MAF=MAF filter, P=perplexity). Each barplot on the page represents one UML algorithm: random forest, visualized with classical and isotonic multidimensional scaling (cMDS and isoMDS) ordination. t-SNE=t-distributed stochastic neighbor embedding with perplexity=15 (P15), and VAE=variational autoencoder. cMDS, isoMDS, and t-SNE were clustered with the partition around medoids (PAM) and hierarchical clustering (H) algorithms, and optimal *K* was chosen using PAM with the gap statistic (GS), H with the highest mean silhouette width (HMSW), and PAM with the HMSW. For VAE, clustering was performed, and optimal *K* chosen, with DBSCAN. *Terrapene* populations are separated by the horizontal white strips. Population IDs (blue left-aligned strip) include the ON=Ornate (*T. ornata ornata*), DS=Desert (*T. o. luteola*), FL=Florida (*T. c. bauri*), CH=Coahuilan (*T. coahuila*), GUMS=Gulf Coast (*T. carolina major*) from Mississippi, GUFL=Gulf Coast from the Florida Panhandle, EA=Woodland (*T. c. carolina*), MX=Mexican (*T. mexicana mexicana*), and TT=Three-toed (*T. m. triunguis*).
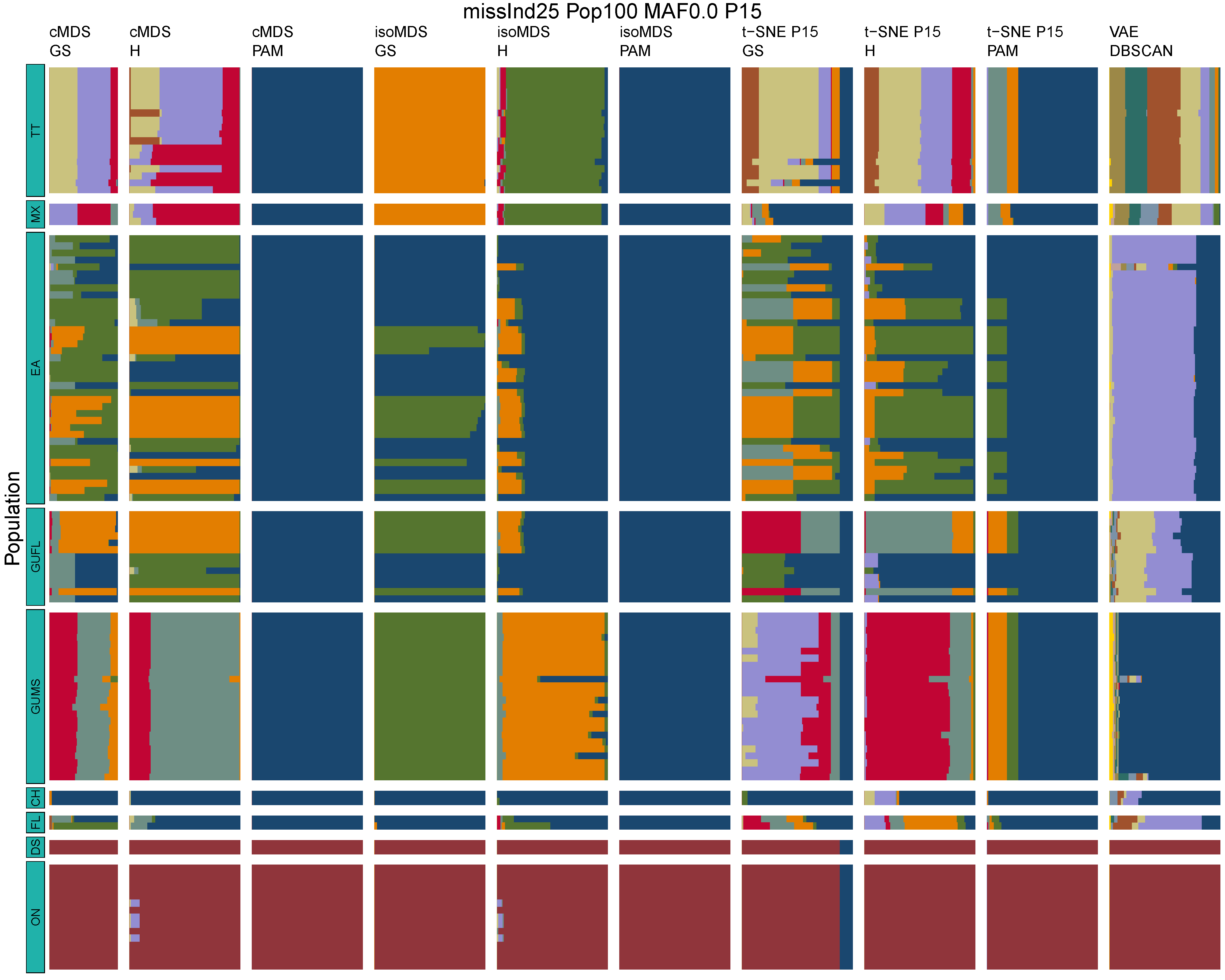

**Figure B13:** *Terrapene* unsupervised machine learning (UML) barplots depicting assignment proportions (x axis) among 100 replicates. Filtering parameters allowed maximum per-individual and per-population missing data and minimum minor allele frequency (MAF) filters. The included maximum missing data and minimum MAF filter proportions are shown in the plot title (missInd=per-individual, Pop=per-population filtering, MAF=MAF filter, P=perplexity). Each barplot on the page represents one UML algorithm: random forest, visualized with classical and isotonic multidimensional scaling (cMDS and isoMDS) ordination. t-SNE=t-distributed stochastic neighbor embedding with perplexity=15 (P15), and VAE=variational autoencoder. cMDS, isoMDS, and t-SNE were clustered with the partition around medoids (PAM) and hierarchical clustering (H) algorithms, and optimal *K* was chosen using PAM with the gap statistic (GS), H with the highest mean silhouette width (HMSW), and PAM with the HMSW. For VAE, clustering was performed, and optimal *K* chosen, with DBSCAN. *Terrapene* populations are separated by the horizontal white strips. Population IDs (blue left-aligned strip) include the ON=Ornate (*T. ornata ornata*), DS=Desert (*T. o. luteola*), FL=Florida (*T. c. bauri*), CH=Coahuilan (*T. coahuila*), GUMS=Gulf Coast (*T. carolina major*) from Mississippi, GUFL=Gulf Coast from the Florida Panhandle, EA=Woodland (*T. c. carolina*), MX=Mexican (*T. mexicana mexicana*), and TT=Three-toed (*T. m. triunguis*).
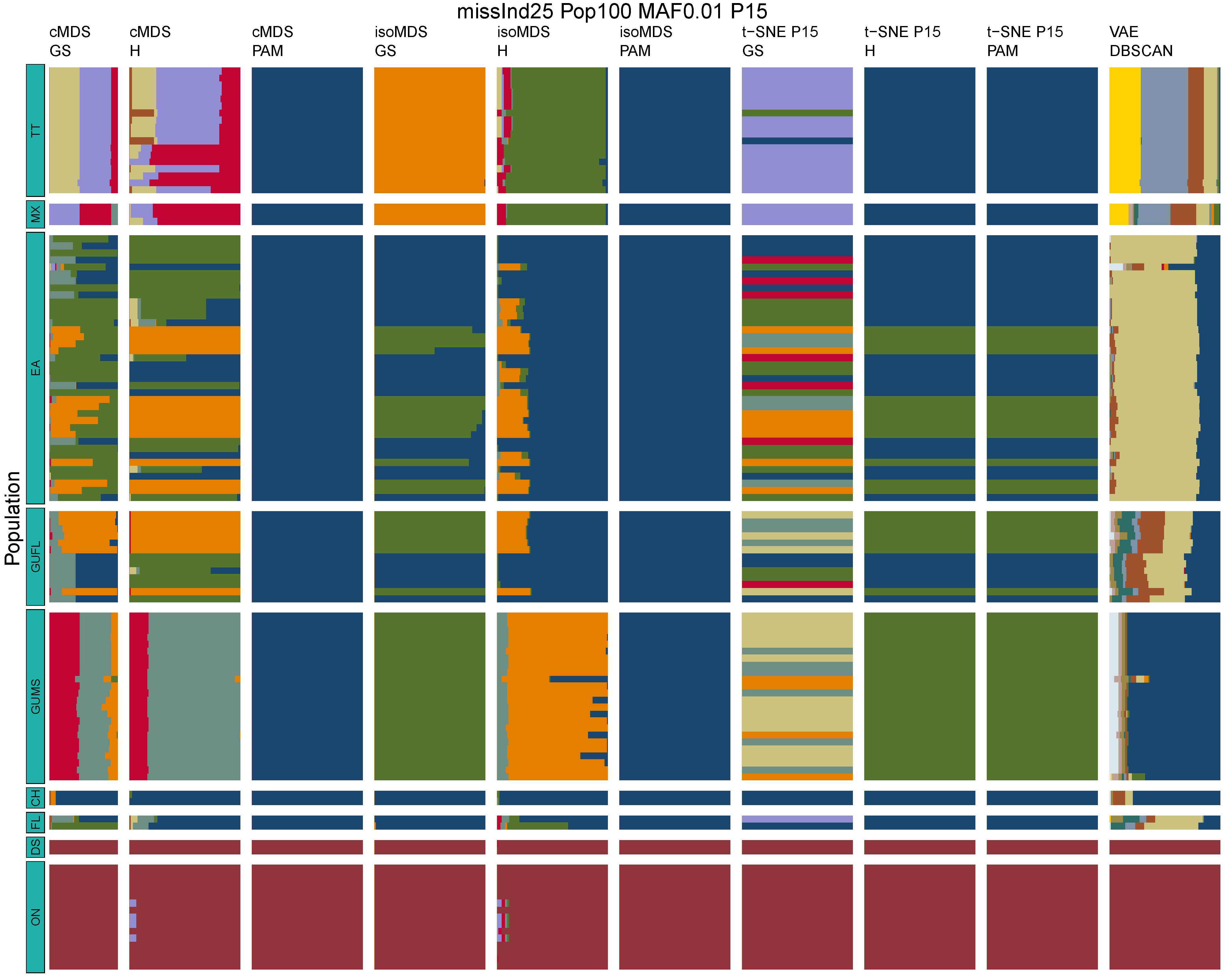

**Figure B14:** *Terrapene* unsupervised machine learning (UML) barplots depicting assignment proportions (x axis) among 100 replicates. Filtering parameters allowed maximum per-individual and per-population missing data and minimum minor allele frequency (MAF) filters. The included maximum missing data and minimum MAF filter proportions are shown in the plot title (missInd=per-individual, Pop=per-population filtering, MAF=MAF filter, P=perplexity). Each barplot on the page represents one UML algorithm: random forest, visualized with classical and isotonic multidimensional scaling (cMDS and isoMDS) ordination. t-SNE=t-distributed stochastic neighbor embedding with perplexity=15 (P15), and VAE=variational autoencoder. cMDS, isoMDS, and t-SNE were clustered with the partition around medoids (PAM) and hierarchical clustering (H) algorithms, and optimal *K* was chosen using PAM with the gap statistic (GS), H with the highest mean silhouette width (HMSW), and PAM with the HMSW. For VAE, clustering was performed, and optimal *K* chosen, with DBSCAN. *Terrapene* populations are separated by the horizontal white strips. Population IDs (blue left-aligned strip) include the ON=Ornate (*T. ornata ornata*), DS=Desert (*T. o. luteola*), FL=Florida (*T. c. bauri*), CH=Coahuilan (*T. coahuila*), GUMS=Gulf Coast (*T. carolina major*) from Mississippi, GUFL=Gulf Coast from the Florida Panhandle, EA=Woodland (*T. c. carolina*), MX=Mexican (*T. mexicana mexicana*), and TT=Three-toed (*T. m. triunguis*).
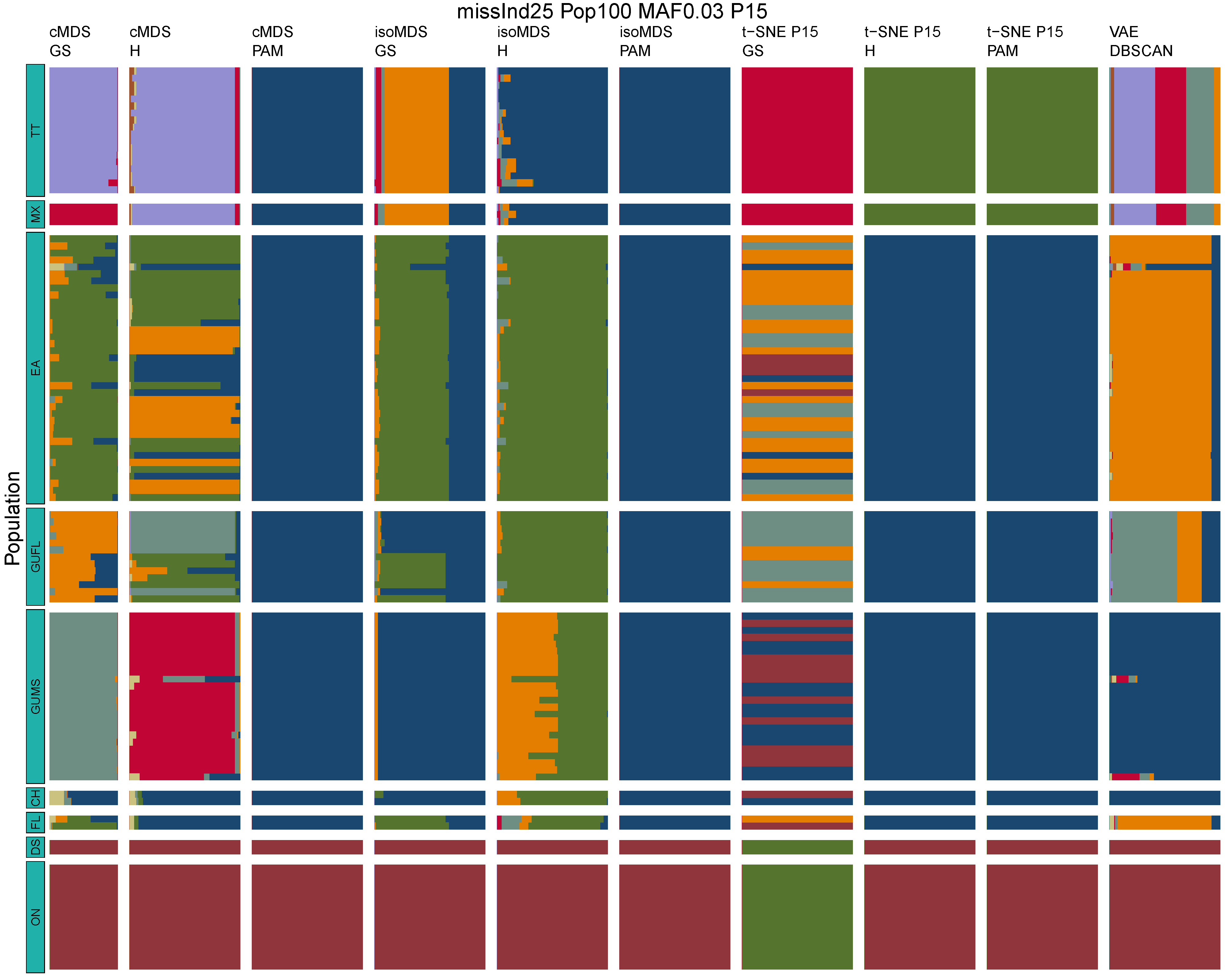

**Figure B15:** *Terrapene* unsupervised machine learning (UML) barplots depicting assignment proportions (x axis) among 100 replicates. Filtering parameters allowed maximum per-individual and per-population missing data and minimum minor allele frequency (MAF) filters. The included maximum missing data and minimum MAF filter proportions are shown in the plot title (missInd=per-individual, Pop=per-population filtering, MAF=MAF filter, P=perplexity). Each barplot on the page represents one UML algorithm: random forest, visualized with classical and isotonic multidimensional scaling (cMDS and isoMDS) ordination. t-SNE=t-distributed stochastic neighbor embedding with perplexity=15 (P15), and VAE=variational autoencoder. cMDS, isoMDS, and t-SNE were clustered with the partition around medoids (PAM) and hierarchical clustering (H) algorithms, and optimal *K* was chosen using PAM with the gap statistic (GS), H with the highest mean silhouette width (HMSW), and PAM with the HMSW. For VAE, clustering was performed, and optimal *K* chosen, with DBSCAN. *Terrapene* populations are separated by the horizontal white strips. Population IDs (blue left-aligned strip) include the ON=Ornate (*T. ornata ornata*), DS=Desert (*T. o. luteola*), FL=Florida (*T. c. bauri*), CH=Coahuilan (*T. coahuila*), GUMS=Gulf Coast (*T. carolina major*) from Mississippi, GUFL=Gulf Coast from the Florida Panhandle, EA=Woodland (*T. c. carolina*), MX=Mexican (*T. mexicana mexicana*), and TT=Three-toed (*T. m. triunguis*).
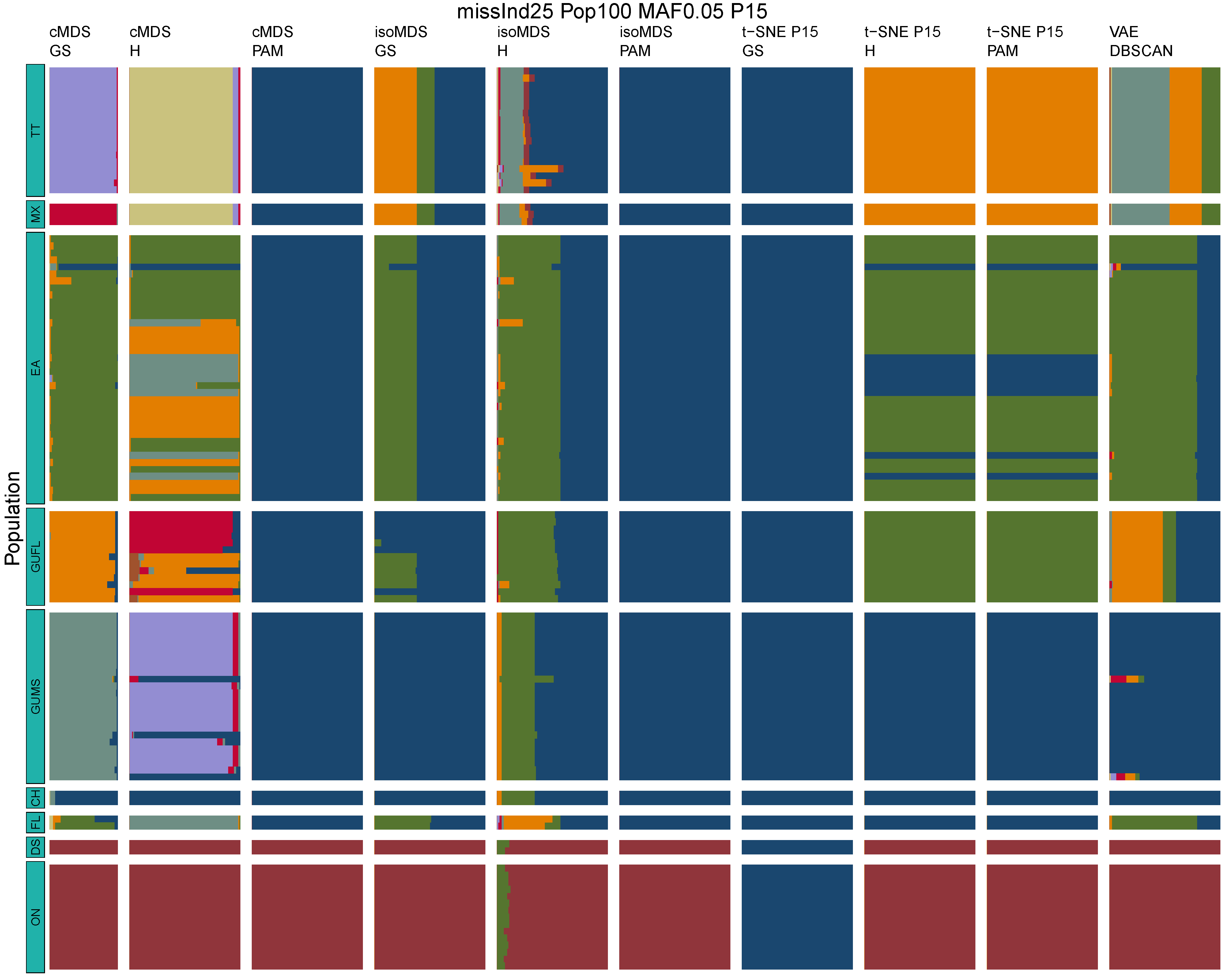

**Figure B16:** *Terrapene* unsupervised machine learning (UML) barplots depicting assignment proportions (x axis) among 100 replicates. Filtering parameters allowed maximum per-individual and per-population missing data and minimum minor allele frequency (MAF) filters. The included maximum missing data and minimum MAF filter proportions are shown in the plot title (missInd=per-individual, Pop=per-population filtering, MAF=MAF filter, P=perplexity). Each barplot on the page represents one UML algorithm: random forest, visualized with classical and isotonic multidimensional scaling (cMDS and isoMDS) ordination. t-SNE=t-distributed stochastic neighbor embedding with perplexity=15 (P15), and VAE=variational autoencoder. cMDS, isoMDS, and t-SNE were clustered with the partition around medoids (PAM) and hierarchical clustering (H) algorithms, and optimal *K* was chosen using PAM with the gap statistic (GS), H with the highest mean silhouette width (HMSW), and PAM with the HMSW. For VAE, clustering was performed, and optimal *K* chosen, with DBSCAN. *Terrapene* populations are separated by the horizontal white strips. Population IDs (blue left-aligned strip) include the ON=Ornate (*T. ornata ornata*), DS=Desert (*T. o. luteola*), FL=Florida (*T. c. bauri*), CH=Coahuilan (*T. coahuila*), GUMS=Gulf Coast (*T. carolina major*) from Mississippi, GUFL=Gulf Coast from the Florida Panhandle, EA=Woodland (*T. c. carolina*), MX=Mexican (*T. mexicana mexicana*), and TT=Three-toed (*T. m. triunguis*).
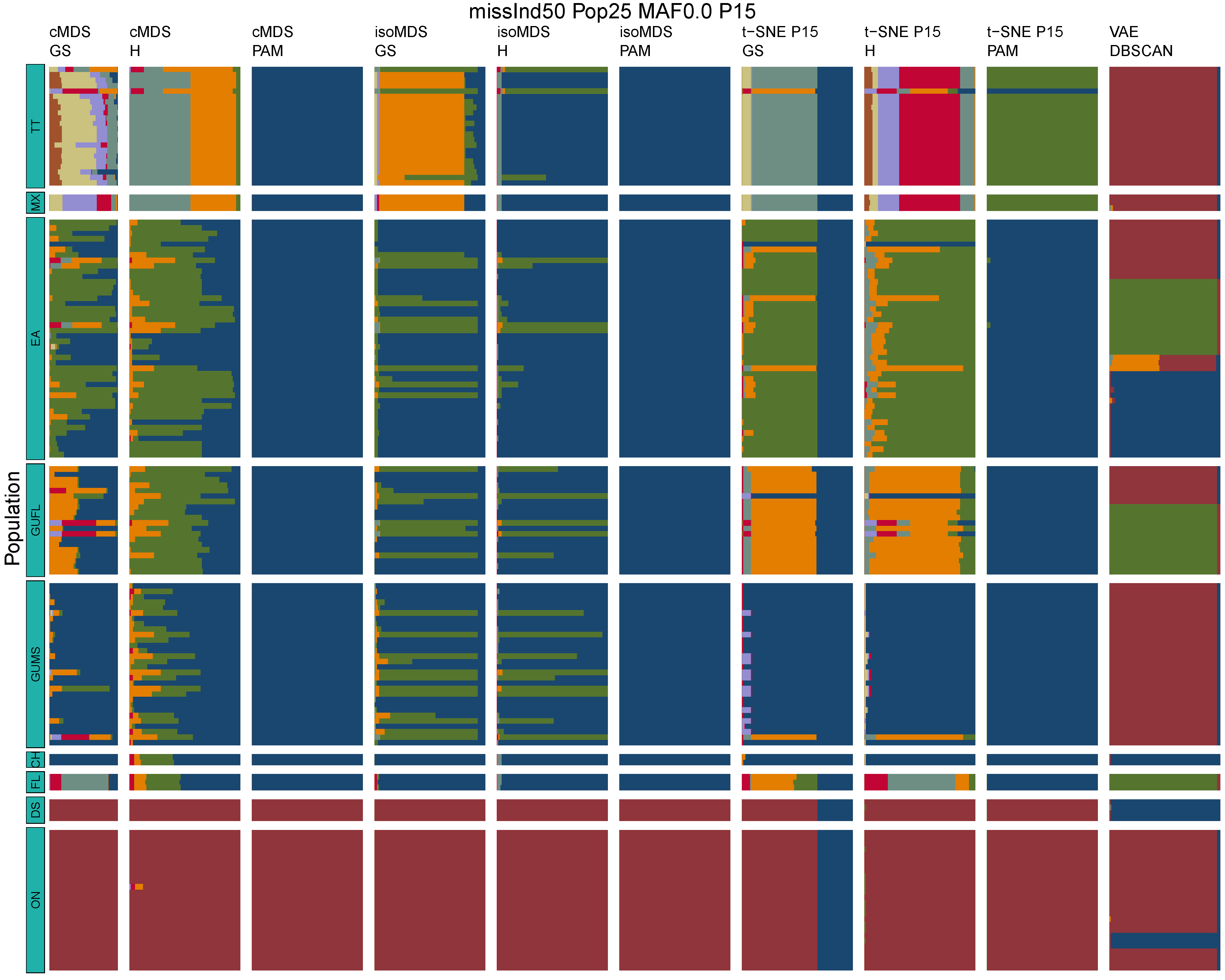

**Figure B17:** *Terrapene* unsupervised machine learning (UML) barplots depicting assignment proportions (x axis) among 100 replicates. Filtering parameters allowed maximum per-individual and per-population missing data and minimum minor allele frequency (MAF) filters. The included maximum missing data and minimum MAF filter proportions are shown in the plot title (missInd=per-individual, Pop=per-population filtering, MAF=MAF filter, P=perplexity). Each barplot on the page represents one UML algorithm: random forest, visualized with classical and isotonic multidimensional scaling (cMDS and isoMDS) ordination. t-SNE=t-distributed stochastic neighbor embedding with perplexity=15 (P15), and VAE=variational autoencoder. cMDS, isoMDS, and t-SNE were clustered with the partition around medoids (PAM) and hierarchical clustering (H) algorithms, and optimal *K* was chosen using PAM with the gap statistic (GS), H with the highest mean silhouette width (HMSW), and PAM with the HMSW. For VAE, clustering was performed, and optimal *K* chosen, with DBSCAN. *Terrapene* populations are separated by the horizontal white strips. Population IDs (blue left-aligned strip) include the ON=Ornate (*T. ornata ornata*), DS=Desert (*T. o. luteola*), FL=Florida (*T. c. bauri*), CH=Coahuilan (*T. coahuila*), GUMS=Gulf Coast (*T. carolina major*) from Mississippi, GUFL=Gulf Coast from the Florida Panhandle, EA=Woodland (*T. c. carolina*), MX=Mexican (*T. mexicana mexicana*), and TT=Three-toed (*T. m. triunguis*).
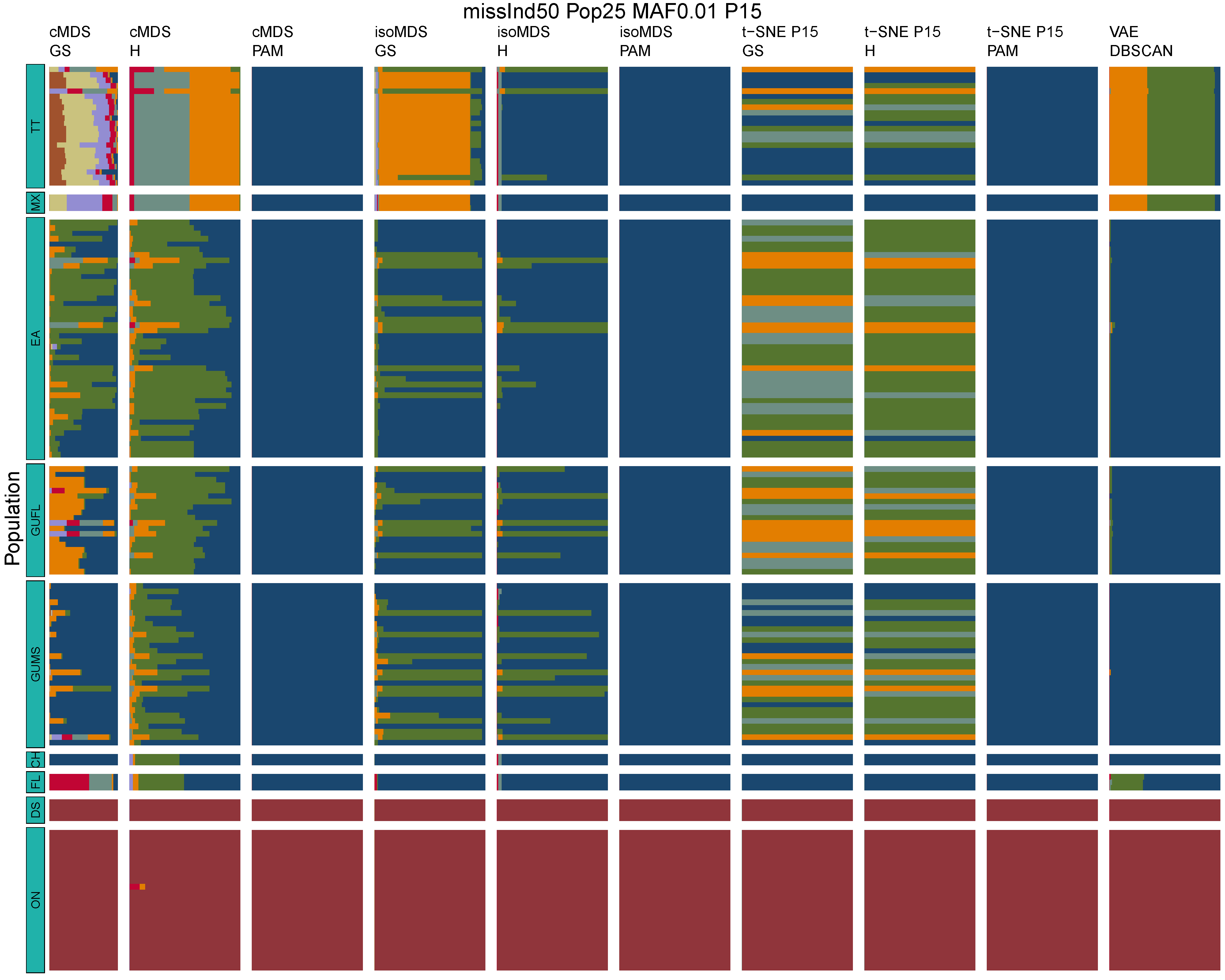

**Figure B18:** *Terrapene* unsupervised machine learning (UML) barplots depicting assignment proportions (x axis) among 100 replicates. Filtering parameters allowed maximum per-individual and per-population missing data and minimum minor allele frequency (MAF) filters. The included maximum missing data and minimum MAF filter proportions are shown in the plot title (missInd=per-individual, Pop=per-population filtering, MAF=MAF filter, P=perplexity). Each barplot on the page represents one UML algorithm: random forest, visualized with classical and isotonic multidimensional scaling (cMDS and isoMDS) ordination. t-SNE=t-distributed stochastic neighbor embedding with perplexity=15 (P15), and VAE=variational autoencoder. cMDS, isoMDS, and t-SNE were clustered with the partition around medoids (PAM) and hierarchical clustering (H) algorithms, and optimal *K* was chosen using PAM with the gap statistic (GS), H with the highest mean silhouette width (HMSW), and PAM with the HMSW. For VAE, clustering was performed, and optimal *K* chosen, with DBSCAN. *Terrapene* populations are separated by the horizontal white strips. Population IDs (blue left-aligned strip) include the ON=Ornate (*T. ornata ornata*), DS=Desert (*T. o. luteola*), FL=Florida (*T. c. bauri*), CH=Coahuilan (*T. coahuila*), GUMS=Gulf Coast (*T. carolina major*) from Mississippi, GUFL=Gulf Coast from the Florida Panhandle, EA=Woodland (*T. c. carolina*), MX=Mexican (*T. mexicana mexicana*), and TT=Three-toed (*T. m. triunguis*).
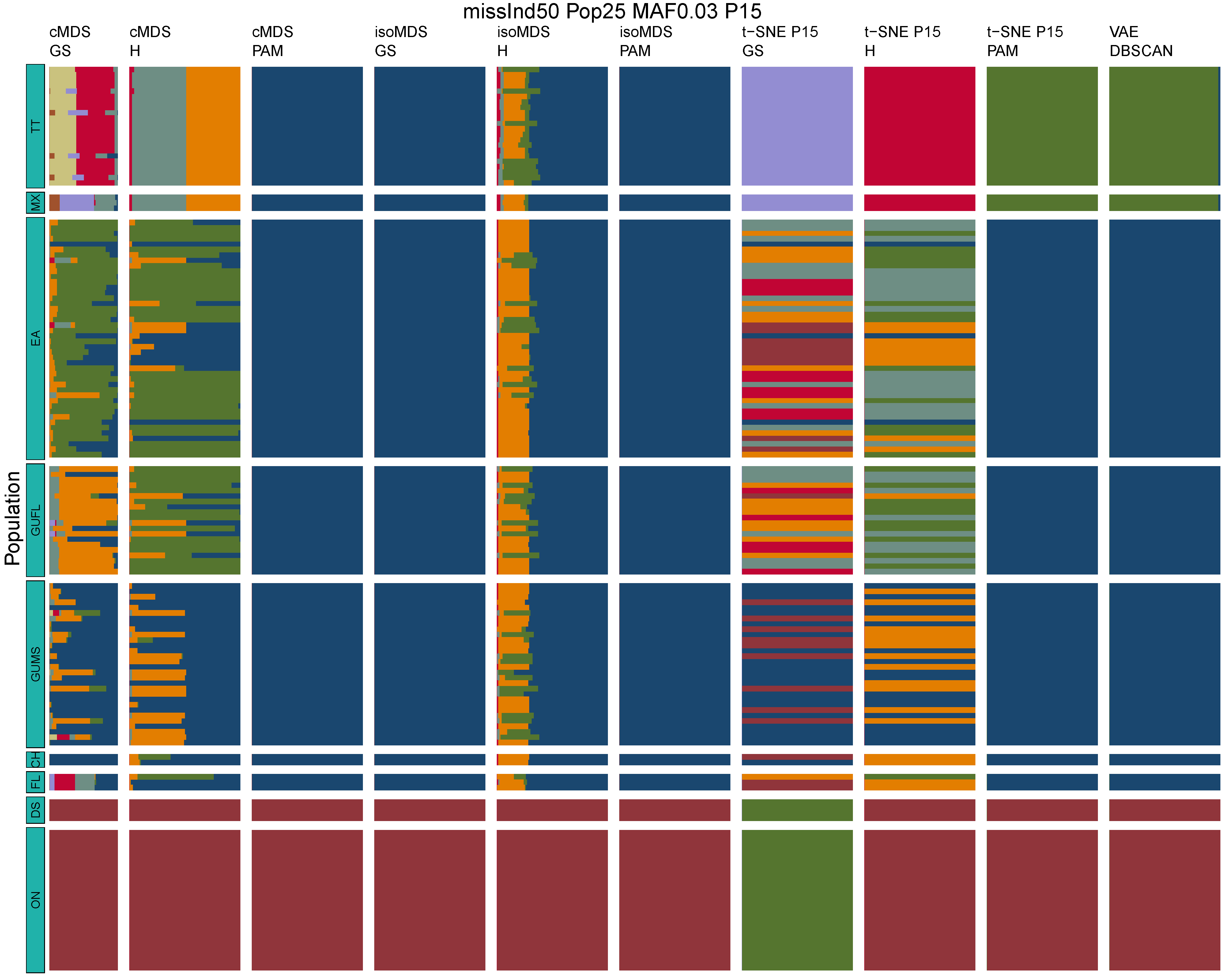

**Figure B19:** *Terrapene* unsupervised machine learning (UML) barplots depicting assignment proportions (x axis) among 100 replicates. Filtering parameters allowed maximum per-individual and per-population missing data and minimum minor allele frequency (MAF) filters. The included maximum missing data and minimum MAF filter proportions are shown in the plot title (missInd=per-individual, Pop=per-population filtering, MAF=MAF filter, P=perplexity). Each barplot on the page represents one UML algorithm: random forest, visualized with classical and isotonic multidimensional scaling (cMDS and isoMDS) ordination. t-SNE=t-distributed stochastic neighbor embedding with perplexity=15 (P15), and VAE=variational autoencoder. cMDS, isoMDS, and t-SNE were clustered with the partition around medoids (PAM) and hierarchical clustering (H) algorithms, and optimal *K* was chosen using PAM with the gap statistic (GS), H with the highest mean silhouette width (HMSW), and PAM with the HMSW. For VAE, clustering was performed, and optimal *K* chosen, with DBSCAN. *Terrapene* populations are separated by the horizontal white strips. Population IDs (blue left-aligned strip) include the ON=Ornate (*T. ornata ornata*), DS=Desert (*T. o. luteola*), FL=Florida (*T. c. bauri*), CH=Coahuilan (*T. coahuila*), GUMS=Gulf Coast (*T. carolina major*) from Mississippi, GUFL=Gulf Coast from the Florida Panhandle, EA=Woodland (*T. c. carolina*), MX=Mexican (*T. mexicana mexicana*), and TT=Three-toed (*T. m. triunguis*).
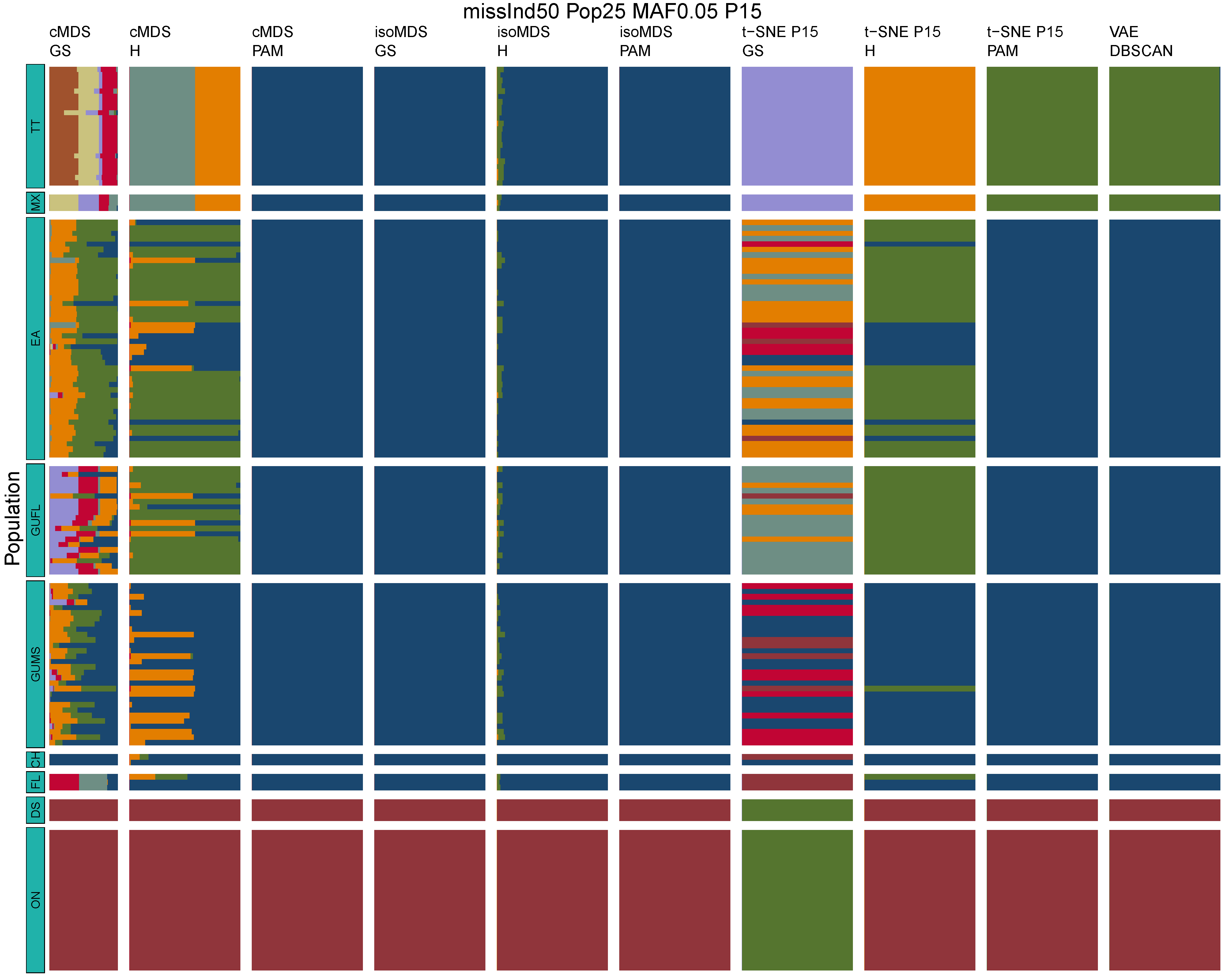

**Figure B20:** *Terrapene* unsupervised machine learning (UML) barplots depicting assignment proportions (x axis) among 100 replicates. Filtering parameters allowed maximum per-individual and per-population missing data and minimum minor allele frequency (MAF) filters. The included maximum missing data and minimum MAF filter proportions are shown in the plot title (missInd=per-individual, Pop=per-population filtering, MAF=MAF filter, P=perplexity). Each barplot on the page represents one UML algorithm: random forest, visualized with classical and isotonic multidimensional scaling (cMDS and isoMDS) ordination. t-SNE=t-distributed stochastic neighbor embedding with perplexity=15 (P15), and VAE=variational autoencoder. cMDS, isoMDS, and t-SNE were clustered with the partition around medoids (PAM) and hierarchical clustering (H) algorithms, and optimal *K* was chosen using PAM with the gap statistic (GS), H with the highest mean silhouette width (HMSW), and PAM with the HMSW. For VAE, clustering was performed, and optimal *K* chosen, with DBSCAN. *Terrapene* populations are separated by the horizontal white strips. Population IDs (blue left-aligned strip) include the ON=Ornate (*T. ornata ornata*), DS=Desert (*T. o. luteola*), FL=Florida (*T. c. bauri*), CH=Coahuilan (*T. coahuila*), GUMS=Gulf Coast (*T. carolina major*) from Mississippi, GUFL=Gulf Coast from the Florida Panhandle, EA=Woodland (*T. c. carolina*), MX=Mexican (*T. mexicana mexicana*), and TT=Three-toed (*T. m. triunguis*).
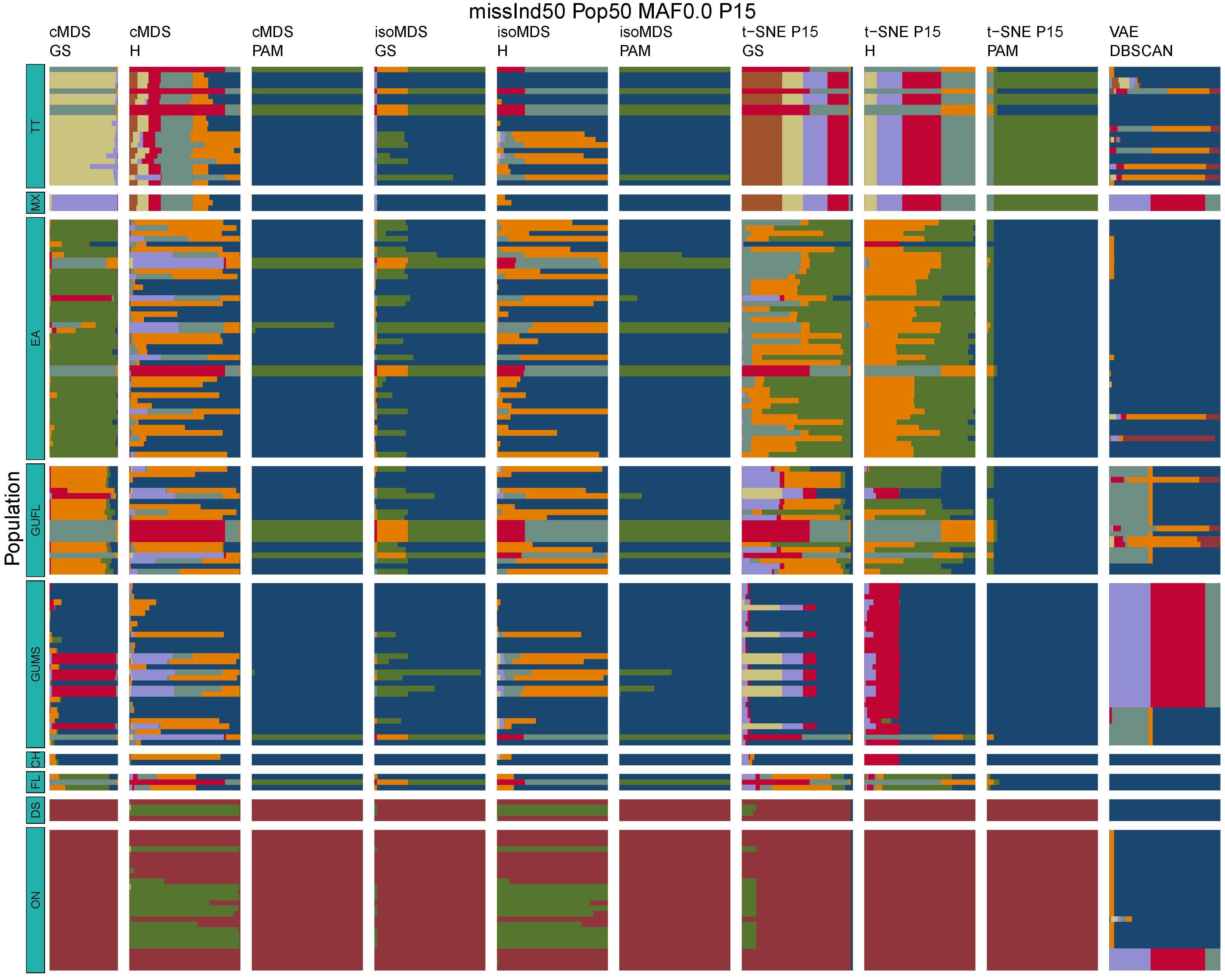

**Figure B21:** *Terrapene* unsupervised machine learning (UML) barplots depicting assignment proportions (x axis) among 100 replicates. Filtering parameters allowed maximum per-individual and per-population missing data and minimum minor allele frequency (MAF) filters. The included maximum missing data and minimum MAF filter proportions are shown in the plot title (missInd=per-individual, Pop=per-population filtering, MAF=MAF filter, P=perplexity). Each barplot on the page represents one UML algorithm: random forest, visualized with classical and isotonic multidimensional scaling (cMDS and isoMDS) ordination. t-SNE=t-distributed stochastic neighbor embedding with perplexity=15 (P15), and VAE=variational autoencoder. cMDS, isoMDS, and t-SNE were clustered with the partition around medoids (PAM) and hierarchical clustering (H) algorithms, and optimal *K* was chosen using PAM with the gap statistic (GS), H with the highest mean silhouette width (HMSW), and PAM with the HMSW. For VAE, clustering was performed, and optimal *K* chosen, with DBSCAN. *Terrapene* populations are separated by the horizontal white strips. Population IDs (blue left-aligned strip) include the ON=Ornate (*T. ornata ornata*), DS=Desert (*T. o. luteola*), FL=Florida (*T. c. bauri*), CH=Coahuilan (*T. coahuila*), GUMS=Gulf Coast (*T. carolina major*) from Mississippi, GUFL=Gulf Coast from the Florida Panhandle, EA=Woodland (*T. c. carolina*), MX=Mexican (*T. mexicana mexicana*), and TT=Three-toed (*T. m. triunguis*).
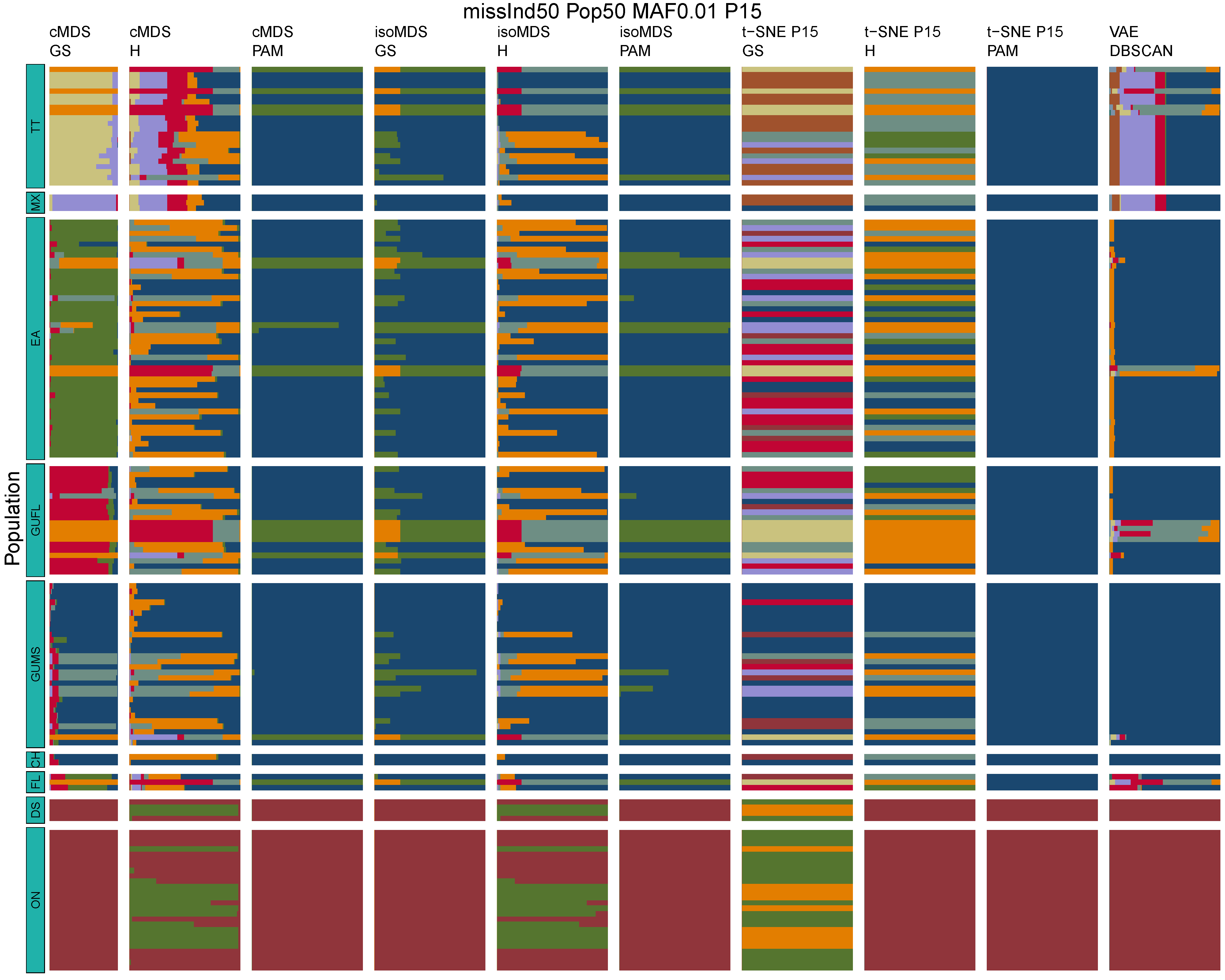

**Figure B22:** *Terrapene* unsupervised machine learning (UML) barplots depicting assignment proportions (x axis) among 100 replicates. Filtering parameters allowed maximum per-individual and per-population missing data and minimum minor allele frequency (MAF) filters. The included maximum missing data and minimum MAF filter proportions are shown in the plot title (missInd=per-individual, Pop=per-population filtering, MAF=MAF filter, P=perplexity). Each barplot on the page represents one UML algorithm: random forest, visualized with classical and isotonic multidimensional scaling (cMDS and isoMDS) ordination. t-SNE=t-distributed stochastic neighbor embedding with perplexity=15 (P15), and VAE=variational autoencoder. cMDS, isoMDS, and t-SNE were clustered with the partition around medoids (PAM) and hierarchical clustering (H) algorithms, and optimal *K* was chosen using PAM with the gap statistic (GS), H with the highest mean silhouette width (HMSW), and PAM with the HMSW. For VAE, clustering was performed, and optimal *K* chosen, with DBSCAN. *Terrapene* populations are separated by the horizontal white strips. Population IDs (blue left-aligned strip) include the ON=Ornate (*T. ornata ornata*), DS=Desert (*T. o. luteola*), FL=Florida (*T. c. bauri*), CH=Coahuilan (*T. coahuila*), GUMS=Gulf Coast (*T. carolina major*) from Mississippi, GUFL=Gulf Coast from the Florida Panhandle, EA=Woodland (*T. c. carolina*), MX=Mexican (*T. mexicana mexicana*), and TT=Three-toed (*T. m. triunguis*).
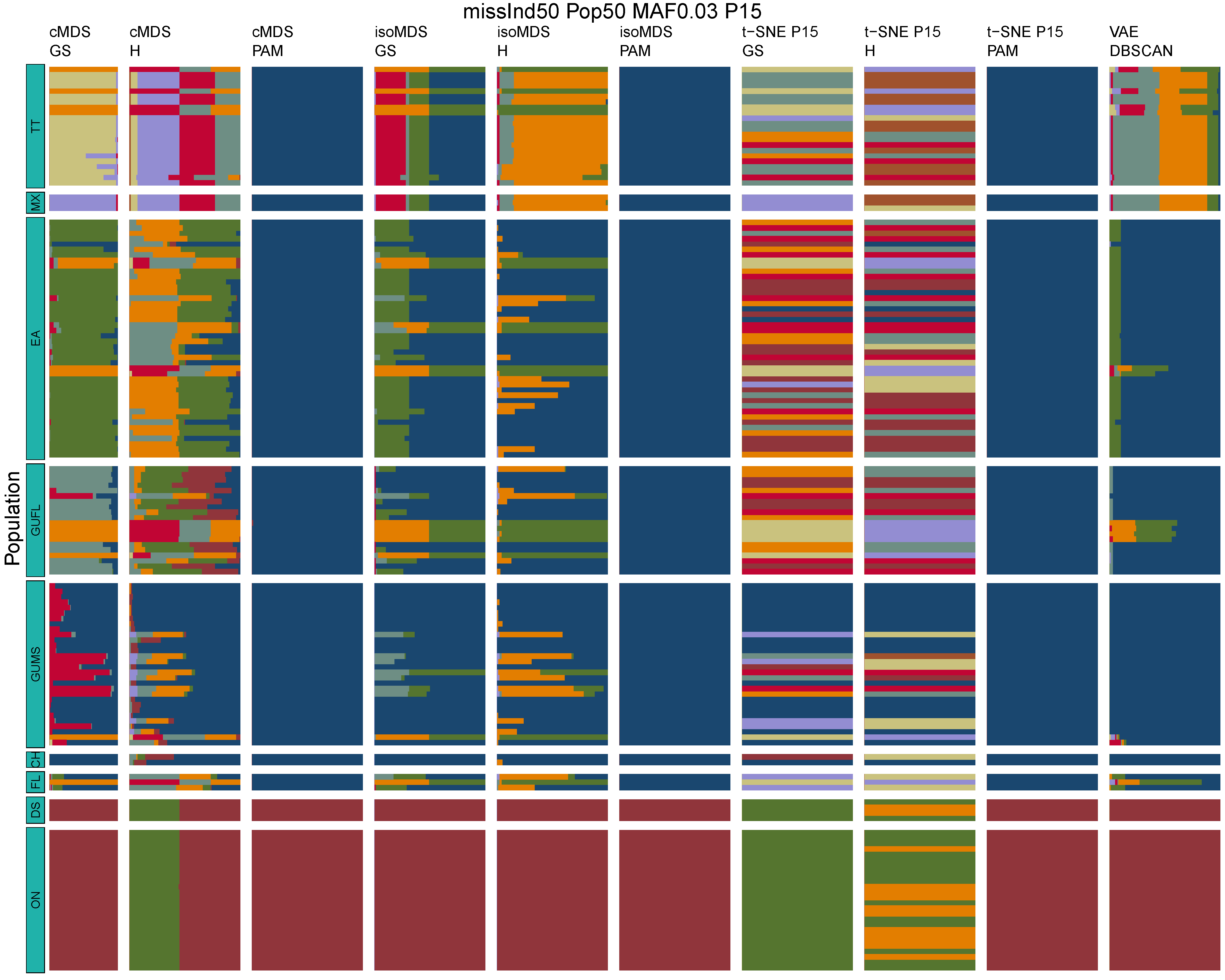

**Figure B23:** *Terrapene* unsupervised machine learning (UML) barplots depicting assignment proportions (x axis) among 100 replicates. Filtering parameters allowed maximum per-individual and per-population missing data and minimum minor allele frequency (MAF) filters. The included maximum missing data and minimum MAF filter proportions are shown in the plot title (missInd=per-individual, Pop=per-population filtering, MAF=MAF filter, P=perplexity). Each barplot on the page represents one UML algorithm: random forest, visualized with classical and isotonic multidimensional scaling (cMDS and isoMDS) ordination. t-SNE=t-distributed stochastic neighbor embedding with perplexity=15 (P15), and VAE=variational autoencoder. cMDS, isoMDS, and t-SNE were clustered with the partition around medoids (PAM) and hierarchical clustering (H) algorithms, and optimal *K* was chosen using PAM with the gap statistic (GS), H with the highest mean silhouette width (HMSW), and PAM with the HMSW. For VAE, clustering was performed, and optimal *K* chosen, with DBSCAN. *Terrapene* populations are separated by the horizontal white strips. Population IDs (blue left-aligned strip) include the ON=Ornate (*T. ornata ornata*), DS=Desert (*T. o. luteola*), FL=Florida (*T. c. bauri*), CH=Coahuilan (*T. coahuila*), GUMS=Gulf Coast (*T. carolina major*) from Mississippi, GUFL=Gulf Coast from the Florida Panhandle, EA=Woodland (*T. c. carolina*), MX=Mexican (*T. mexicana mexicana*), and TT=Three-toed (*T. m. triunguis*).
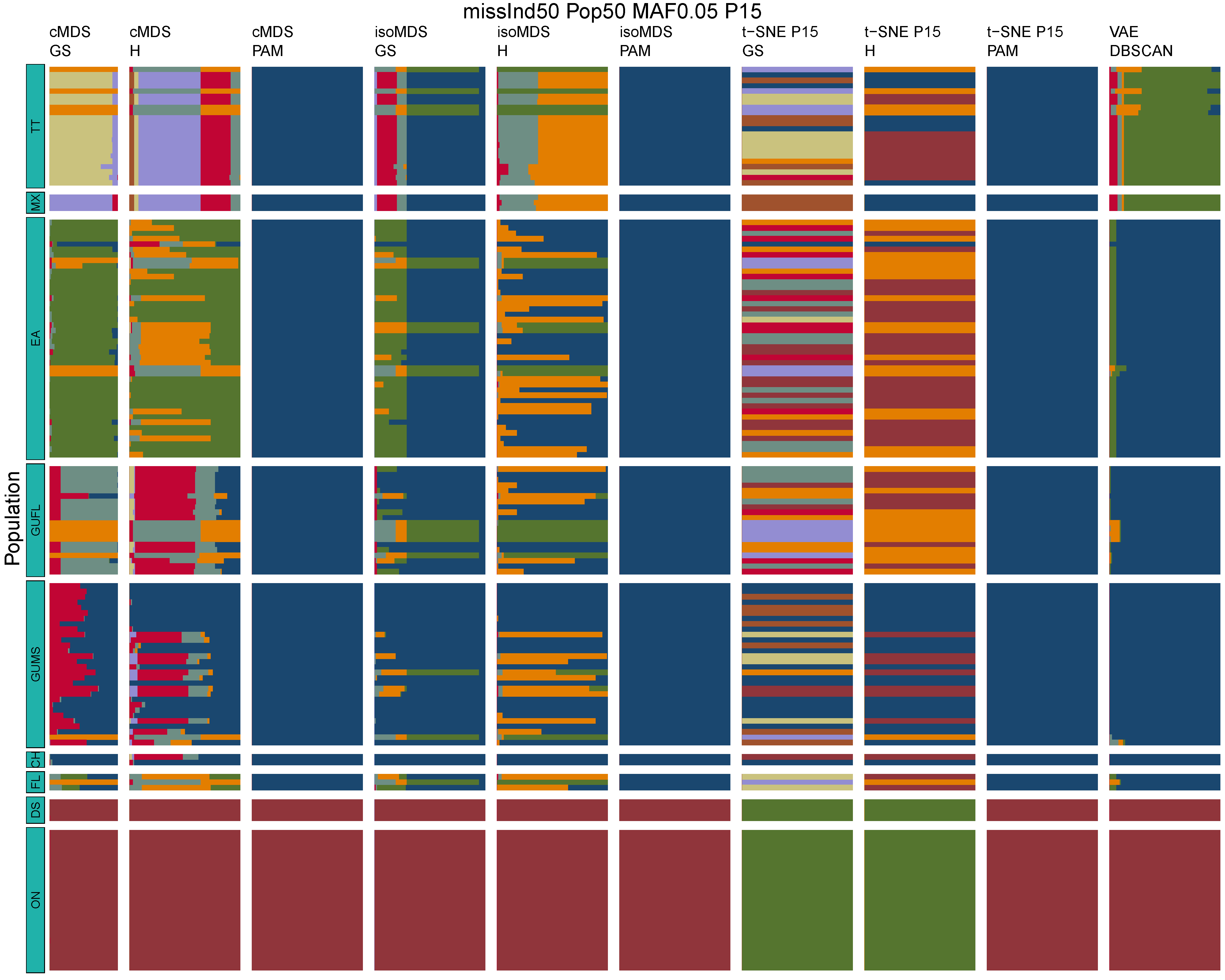

**Figure B24:** *Terrapene* unsupervised machine learning (UML) barplots depicting assignment proportions (x axis) among 100 replicates. Filtering parameters allowed maximum per-individual and per-population missing data and minimum minor allele frequency (MAF) filters. The included maximum missing data and minimum MAF filter proportions are shown in the plot title (missInd=per-individual, Pop=per-population filtering, MAF=MAF filter, P=perplexity). Each barplot on the page represents one UML algorithm: random forest, visualized with classical and isotonic multidimensional scaling (cMDS and isoMDS) ordination. t-SNE=t-distributed stochastic neighbor embedding with perplexity=15 (P15), and VAE=variational autoencoder. cMDS, isoMDS, and t-SNE were clustered with the partition around medoids (PAM) and hierarchical clustering (H) algorithms, and optimal *K* was chosen using PAM with the gap statistic (GS), H with the highest mean silhouette width (HMSW), and PAM with the HMSW. For VAE, clustering was performed, and optimal *K* chosen, with DBSCAN. *Terrapene* populations are separated by the horizontal white strips. Population IDs (blue left-aligned strip) include the ON=Ornate (*T. ornata ornata*), DS=Desert (*T. o. luteola*), FL=Florida (*T. c. bauri*), CH=Coahuilan (*T. coahuila*), GUMS=Gulf Coast (*T. carolina major*) from Mississippi, GUFL=Gulf Coast from the Florida Panhandle, EA=Woodland (*T. c. carolina*), MX=Mexican (*T. mexicana mexicana*), and TT=Three-toed (*T. m. triunguis*).
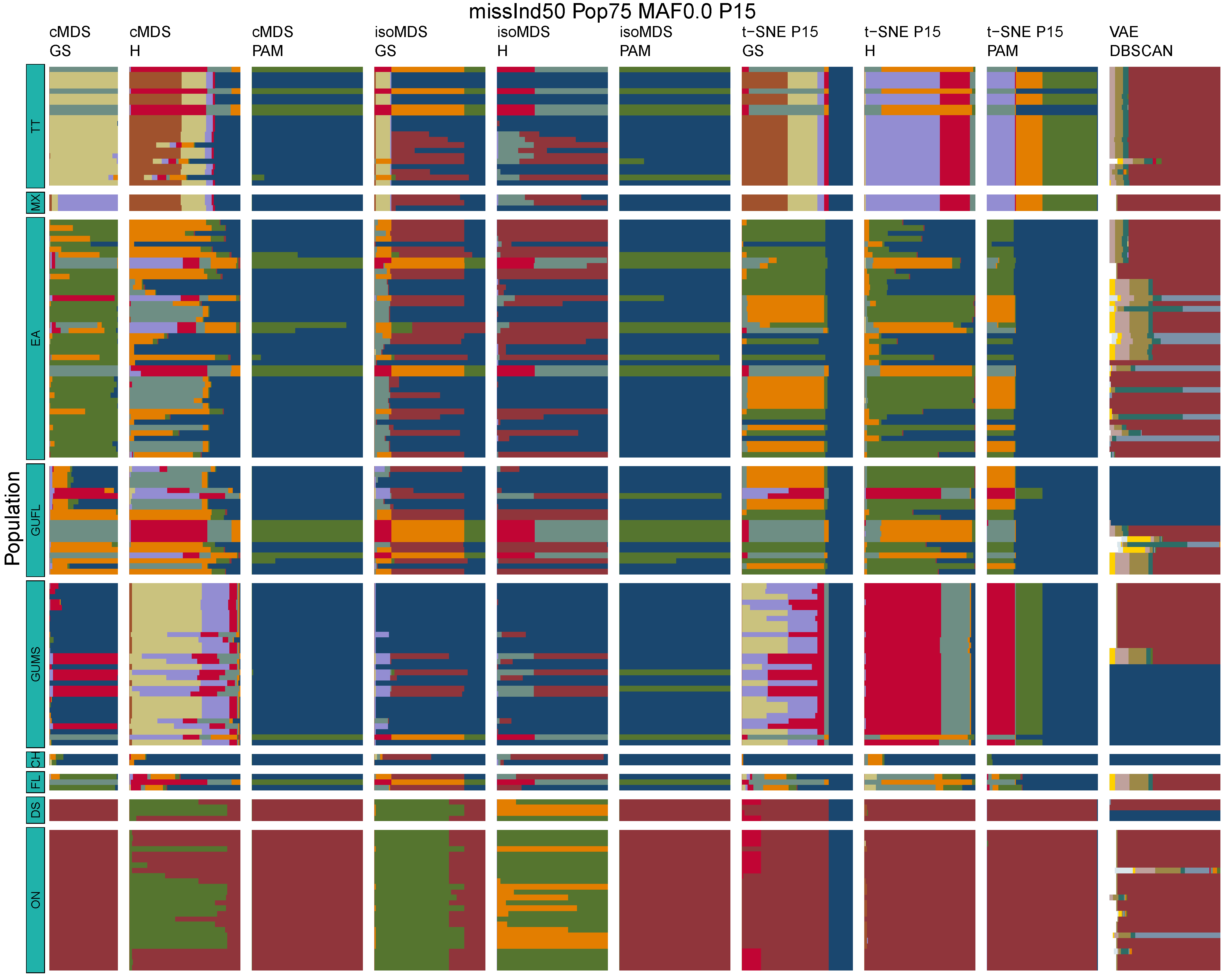

**Figure B25:** *Terrapene* unsupervised machine learning (UML) barplots depicting assignment proportions (x axis) among 100 replicates. Filtering parameters allowed maximum per-individual and per-population missing data and minimum minor allele frequency (MAF) filters. The included maximum missing data and minimum MAF filter proportions are shown in the plot title (missInd=per-individual, Pop=per-population filtering, MAF=MAF filter, P=perplexity). Each barplot on the page represents one UML algorithm: random forest, visualized with classical and isotonic multidimensional scaling (cMDS and isoMDS) ordination. t-SNE=t-distributed stochastic neighbor embedding with perplexity=15 (P15), and VAE=variational autoencoder. cMDS, isoMDS, and t-SNE were clustered with the partition around medoids (PAM) and hierarchical clustering (H) algorithms, and optimal *K* was chosen using PAM with the gap statistic (GS), H with the highest mean silhouette width (HMSW), and PAM with the HMSW. For VAE, clustering was performed, and optimal *K* chosen, with DBSCAN. *Terrapene* populations are separated by the horizontal white strips. Population IDs (blue left-aligned strip) include the ON=Ornate (*T. ornata ornata*), DS=Desert (*T. o. luteola*), FL=Florida (*T. c. bauri*), CH=Coahuilan (*T. coahuila*), GUMS=Gulf Coast (*T. carolina major*) from Mississippi, GUFL=Gulf Coast from the Florida Panhandle, EA=Woodland (*T. c. carolina*), MX=Mexican (*T. mexicana mexicana*), and TT=Three-toed (*T. m. triunguis*).
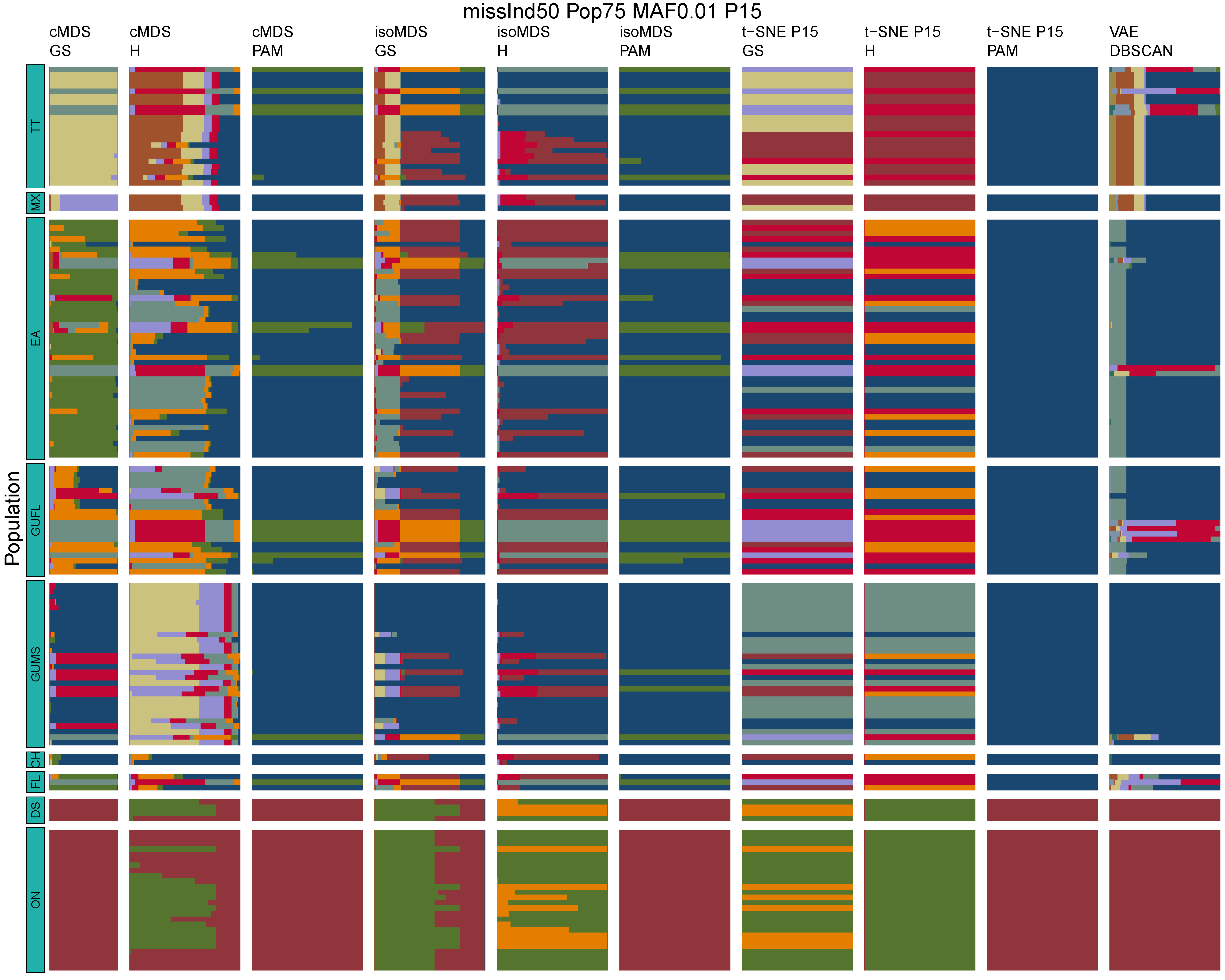

**Figure B26:** *Terrapene* unsupervised machine learning (UML) barplots depicting assignment proportions (x axis) among 100 replicates. Filtering parameters allowed maximum per-individual and per-population missing data and minimum minor allele frequency (MAF) filters. The included maximum missing data and minimum MAF filter proportions are shown in the plot title (missInd=per-individual, Pop=per-population filtering, MAF=MAF filter, P=perplexity). Each barplot on the page represents one UML algorithm: random forest, visualized with classical and isotonic multidimensional scaling (cMDS and isoMDS) ordination. t-SNE=t-distributed stochastic neighbor embedding with perplexity=15 (P15), and VAE=variational autoencoder. cMDS, isoMDS, and t-SNE were clustered with the partition around medoids (PAM) and hierarchical clustering (H) algorithms, and optimal *K* was chosen using PAM with the gap statistic (GS), H with the highest mean silhouette width (HMSW), and PAM with the HMSW. For VAE, clustering was performed, and optimal *K* chosen, with DBSCAN. *Terrapene* populations are separated by the horizontal white strips. Population IDs (blue left-aligned strip) include the ON=Ornate (*T. ornata ornata*), DS=Desert (*T. o. luteola*), FL=Florida (*T. c. bauri*), CH=Coahuilan (*T. coahuila*), GUMS=Gulf Coast (*T. carolina major*) from Mississippi, GUFL=Gulf Coast from the Florida Panhandle, EA=Woodland (*T. c. carolina*), MX=Mexican (*T. mexicana mexicana*), and TT=Three-toed (*T. m. triunguis*).
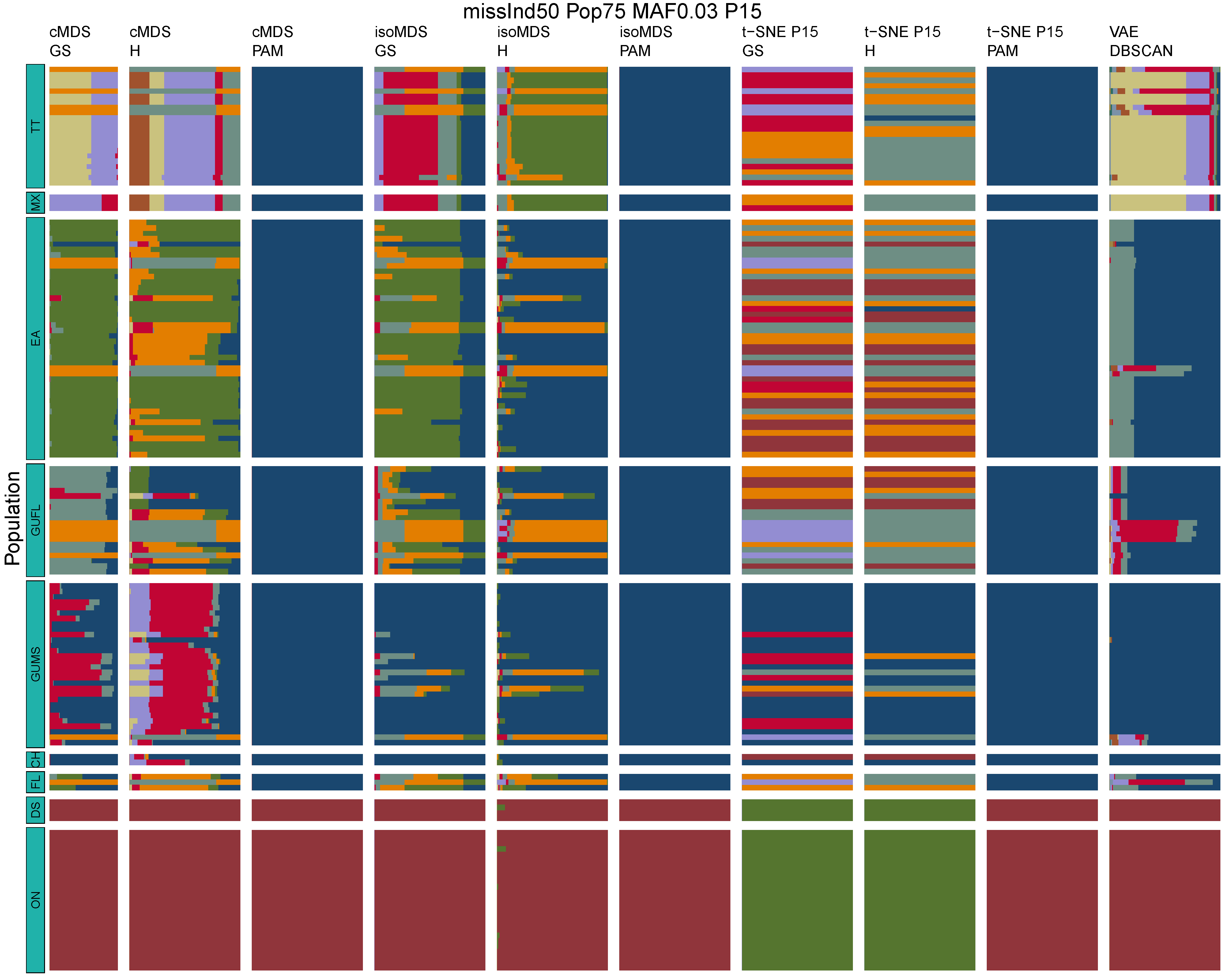

**Figure B27:** *Terrapene* unsupervised machine learning (UML) barplots depicting assignment proportions (x axis) among 100 replicates. Filtering parameters allowed maximum per-individual and per-population missing data and minimum minor allele frequency (MAF) filters. The included maximum missing data and minimum MAF filter proportions are shown in the plot title (missInd=per-individual, Pop=per-population filtering, MAF=MAF filter, P=perplexity). Each barplot on the page represents one UML algorithm: random forest, visualized with classical and isotonic multidimensional scaling (cMDS and isoMDS) ordination. t-SNE=t-distributed stochastic neighbor embedding with perplexity=15 (P15), and VAE=variational autoencoder. cMDS, isoMDS, and t-SNE were clustered with the partition around medoids (PAM) and hierarchical clustering (H) algorithms, and optimal *K* was chosen using PAM with the gap statistic (GS), H with the highest mean silhouette width (HMSW), and PAM with the HMSW. For VAE, clustering was performed, and optimal *K* chosen, with DBSCAN. *Terrapene* populations are separated by the horizontal white strips. Population IDs (blue left-aligned strip) include the ON=Ornate (*T. ornata ornata*), DS=Desert (*T. o. luteola*), FL=Florida (*T. c. bauri*), CH=Coahuilan (*T. coahuila*), GUMS=Gulf Coast (*T. carolina major*) from Mississippi, GUFL=Gulf Coast from the Florida Panhandle, EA=Woodland (*T. c. carolina*), MX=Mexican (*T. mexicana mexicana*), and TT=Three-toed (*T. m. triunguis*).
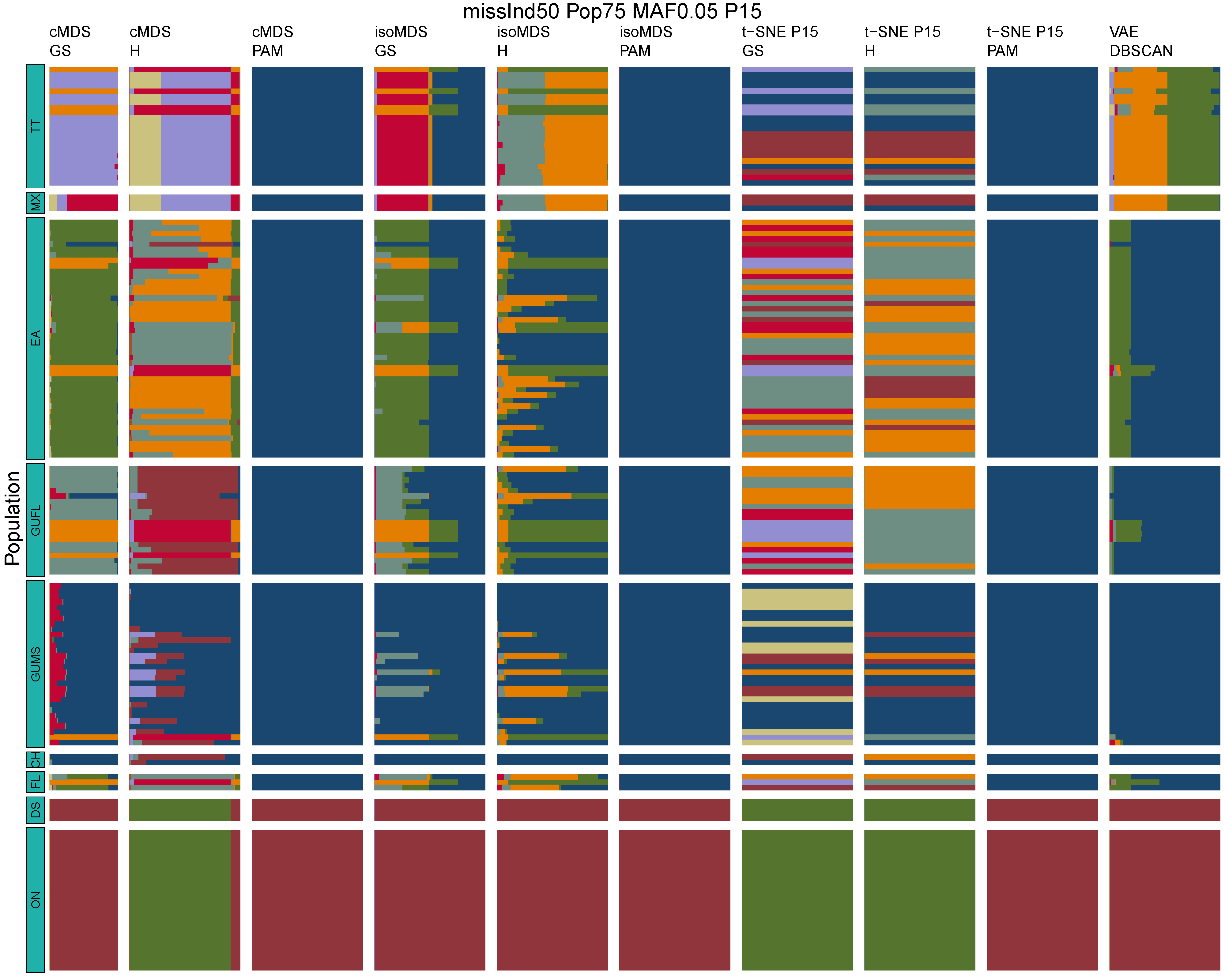

**Figure B28:** *Terrapene* unsupervised machine learning (UML) barplots depicting assignment proportions (x axis) among 100 replicates. Filtering parameters allowed maximum per-individual and per-population missing data and minimum minor allele frequency (MAF) filters. The included maximum missing data and minimum MAF filter proportions are shown in the plot title (missInd=per-individual, Pop=per-population filtering, MAF=MAF filter, P=perplexity). Each barplot on the page represents one UML algorithm: random forest, visualized with classical and isotonic multidimensional scaling (cMDS and isoMDS) ordination. t-SNE=t-distributed stochastic neighbor embedding with perplexity=15 (P15), and VAE=variational autoencoder. cMDS, isoMDS, and t-SNE were clustered with the partition around medoids (PAM) and hierarchical clustering (H) algorithms, and optimal *K* was chosen using PAM with the gap statistic (GS), H with the highest mean silhouette width (HMSW), and PAM with the HMSW. For VAE, clustering was performed, and optimal *K* chosen, with DBSCAN. *Terrapene* populations are separated by the horizontal white strips. Population IDs (blue left-aligned strip) include the ON=Ornate (*T. ornata ornata*), DS=Desert (*T. o. luteola*), FL=Florida (*T. c. bauri*), CH=Coahuilan (*T. coahuila*), GUMS=Gulf Coast (*T. carolina major*) from Mississippi, GUFL=Gulf Coast from the Florida Panhandle, EA=Woodland (*T. c. carolina*), MX=Mexican (*T. mexicana mexicana*), and TT=Three-toed (*T. m. triunguis*).
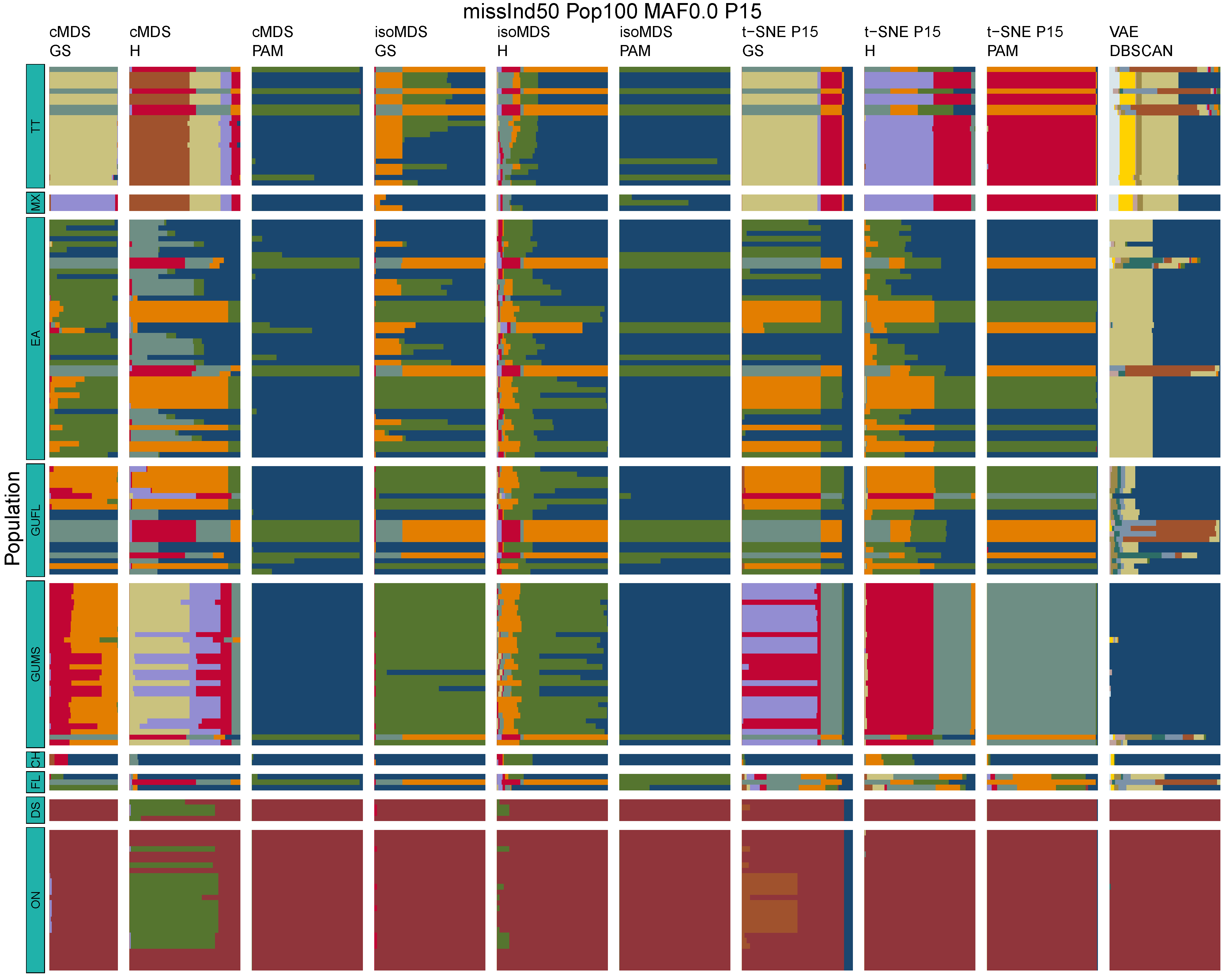

**Figure B29:** *Terrapene* unsupervised machine learning (UML) barplots depicting assignment proportions (x axis) among 100 replicates. Filtering parameters allowed maximum per-individual and per-population missing data and minimum minor allele frequency (MAF) filters. The included maximum missing data and minimum MAF filter proportions are shown in the plot title (missInd=per-individual, Pop=per-population filtering, MAF=MAF filter, P=perplexity). Each barplot on the page represents one UML algorithm: random forest, visualized with classical and isotonic multidimensional scaling (cMDS and isoMDS) ordination. t-SNE=t-distributed stochastic neighbor embedding with perplexity=15 (P15), and VAE=variational autoencoder. cMDS, isoMDS, and t-SNE were clustered with the partition around medoids (PAM) and hierarchical clustering (H) algorithms, and optimal *K* was chosen using PAM with the gap statistic (GS), H with the highest mean silhouette width (HMSW), and PAM with the HMSW. For VAE, clustering was performed, and optimal *K* chosen, with DBSCAN. *Terrapene* populations are separated by the horizontal white strips. Population IDs (blue left-aligned strip) include the ON=Ornate (*T. ornata ornata*), DS=Desert (*T. o. luteola*), FL=Florida (*T. c. bauri*), CH=Coahuilan (*T. coahuila*), GUMS=Gulf Coast (*T. carolina major*) from Mississippi, GUFL=Gulf Coast from the Florida Panhandle, EA=Woodland (*T. c. carolina*), MX=Mexican (*T. mexicana mexicana*), and TT=Three-toed (*T. m. triunguis*).
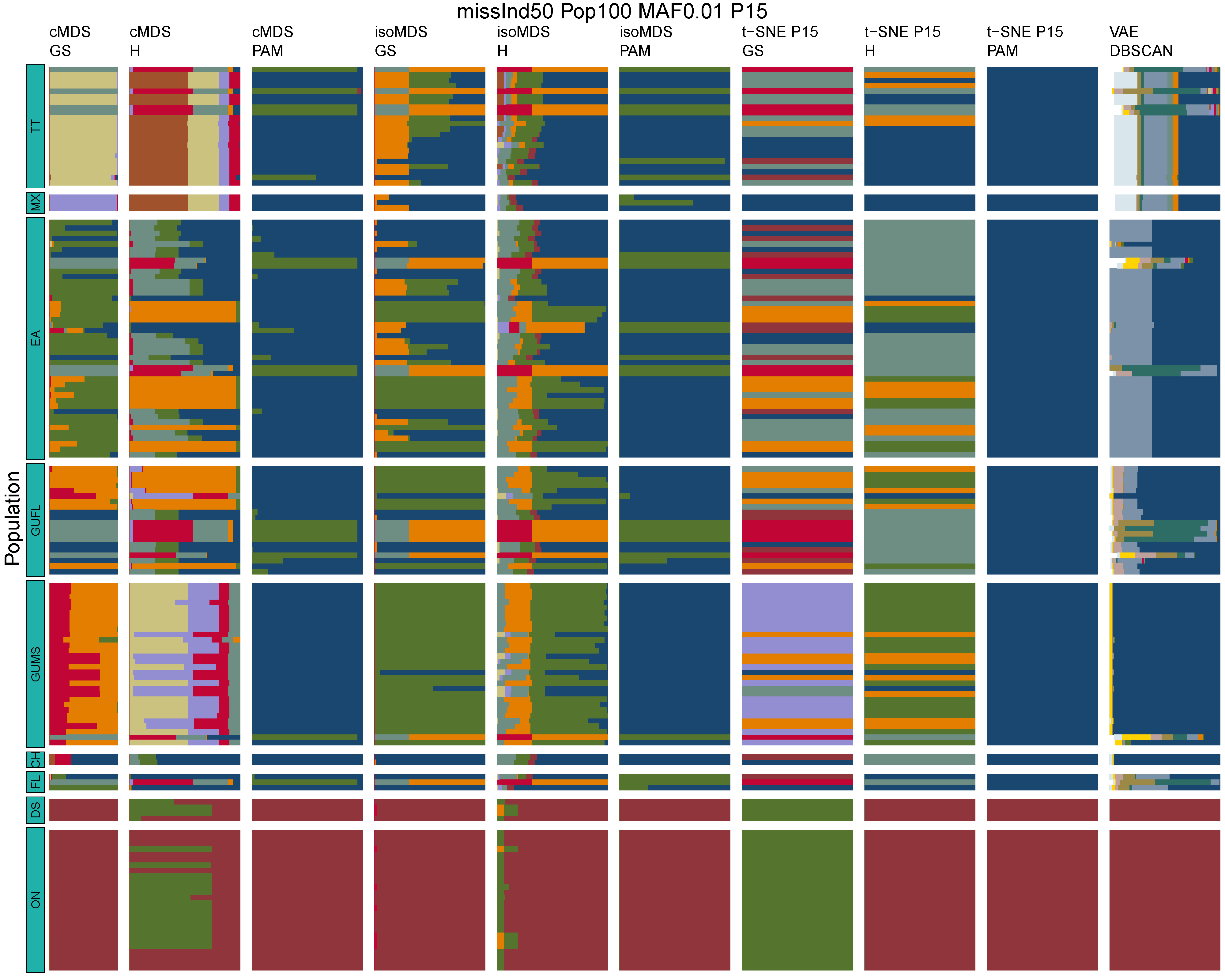

**Figure B30:** *Terrapene* unsupervised machine learning (UML) barplots depicting assignment proportions (x axis) among 100 replicates. Filtering parameters allowed maximum per-individual and per-population missing data and minimum minor allele frequency (MAF) filters. The included maximum missing data and minimum MAF filter proportions are shown in the plot title (missInd=per-individual, Pop=per-population filtering, MAF=MAF filter, P=perplexity). Each barplot on the page represents one UML algorithm: random forest, visualized with classical and isotonic multidimensional scaling (cMDS and isoMDS) ordination. t-SNE=t-distributed stochastic neighbor embedding with perplexity=15 (P15), and VAE=variational autoencoder. cMDS, isoMDS, and t-SNE were clustered with the partition around medoids (PAM) and hierarchical clustering (H) algorithms, and optimal *K* was chosen using PAM with the gap statistic (GS), H with the highest mean silhouette width (HMSW), and PAM with the HMSW. For VAE, clustering was performed, and optimal *K* chosen, with DBSCAN. *Terrapene* populations are separated by the horizontal white strips. Population IDs (blue left-aligned strip) include the ON=Ornate (*T. ornata ornata*), DS=Desert (*T. o. luteola*), FL=Florida (*T. c. bauri*), CH=Coahuilan (*T. coahuila*), GUMS=Gulf Coast (*T. carolina major*) from Mississippi, GUFL=Gulf Coast from the Florida Panhandle, EA=Woodland (*T. c. carolina*), MX=Mexican (*T. mexicana mexicana*), and TT=Three-toed (*T. m. triunguis*).

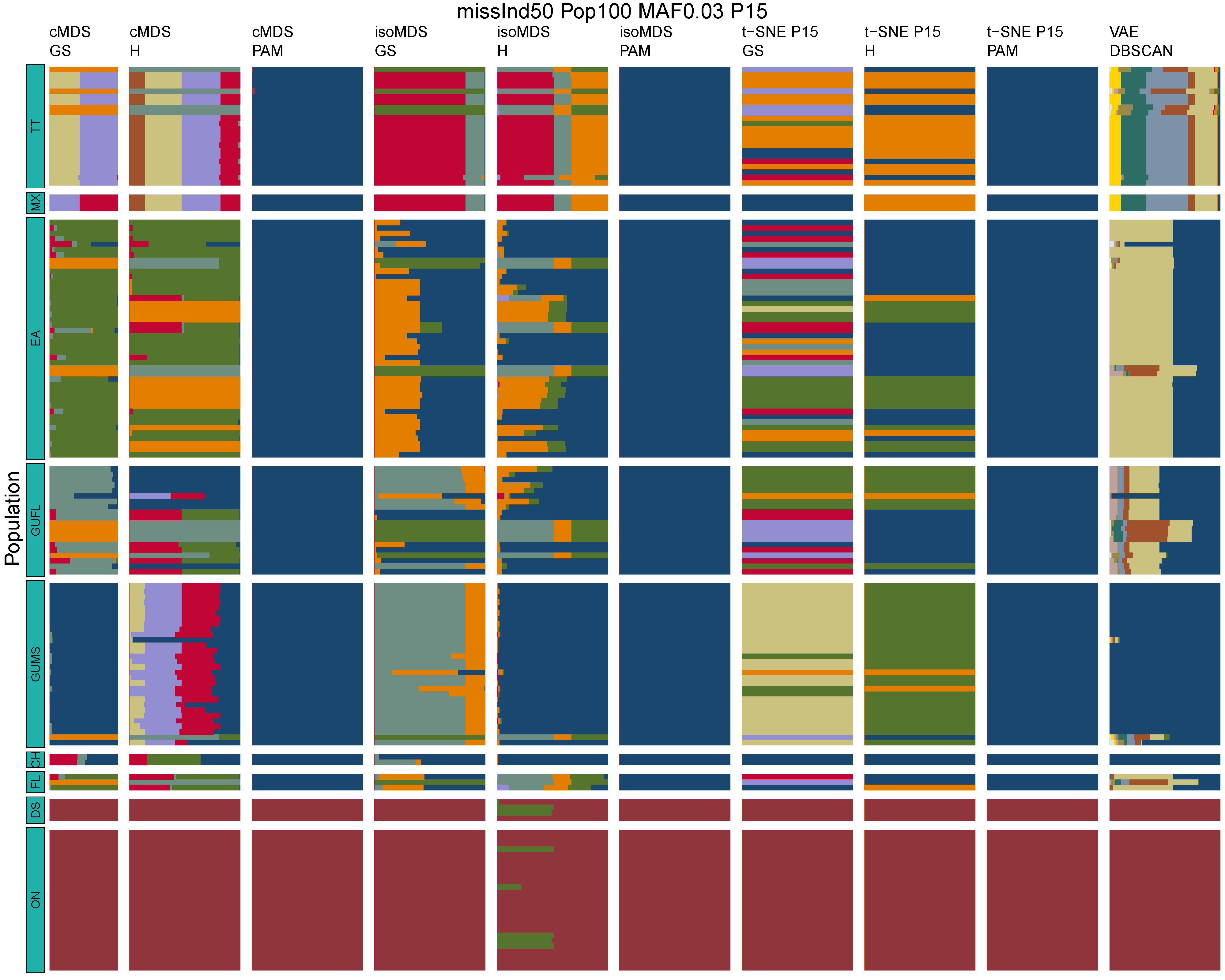

**Figure B31:** *Terrapene* unsupervised machine learning (UML) barplots depicting assignment proportions (x axis) among 100 replicates. Filtering parameters allowed maximum per-individual and per-population missing data and minimum minor allele frequency (MAF) filters. The included maximum missing data and minimum MAF filter proportions are shown in the plot title (missInd=per-individual, Pop=per-population filtering, MAF=MAF filter, P=perplexity). Each barplot on the page represents one UML algorithm: random forest, visualized with classical and isotonic multidimensional scaling (cMDS and isoMDS) ordination. t-SNE=t-distributed stochastic neighbor embedding with perplexity=15 (P15), and VAE=variational autoencoder. cMDS, isoMDS, and t-SNE were clustered with the partition around medoids (PAM) and hierarchical clustering (H) algorithms, and optimal *K* was chosen using PAM with the gap statistic (GS), H with the highest mean silhouette width (HMSW), and PAM with the HMSW. For VAE, clustering was performed, and optimal *K* chosen, with DBSCAN. *Terrapene* populations are separated by the horizontal white strips. Population IDs (blue left-aligned strip) include the ON=Ornate (*T. ornata ornata*), DS=Desert (*T. o. luteola*), FL=Florida (*T. c. bauri*), CH=Coahuilan (*T. coahuila*), GUMS=Gulf Coast (*T. carolina major*) from Mississippi, GUFL=Gulf Coast from the Florida Panhandle, EA=Woodland (*T. c. carolina*), MX=Mexican (*T. mexicana mexicana*), and TT=Three-toed (*T. m. triunguis*).
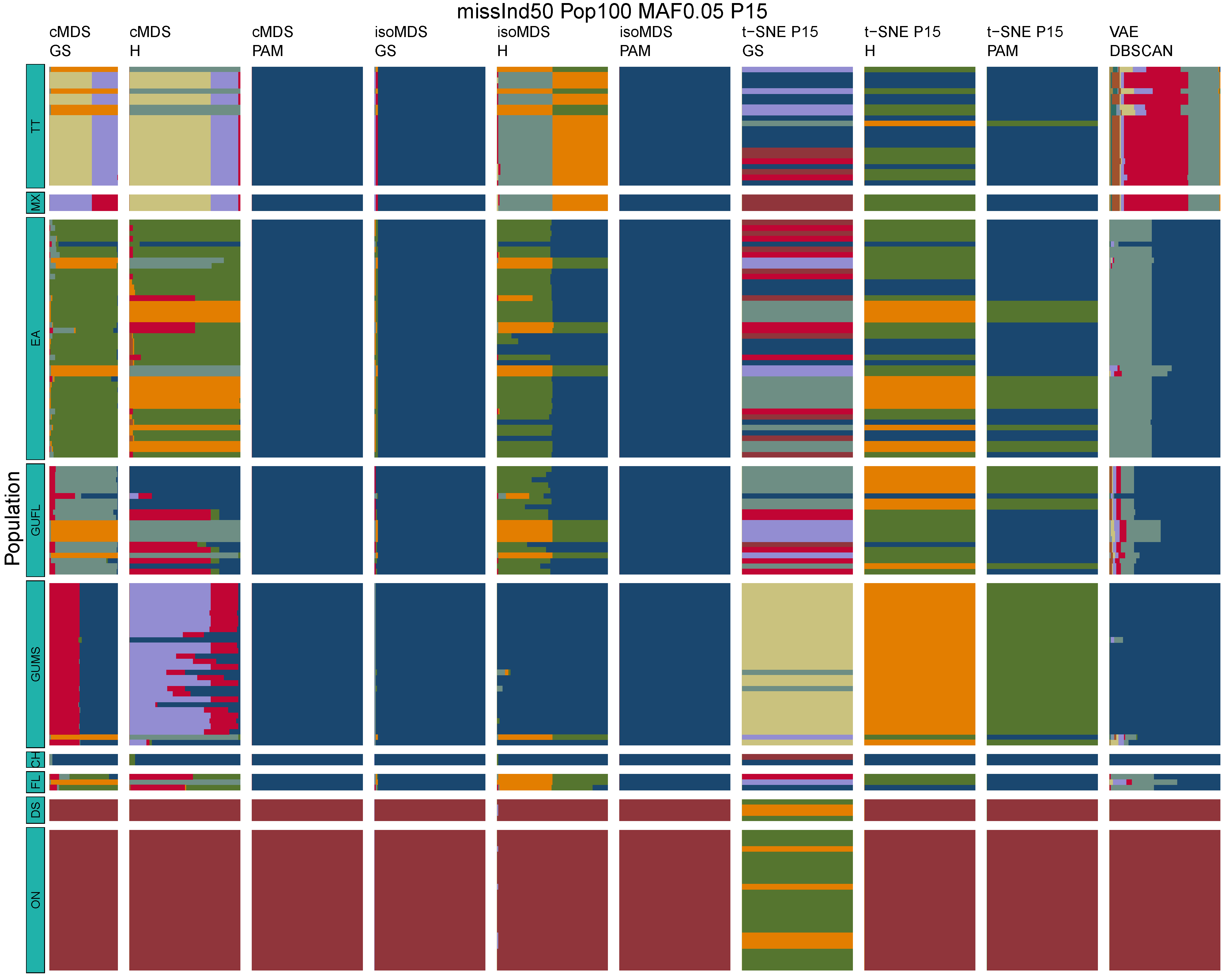

**Figure B32:** *Terrapene* unsupervised machine learning (UML) barplots depicting assignment proportions (x axis) among 100 replicates. Filtering parameters allowed maximum per-individual and per-population missing data and minimum minor allele frequency (MAF) filters. The included maximum missing data and minimum MAF filter proportions are shown in the plot title (missInd=per-individual, Pop=per-population filtering, MAF=MAF filter, P=perplexity). Each barplot on the page represents one UML algorithm: random forest, visualized with classical and isotonic multidimensional scaling (cMDS and isoMDS) ordination. t-SNE=t-distributed stochastic neighbor embedding with perplexity=15 (P15), and VAE=variational autoencoder. cMDS, isoMDS, and t-SNE were clustered with the partition around medoids (PAM) and hierarchical clustering (H) algorithms, and optimal *K* was chosen using PAM with the gap statistic (GS), H with the highest mean silhouette width (HMSW), and PAM with the HMSW. For VAE, clustering was performed, and optimal *K* chosen, with DBSCAN. *Terrapene* populations are separated by the horizontal white strips. Population IDs (blue left-aligned strip) include the ON=Ornate (*T. ornata ornata*), DS=Desert (*T. o. luteola*), FL=Florida (*T. c. bauri*), CH=Coahuilan (*T. coahuila*), GUMS=Gulf Coast (*T. carolina major*) from Mississippi, GUFL=Gulf Coast from the Florida Panhandle, EA=Woodland (*T. c. carolina*), MX=Mexican (*T. mexicana mexicana*), and TT=Three-toed (*T. m. triunguis*).
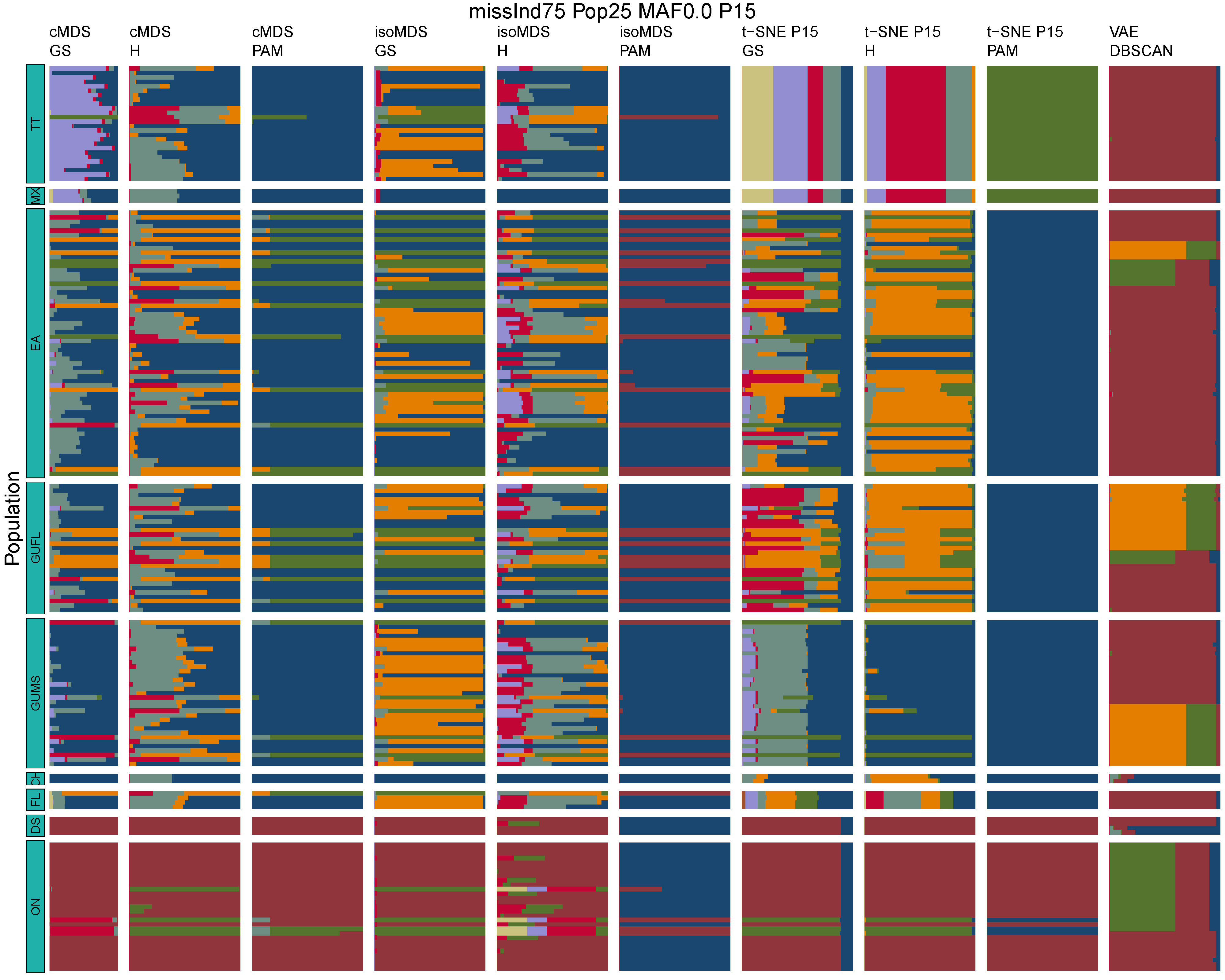

**Figure B33:** *Terrapene* unsupervised machine learning (UML) barplots depicting assignment proportions (x axis) among 100 replicates. Filtering parameters allowed maximum per-individual and per-population missing data and minimum minor allele frequency (MAF) filters. The included maximum missing data and minimum MAF filter proportions are shown in the plot title (missInd=per-individual, Pop=per-population filtering, MAF=MAF filter, P=perplexity). Each barplot on the page represents one UML algorithm: random forest, visualized with classical and isotonic multidimensional scaling (cMDS and isoMDS) ordination. t-SNE=t-distributed stochastic neighbor embedding with perplexity=15 (P15), and VAE=variational autoencoder. cMDS, isoMDS, and t-SNE were clustered with the partition around medoids (PAM) and hierarchical clustering (H) algorithms, and optimal *K* was chosen using PAM with the gap statistic (GS), H with the highest mean silhouette width (HMSW), and PAM with the HMSW. For VAE, clustering was performed, and optimal *K* chosen, with DBSCAN. *Terrapene* populations are separated by the horizontal white strips. Population IDs (blue left-aligned strip) include the ON=Ornate (*T. ornata ornata*), DS=Desert (*T. o. luteola*), FL=Florida (*T. c. bauri*), CH=Coahuilan (*T. coahuila*), GUMS=Gulf Coast (*T. carolina major*) from Mississippi, GUFL=Gulf Coast from the Florida Panhandle, EA=Woodland (*T. c. carolina*), MX=Mexican (*T. mexicana mexicana*), and TT=Three-toed (*T. m. triunguis*).
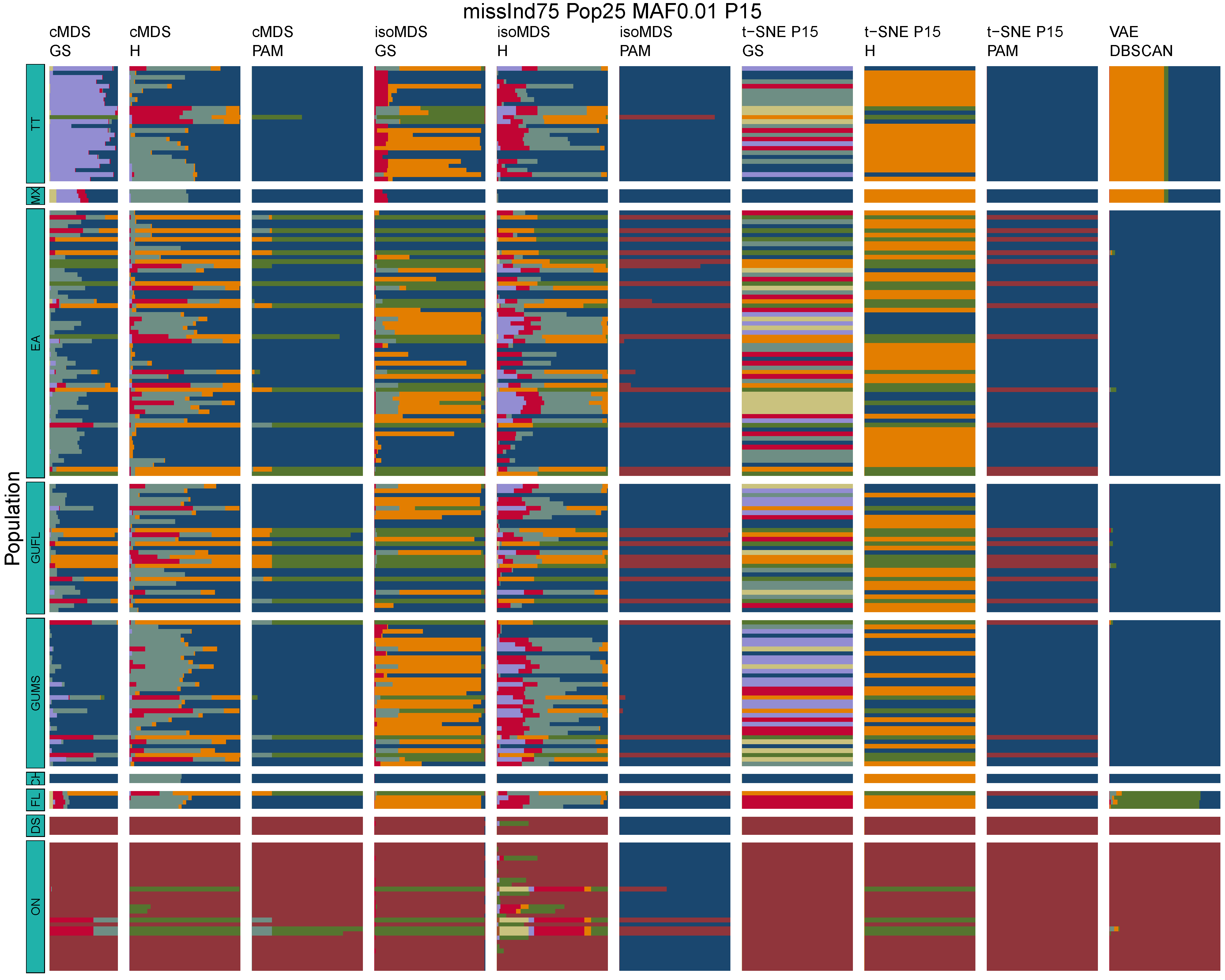

**Figure B34:** *Terrapene* unsupervised machine learning (UML) barplots depicting assignment proportions (x axis) among 100 replicates. Filtering parameters allowed maximum per-individual and per-population missing data and minimum minor allele frequency (MAF) filters. The included maximum missing data and minimum MAF filter proportions are shown in the plot title (missInd=per-individual, Pop=per-population filtering, MAF=MAF filter, P=perplexity). Each barplot on the page represents one UML algorithm: random forest, visualized with classical and isotonic multidimensional scaling (cMDS and isoMDS) ordination. t-SNE=t-distributed stochastic neighbor embedding with perplexity=15 (P15), and VAE=variational autoencoder. cMDS, isoMDS, and t-SNE were clustered with the partition around medoids (PAM) and hierarchical clustering (H) algorithms, and optimal *K* was chosen using PAM with the gap statistic (GS), H with the highest mean silhouette width (HMSW), and PAM with the HMSW. For VAE, clustering was performed, and optimal *K* chosen, with DBSCAN. *Terrapene* populations are separated by the horizontal white strips. Population IDs (blue left-aligned strip) include the ON=Ornate (*T. ornata ornata*), DS=Desert (*T. o. luteola*), FL=Florida (*T. c. bauri*), CH=Coahuilan (*T. coahuila*), GUMS=Gulf Coast (*T. carolina major*) from Mississippi, GUFL=Gulf Coast from the Florida Panhandle, EA=Woodland (*T. c. carolina*), MX=Mexican (*T. mexicana mexicana*), and TT=Three-toed (*T. m. triunguis*).
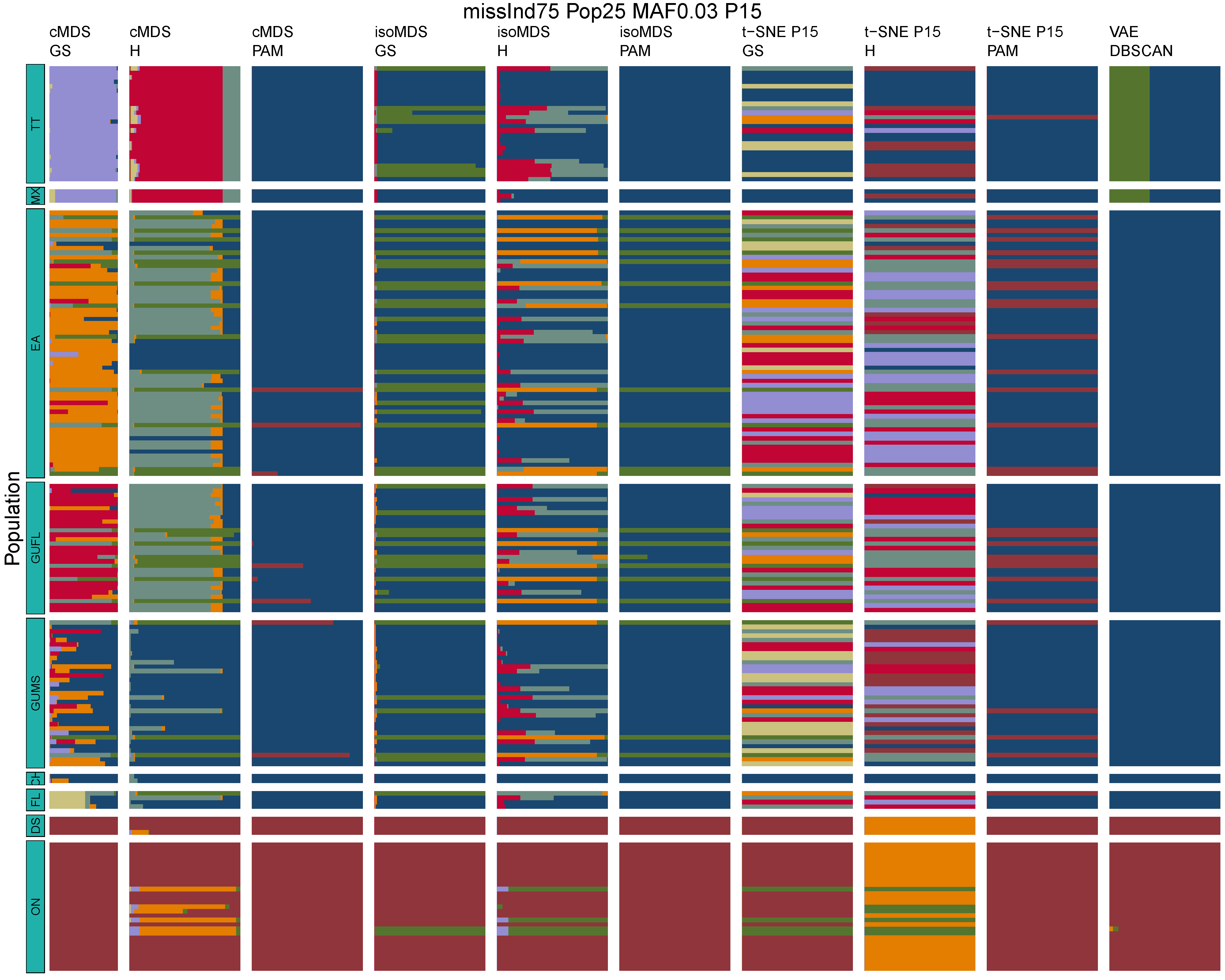

**Figure B35:** *Terrapene* unsupervised machine learning (UML) barplots depicting assignment proportions (x axis) among 100 replicates. Filtering parameters allowed maximum per-individual and per-population missing data and minimum minor allele frequency (MAF) filters. The included maximum missing data and minimum MAF filter proportions are shown in the plot title (missInd=per-individual, Pop=per-population filtering, MAF=MAF filter, P=perplexity). Each barplot on the page represents one UML algorithm: random forest, visualized with classical and isotonic multidimensional scaling (cMDS and isoMDS) ordination. t-SNE=t-distributed stochastic neighbor embedding with perplexity=15 (P15), and VAE=variational autoencoder. cMDS, isoMDS, and t-SNE were clustered with the partition around medoids (PAM) and hierarchical clustering (H) algorithms, and optimal *K* was chosen using PAM with the gap statistic (GS), H with the highest mean silhouette width (HMSW), and PAM with the HMSW. For VAE, clustering was performed, and optimal *K* chosen, with DBSCAN. *Terrapene* populations are separated by the horizontal white strips. Population IDs (blue left-aligned strip) include the ON=Ornate (*T. ornata ornata*), DS=Desert (*T. o. luteola*), FL=Florida (*T. c. bauri*), CH=Coahuilan (*T. coahuila*), GUMS=Gulf Coast (*T. carolina major*) from Mississippi, GUFL=Gulf Coast from the Florida Panhandle, EA=Woodland (*T. c. carolina*), MX=Mexican (*T. mexicana mexicana*), and TT=Three-toed (*T. m. triunguis*).
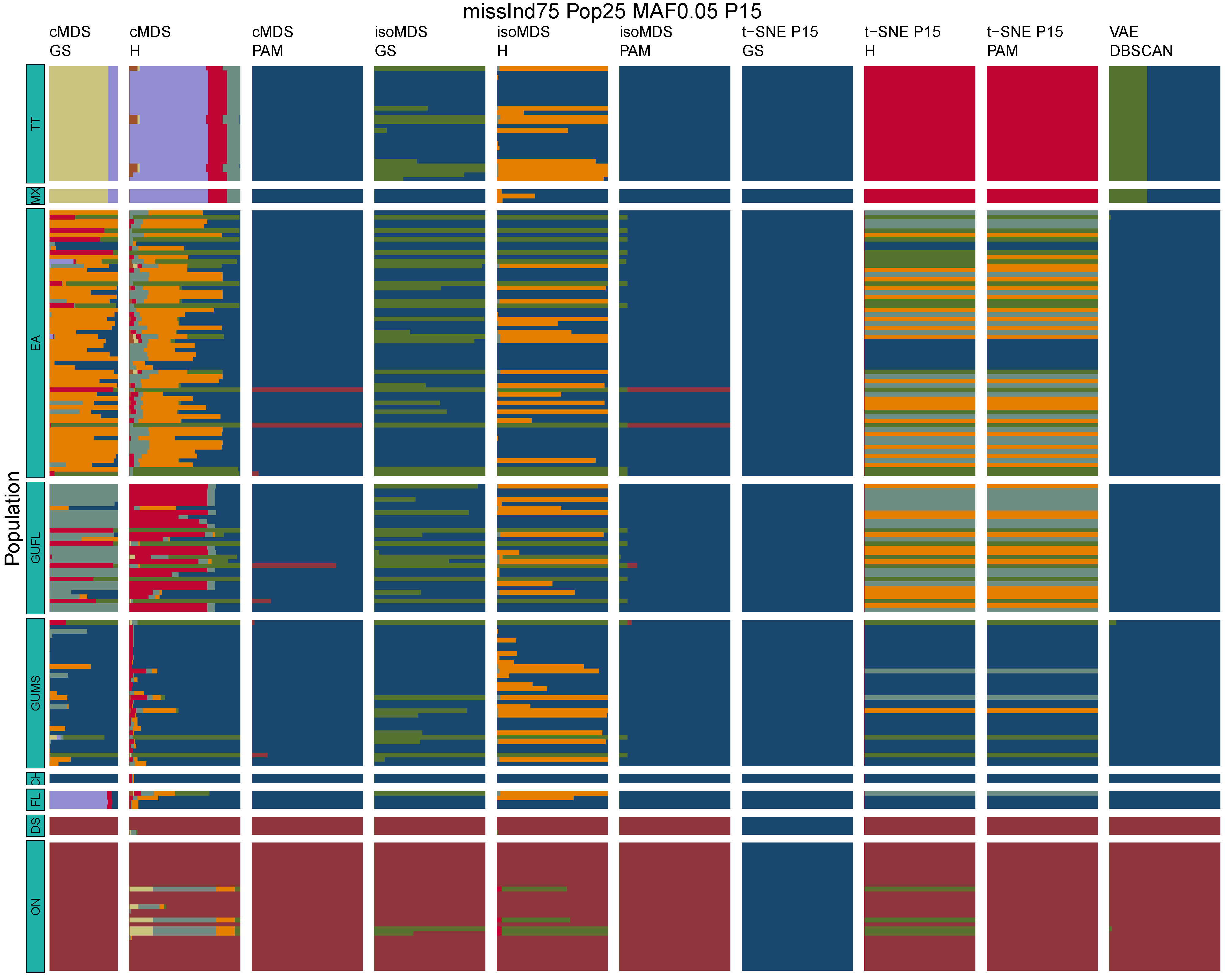

**Figure B36:** *Terrapene* unsupervised machine learning (UML) barplots depicting assignment proportions (x axis) among 100 replicates. Filtering parameters allowed maximum per-individual and per-population missing data and minimum minor allele frequency (MAF) filters. The included maximum missing data and minimum MAF filter proportions are shown in the plot title (missInd=per-individual, Pop=per-population filtering, MAF=MAF filter, P=perplexity). Each barplot on the page represents one UML algorithm: random forest, visualized with classical and isotonic multidimensional scaling (cMDS and isoMDS) ordination. t-SNE=t-distributed stochastic neighbor embedding with perplexity=15 (P15), and VAE=variational autoencoder. cMDS, isoMDS, and t-SNE were clustered with the partition around medoids (PAM) and hierarchical clustering (H) algorithms, and optimal *K* was chosen using PAM with the gap statistic (GS), H with the highest mean silhouette width (HMSW), and PAM with the HMSW. For VAE, clustering was performed, and optimal *K* chosen, with DBSCAN. *Terrapene* populations are separated by the horizontal white strips. Population IDs (blue left-aligned strip) include the ON=Ornate (*T. ornata ornata*), DS=Desert (*T. o. luteola*), FL=Florida (*T. c. bauri*), CH=Coahuilan (*T. coahuila*), GUMS=Gulf Coast (*T. carolina major*) from Mississippi, GUFL=Gulf Coast from the Florida Panhandle, EA=Woodland (*T. c. carolina*), MX=Mexican (*T. mexicana mexicana*), and TT=Three-toed (*T. m. triunguis*).
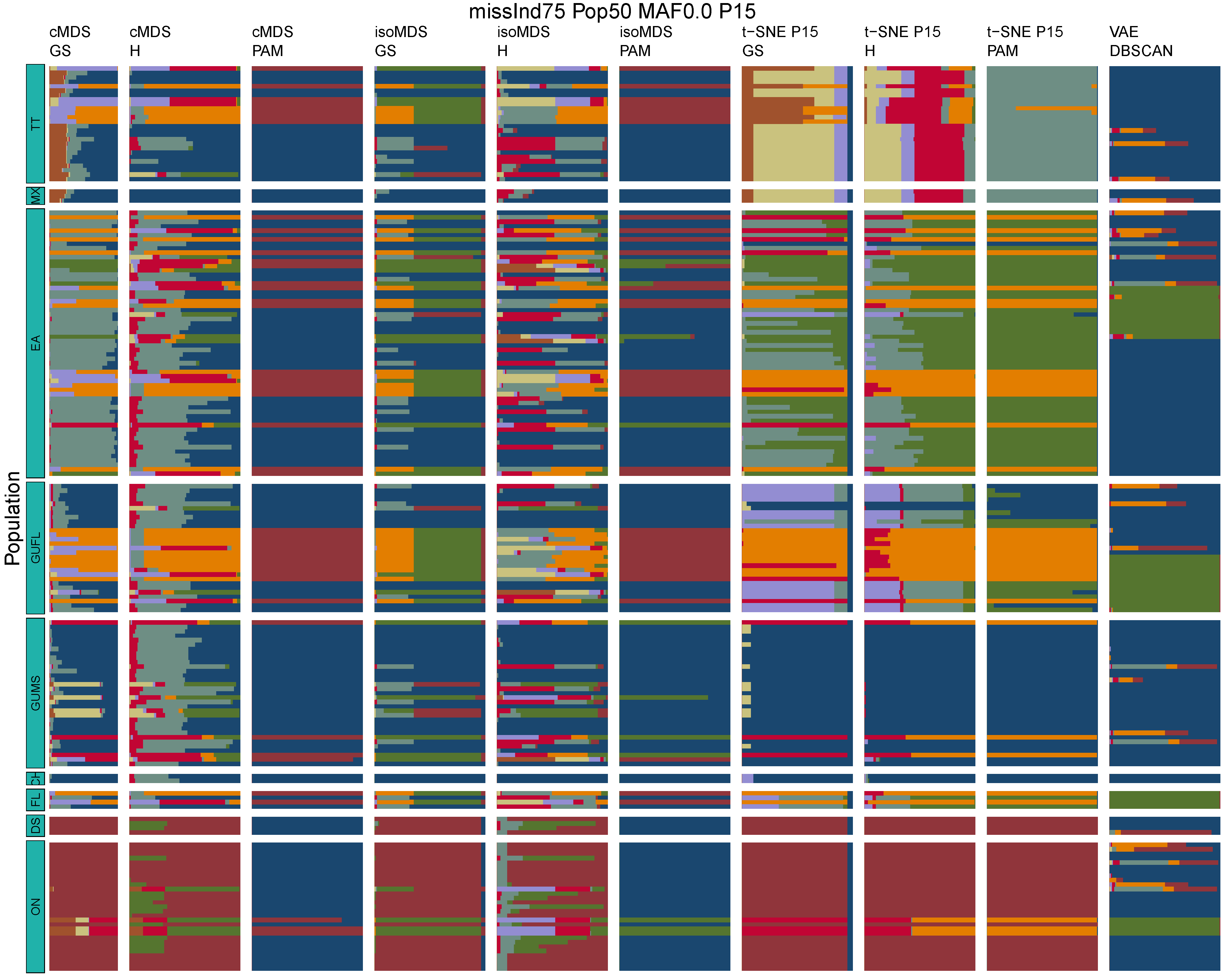

**Figure B37:** *Terrapene* unsupervised machine learning (UML) barplots depicting assignment proportions (x axis) among 100 replicates. Filtering parameters allowed maximum per-individual and per-population missing data and minimum minor allele frequency (MAF) filters. The included maximum missing data and minimum MAF filter proportions are shown in the plot title (missInd=per-individual, Pop=per-population filtering, MAF=MAF filter, P=perplexity). Each barplot on the page represents one UML algorithm: random forest, visualized with classical and isotonic multidimensional scaling (cMDS and isoMDS) ordination. t-SNE=t-distributed stochastic neighbor embedding with perplexity=15 (P15), and VAE=variational autoencoder. cMDS, isoMDS, and t-SNE were clustered with the partition around medoids (PAM) and hierarchical clustering (H) algorithms, and optimal *K* was chosen using PAM with the gap statistic (GS), H with the highest mean silhouette width (HMSW), and PAM with the HMSW. For VAE, clustering was performed, and optimal *K* chosen, with DBSCAN. *Terrapene* populations are separated by the horizontal white strips. Population IDs (blue left-aligned strip) include the ON=Ornate (*T. ornata ornata*), DS=Desert (*T. o. luteola*), FL=Florida (*T. c. bauri*), CH=Coahuilan (*T. coahuila*), GUMS=Gulf Coast (*T. carolina major*) from Mississippi, GUFL=Gulf Coast from the Florida Panhandle, EA=Woodland (*T. c. carolina*), MX=Mexican (*T. mexicana mexicana*), and TT=Three-toed (*T. m. triunguis*).
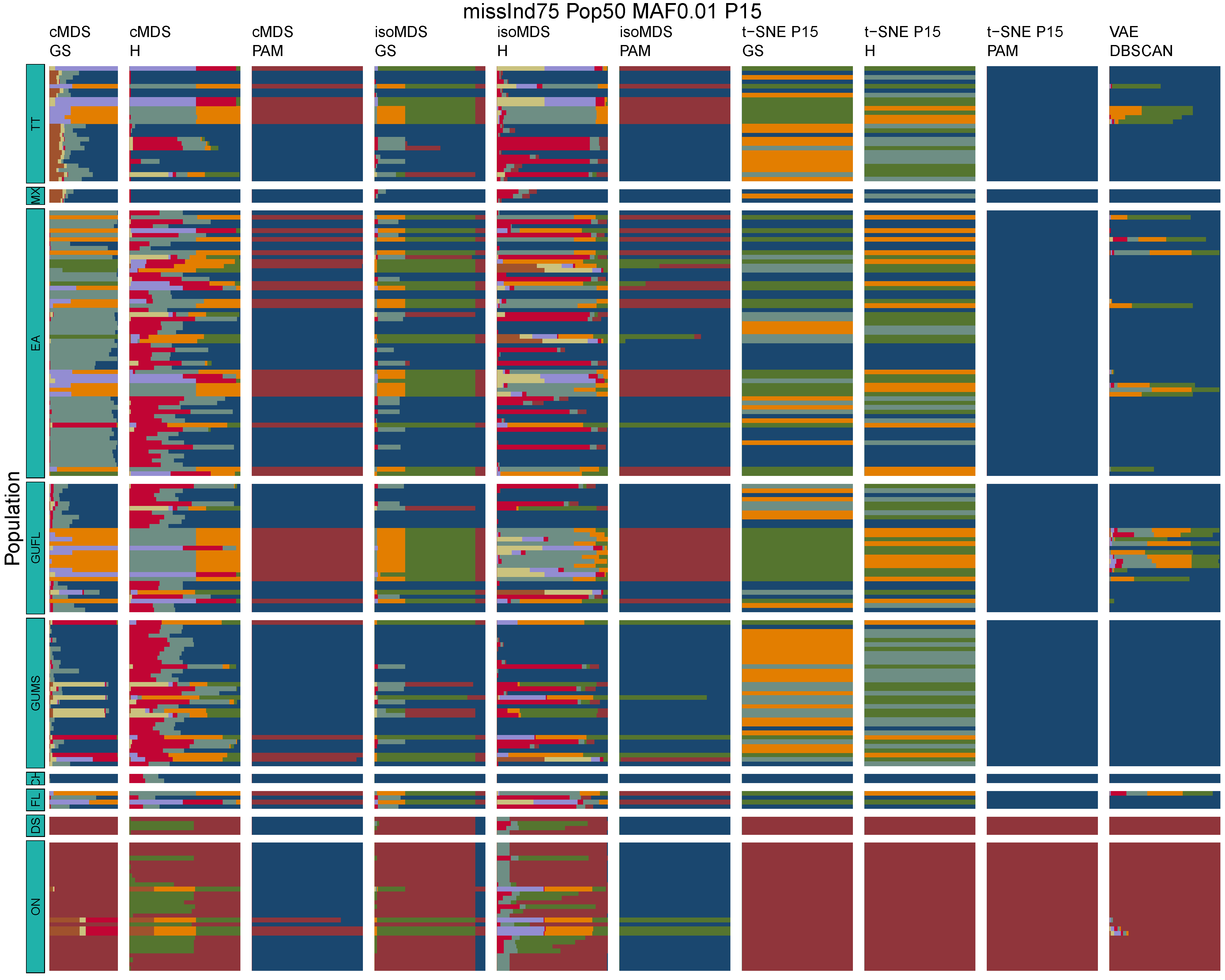

**Figure B38:** *Terrapene* unsupervised machine learning (UML) barplots depicting assignment proportions (x axis) among 100 replicates. Filtering parameters allowed maximum per-individual and per-population missing data and minimum minor allele frequency (MAF) filters. The included maximum missing data and minimum MAF filter proportions are shown in the plot title (missInd=per-individual, Pop=per-population filtering, MAF=MAF filter, P=perplexity). Each barplot on the page represents one UML algorithm: random forest, visualized with classical and isotonic multidimensional scaling (cMDS and isoMDS) ordination. t-SNE=t-distributed stochastic neighbor embedding with perplexity=15 (P15), and VAE=variational autoencoder. cMDS, isoMDS, and t-SNE were clustered with the partition around medoids (PAM) and hierarchical clustering (H) algorithms, and optimal *K* was chosen using PAM with the gap statistic (GS), H with the highest mean silhouette width (HMSW), and PAM with the HMSW. For VAE, clustering was performed, and optimal *K* chosen, with DBSCAN. *Terrapene* populations are separated by the horizontal white strips. Population IDs (blue left-aligned strip) include the ON=Ornate (*T. ornata ornata*), DS=Desert (*T. o. luteola*), FL=Florida (*T. c. bauri*), CH=Coahuilan (*T. coahuila*), GUMS=Gulf Coast (*T. carolina major*) from Mississippi, GUFL=Gulf Coast from the Florida Panhandle, EA=Woodland (*T. c. carolina*), MX=Mexican (*T. mexicana mexicana*), and TT=Three-toed (*T. m. triunguis*).
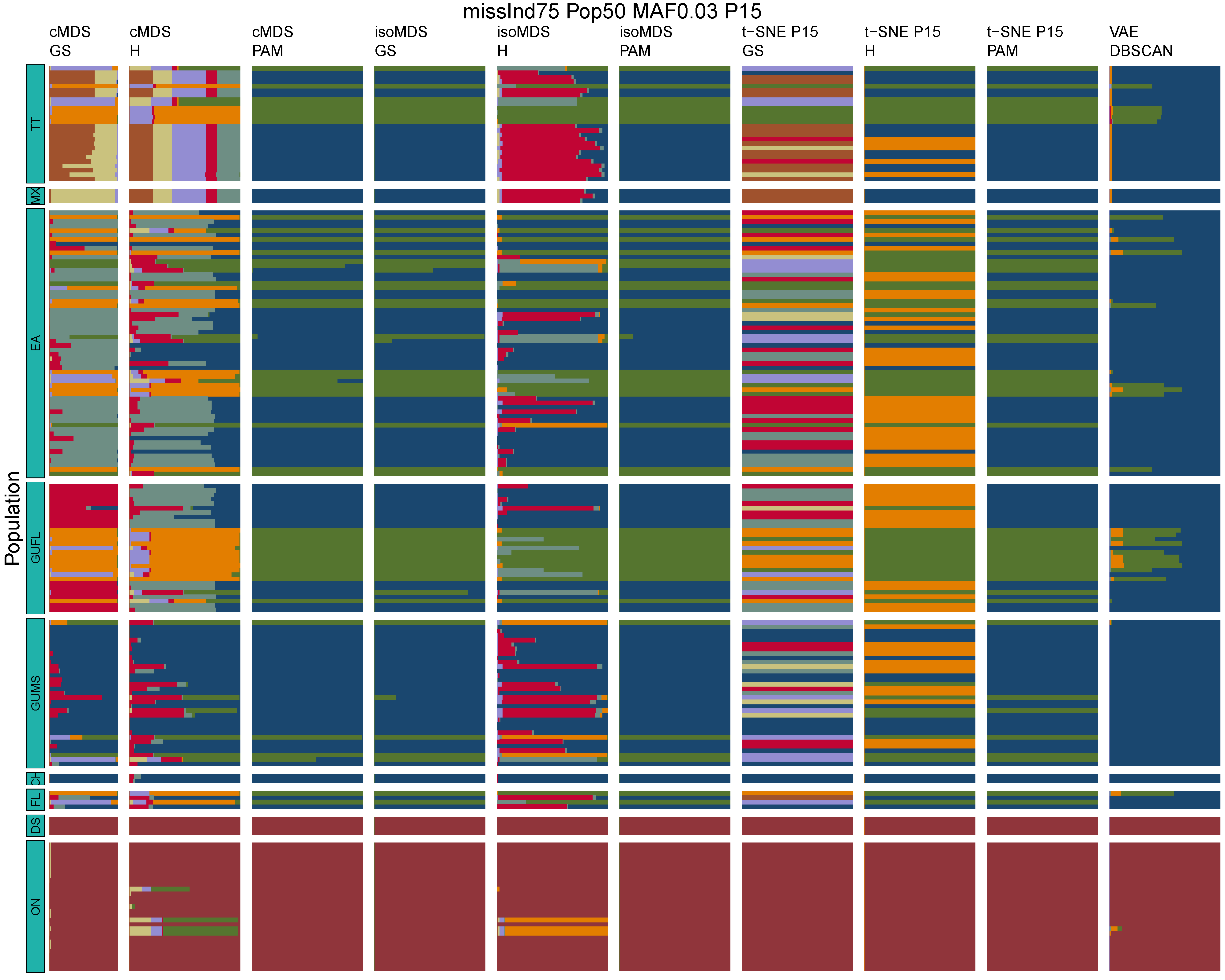

**Figure B39:** *Terrapene* unsupervised machine learning (UML) barplots depicting assignment proportions (x axis) among 100 replicates. Filtering parameters allowed maximum per-individual and per-population missing data and minimum minor allele frequency (MAF) filters. The included maximum missing data and minimum MAF filter proportions are shown in the plot title (missInd=per-individual, Pop=per-population filtering, MAF=MAF filter, P=perplexity). Each barplot on the page represents one UML algorithm: random forest, visualized with classical and isotonic multidimensional scaling (cMDS and isoMDS) ordination. t-SNE=t-distributed stochastic neighbor embedding with perplexity=15 (P15), and VAE=variational autoencoder. cMDS, isoMDS, and t-SNE were clustered with the partition around medoids (PAM) and hierarchical clustering (H) algorithms, and optimal *K* was chosen using PAM with the gap statistic (GS), H with the highest mean silhouette width (HMSW), and PAM with the HMSW. For VAE, clustering was performed, and optimal *K* chosen, with DBSCAN. *Terrapene* populations are separated by the horizontal white strips. Population IDs (blue left-aligned strip) include the ON=Ornate (*T. ornata ornata*), DS=Desert (*T. o. luteola*), FL=Florida (*T. c. bauri*), CH=Coahuilan (*T. coahuila*), GUMS=Gulf Coast (*T. carolina major*) from Mississippi, GUFL=Gulf Coast from the Florida Panhandle, EA=Woodland (*T. c. carolina*), MX=Mexican (*T. mexicana mexicana*), and TT=Three-toed (*T. m. triunguis*).
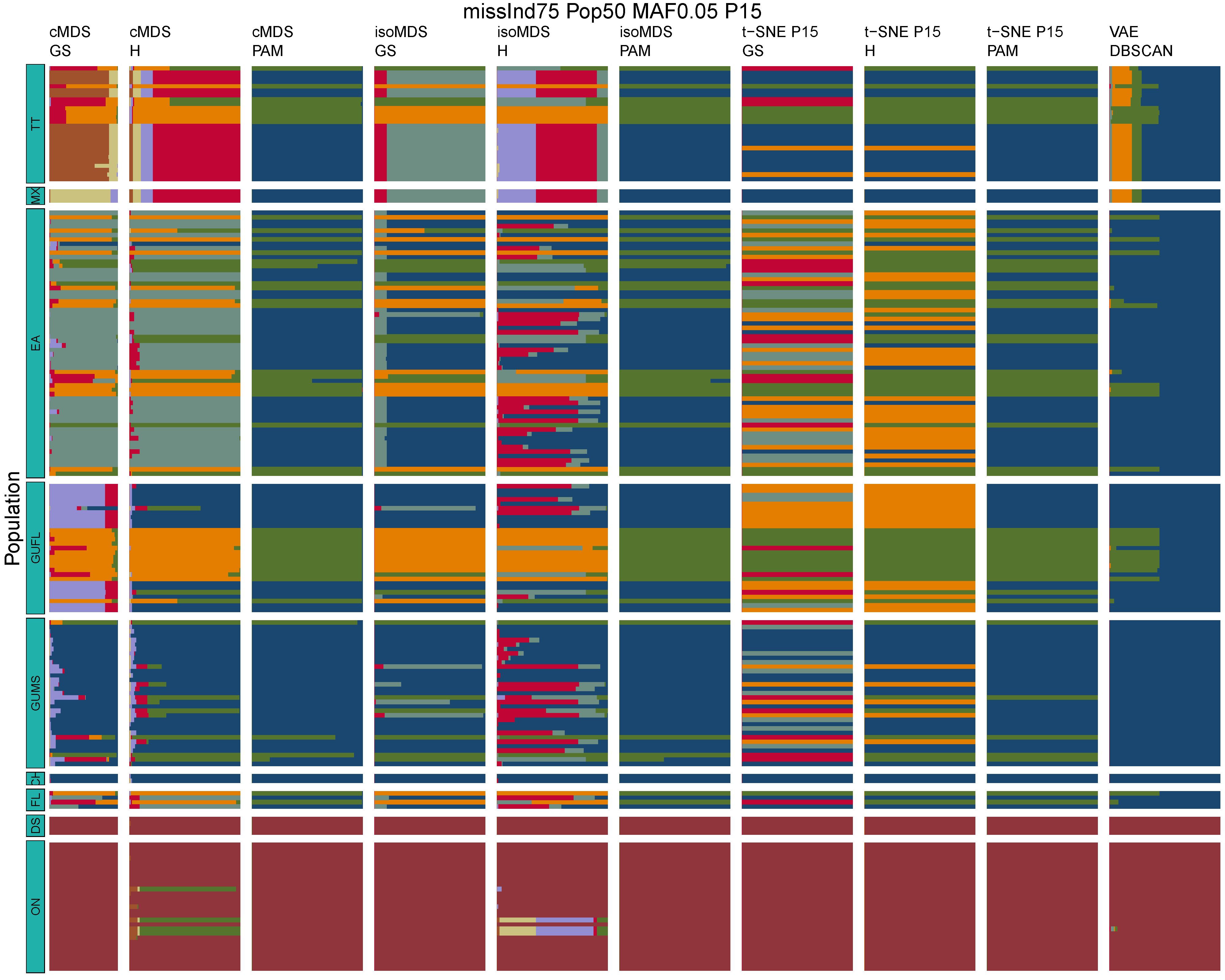

**Figure B40:** *Terrapene* unsupervised machine learning (UML) barplots depicting assignment proportions (x axis) among 100 replicates. Filtering parameters allowed maximum per-individual and per-population missing data and minimum minor allele frequency (MAF) filters. The included maximum missing data and minimum MAF filter proportions are shown in the plot title (missInd=per-individual, Pop=per-population filtering, MAF=MAF filter, P=perplexity). Each barplot on the page represents one UML algorithm: random forest, visualized with classical and isotonic multidimensional scaling (cMDS and isoMDS) ordination. t-SNE=t-distributed stochastic neighbor embedding with perplexity=15 (P15), and VAE=variational autoencoder. cMDS, isoMDS, and t-SNE were clustered with the partition around medoids (PAM) and hierarchical clustering (H) algorithms, and optimal *K* was chosen using PAM with the gap statistic (GS), H with the highest mean silhouette width (HMSW), and PAM with the HMSW. For VAE, clustering was performed, and optimal *K* chosen, with DBSCAN. *Terrapene* populations are separated by the horizontal white strips. Population IDs (blue left-aligned strip) include the ON=Ornate (*T. ornata ornata*), DS=Desert (*T. o. luteola*), FL=Florida (*T. c. bauri*), CH=Coahuilan (*T. coahuila*), GUMS=Gulf Coast (*T. carolina major*) from Mississippi, GUFL=Gulf Coast from the Florida Panhandle, EA=Woodland (*T. c. carolina*), MX=Mexican (*T. mexicana mexicana*), and TT=Three-toed (*T. m. triunguis*).
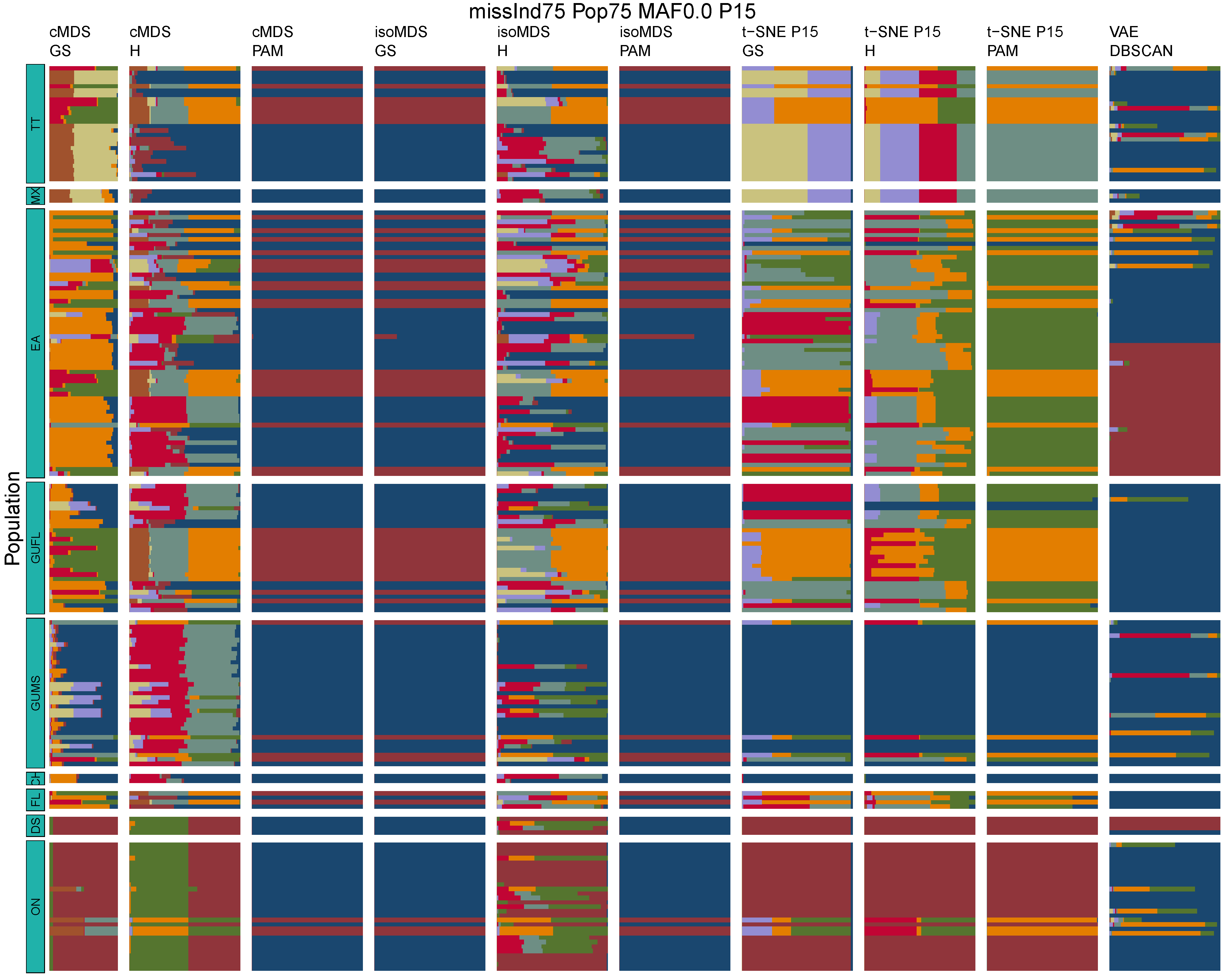

**Figure B41:** *Terrapene* unsupervised machine learning (UML) barplots depicting assignment proportions (x axis) among 100 replicates. Filtering parameters allowed maximum per-individual and per-population missing data and minimum minor allele frequency (MAF) filters. The included maximum missing data and minimum MAF filter proportions are shown in the plot title (missInd=per-individual, Pop=per-population filtering, MAF=MAF filter, P=perplexity). Each barplot on the page represents one UML algorithm: random forest, visualized with classical and isotonic multidimensional scaling (cMDS and isoMDS) ordination. t-SNE=t-distributed stochastic neighbor embedding with perplexity=15 (P15), and VAE=variational autoencoder. cMDS, isoMDS, and t-SNE were clustered with the partition around medoids (PAM) and hierarchical clustering (H) algorithms, and optimal *K* was chosen using PAM with the gap statistic (GS), H with the highest mean silhouette width (HMSW), and PAM with the HMSW. For VAE, clustering was performed, and optimal *K* chosen, with DBSCAN. *Terrapene* populations are separated by the horizontal white strips. Population IDs (blue left-aligned strip) include the ON=Ornate (*T. ornata ornata*), DS=Desert (*T. o. luteola*), FL=Florida (*T. c. bauri*), CH=Coahuilan (*T. coahuila*), GUMS=Gulf Coast (*T. carolina major*) from Mississippi, GUFL=Gulf Coast from the Florida Panhandle, EA=Woodland (*T. c. carolina*), MX=Mexican (*T. mexicana mexicana*), and TT=Three-toed (*T. m. triunguis*).
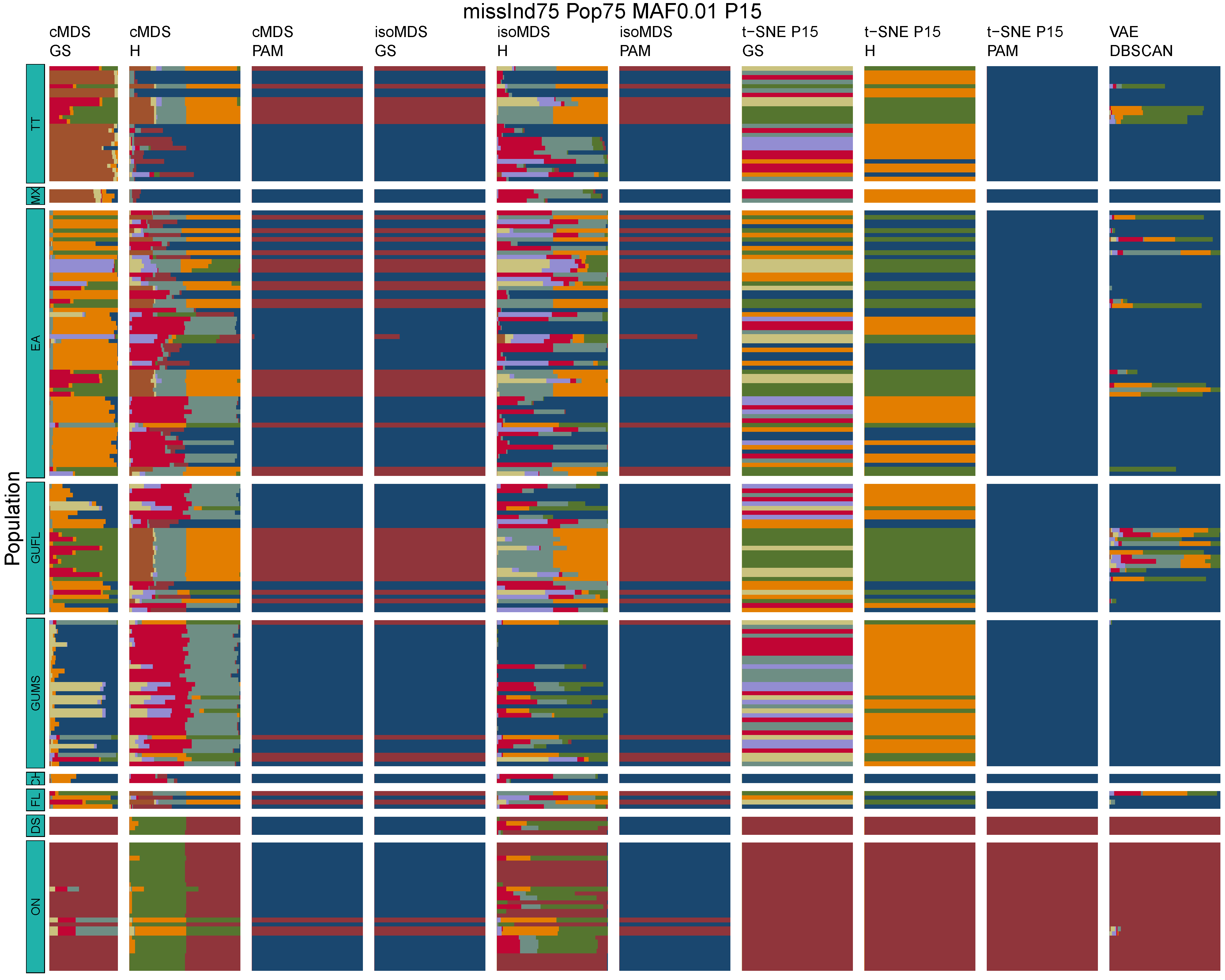

**Figure B42:** *Terrapene* unsupervised machine learning (UML) barplots depicting assignment proportions (x axis) among 100 replicates. Filtering parameters allowed maximum per-individual and per-population missing data and minimum minor allele frequency (MAF) filters. The included maximum missing data and minimum MAF filter proportions are shown in the plot title (missInd=per-individual, Pop=per-population filtering, MAF=MAF filter, P=perplexity). Each barplot on the page represents one UML algorithm: random forest, visualized with classical and isotonic multidimensional scaling (cMDS and isoMDS) ordination. t-SNE=t-distributed stochastic neighbor embedding with perplexity=15 (P15), and VAE=variational autoencoder. cMDS, isoMDS, and t-SNE were clustered with the partition around medoids (PAM) and hierarchical clustering (H) algorithms, and optimal *K* was chosen using PAM with the gap statistic (GS), H with the highest mean silhouette width (HMSW), and PAM with the HMSW. For VAE, clustering was performed, and optimal *K* chosen, with DBSCAN. *Terrapene* populations are separated by the horizontal white strips. Population IDs (blue left-aligned strip) include the ON=Ornate (*T. ornata ornata*), DS=Desert (*T. o. luteola*), FL=Florida (*T. c. bauri*), CH=Coahuilan (*T. coahuila*), GUMS=Gulf Coast (*T. carolina major*) from Mississippi, GUFL=Gulf Coast from the Florida Panhandle, EA=Woodland (*T. c. carolina*), MX=Mexican (*T. mexicana mexicana*), and TT=Three-toed (*T. m. triunguis*).
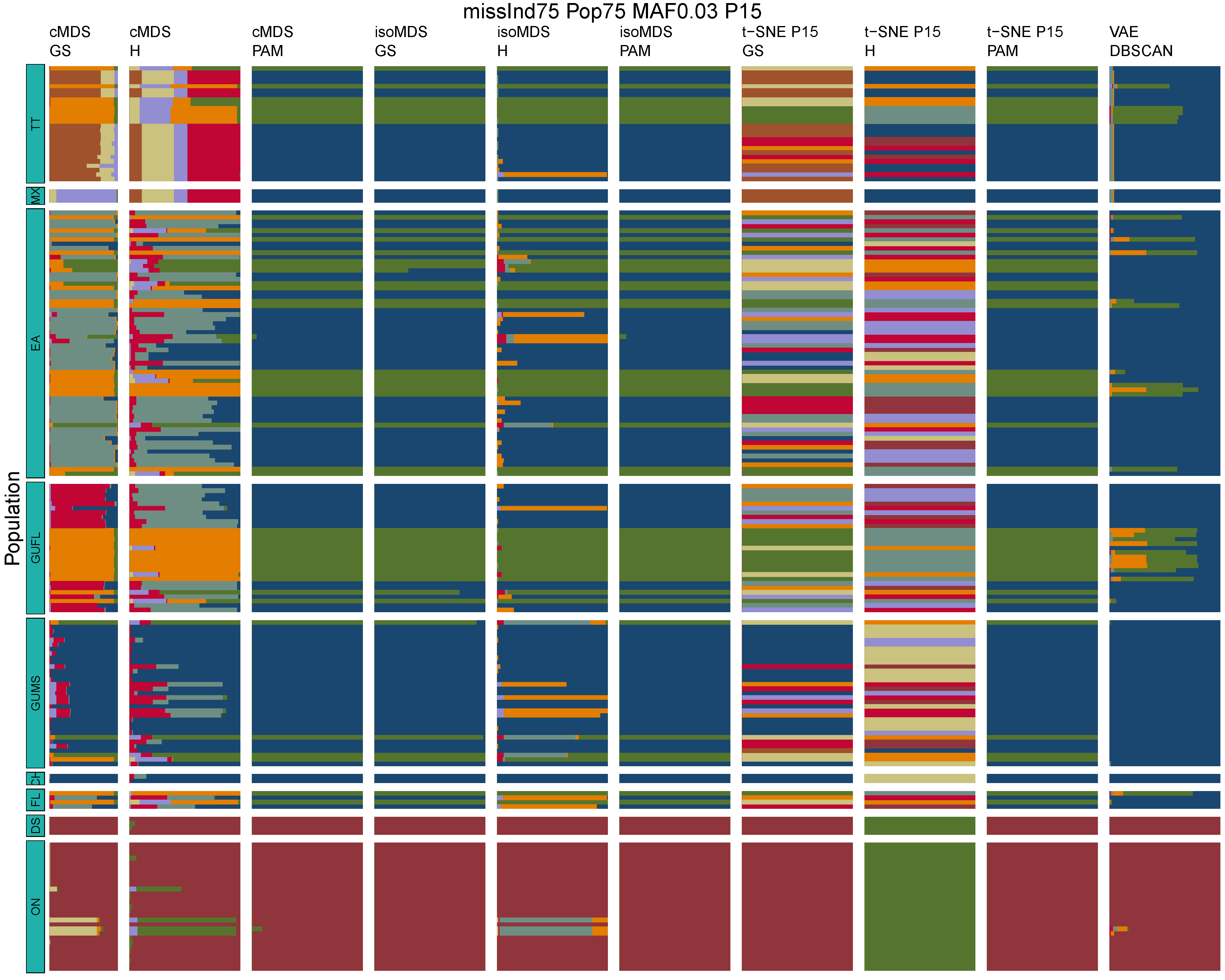

**Figure B43:** *Terrapene* unsupervised machine learning (UML) barplots depicting assignment proportions (x axis) among 100 replicates. Filtering parameters allowed maximum per-individual and per-population missing data and minimum minor allele frequency (MAF) filters. The included maximum missing data and minimum MAF filter proportions are shown in the plot title (missInd=per-individual, Pop=per-population filtering, MAF=MAF filter, P=perplexity). Each barplot on the page represents one UML algorithm: random forest, visualized with classical and isotonic multidimensional scaling (cMDS and isoMDS) ordination. t-SNE=t-distributed stochastic neighbor embedding with perplexity=15 (P15), and VAE=variational autoencoder. cMDS, isoMDS, and t-SNE were clustered with the partition around medoids (PAM) and hierarchical clustering (H) algorithms, and optimal *K* was chosen using PAM with the gap statistic (GS), H with the highest mean silhouette width (HMSW), and PAM with the HMSW. For VAE, clustering was performed, and optimal *K* chosen, with DBSCAN. *Terrapene* populations are separated by the horizontal white strips. Population IDs (blue left-aligned strip) include the ON=Ornate (*T. ornata ornata*), DS=Desert (*T. o. luteola*), FL=Florida (*T. c. bauri*), CH=Coahuilan (*T. coahuila*), GUMS=Gulf Coast (*T. carolina major*) from Mississippi, GUFL=Gulf Coast from the Florida Panhandle, EA=Woodland (*T. c. carolina*), MX=Mexican (*T. mexicana mexicana*), and TT=Three-toed (*T. m. triunguis*).
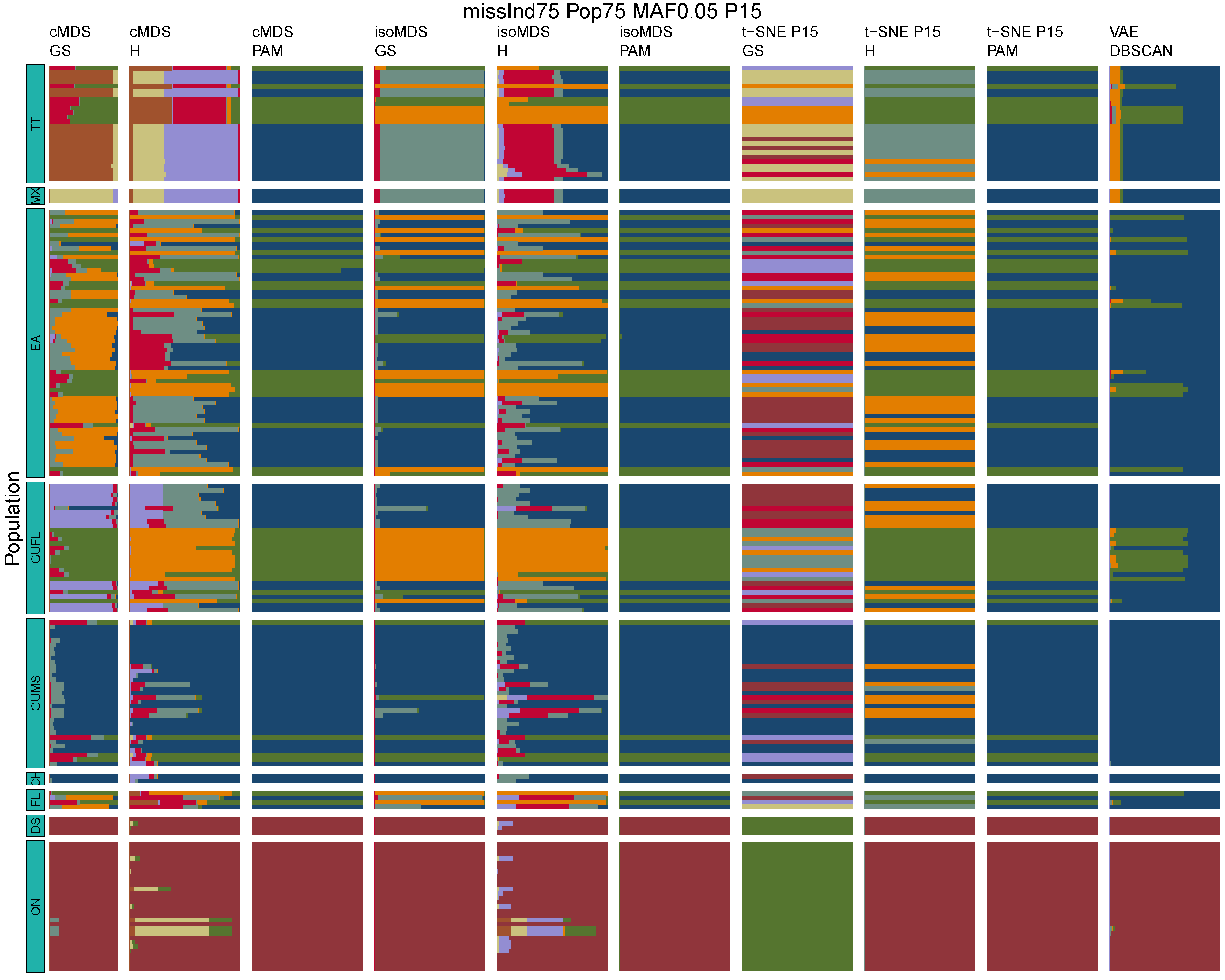

**Figure B44:** *Terrapene* unsupervised machine learning (UML) barplots depicting assignment proportions (x axis) among 100 replicates. Filtering parameters allowed maximum per-individual and per-population missing data and minimum minor allele frequency (MAF) filters. The included maximum missing data and minimum MAF filter proportions are shown in the plot title (missInd=per-individual, Pop=per-population filtering, MAF=MAF filter, P=perplexity). Each barplot on the page represents one UML algorithm: random forest, visualized with classical and isotonic multidimensional scaling (cMDS and isoMDS) ordination. t-SNE=t-distributed stochastic neighbor embedding with perplexity=15 (P15), and VAE=variational autoencoder. cMDS, isoMDS, and t-SNE were clustered with the partition around medoids (PAM) and hierarchical clustering (H) algorithms, and optimal *K* was chosen using PAM with the gap statistic (GS), H with the highest mean silhouette width (HMSW), and PAM with the HMSW. For VAE, clustering was performed, and optimal *K* chosen, with DBSCAN. *Terrapene* populations are separated by the horizontal white strips. Population IDs (blue left-aligned strip) include the ON=Ornate (*T. ornata ornata*), DS=Desert (*T. o. luteola*), FL=Florida (*T. c. bauri*), CH=Coahuilan (*T. coahuila*), GUMS=Gulf Coast (*T. carolina major*) from Mississippi, GUFL=Gulf Coast from the Florida Panhandle, EA=Woodland (*T. c. carolina*), MX=Mexican (*T. mexicana mexicana*), and TT=Three-toed (*T. m. triunguis*).
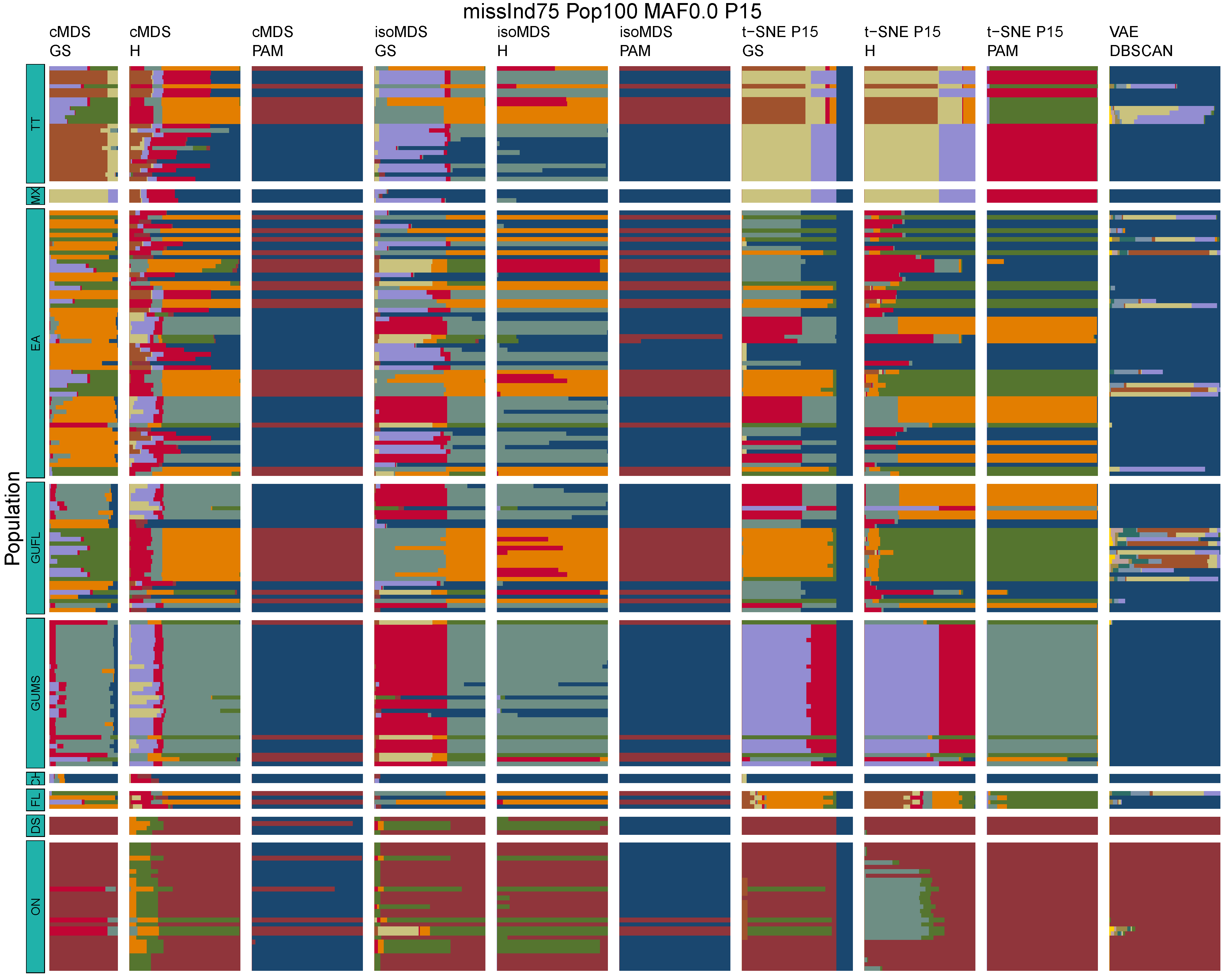

**Figure B45:** *Terrapene* unsupervised machine learning (UML) barplots depicting assignment proportions (x axis) among 100 replicates. Filtering parameters allowed maximum per-individual and per-population missing data and minimum minor allele frequency (MAF) filters. The included maximum missing data and minimum MAF filter proportions are shown in the plot title (missInd=per-individual, Pop=per-population filtering, MAF=MAF filter, P=perplexity). Each barplot on the page represents one UML algorithm: random forest, visualized with classical and isotonic multidimensional scaling (cMDS and isoMDS) ordination. t-SNE=t-distributed stochastic neighbor embedding with perplexity=15 (P15), and VAE=variational autoencoder. cMDS, isoMDS, and t-SNE were clustered with the partition around medoids (PAM) and hierarchical clustering (H) algorithms, and optimal *K* was chosen using PAM with the gap statistic (GS), H with the highest mean silhouette width (HMSW), and PAM with the HMSW. For VAE, clustering was performed, and optimal *K* chosen, with DBSCAN. *Terrapene* populations are separated by the horizontal white strips. Population IDs (blue left-aligned strip) include the ON=Ornate (*T. ornata ornata*), DS=Desert (*T. o. luteola*), FL=Florida (*T. c. bauri*), CH=Coahuilan (*T. coahuila*), GUMS=Gulf Coast (*T. carolina major*) from Mississippi, GUFL=Gulf Coast from the Florida Panhandle, EA=Woodland (*T. c. carolina*), MX=Mexican (*T. mexicana mexicana*), and TT=Three-toed (*T. m. triunguis*).
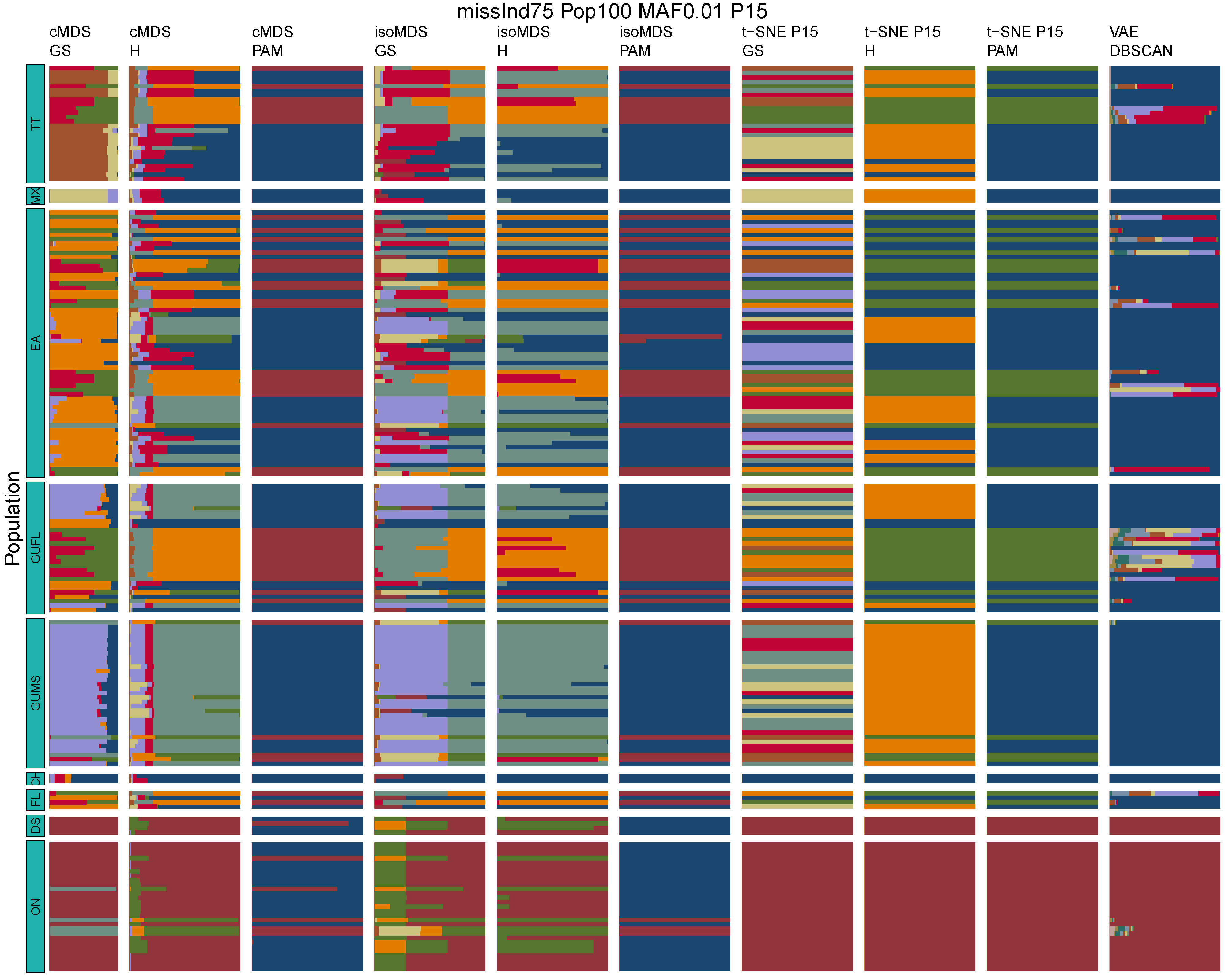

**Figure B46:** *Terrapene* unsupervised machine learning (UML) barplots depicting assignment proportions (x axis) among 100 replicates. Filtering parameters allowed maximum per-individual and per-population missing data and minimum minor allele frequency (MAF) filters. The included maximum missing data and minimum MAF filter proportions are shown in the plot title (missInd=per-individual, Pop=per-population filtering, MAF=MAF filter, P=perplexity). Each barplot on the page represents one UML algorithm: random forest, visualized with classical and isotonic multidimensional scaling (cMDS and isoMDS) ordination. t-SNE=t-distributed stochastic neighbor embedding with perplexity=15 (P15), and VAE=variational autoencoder. cMDS, isoMDS, and t-SNE were clustered with the partition around medoids (PAM) and hierarchical clustering (H) algorithms, and optimal *K* was chosen using PAM with the gap statistic (GS), H with the highest mean silhouette width (HMSW), and PAM with the HMSW. For VAE, clustering was performed, and optimal *K* chosen, with DBSCAN. *Terrapene* populations are separated by the horizontal white strips. Population IDs (blue left-aligned strip) include the ON=Ornate (*T. ornata ornata*), DS=Desert (*T. o. luteola*), FL=Florida (*T. c. bauri*), CH=Coahuilan (*T. coahuila*), GUMS=Gulf Coast (*T. carolina major*) from Mississippi, GUFL=Gulf Coast from the Florida Panhandle, EA=Woodland (*T. c. carolina*), MX=Mexican (*T. mexicana mexicana*), and TT=Three-toed (*T. m. triunguis*).
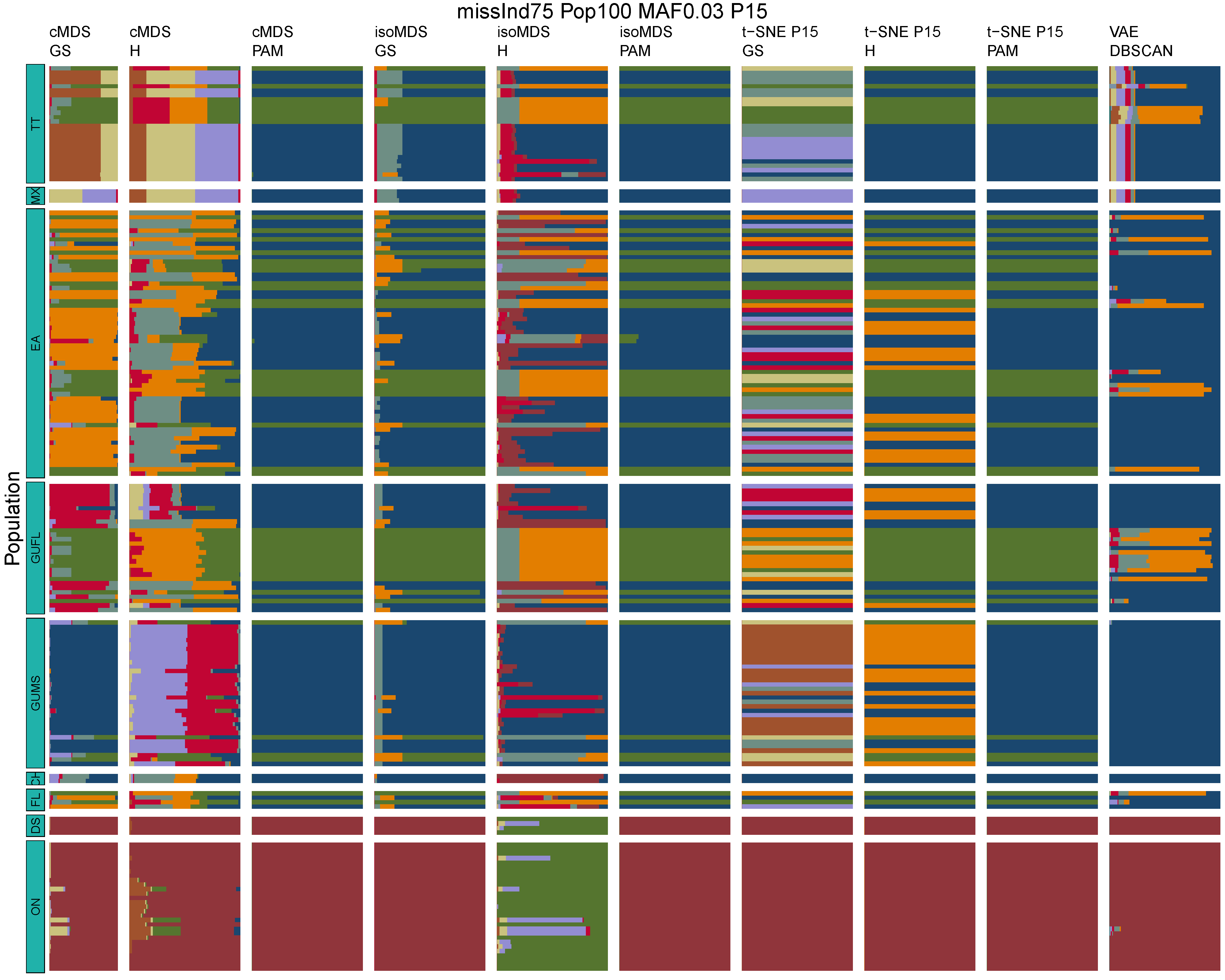

**Figure B47:** *Terrapene* unsupervised machine learning (UML) barplots depicting assignment proportions (x axis) among 100 replicates. Filtering parameters allowed maximum per-individual and per-population missing data and minimum minor allele frequency (MAF) filters. The included maximum missing data and minimum MAF filter proportions are shown in the plot title (missInd=per-individual, Pop=per-population filtering, MAF=MAF filter, P=perplexity). Each barplot on the page represents one UML algorithm: random forest, visualized with classical and isotonic multidimensional scaling (cMDS and isoMDS) ordination. t-SNE=t-distributed stochastic neighbor embedding with perplexity=15 (P15), and VAE=variational autoencoder. cMDS, isoMDS, and t-SNE were clustered with the partition around medoids (PAM) and hierarchical clustering (H) algorithms, and optimal *K* was chosen using PAM with the gap statistic (GS), H with the highest mean silhouette width (HMSW), and PAM with the HMSW. For VAE, clustering was performed, and optimal *K* chosen, with DBSCAN. *Terrapene* populations are separated by the horizontal white strips. Population IDs (blue left-aligned strip) include the ON=Ornate (*T. ornata ornata*), DS=Desert (*T. o. luteola*), FL=Florida (*T. c. bauri*), CH=Coahuilan (*T. coahuila*), GUMS=Gulf Coast (*T. carolina major*) from Mississippi, GUFL=Gulf Coast from the Florida Panhandle, EA=Woodland (*T. c. carolina*), MX=Mexican (*T. mexicana mexicana*), and TT=Three-toed (*T. m. triunguis*).
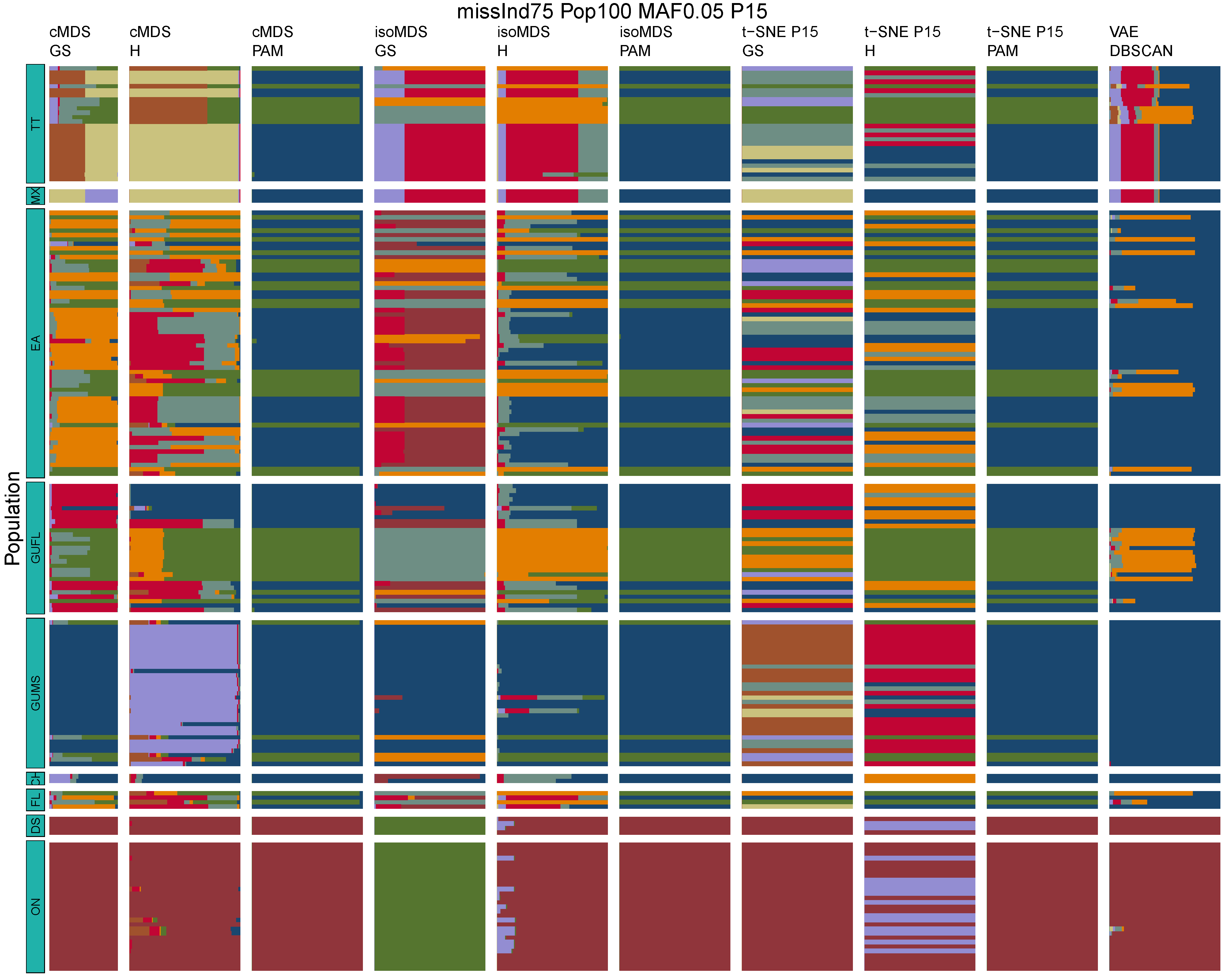

**Figure B48:** *Terrapene* unsupervised machine learning (UML) barplots depicting assignment proportions (x axis) among 100 replicates. Filtering parameters allowed maximum per-individual and per-population missing data and minimum minor allele frequency (MAF) filters. The included maximum missing data and minimum MAF filter proportions are shown in the plot title (missInd=per-individual, Pop=per-population filtering, MAF=MAF filter, P=perplexity). Each barplot on the page represents one UML algorithm: random forest, visualized with classical and isotonic multidimensional scaling (cMDS and isoMDS) ordination. t-SNE=t-distributed stochastic neighbor embedding with perplexity=15 (P15), and VAE=variational autoencoder. cMDS, isoMDS, and t-SNE were clustered with the partition around medoids (PAM) and hierarchical clustering (H) algorithms, and optimal *K* was chosen using PAM with the gap statistic (GS), H with the highest mean silhouette width (HMSW), and PAM with the HMSW. For VAE, clustering was performed, and optimal *K* chosen, with DBSCAN. *Terrapene* populations are separated by the horizontal white strips. Population IDs (blue left-aligned strip) include the ON=Ornate (*T. ornata ornata*), DS=Desert (*T. o. luteola*), FL=Florida (*T. c. bauri*), CH=Coahuilan (*T. coahuila*), GUMS=Gulf Coast (*T. carolina major*) from Mississippi, GUFL=Gulf Coast from the Florida Panhandle, EA=Woodland (*T. c. carolina*), MX=Mexican (*T. mexicana mexicana*), and TT=Three-toed (*T. m. triunguis*).
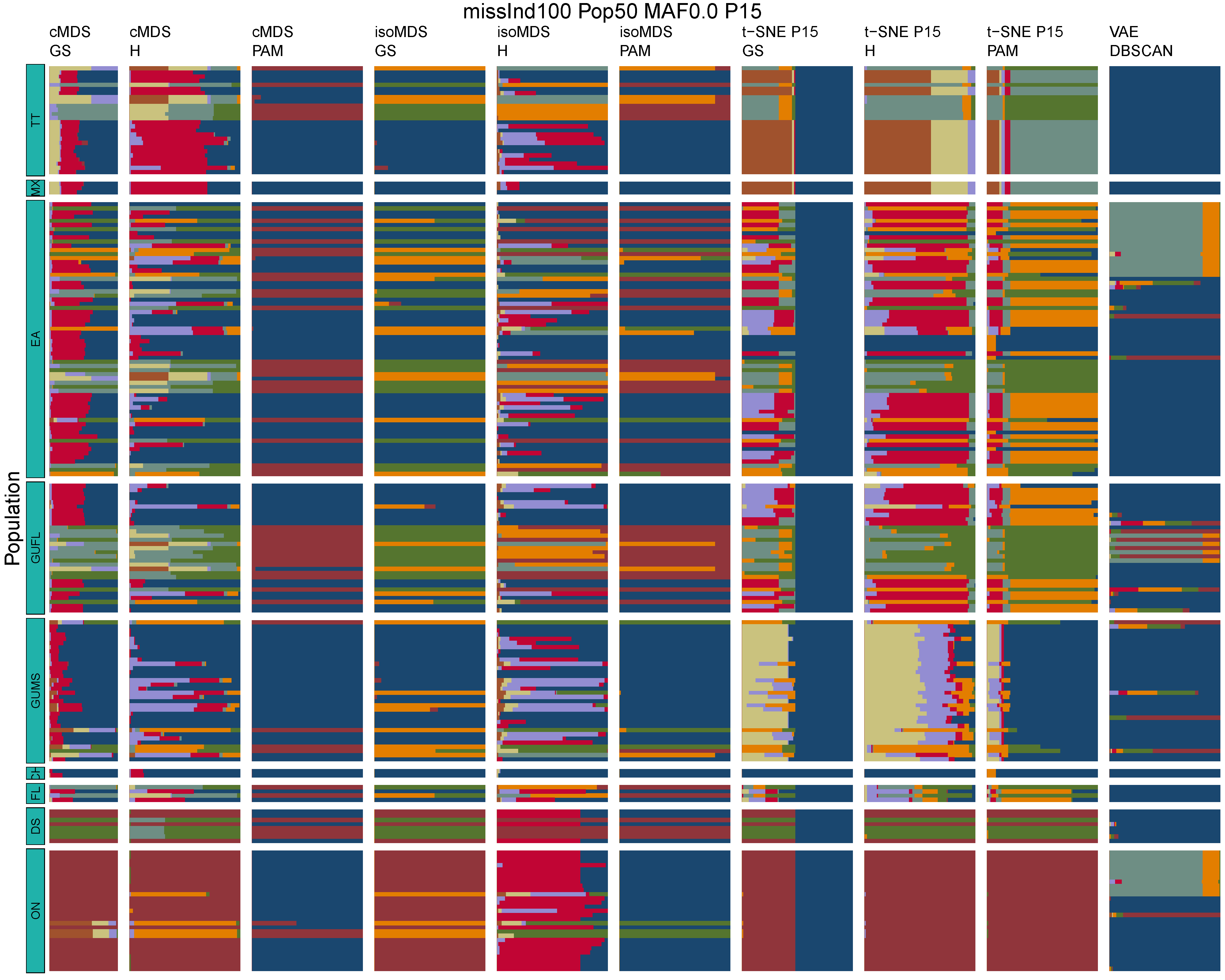

**Figure B49:** *Terrapene* unsupervised machine learning (UML) barplots depicting assignment proportions (x axis) among 100 replicates. Filtering parameters allowed maximum per-individual and per-population missing data and minimum minor allele frequency (MAF) filters. The included maximum missing data and minimum MAF filter proportions are shown in the plot title (missInd=per-individual, Pop=per-population filtering, MAF=MAF filter, P=perplexity). Each barplot on the page represents one UML algorithm: random forest, visualized with classical and isotonic multidimensional scaling (cMDS and isoMDS) ordination. t-SNE=t-distributed stochastic neighbor embedding with perplexity=15 (P15), and VAE=variational autoencoder. cMDS, isoMDS, and t-SNE were clustered with the partition around medoids (PAM) and hierarchical clustering (H) algorithms, and optimal *K* was chosen using PAM with the gap statistic (GS), H with the highest mean silhouette width (HMSW), and PAM with the HMSW. For VAE, clustering was performed, and optimal *K* chosen, with DBSCAN. *Terrapene* populations are separated by the horizontal white strips. Population IDs (blue left-aligned strip) include the ON=Ornate (*T. ornata ornata*), DS=Desert (*T. o. luteola*), FL=Florida (*T. c. bauri*), CH=Coahuilan (*T. coahuila*), GUMS=Gulf Coast (*T. carolina major*) from Mississippi, GUFL=Gulf Coast from the Florida Panhandle, EA=Woodland (*T. c. carolina*), MX=Mexican (*T. mexicana mexicana*), and TT=Three-toed (*T. m. triunguis*).
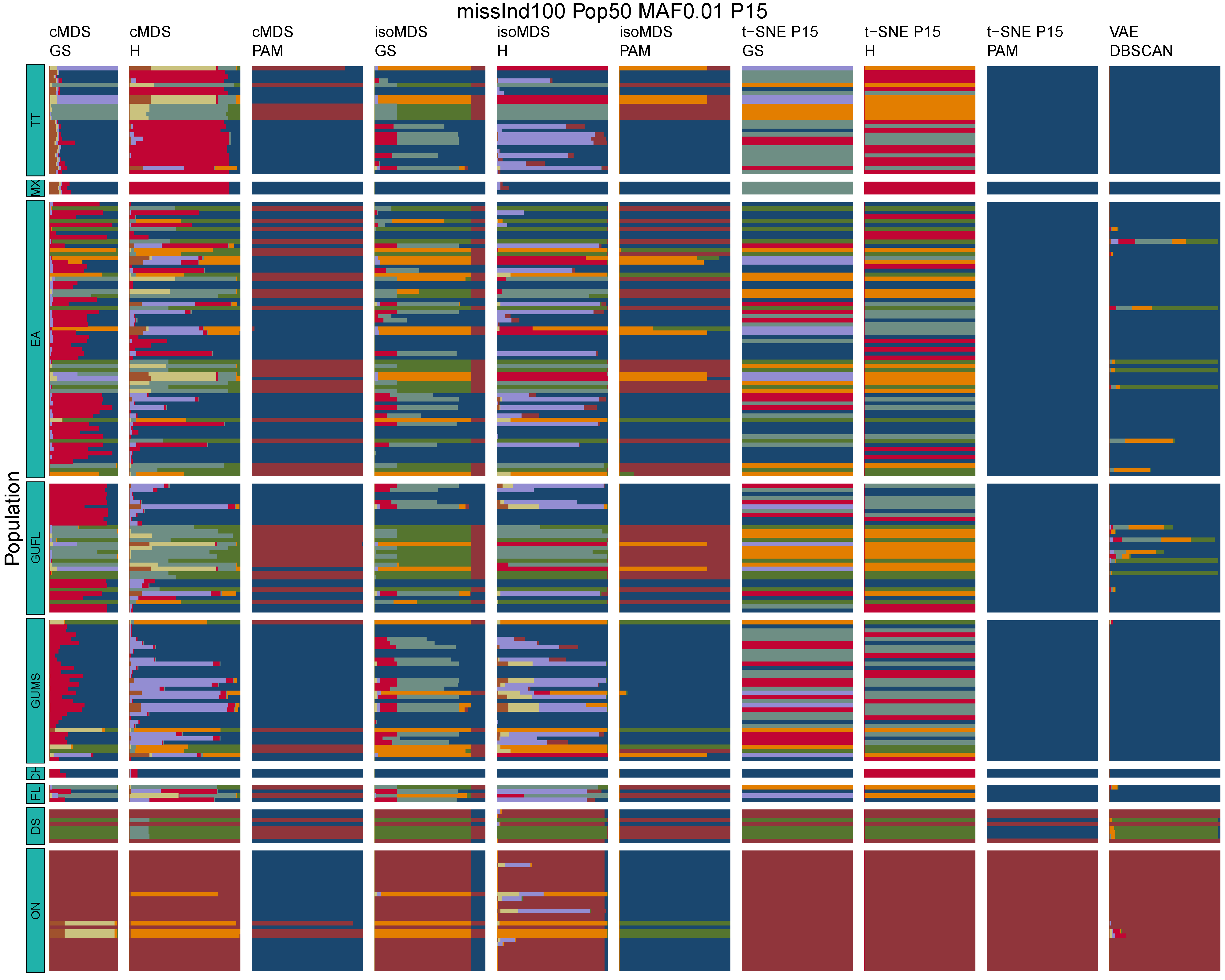

**Figure B50:** *Terrapene* unsupervised machine learning (UML) barplots depicting assignment proportions (x axis) among 100 replicates. Filtering parameters allowed maximum per-individual and per-population missing data and minimum minor allele frequency (MAF) filters. The included maximum missing data and minimum MAF filter proportions are shown in the plot title (missInd=per-individual, Pop=per-population filtering, MAF=MAF filter, P=perplexity). Each barplot on the page represents one UML algorithm: random forest, visualized with classical and isotonic multidimensional scaling (cMDS and isoMDS) ordination. t-SNE=t-distributed stochastic neighbor embedding with perplexity=15 (P15), and VAE=variational autoencoder. cMDS, isoMDS, and t-SNE were clustered with the partition around medoids (PAM) and hierarchical clustering (H) algorithms, and optimal *K* was chosen using PAM with the gap statistic (GS), H with the highest mean silhouette width (HMSW), and PAM with the HMSW. For VAE, clustering was performed, and optimal *K* chosen, with DBSCAN. *Terrapene* populations are separated by the horizontal white strips. Population IDs (blue left-aligned strip) include the ON=Ornate (*T. ornata ornata*), DS=Desert (*T. o. luteola*), FL=Florida (*T. c. bauri*), CH=Coahuilan (*T. coahuila*), GUMS=Gulf Coast (*T. carolina major*) from Mississippi, GUFL=Gulf Coast from the Florida Panhandle, EA=Woodland (*T. c. carolina*), MX=Mexican (*T. mexicana mexicana*), and TT=Three-toed (*T. m. triunguis*).
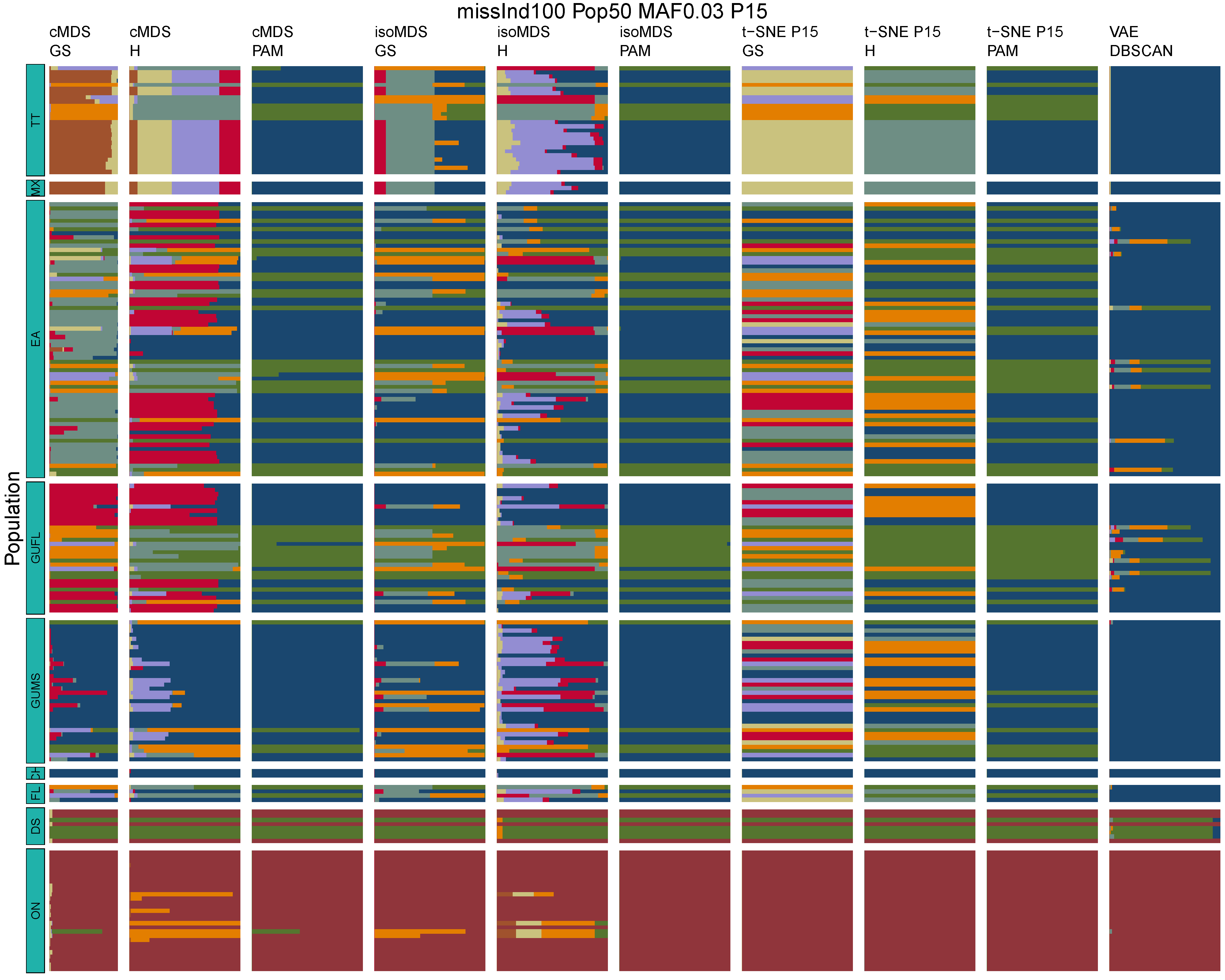

**Figure B51:** *Terrapene* unsupervised machine learning (UML) barplots depicting assignment proportions (x axis) among 100 replicates. Filtering parameters allowed maximum per-individual and per-population missing data and minimum minor allele frequency (MAF) filters. The included maximum missing data and minimum MAF filter proportions are shown in the plot title (missInd=per-individual, Pop=per-population filtering, MAF=MAF filter, P=perplexity). Each barplot on the page represents one UML algorithm: random forest, visualized with classical and isotonic multidimensional scaling (cMDS and isoMDS) ordination. t-SNE=t-distributed stochastic neighbor embedding with perplexity=15 (P15), and VAE=variational autoencoder. cMDS, isoMDS, and t-SNE were clustered with the partition around medoids (PAM) and hierarchical clustering (H) algorithms, and optimal *K* was chosen using PAM with the gap statistic (GS), H with the highest mean silhouette width (HMSW), and PAM with the HMSW. For VAE, clustering was performed, and optimal *K* chosen, with DBSCAN. *Terrapene* populations are separated by the horizontal white strips. Population IDs (blue left-aligned strip) include the ON=Ornate (*T. ornata ornata*), DS=Desert (*T. o. luteola*), FL=Florida (*T. c. bauri*), CH=Coahuilan (*T. coahuila*), GUMS=Gulf Coast (*T. carolina major*) from Mississippi, GUFL=Gulf Coast from the Florida Panhandle, EA=Woodland (*T. c. carolina*), MX=Mexican (*T. mexicana mexicana*), and TT=Three-toed (*T. m. triunguis*).
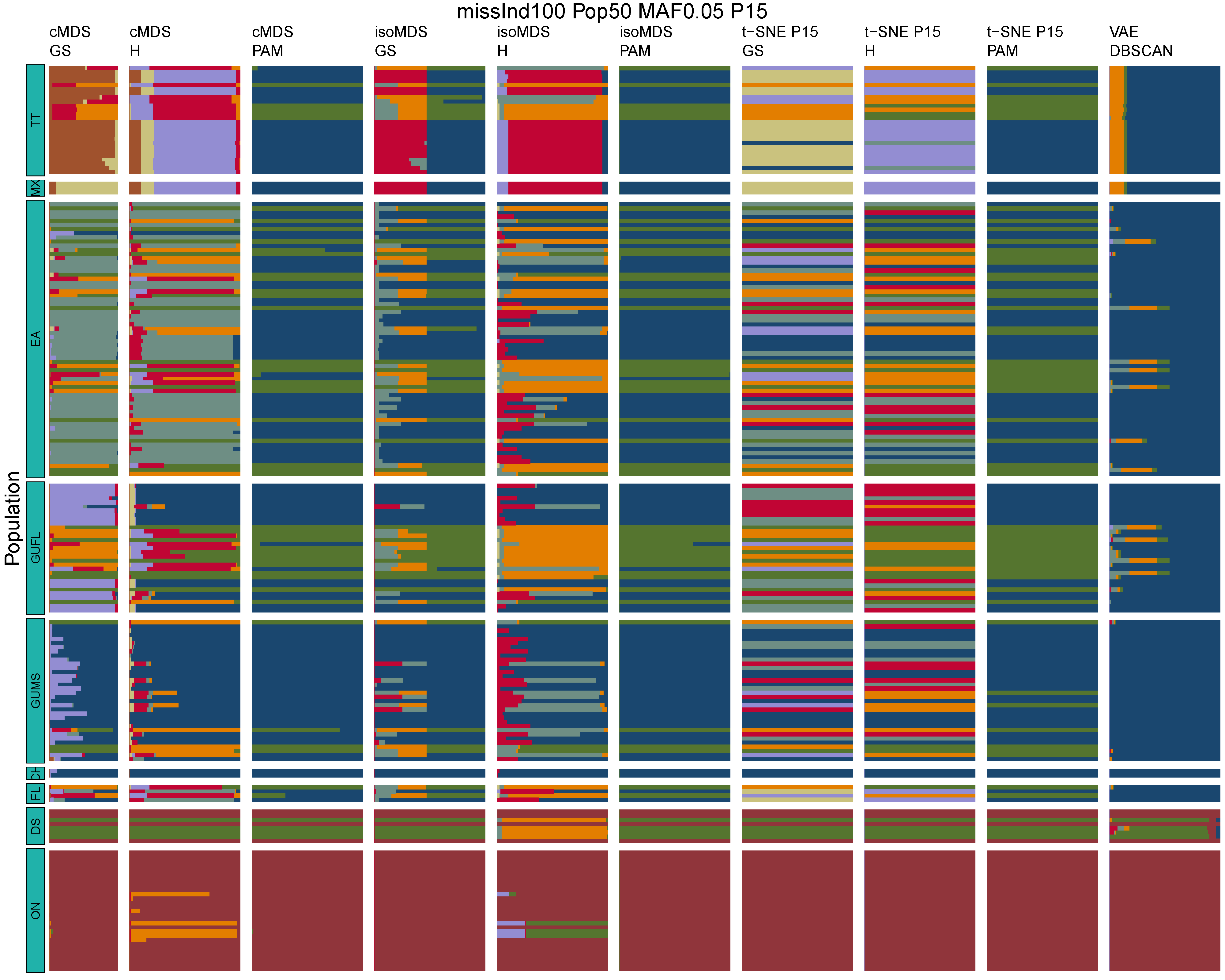

**Figure B52:** *Terrapene* unsupervised machine learning (UML) barplots depicting assignment proportions (x axis) among 100 replicates. Filtering parameters allowed maximum per-individual and per-population missing data and minimum minor allele frequency (MAF) filters. The included maximum missing data and minimum MAF filter proportions are shown in the plot title (missInd=per-individual, Pop=per-population filtering, MAF=MAF filter, P=perplexity). Each barplot on the page represents one UML algorithm: random forest, visualized with classical and isotonic multidimensional scaling (cMDS and isoMDS) ordination. t-SNE=t-distributed stochastic neighbor embedding with perplexity=15 (P15), and VAE=variational autoencoder. cMDS, isoMDS, and t-SNE were clustered with the partition around medoids (PAM) and hierarchical clustering (H) algorithms, and optimal *K* was chosen using PAM with the gap statistic (GS), H with the highest mean silhouette width (HMSW), and PAM with the HMSW. For VAE, clustering was performed, and optimal *K* chosen, with DBSCAN. *Terrapene* populations are separated by the horizontal white strips. Population IDs (blue left-aligned strip) include the ON=Ornate (*T. ornata ornata*), DS=Desert (*T. o. luteola*), FL=Florida (*T. c. bauri*), CH=Coahuilan (*T. coahuila*), GUMS=Gulf Coast (*T. carolina major*) from Mississippi, GUFL=Gulf Coast from the Florida Panhandle, EA=Woodland (*T. c. carolina*), MX=Mexican (*T. mexicana mexicana*), and TT=Three-toed (*T. m. triunguis*).
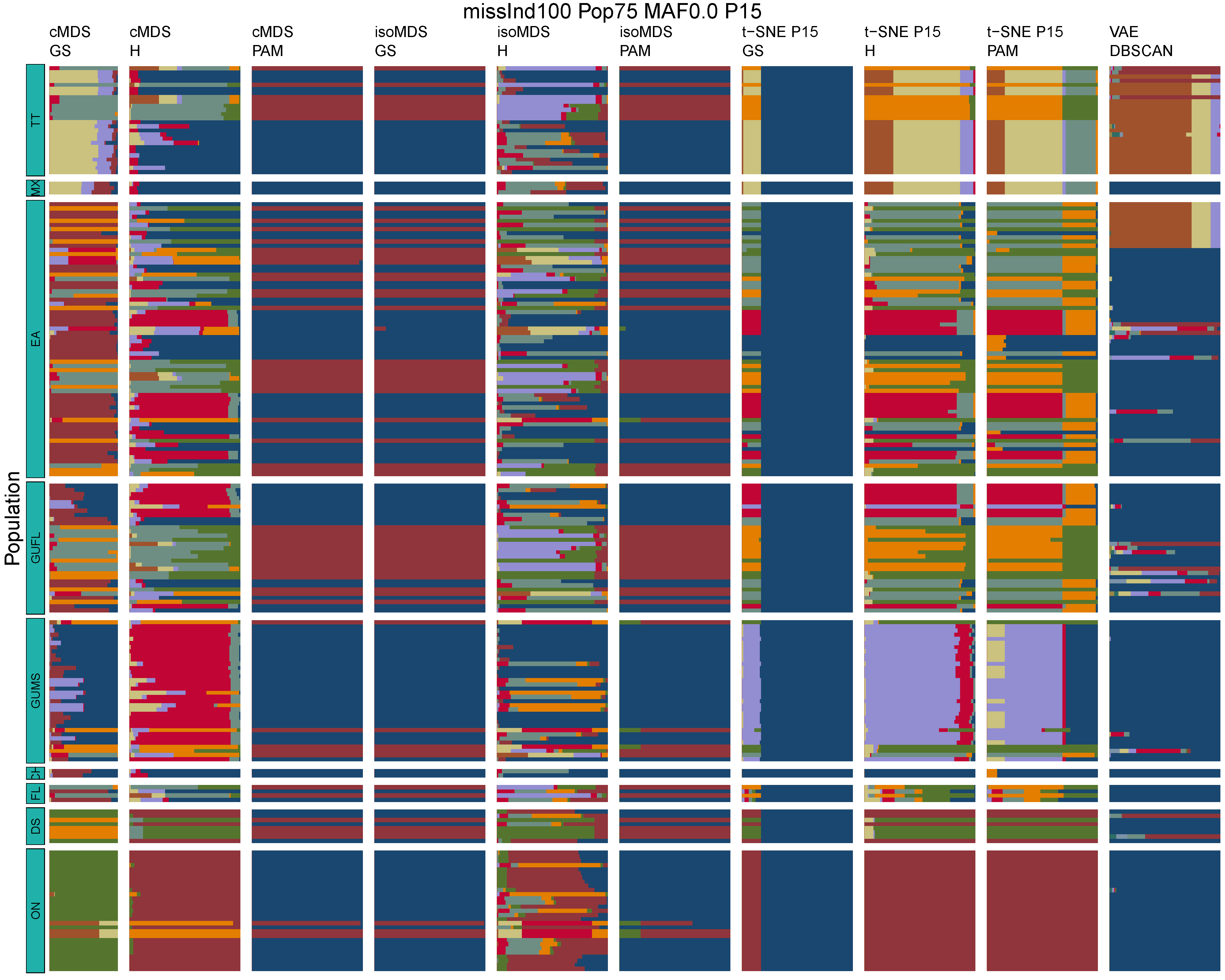

**Figure B53:** *Terrapene* unsupervised machine learning (UML) barplots depicting assignment proportions (x axis) among 100 replicates. Filtering parameters allowed maximum per-individual and per-population missing data and minimum minor allele frequency (MAF) filters. The included maximum missing data and minimum MAF filter proportions are shown in the plot title (missInd=per-individual, Pop=per-population filtering, MAF=MAF filter, P=perplexity). Each barplot on the page represents one UML algorithm: random forest, visualized with classical and isotonic multidimensional scaling (cMDS and isoMDS) ordination. t-SNE=t-distributed stochastic neighbor embedding with perplexity=15 (P15), and VAE=variational autoencoder. cMDS, isoMDS, and t-SNE were clustered with the partition around medoids (PAM) and hierarchical clustering (H) algorithms, and optimal *K* was chosen using PAM with the gap statistic (GS), H with the highest mean silhouette width (HMSW), and PAM with the HMSW. For VAE, clustering was performed, and optimal *K* chosen, with DBSCAN. *Terrapene* populations are separated by the horizontal white strips. Population IDs (blue left-aligned strip) include the ON=Ornate (*T. ornata ornata*), DS=Desert (*T. o. luteola*), FL=Florida (*T. c. bauri*), CH=Coahuilan (*T. coahuila*), GUMS=Gulf Coast (*T. carolina major*) from Mississippi, GUFL=Gulf Coast from the Florida Panhandle, EA=Woodland (*T. c. carolina*), MX=Mexican (*T. mexicana mexicana*), and TT=Three-toed (*T. m. triunguis*).
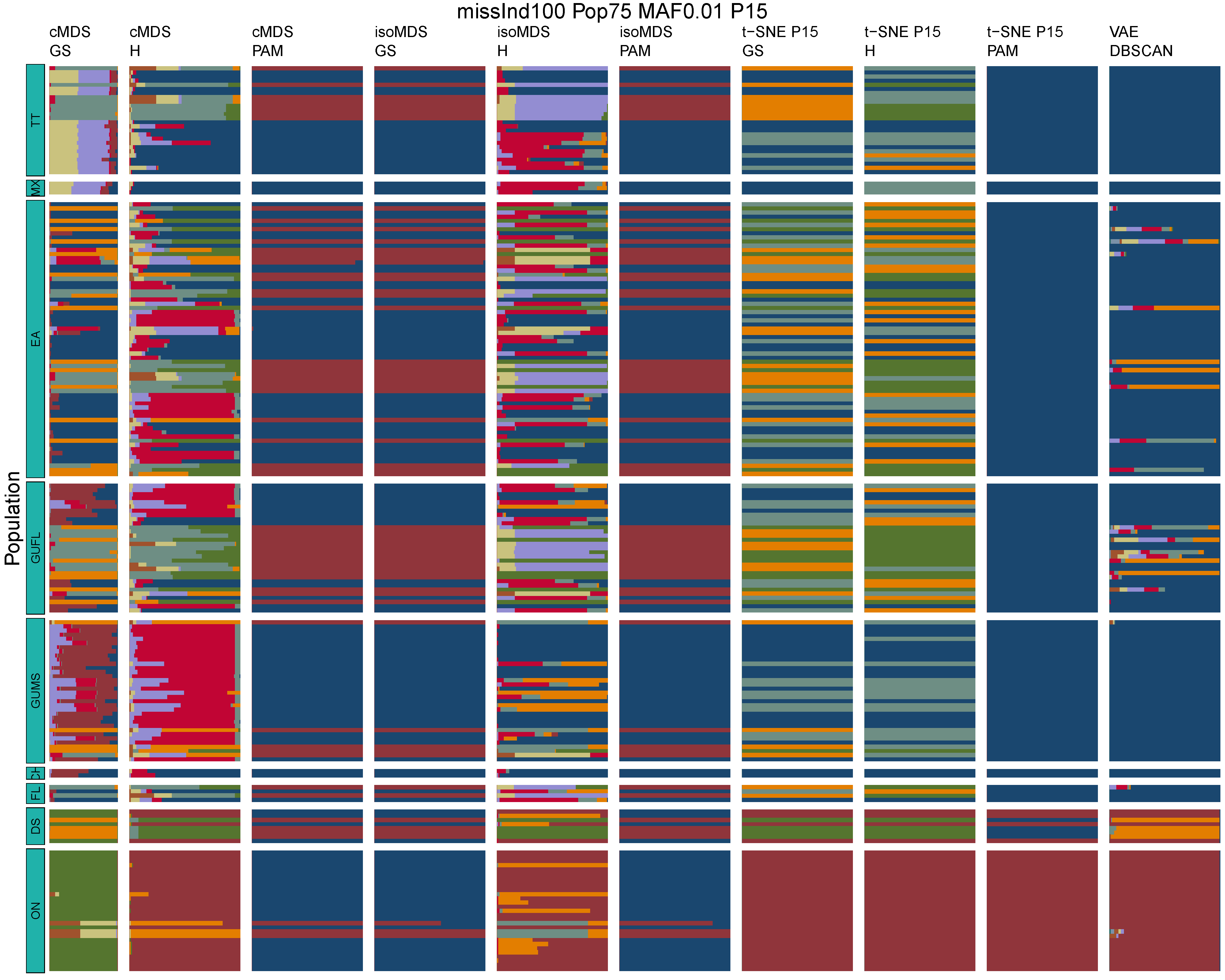

**Figure B54:** *Terrapene* unsupervised machine learning (UML) barplots depicting assignment proportions (x axis) among 100 replicates. Filtering parameters allowed maximum per-individual and per-population missing data and minimum minor allele frequency (MAF) filters. The included maximum missing data and minimum MAF filter proportions are shown in the plot title (missInd=per-individual, Pop=per-population filtering, MAF=MAF filter, P=perplexity). Each barplot on the page represents one UML algorithm: random forest, visualized with classical and isotonic multidimensional scaling (cMDS and isoMDS) ordination. t-SNE=t-distributed stochastic neighbor embedding with perplexity=15 (P15), and VAE=variational autoencoder. cMDS, isoMDS, and t-SNE were clustered with the partition around medoids (PAM) and hierarchical clustering (H) algorithms, and optimal *K* was chosen using PAM with the gap statistic (GS), H with the highest mean silhouette width (HMSW), and PAM with the HMSW. For VAE, clustering was performed, and optimal *K* chosen, with DBSCAN. *Terrapene* populations are separated by the horizontal white strips. Population IDs (blue left-aligned strip) include the ON=Ornate (*T. ornata ornata*), DS=Desert (*T. o. luteola*), FL=Florida (*T. c. bauri*), CH=Coahuilan (*T. coahuila*), GUMS=Gulf Coast (*T. carolina major*) from Mississippi, GUFL=Gulf Coast from the Florida Panhandle, EA=Woodland (*T. c. carolina*), MX=Mexican (*T. mexicana mexicana*), and TT=Three-toed (*T. m. triunguis*).
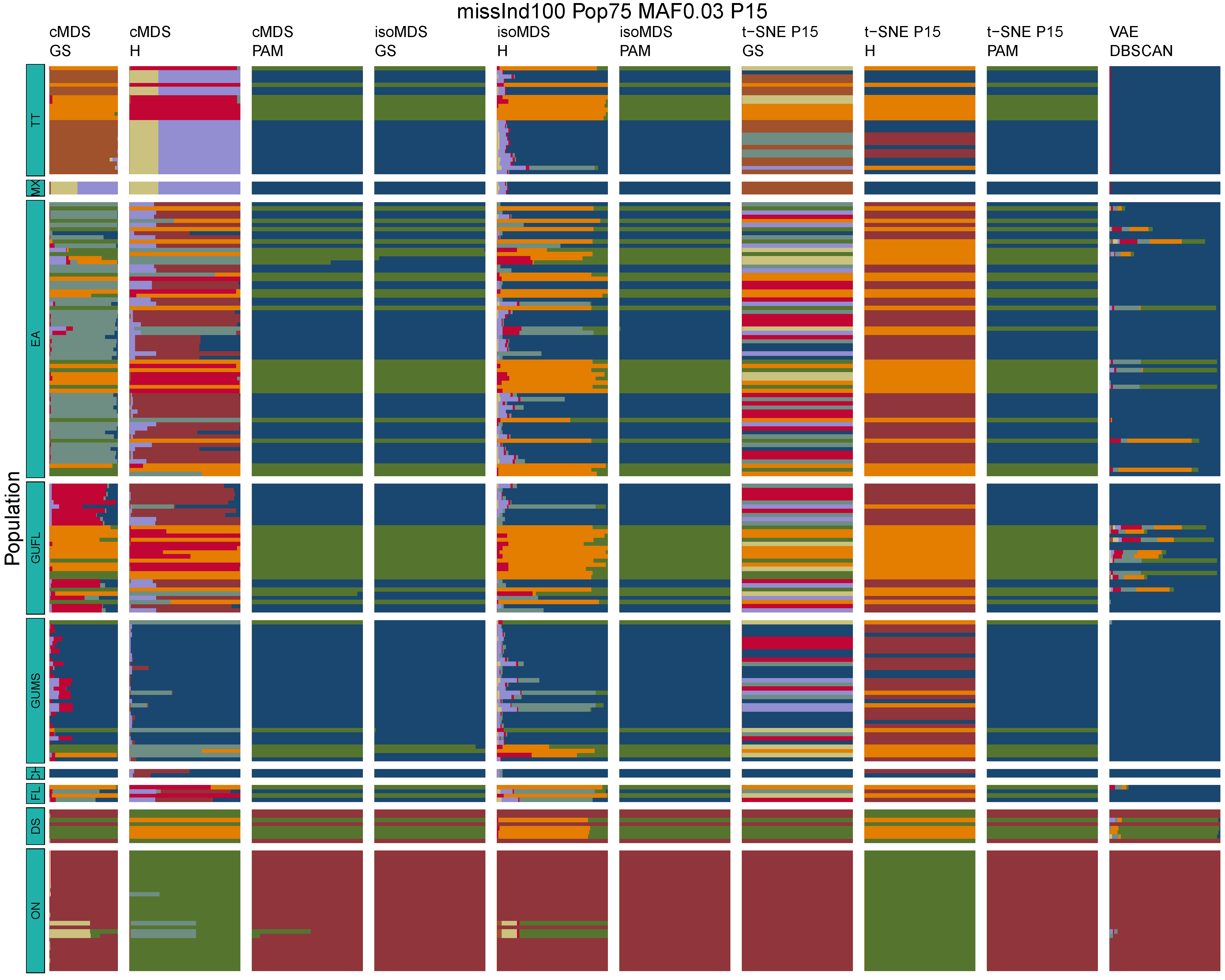

**Figure B55:** *Terrapene* unsupervised machine learning (UML) barplots depicting assignment proportions (x axis) among 100 replicates. Filtering parameters allowed maximum per-individual and per-population missing data and minimum minor allele frequency (MAF) filters. The included maximum missing data and minimum MAF filter proportions are shown in the plot title (missInd=per-individual, Pop=per-population filtering, MAF=MAF filter, P=perplexity). Each barplot on the page represents one UML algorithm: random forest, visualized with classical and isotonic multidimensional scaling (cMDS and isoMDS) ordination. t-SNE=t-distributed stochastic neighbor embedding with perplexity=15 (P15), and VAE=variational autoencoder. cMDS, isoMDS, and t-SNE were clustered with the partition around medoids (PAM) and hierarchical clustering (H) algorithms, and optimal *K* was chosen using PAM with the gap statistic (GS), H with the highest mean silhouette width (HMSW), and PAM with the HMSW. For VAE, clustering was performed, and optimal *K* chosen, with DBSCAN. *Terrapene* populations are separated by the horizontal white strips. Population IDs (blue left-aligned strip) include the ON=Ornate (*T. ornata ornata*), DS=Desert (*T. o. luteola*), FL=Florida (*T. c. bauri*), CH=Coahuilan (*T. coahuila*), GUMS=Gulf Coast (*T. carolina major*) from Mississippi, GUFL=Gulf Coast from the Florida Panhandle, EA=Woodland (*T. c. carolina*), MX=Mexican (*T. mexicana mexicana*), and TT=Three-toed (*T. m. triunguis*).
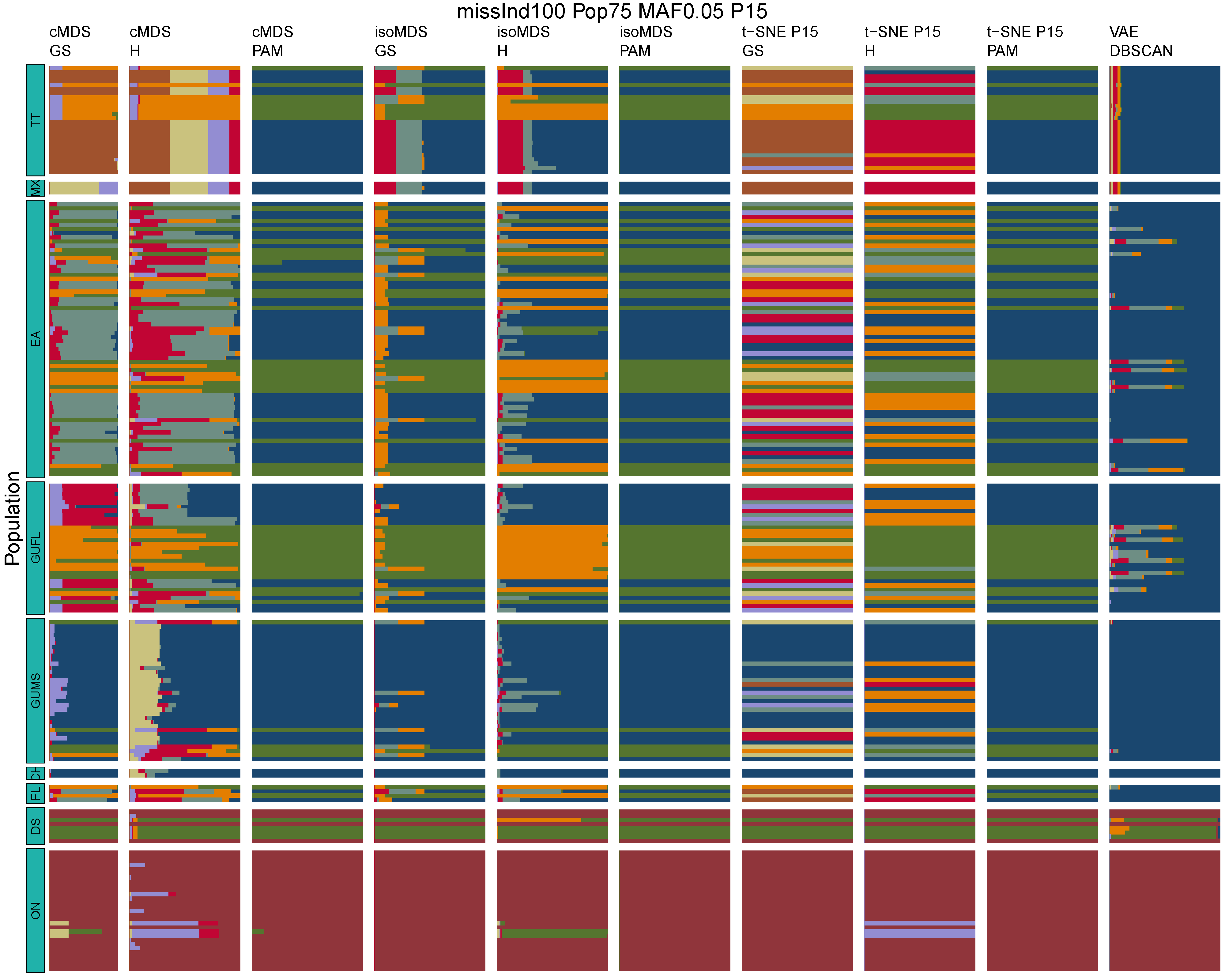

**Figure B56:** *Terrapene* unsupervised machine learning (UML) barplots depicting assignment proportions (x axis) among 100 replicates. Filtering parameters allowed maximum per-individual and per-population missing data and minimum minor allele frequency (MAF) filters. The included maximum missing data and minimum MAF filter proportions are shown in the plot title (missInd=per-individual, Pop=per-population filtering, MAF=MAF filter, P=perplexity). Each barplot on the page represents one UML algorithm: random forest, visualized with classical and isotonic multidimensional scaling (cMDS and isoMDS) ordination. t-SNE=t-distributed stochastic neighbor embedding with perplexity=15 (P15), and VAE=variational autoencoder. cMDS, isoMDS, and t-SNE were clustered with the partition around medoids (PAM) and hierarchical clustering (H) algorithms, and optimal *K* was chosen using PAM with the gap statistic (GS), H with the highest mean silhouette width (HMSW), and PAM with the HMSW. For VAE, clustering was performed, and optimal *K* chosen, with DBSCAN. *Terrapene* populations are separated by the horizontal white strips. Population IDs (blue left-aligned strip) include the ON=Ornate (*T. ornata ornata*), DS=Desert (*T. o. luteola*), FL=Florida (*T. c. bauri*), CH=Coahuilan (*T. coahuila*), GUMS=Gulf Coast (*T. carolina major*) from Mississippi, GUFL=Gulf Coast from the Florida Panhandle, EA=Woodland (*T. c. carolina*), MX=Mexican (*T. mexicana mexicana*), and TT=Three-toed (*T. m. triunguis*).
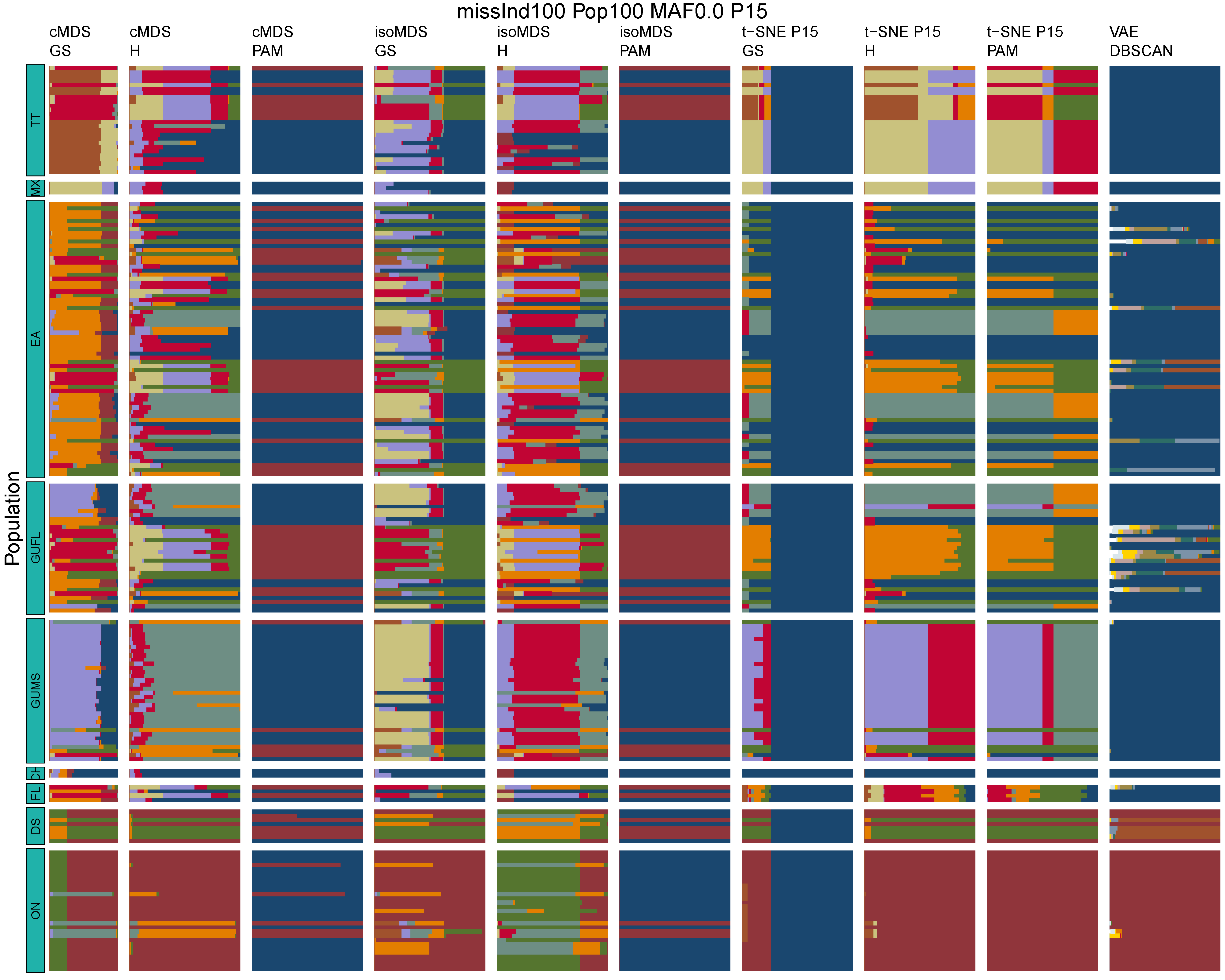

**Figure B57:** *Terrapene* unsupervised machine learning (UML) barplots depicting assignment proportions (x axis) among 100 replicates. Filtering parameters allowed maximum per-individual and per-population missing data and minimum minor allele frequency (MAF) filters. The included maximum missing data and minimum MAF filter proportions are shown in the plot title (missInd=per-individual, Pop=per-population filtering, MAF=MAF filter, P=perplexity). Each barplot on the page represents one UML algorithm: random forest, visualized with classical and isotonic multidimensional scaling (cMDS and isoMDS) ordination. t-SNE=t-distributed stochastic neighbor embedding with perplexity=15 (P15), and VAE=variational autoencoder. cMDS, isoMDS, and t-SNE were clustered with the partition around medoids (PAM) and hierarchical clustering (H) algorithms, and optimal *K* was chosen using PAM with the gap statistic (GS), H with the highest mean silhouette width (HMSW), and PAM with the HMSW. For VAE, clustering was performed, and optimal *K* chosen, with DBSCAN. *Terrapene* populations are separated by the horizontal white strips. Population IDs (blue left-aligned strip) include the ON=Ornate (*T. ornata ornata*), DS=Desert (*T. o. luteola*), FL=Florida (*T. c. bauri*), CH=Coahuilan (*T. coahuila*), GUMS=Gulf Coast (*T. carolina major*) from Mississippi, GUFL=Gulf Coast from the Florida Panhandle, EA=Woodland (*T. c. carolina*), MX=Mexican (*T. mexicana mexicana*), and TT=Three-toed (*T. m. triunguis*).
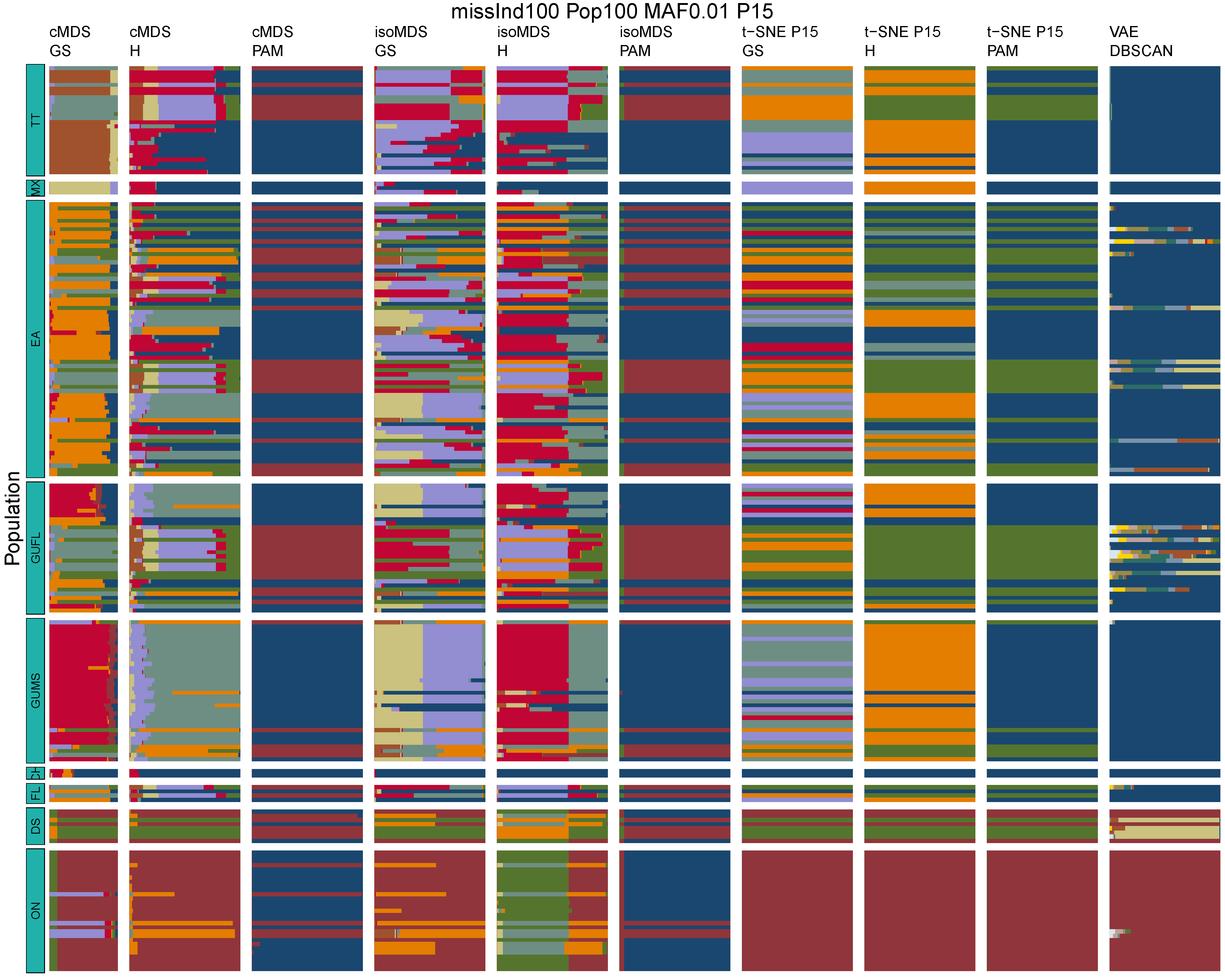

**Figure B58:** *Terrapene* unsupervised machine learning (UML) barplots depicting assignment proportions (x axis) among 100 replicates. Filtering parameters allowed maximum per-individual and per-population missing data and minimum minor allele frequency (MAF) filters. The included maximum missing data and minimum MAF filter proportions are shown in the plot title (missInd=per-individual, Pop=per-population filtering, MAF=MAF filter, P=perplexity). Each barplot on the page represents one UML algorithm: random forest, visualized with classical and isotonic multidimensional scaling (cMDS and isoMDS) ordination. t-SNE=t-distributed stochastic neighbor embedding with perplexity=15 (P15), and VAE=variational autoencoder. cMDS, isoMDS, and t-SNE were clustered with the partition around medoids (PAM) and hierarchical clustering (H) algorithms, and optimal *K* was chosen using PAM with the gap statistic (GS), H with the highest mean silhouette width (HMSW), and PAM with the HMSW. For VAE, clustering was performed, and optimal *K* chosen, with DBSCAN. *Terrapene* populations are separated by the horizontal white strips. Population IDs (blue left-aligned strip) include the ON=Ornate (*T. ornata ornata*), DS=Desert (*T. o. luteola*), FL=Florida (*T. c. bauri*), CH=Coahuilan (*T. coahuila*), GUMS=Gulf Coast (*T. carolina major*) from Mississippi, GUFL=Gulf Coast from the Florida Panhandle, EA=Woodland (*T. c. carolina*), MX=Mexican (*T. mexicana mexicana*), and TT=Three-toed (*T. m. triunguis*).
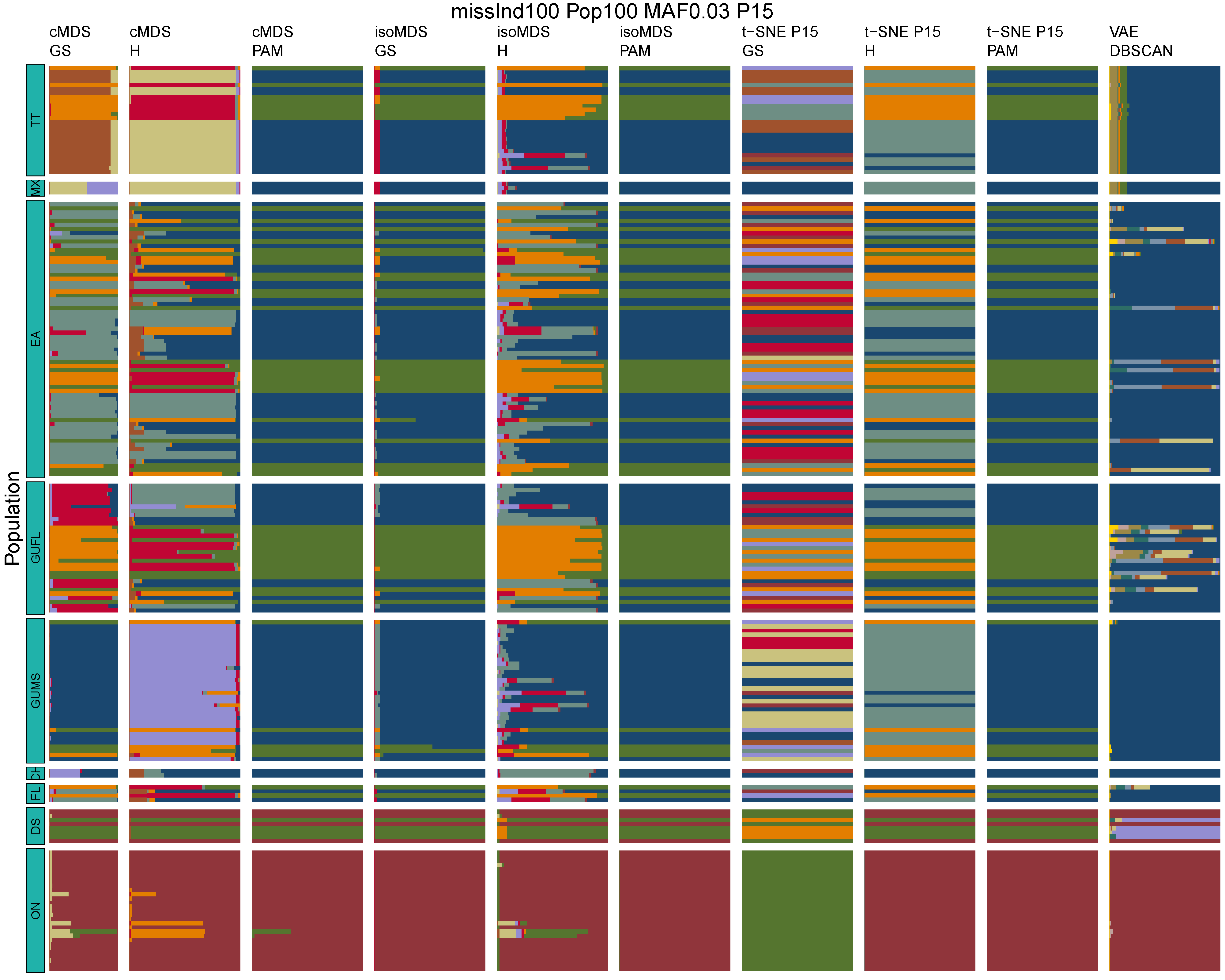

**Figure B59:** *Terrapene* unsupervised machine learning (UML) barplots depicting assignment proportions (x axis) among 100 replicates. Filtering parameters allowed maximum per-individual and per-population missing data and minimum minor allele frequency (MAF) filters. The included maximum missing data and minimum MAF filter proportions are shown in the plot title (missInd=per-individual, Pop=per-population filtering, MAF=MAF filter, P=perplexity). Each barplot on the page represents one UML algorithm: random forest, visualized with classical and isotonic multidimensional scaling (cMDS and isoMDS) ordination. t-SNE=t-distributed stochastic neighbor embedding with perplexity=15 (P15), and VAE=variational autoencoder. cMDS, isoMDS, and t-SNE were clustered with the partition around medoids (PAM) and hierarchical clustering (H) algorithms, and optimal *K* was chosen using PAM with the gap statistic (GS), H with the highest mean silhouette width (HMSW), and PAM with the HMSW. For VAE, clustering was performed, and optimal *K* chosen, with DBSCAN. *Terrapene* populations are separated by the horizontal white strips. Population IDs (blue left-aligned strip) include the ON=Ornate (*T. ornata ornata*), DS=Desert (*T. o. luteola*), FL=Florida (*T. c. bauri*), CH=Coahuilan (*T. coahuila*), GUMS=Gulf Coast (*T. carolina major*) from Mississippi, GUFL=Gulf Coast from the Florida Panhandle, EA=Woodland (*T. c. carolina*), MX=Mexican (*T. mexicana mexicana*), and TT=Three-toed (*T. m. triunguis*).
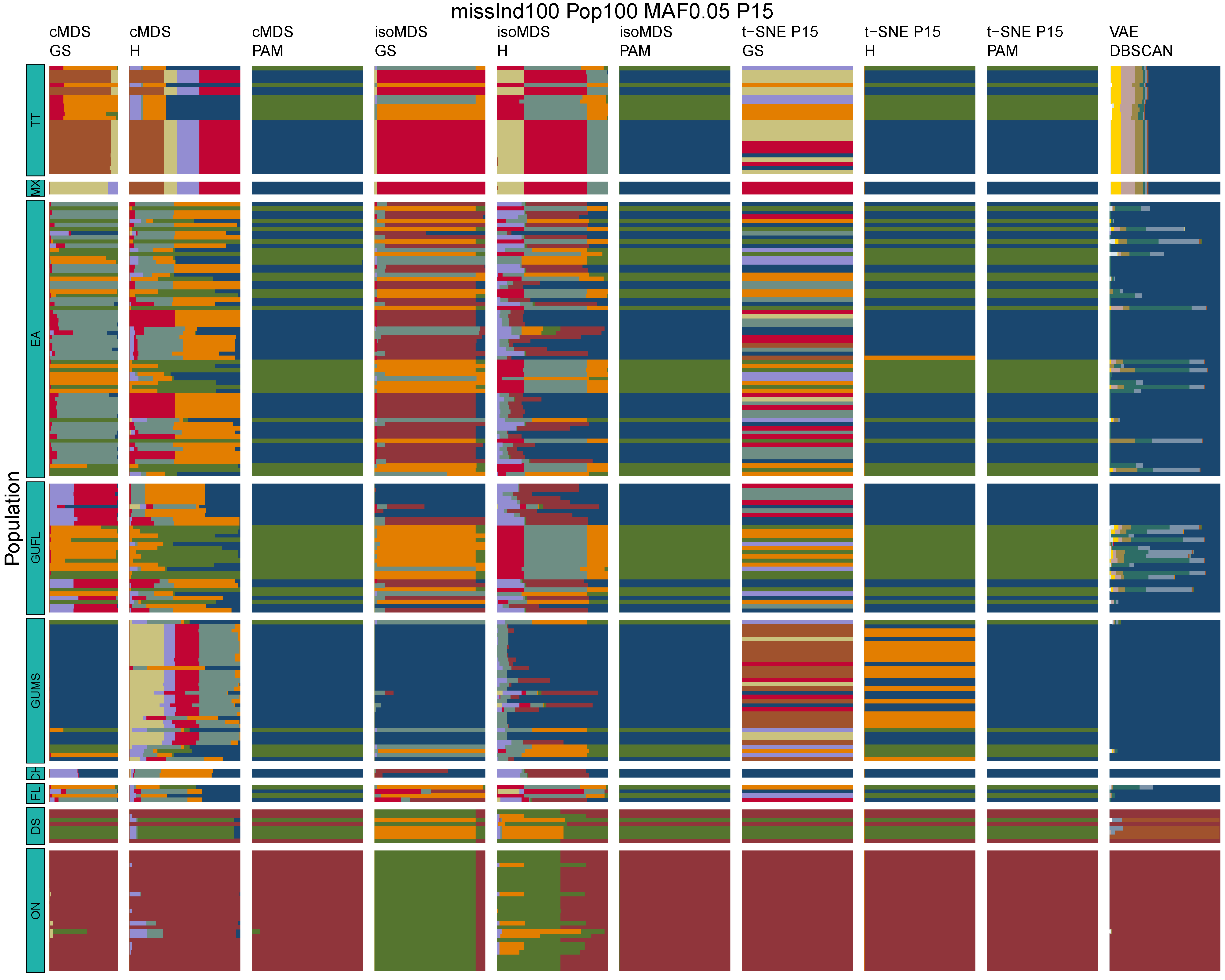

**Figure B60:** *Terrapene* unsupervised machine learning (UML) barplots depicting assignment proportions (x axis) among 100 replicates. Filtering parameters allowed maximum per-individual and per-population missing data and minimum minor allele frequency (MAF) filters. The included maximum missing data and minimum MAF filter proportions are shown in the plot title (missInd=per-individual, Pop=per-population filtering, MAF=MAF filter, P=perplexity). Each barplot on the page represents one UML algorithm: random forest, visualized with classical and isotonic multidimensional scaling (cMDS and isoMDS) ordination. t-SNE=t-distributed stochastic neighbor embedding with perplexity=15 (P15), and VAE=variational autoencoder. cMDS, isoMDS, and t-SNE were clustered with the partition around medoids (PAM) and hierarchical clustering (H) algorithms, and optimal *K* was chosen using PAM with the gap statistic (GS), H with the highest mean silhouette width (HMSW), and PAM with the HMSW. For VAE, clustering was performed, and optimal *K* chosen, with DBSCAN. *Terrapene* populations are separated by the horizontal white strips. Population IDs (blue left-aligned strip) include the ON=Ornate (*T. ornata ornata*), DS=Desert (*T. o. luteola*), FL=Florida (*T. c. bauri*), CH=Coahuilan (*T. coahuila*), GUMS=Gulf Coast (*T. carolina major*) from Mississippi, GUFL=Gulf Coast from the Florida Panhandle, EA=Woodland (*T. c. carolina*), MX=Mexican (*T. mexicana mexicana*), and TT=Three-toed (*T. m. triunguis*).
