## Supplementary Information S Tables and Figures for "The choices we make and the impacts they have: Machine learning and species delimitation in North American box turtles (*Terrapene* spp.)"

**Table S2:** Summary statistics from twenty BFD* models (Bayes Factor Delimitation, *with genomic data) among 37 North American box turtle (*Terrapene* spp.) samples and 179 unlinked ddRADseq single nucleotide polymorphism (SNP) variants. The standard deviations reflect error in the calculation of the marginal likelihood estimate (MLE) from ten path sampling cross-validation runs.

| **BFD Model** | **ESS Mean** | **ESS Median (Min, Max)** | **MLE** | **Std. Dev.** |
| --- | --- | --- | --- | --- |
| run1, East/West | 649.18 | 506.72755 (122.6519, 1334) | -2926.56 | 0.06 |
| run2, EA+FL+CH+GUMS+GUFL+TT+MX | 515.20 | 422.0864 (105.0803, 1289.1458) | -2926.20 | 0.08 |
| run3, ON+DS/EA+TT+MX+CH+GUMS+GUFL/FL | 593.63 | 459.24485 (71.6897, 1334) | -2720.23 | 0.07 |
| run4, EA+FL+CH+GUMS+GUFL+TT | 487.48 | 372.5475 (21.7602, 1264.0155) | -2800.56 | 0.09 |
| run5, EA+CH+GUMS+GUFL+TT+MX | 493.91 | 492.88135 (56.1857, 1115.7808) | -2719.02 | 0.08 |
| run6, EA+FL+CH+GUMS+GUFL | 484.56 | 433.14245 (74.3832, 1236.2148) | -2693.37 | 0.10 |
| run7, EA+CH+GUMS+GUFL+MX | 478.60 | 380.1236 (68.1691, 1153.8154) | -2657.48 | 0.09 |
| run8, EA+CH+GUMS+GUFL+TT | 455.44 | 379.51645 (28.2256, 1217.8904) | -2607.72 | 0.10 |
| run9, GUMS+GUFL+CH+EA | 420.68 | 369.88705 (28.1414, 1295.5895) | -2514.86 | 0.11 |
| run10, EA+FL+GUFL/CH+GUMS | 407.49 | 322.56155 (38.1882, 1181.6271) | -2594.91 | 0.11 |
| run11, GUMS+GUFL+CH | 367.29 | 305.75535 (62.5265, 1097.3091) | -2489.62 | 0.12 |
| run12, GUMS+CH/GUFL+EA | 340.61 | 285.44545 (42.0809, 968.308) | -2461.28 | 0.13 |
| run13, EA+FL/CH+GUMS | 365.06 | 307.52505 (73.946, 913.1602) | -2555.16 | 0.13 |
| run14, EA+FL+GUFL | 321.74 | 251.91375 (31.6129, 1127.1272) | -2552.22 | 0.11 |
| run15, GUMS+GUFL | 288.79 | 274.3423 (59.7872, 815.3268) | -2427.58 | 0.12 |
| run16, GUMS+CH | 330.86 | 261.6958 (68.1546, 880.6262) | -2448.61 | 0.13 |
| run17, EA+GUFL | 314.11 | 212.32105 (49.5533, 1158.6918) | -2417.84 | 0.12 |
| run18, EA+FL | 343.54 | 299.22845 (28.5148, 1291.1445) | -2511.83 | 0.14 |
| run19, DS+ON | 386.57 | 265.64725 (54.6711, 1181.5824) | -2404.34 | 0.12 |
| run20, All Separate | 305.72 | 223.09945 (81.8985, 1066.5475) | -2403.39 | 0.13 |

ESS = Effective sample size

MLE = Marginal likelihood estimate

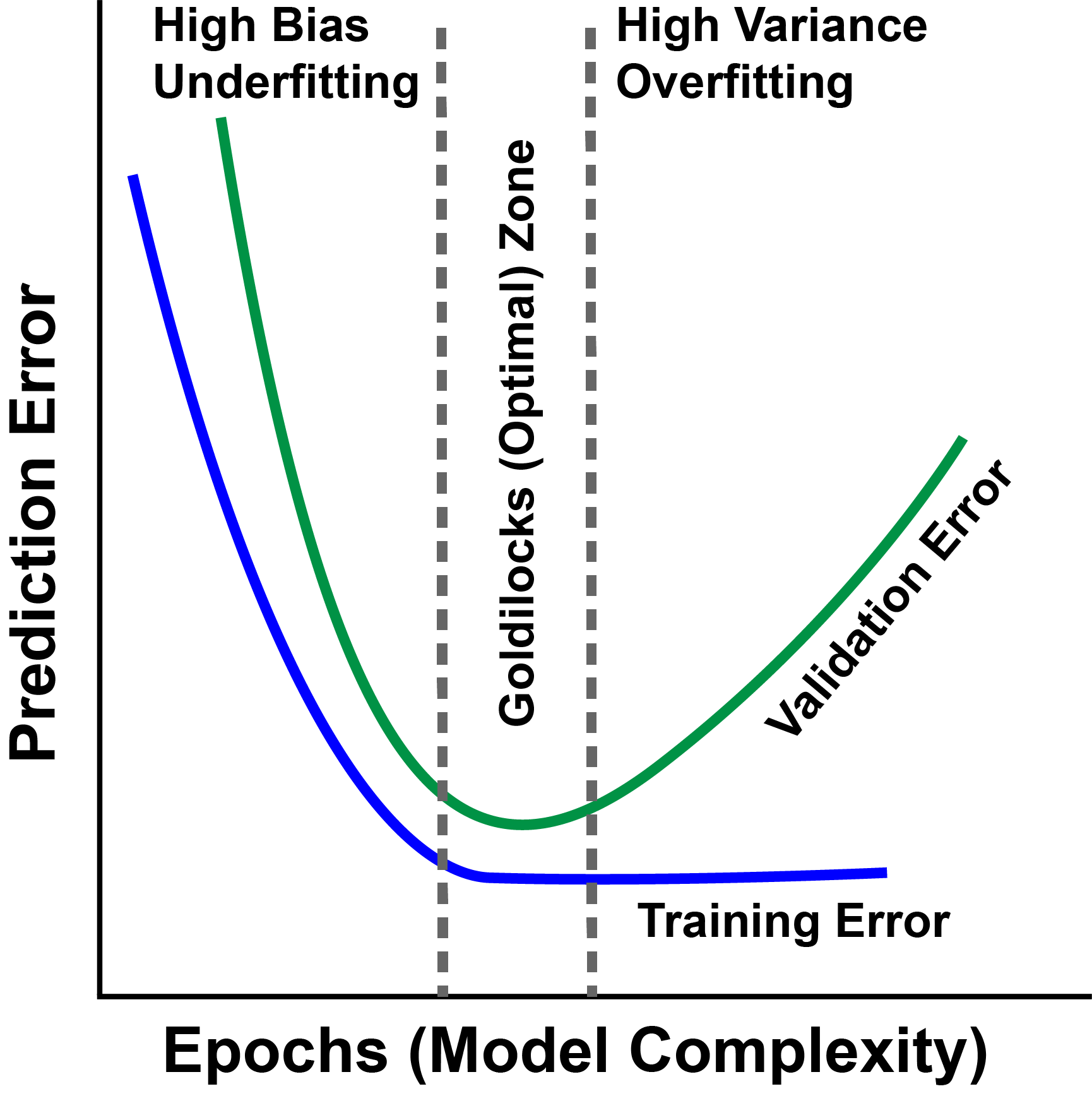

**Figure S1:** Prediction error *versus* model complexity for machine learning. Ideally, training should stop when the validation and training prediction error are at their lowest point [‘Goldilocks zone’, indicated by gray dashed lines; (Al’Aref *et al.* 2019)]. High bias and underfitting occur when training ends prior to reaching the Goldilocks zone, and high variance and overfitting occur when validation error begins to climb.

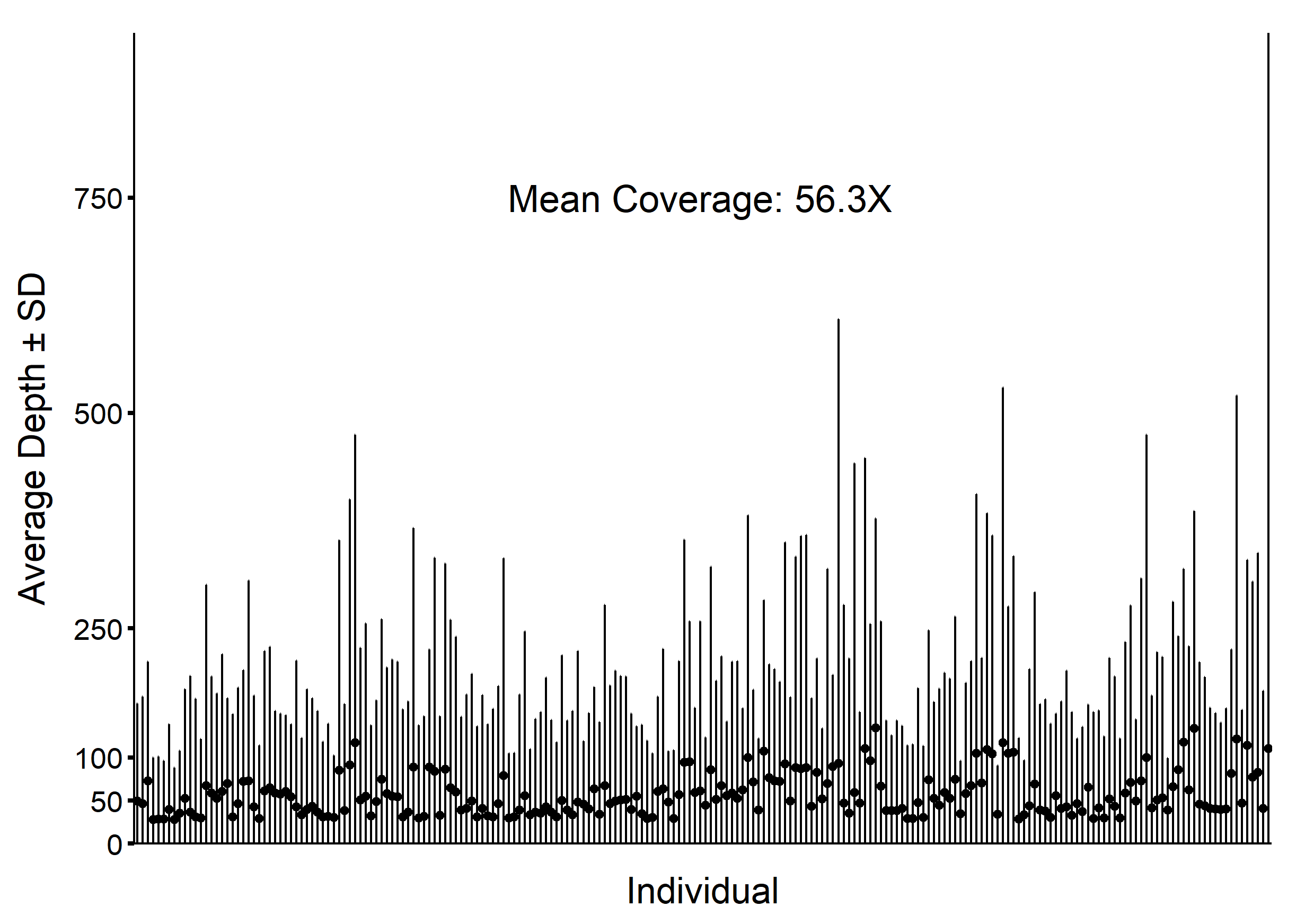

**Figure S2:** Mean per-individual read depth (points) ± standard deviation (gray bars) for 214 North American box turtle (*Terrapene* spp.) ddRAD sequencing samples.

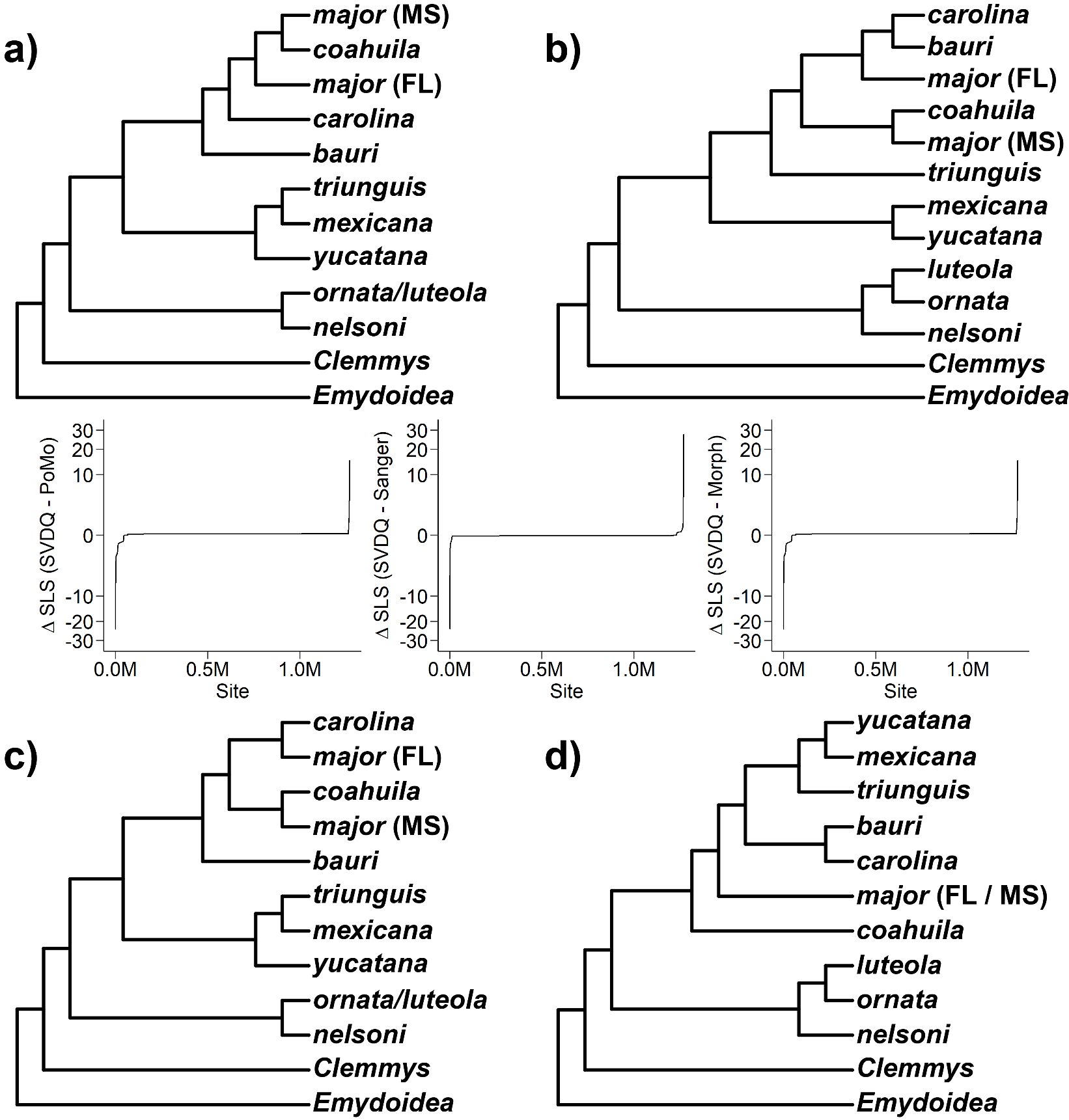

**Figure S3:** *Terrapene* constraint trees and respective changes in site-likelihood scores (**Δ**SLS) representing genome-wide support for the a) SVDquartets, b) PoMo, c) Sanger, and d) Morphological phylogenetic hypotheses. The SVDquartets and PoMo trees are derived in this study, whereas the Sanger and morphological hypotheses are results previously published (Minx 1996; Martin *et al.* 2013).

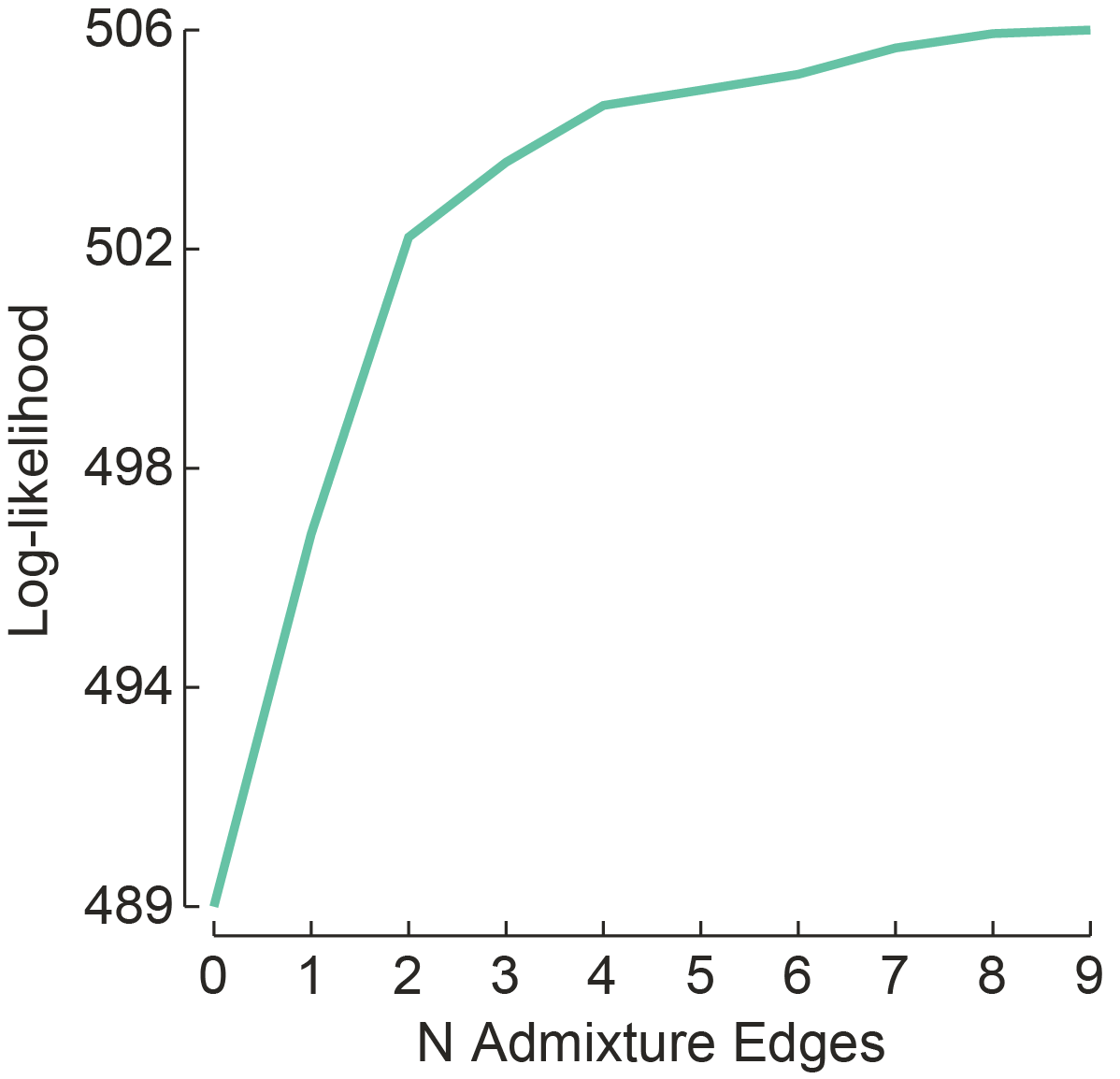

**Figure S4:** Log-likelihoods for the number (N) of TreeMix admixture edges.

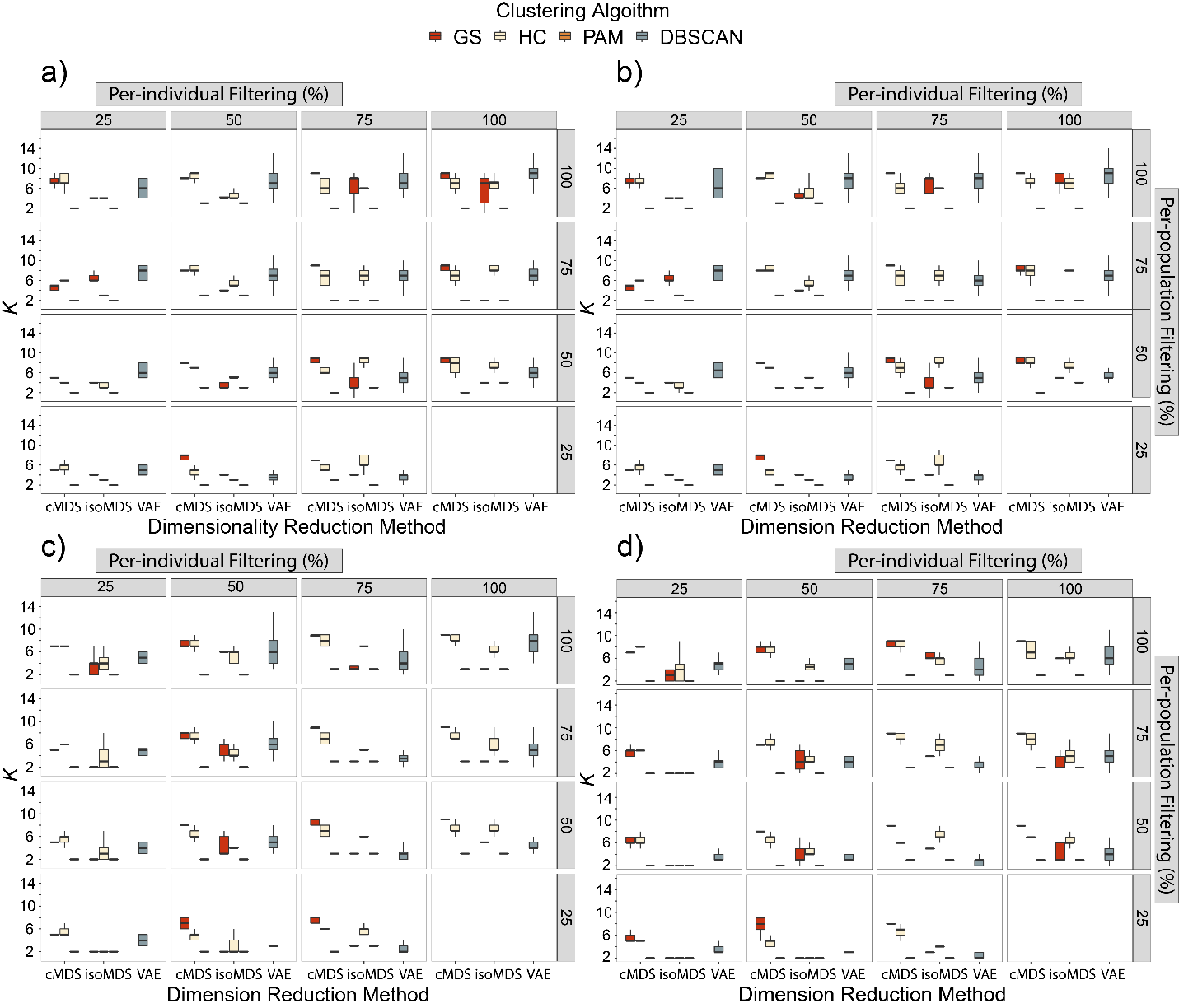

**Figure S5:** Tukey-style box and whiskers plots for 100 unsupervised machine learning (UML) species delimitation replicates. Data were filtered using per-individual (panel columns) and per-population (panel rows) missing data filters (25%=most stringent; 100%=no filtering) and a) 0%, b) 1%, c) 3%, and d) 5% minor allele frequency filters. Black bars on the boxplots indicate the median, lower and upper hinges represent the 25^th^ and 75^th^ percentiles, and the whisker range includes 1.5 times the inter-quartile range (IQR) past the hinges. cMDS and isoMDS = random forest classification visualized with classical and isotonic multidimensional scaling; VAE=variational autoencoder. Optimal *K* among cMDS and isoMDS was determined in three ways among two clustering algorithms: partition around medoids (PAM) with the gap statistic (GS), hierarchical clustering with the highest mean silhouette width (HMSW), and PAM with HMSW. Optimal *K* for VAE was determined using DBSCAN.

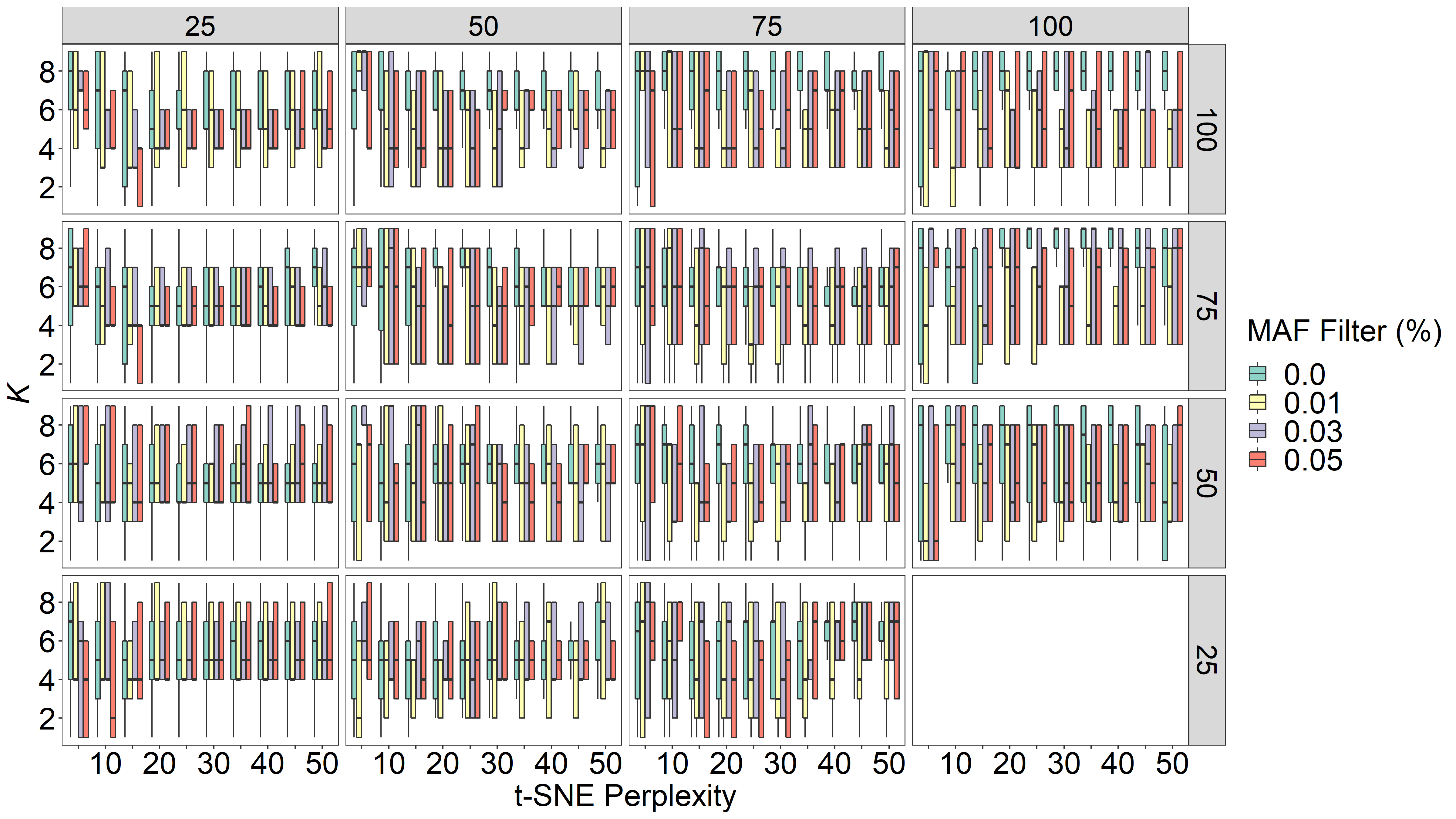

**Figure S6:** Tukey-style box and whiskers plots depicting variation in mean optimal K among a t-SNE perplexity grid search (x axes). Data were filtered with 25%, 50%, 75%, and 100% (no filtering) per-individual (panel columns) and per-population (panel rows) missing data filters and 0%, 1%, 3%, and 5% minor allele frequency (MAF) filters. Black bars on the boxplots indicate the median, lower and upper hinges represent the 25^th^ and 75^th^ percentiles, and the whisker range includes 1.5 times the inter-quartile range (IQR) past the hinges. t-SNE = t-distributed stochastic neighbor embedding.

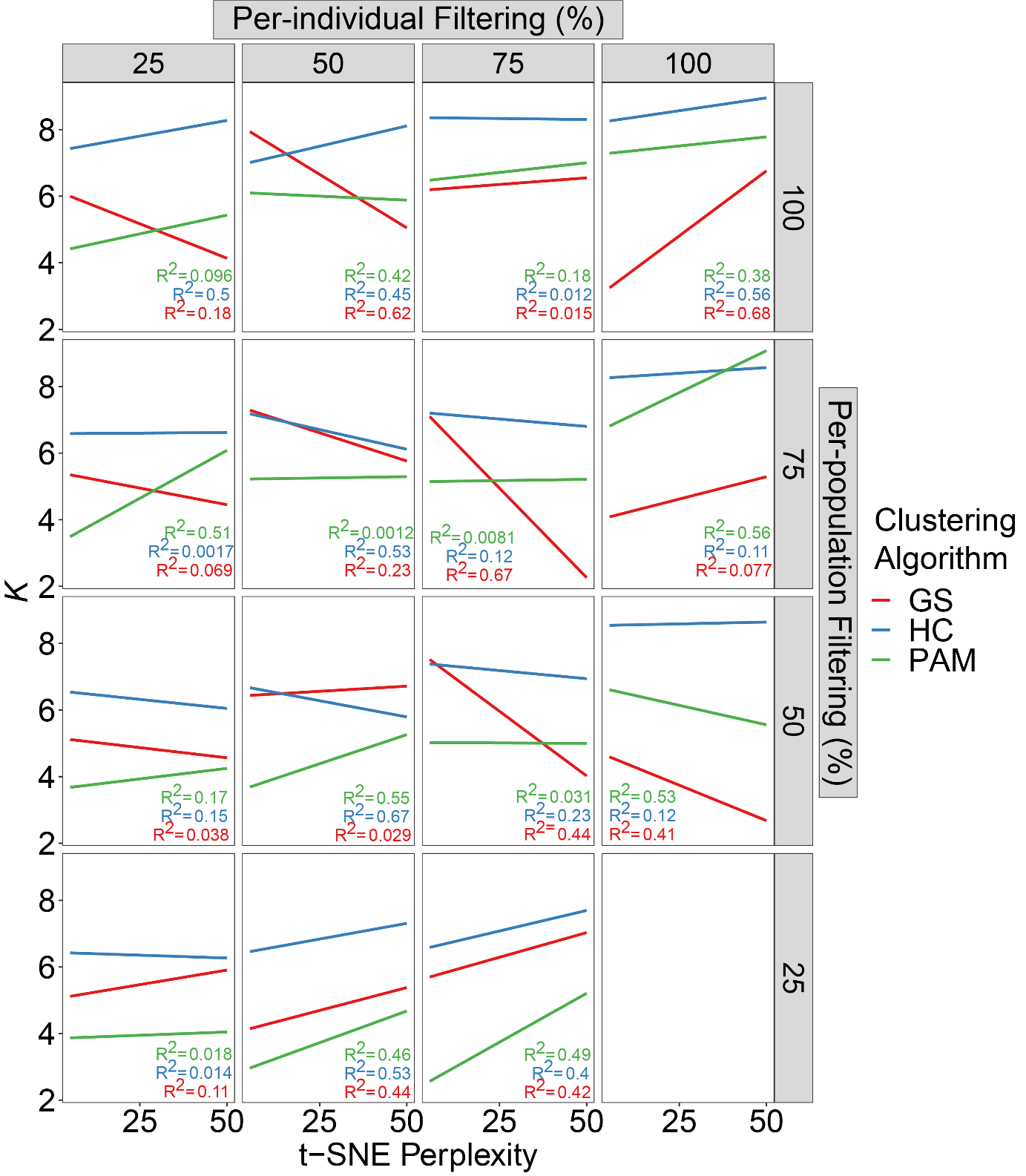

**Figure S7:** Regressions showing mean optimal *K* among ten t-SNE perplexity settings and three clustering algorithms for a minor allele frequency (MAF) filter=0% (no filtering). Panel columns and rows represent per-individual and per-population missing data filters, respectively. Mean optimal *K* was chosen using three clustering algorithms (fill colors): 1) Partition around medoids (PAM) with the gap statistic (GS), hierarchical clustering (HC) with the highest mean silhouette width (HMSW), and PAM with HMSW. R^2^ values show correlations per clustering algorithm.

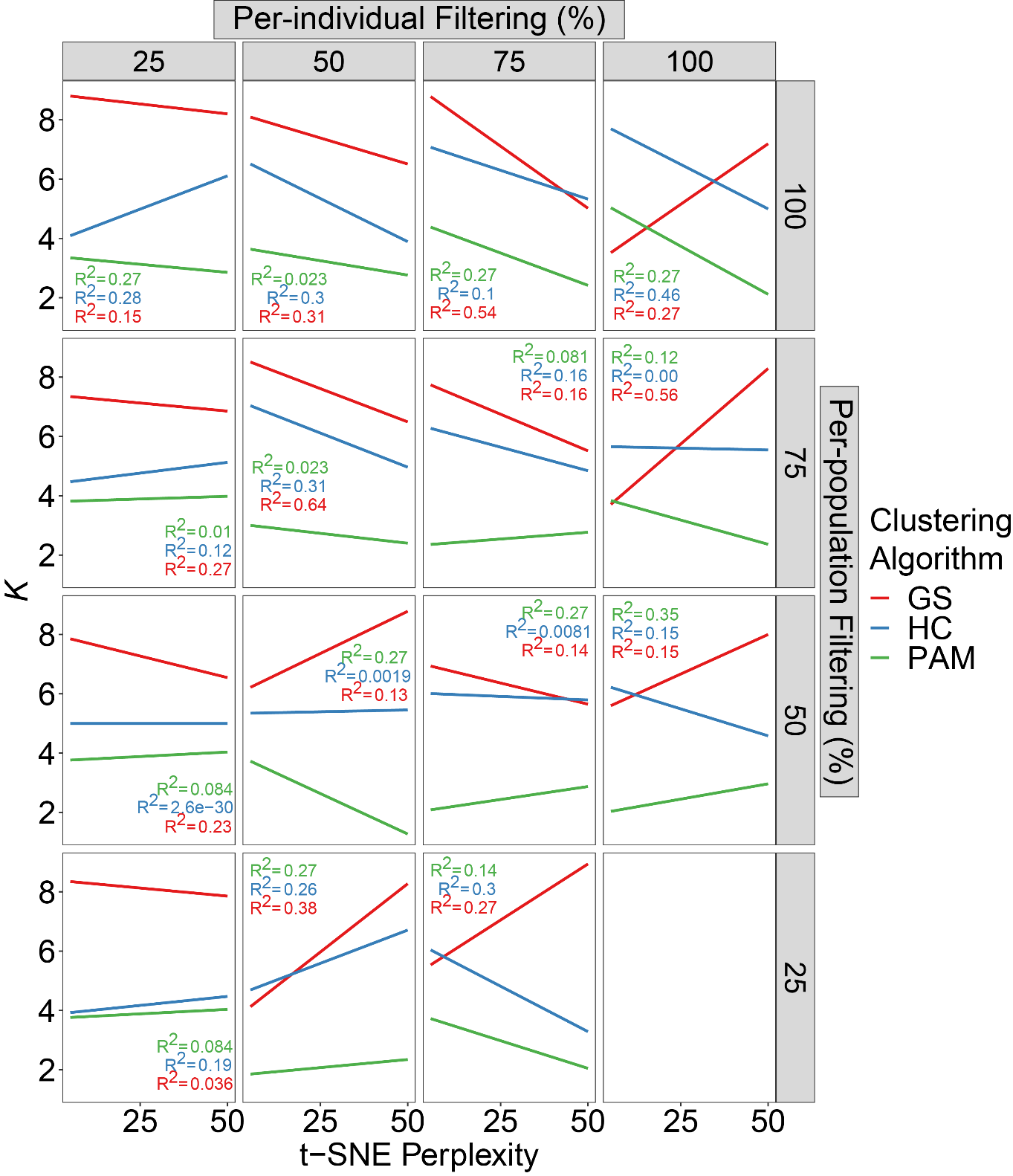

**Figure S8:** Regressions showing mean optimal *K* among ten t-SNE perplexity settings and three clustering algorithms for a minor allele frequency (MAF) filter=1%. Panel columns and rows represent per-individual and per-population missing data filters, respectively. Mean optimal *K* was chosen using three clustering algorithms (fill colors): 1) Partition around medoids (PAM) with the gap statistic (GS), hierarchical clustering (HC) with the highest mean silhouette width (HMSW), and PAM with HMSW. R^2^ values show correlations per clustering algorithm.

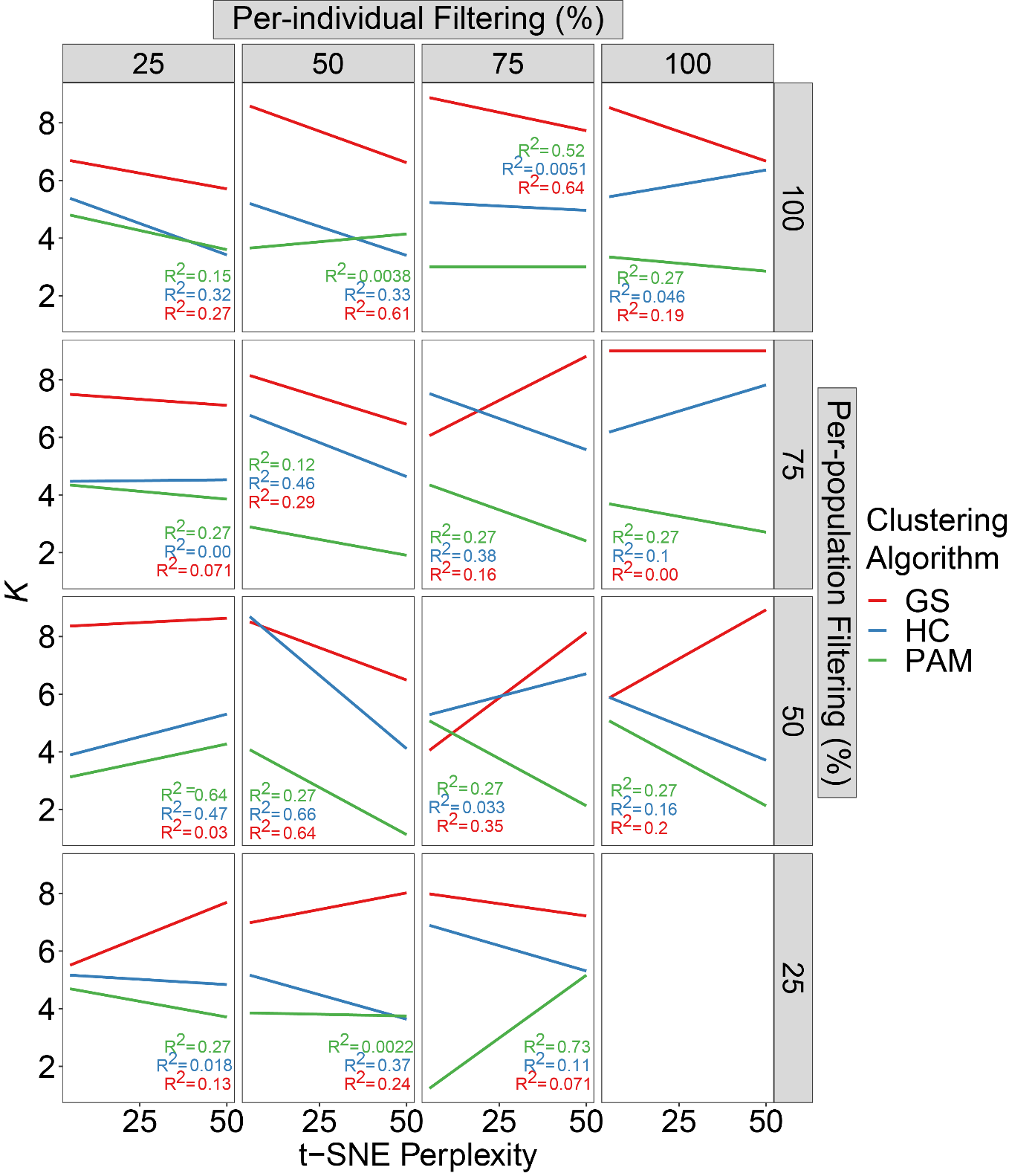

**Figure S9:** Regressions showing mean optimal *K* among ten t-SNE perplexity settings and three clustering algorithms for a minor allele frequency (MAF) filter=3%. Panel columns and rows represent per-individual and per-population missing data filters, respectively. Mean optimal *K* was chosen using three clustering algorithms (fill colors): 1) Partition around medoids (PAM) with the gap statistic (GS), hierarchical clustering (HC) with the highest mean silhouette width (HMSW), and PAM with HMSW. R^2^ values show correlations per clustering algorithm.

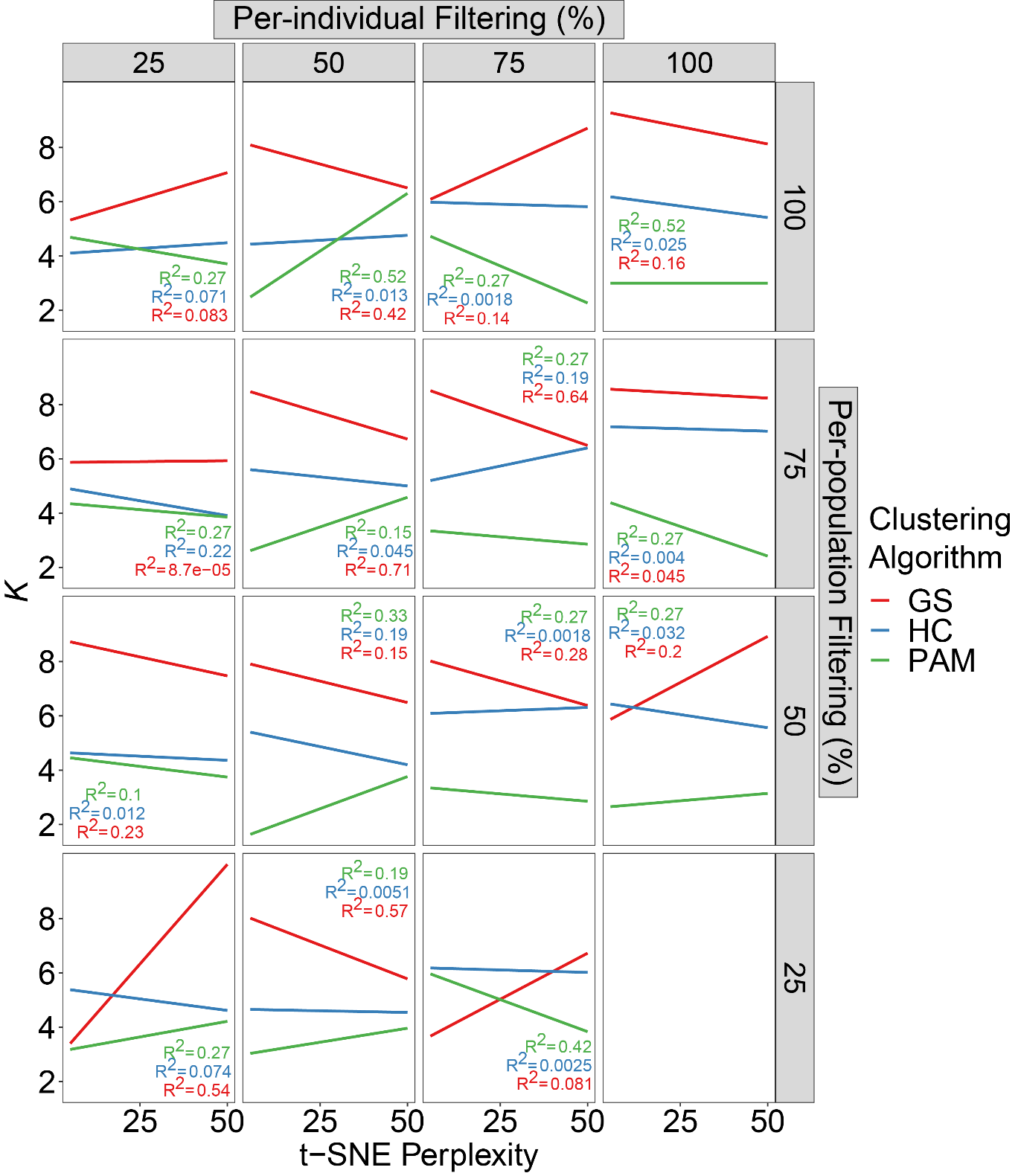

**Figure S10:** Regressions showing mean optimal *K* among ten t-SNE perplexity settings and three clustering algorithms for a minor allele frequency (MAF) filter=5%. Panel columns and rows represent per-individual and per-population missing data filters, respectively. Mean optimal *K* was chosen using three clustering algorithms (fill colors): 1) Partition around medoids (PAM) with the gap statistic (GS), hierarchical clustering (HC) with the highest mean silhouette width (HMSW), and PAM with HMSW. R^2^ values show correlations per clustering algorithm.
